# Metabolic engineering and late-stage functionalization expand the chemical space of the antimalarial premarineosin A

**DOI:** 10.1101/2025.01.28.635342

**Authors:** Christina M. McBride, Morgan McCauley, Natalia R. Harris, Sahar Amin, Brian J. Curtis, Linnea Verhey-Henke, Awet A. Teklemichael, Erin N. Oliphant, Patricia Dranchak, Katherine L. Lev, Fengrui Qu, Harrison M. Snodgrass, Jared C. Lewis, James Inglese, Xin-zhuan Su, Filipa Pereira, David H. Sherman

## Abstract

Diversification of structurally complex natural products remains a key challenge in the discovery of next-generation therapeutics. Premarineosin A, a potent and selective antimalarial natural product, is a promising yet underexplored scaffold due to limited availability and synthetic complexity. In this work, we overcome both barriers by coupling metabolic engineering with late-stage derivatization, enabling the first systematic exploration of the premarineosin A scaffold. Rational engineering of *Streptomyces eitanensis*, encoding a (−)-premarineosin A biosynthetic gene cluster, increased titers over 200-fold. Sustainable production of (−)-premarineosin A enabled a unique semi-synthetic and biocatalytic derivatization campaign. In this first structure-activity relationship study of premarineosin A, we accessed a suite of novel analogs, including a C12-brominated derivative with nanomolar potency (EC50 < 5 nM). This work establishes (−)-premarineosin A as a tractable and evolvable antimalarial scaffold, demonstrating how chemical biology approaches can unlock new structural and pharmacological space from complex microbial metabolites.

## Introduction

Malaria is a life-threatening disease that poses severe health risks for nearly half of the global population. In 2023 alone, malaria was responsible for an estimated 263 million cases and 597,000 deaths^1^. As the causative *Plasmodium* parasites are becoming increasingly drug resistant, developing new antimalarials that match or exceed the efficacy of current therapeutics is paramount^2–4^. The prodiginine class of microbial natural products, distinguished by a core tripyrrole moiety^5^, has been explored for potent antiparasitic activity and selectivity^6–11^. In addition to their antimalarial activity^6–9,12^, these metabolites have exhibited remarkable anticancer^13–15^, antibacterial^16,17^, antifungal^17^, and immunosuppressant^14^ properties. Cyclic prodiginines are particularly notable, as their constrained conformations encourage the binding of charged ions, leading to enhanced bioactivity compared to their linear counterparts^15,18,19^. For instance, marineosin A, a cyclic prodiginine isolated from *Streptomyces* CNQ-617, incorporates a unique spiroaminal structure that confers potent cytotoxicity^20^. In a key late-stage step in marineosin A biosynthesis, the linear 23-hydroxyundecylprodiginine (23-HUP, **1**) undergoes an oxidative bicyclization at the C8-C9 double bond catalyzed by a Rieske oxygenase (MarG), yielding premarineosin A^12^. Finally, premarineosin A is converted to marineosin A by a proposed MarA-catalyzed reduction of the C6-C7 double bond^12,20^. Premarineosin A is of particular interest, as it exhibits single-digit nanomolar antimalarial activity *in vitro* and is notably less cytotoxic than marineosin A^12^.

Derivatization is an effective strategy for enhancing the antiplasmodial activity and selectivity of prodiginine natural products^6–11^, as functionalization can improve potency, reduce toxicity, optimize metabolism, or increase oral bioavailability in comparison to the native scaffold^21–25^. Most often, total synthesis has been employed to derivatize linear prodiginines, with modifications occurring early in the route through the functionalization of simple precursors^8^. However, because of the synthetic complexity of the spirocyclic core, premarineosin A is significantly less amenable to total synthesis, as evidenced by all prior efforts resulting in low overall yields^26,27^. Synthesis of premarineosin A analogs has been attempted via bioconversion of 23-HUP, but was limited by low conversion rates^6^. Late-stage functionalization is a promising alternative strategy, but can be challenging for natural product scaffolds due to their complex ring structures and diverse functional groups, which can hinder selectivity and reactive site accessibility^28,29^. Hence, identifying late-stage diversification approaches that selectively target sites like the highly reactive pyrrolic carbons would enable rapid functionalization of premarineosin A.

Securing access to premarineosin A is also essential to our diversification efforts. While heterologous expression of the marineosin biosynthetic cluster disrupted in *marA* enabled the isolation of 0.5 mg of premarineosin A^12^, no production was observed in cultures >100 mL, limiting further structural characterization, derivatization, and drug development efforts. Metabolic engineering is a proven approach for enhancing the production of natural products^30,31^, including prodiginines^32–36^. In this work, we report a previously unexplored (−)-premarineosin A (**2**) (*pma*) biosynthetic gene cluster (BGC) in *Streptomyces eitanensis* and employed metabolic engineering approaches to improve its production. Access to a sustainable source of **2** enabled us to define its absolute stereochemistry and develop high-yield strategies for late-stage derivatization of (−)-premarineosin A via unique semi-synthetic and biocatalytic approaches. The new (−)-premarineosin A derivatives were investigated for potency and cytotoxicity *in vitro*, which identified a novel brominated analog with high potency against drug-sensitive and drug-resistant *Plasmodium falciparum* parasites.

## Results

### Genome and metabolome analysis identified a (−)-premarineosin A biosynthetic gene cluster in *S. eitanensis*

Recently, we identified 23-HUP in *Streptomyces eitanensis*^30^. A detailed review of its genome revealed a biosynthetic gene cluster (BGC) with 90% similarity to the canonical marineosin BGC (MIBiG: BGC0000091)^12,37^. This newly identified cluster does not encode the putative acyltransferase (MarE) and the reductase (MarA) needed to generate the 1-pyrroline B-ring in marineosin biosynthesis (**Fig. 1a**)^12^. However, it does encode the putative Rieske oxygenase PmaG, a homolog (85% similarity, 77% identity) of the MarG enzyme that catalyzes bicyclization of 23-HUP into the marineosin precursor, premarineosin A (**Fig. 1b, c**). Closer inspection of the *S. eitanensis* metabolome detected a compound with the same molecular weight and fragmentation pattern as premarineosin A (**Supplementary Fig. 1**). As marineosin A production was not observed in this strain, we reasoned that the identified BGC (*pma* BGC) likely encodes premarineosin A as the terminal product, suggesting that *S. eitanensis* could serve as an ideal chassis strain for optimizing its large-scale production.

**Figure 1.**
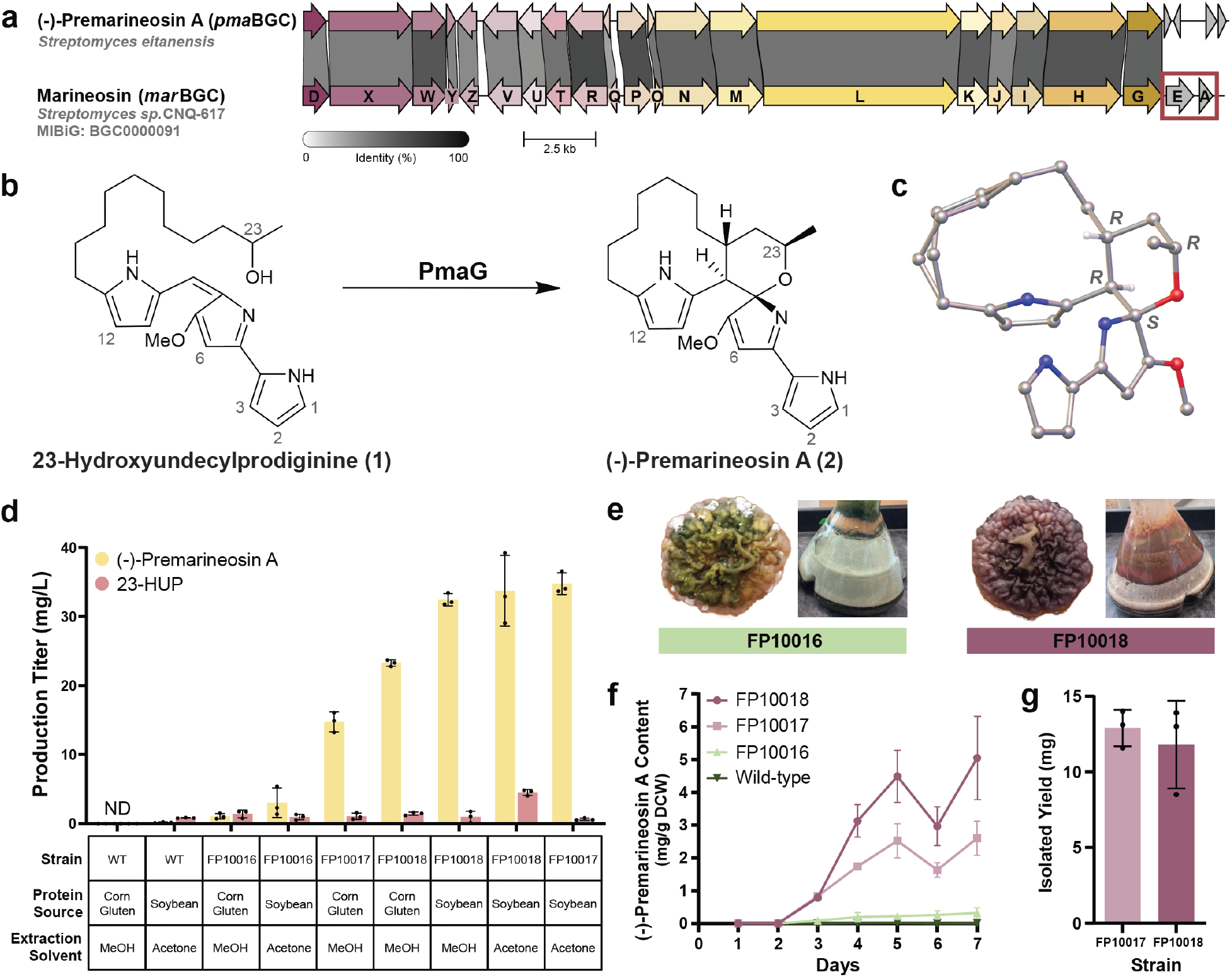
Metabolic engineering of *S. eitanensis* enhanced (−)-premarineosin A production. **a**. Comparison of the *S. eitanensis* (−)-premarineosin A BGC and the *Streptomyces sp*. CNQ617 marineosin BGC using clinker^43^. The (−)-premarineosin A BGC lacks genes with homology to *marA* and *marE* (red box). **b**. The final biosynthetic step of the (−)-premarineosin A pathway. **c**. Crystal structure of (−)-premarineosin A. **d**. (−)-Premarineosin A (**2**) and 23-HUP (**1**) production (mg/L), quantified by AUC (HPLC) against standard curves of each compound. Dots represent three independent culture replicates. Error bars indicate standard deviation (SD). ND: Not Detected, the measured production was below the limit of detection. **e**. Colonies of engineered FP10016 (green) and FP10018 (red) strains cultivated in solid GICYE. **f**. (−)-Premarineosin A production (mg) per gram of dry cell weight (DCW, g) over time by wild-type and engineered FP10016, FP10017 and FP10018 strains. Error bars indicate standard deviation (SD) for n=3 replicates. **g**. Isolated titers of pure (−)-premarineosin A from the engineered FP10017 and FP10018 strains. Dots represent three independent culture replicates. Error bars indicate standard deviation (SD).

### Overexpression of the *pma* cluster-situated regulator (*pmaD*) and Rieske oxygenase (*pmaG)* improved (−)-premarineosin A production in *S. eitanensis*

Trace amounts of (−)-premarineosin A were detected in wild-type *S. eitanensis* (**Fig. 1d, Supplementary Fig. 1**). Annotation of the *pma* BGC (**Supplementary Table 1**) revealed PmaD, a homolog (40% protein identity, 52% similarity) of the RedD transcriptional activator from the undecylprodigiosin (UDP) pathway in *Streptomyces coelicolor* A3(2)^38–40^, suggesting that PmaD may similarly regulate (−)-premarineosin A production in *S. eitanensis*. Thus, we engineered *S. eitanensis* for enhanced (−)-premarineosin A production using a regulatory gene expression strategy^30^. The *pmaD* gene, amplified from the *S. eitanensis* genome, was assembled in pSET152k under the strong constitutive promoter KasOp*, yielding pSET152k-*pmaD*. This plasmid was integrated at the ΦC31 actinophage integrase *attB* site in the *S. eitanensis* genome, generating the *pmaD* overexpressing strain FP10017 (**Supplementary Fig. 2**). To ensure that the producing phenotype was directly attributed to overexpression of *pmaD*, we integrated the empty vector into the *S. eitanensis* genome, generating the strain FP10016. Under non-optimized culture conditions (corn gluten meal, GICYE), the FP10017 strain produced significantly more (−)-premarineosin A (14.7 ± 1.4 mg/L) than the vector control FP10016 strain (1.0 ± 0.4 mg/L) and the wild-type strain (below the limit of quantification) (**Fig. 1d**).

In the last step of (−)-premarineosin A biosynthesis, the putative Rieske oxygenase PmaG catalyzes the bicyclization of 23-HUP into **2**^6,12^. As accumulation of 23-HUP was observed in the engineered FP10017 strain (**Fig. 1d**), we designed pSET152k-*pmaDG* for constitutive overexpression of *pmaG* and *pmaD in S. eitanensis*. This plasmid was integrated into *S. eitanensis* wild-type, yielding the FP10018 strain. Constitutive overexpression of *pmaG* and *pmaD* significantly enhanced (−)-premarineosin A production (23.3 ± 0.5 mg/L) in FP10018, a 1.6-fold increase compared to the overexpression of *pmaD* alone. The metabolic shift toward the production of **2** is also reflected by the difference in color between the *pmaG* and *pmaD* overexpressing strain FP10018 (red) and the FP10016 (green) strain (**Fig. 1e**).

In addition to gene overexpression strategies, optimization of media composition and culture conditions can enhance secondary metabolite production in *Streptomyces*^30,36,41,42^. Cultivation of the engineered FP10018 strain in soybean meal medium (GISYE) significantly (p < 0.0001) enhanced (−)-premarineosin A production by 1.39-fold compared to cultivation in GICYE (**Fig. 1d, Supplementary Fig. 3**). Production of **2** and **1** was assessed throughout cultivation, and while maximum titers of (−)-premarineosin A were obtained around day six, the titers of 23-HUP were lowest on day seven (**Fig. 1f, Supplementary Fig. 4**). Both acetone and methanol were investigated as extraction solvents, with acetone showing, on average, higher (−)-premarineosin A and 23-HUP recovery (**Fig. 1d, Supplementary Fig. 5**). Under optimal conditions (GISYE at 22 °C for seven days followed by extraction with acetone), (−)-premarineosin A production increased over 200-fold in the engineered strains FP10017 and FP10018 (34.8 ± 1.6 mg/L v 33.7 ± 5.1 mg/L, respectively) when compared to wild-type (0.2 ± 0.1 mg/L) (**Fig. 1d**). In GISYE, the vector control strain FP10016 produced 20-fold more **2** (3.0 ± 2.1 mg/L) compared to wild-type. The FP10018 strain produces nearly twice (p < 0.05) as much **2** per gram of dry cell weight (5.1 mg/g DCW) as strain FP10017 (2.6 mg/g DCW) (**Fig. 1f**). Production of 23-HUP was also increased in the engineered strains, with significantly (p < 0.0005) increased production in the FP10018 strain compared to the wild-type when cultured in GISYE (**Fig. 1d**).

Efforts to maximize (−)-premarineosin A isolation and purification from the engineered strains FP10017 and FP10018 yielded 12.9 ± 1.2 mg/L and 11.8 ± 2.9 mg/L, respectively, with >95% purity (**Fig. 1g, Supplementary Fig. 6**). Structural characterization was conducted using NMR spectroscopy (**Supplementary Fig. 11-16**)^12^, X-ray crystallography (**Fig. 1c**), and polarimetry. While the relative stereochemistry of (−)-premarineosin A confirms previously described reports^7,13,14^, our X-ray crystal data (**Fig. 1c**) revealed that its absolute stereochemistry did not agree with the premarineosin A structure obtained by total synthesis^26,27^. As the optical rotation value for the compound (**2**) isolated in this work was [α]_D_^24^ (c 0.4758 in MeOH) = -104.0 ± 0.1°, we refer to the compound as (−)-premarineosin A.

Isolated compound **2** was tested against the chloroquine (CQ)-sensitive 3D7 and CQ-resistant Dd2 *P. falciparum* parasites in an *in vitro* assay (**Fig. 2**). The compound was concomitantly evaluated for cytotoxicity against the HEK293 embryonic kidney and MOLT4 T-cell leukemia cell lines. (−)-Premarineosin A exhibits nanomolar potency against the 3D7 (EC_50_ = 4.0 ± 0.1 nM) and Dd2 (EC_50_ = 1.0 ± 0.7 nM) *P. falciparum* parasites with minimal toxicity to mammalian cells (**Fig. 2**). In comparison, **2** is over 10- (3D7) and 350-fold (Dd2) more potent than chloroquine and is 6-(3D7) and 8-fold (Dd2) more potent than artesunate (**Fig. 2**).

**Figure 2.**
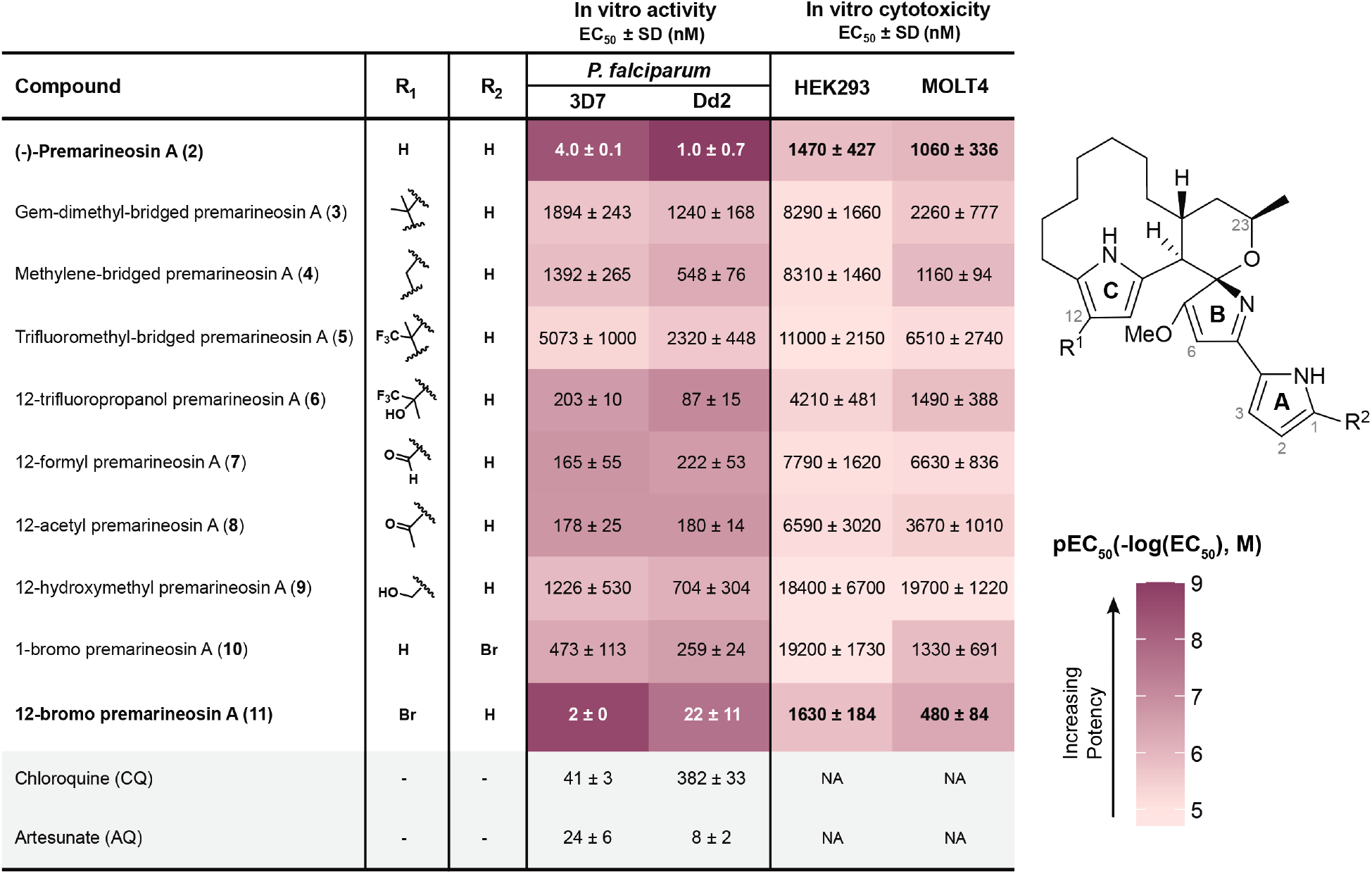
*In vitro* bioactivities of (−)-premarineosin A (2) and derivatives (3-11) show selectivity against *Plasmodium falciparum*. Compounds were evaluated against 3D7 (chloroquine-sensitive) and Dd2 (multidrug-resistant) *P. falciparum* strains for antimalarial activity and the HEK293 embryonic kidney and MOLT4 T-cell leukemia lines for mammalian cell cytotoxicity (72 h treatment). EC50: half-maximal effective concentration. NA: Not assessed. SD: Standard Deviation

### Acid-catalyzed electrophilic aromatic substitution enabled late-stage derivatization of (−)-premarineosin A

With access to sufficient quantities of (−)-premarineosin A, we pursued a semi-synthetic approach to functionalize its scaffold for the first structure-activity relationship analysis. Dimerization of natural products is a known approach to increase their structural complexity and biological activity^44,45^. Previous synthetic studies established that simple 2,5-dimethylpyrroles are susceptible to acid-catalyzed condensation with ketones, such as acetone, to form dimerized substrates at the 3-position in an electrophilic aromatic substitution-like reaction^46^. To our knowledge, this synthetic approach has not been previously applied to any natural product scaffold. Thus, we reasoned that exploration of the breadth and flexibility of this chemistry may enable access to previously untapped chemical space. Applying similar methodologies, we found that (−)-premarineosin A dimerizes under acidic conditions with both ketones and aldehydes at the C12 position of its C-ring, forming premarineosin A dimers with a gem-dimethyl-bridge (**3**), a methylene-bridge (**4**), and a trifluoromethyl-bridge (**5**) (**Fig. 3a, top**). A crystal structure of **3** was obtained to confirm its unique structure, stereochemistry, and absolute configuration (**Fig. 3b**). Functionalization of **2** with trifluoroacetone also yielded a monomeric derivative (12-trifluoropropanol premarineosin A, (**6**)), likely due to early reaction termination before complete substrate consumption (**Supplementary Fig. 7**). Premature termination using acetone or formaldehyde as the electrophile did not generate monomeric intermediates, suggesting that isolating these monomeric (−)-premarineosin A derivatives under electrophilic aromatic substitution conditions is only possible when using electron-withdrawing groups^46^. To synthesize other monomeric derivatives of **2**, we used acid chlorides as the electrophile for Friedel-Crafts acylation to form 12-formyl premarineosin A (**7**) and 12-acetyl premarineosin A (**8**). Further reduction of **7** with sodium borohydride provided the desired primary alcohol, forming 12-hydroxymethyl-premarineosin A (**9**) (**Fig. 3a, bottom**).

**Figure 3.**
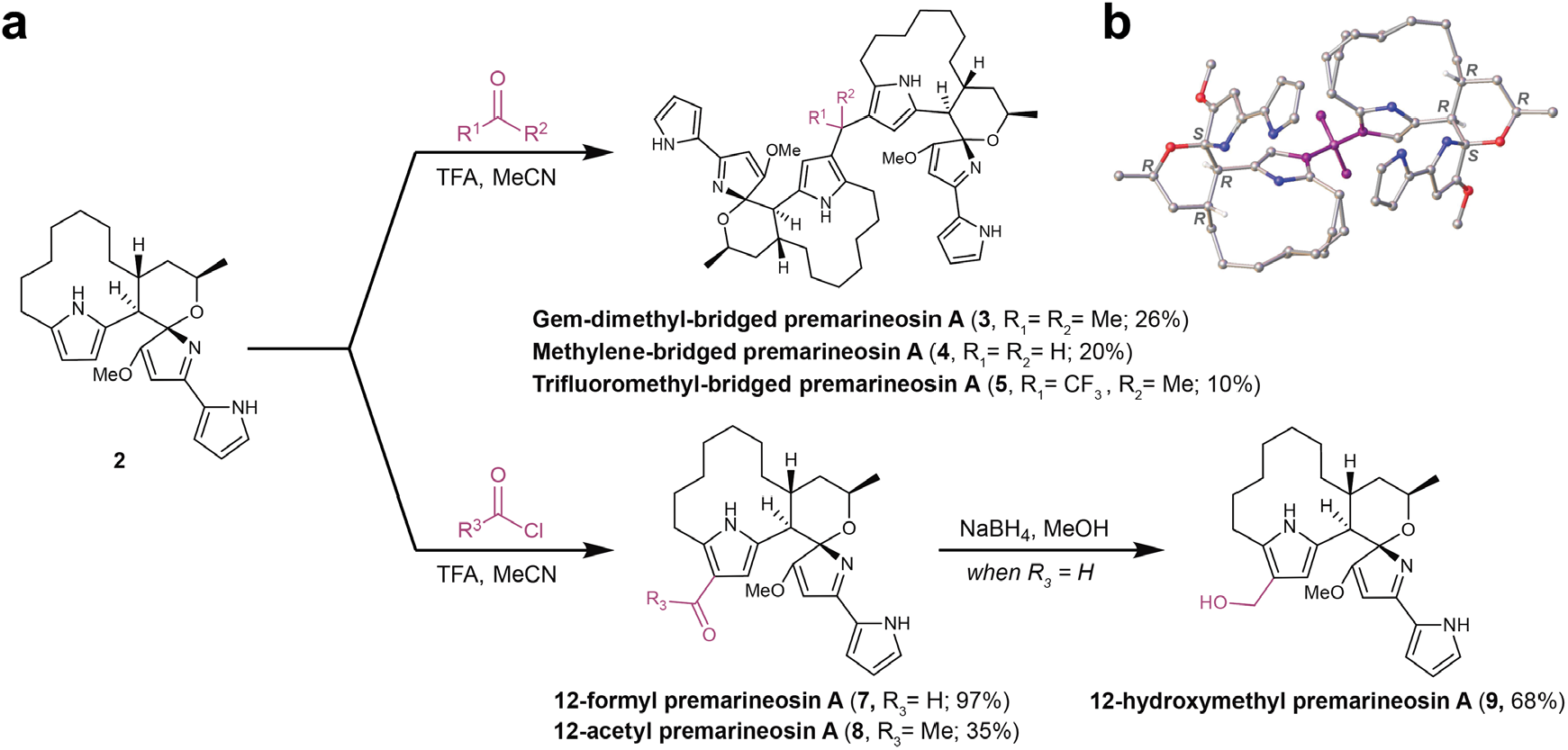
Semi-synthetic electrophilic aromatic substitution at the C12 position facilitated robust derivatization of the (−)-premarineosin A scaffold. **a**. Synthesis of dimeric (top) and monomeric (bottom) premarineosin A analogs. **b**. X-ray crystal structure of **3**.

The antiparasitic activity and cytotoxicity of these synthetic derivatives were evaluated in comparison to **2** (**Fig. 3**). The monomeric derivatives **6, 7**, and **8** display sub-micromolar activity against the 3D7 and Dd2 *P. falciparum* parasites. Despite the reduced activity compared to (−)-premarineosin A, derivatives **6, 7**, and **8** perform better than chloroquine (EC_50_ = 382 ± 22 nM) against Dd2. Compound **6** is the most promising semi-synthetic derivative in terms of selectivity, displaying moderate potency against Dd2 (EC_50_ = 87 ± 15 nM) with limited apparent toxicity to HEK293 cells (EC_50_ = 4210 ± 481 nM). Compound **4** (EC_50_ = 548 ± 76 nM) and **9** (EC_50_ = 704 ± 304 nM) also exhibited sub-micromolar potency against the multidrug-resistant *P. falciparum* strain yet had limited potency against the drug-sensitive strain. While these synthetic modifications at the C-ring reduced the antimalarial potency of the premarineosin A scaffold, the analogs broadly exhibit decreased cytotoxicity.

### Biocatalytic and semi-synthetic bromination enabled late-stage C-H functionalization of (−)-premarineosin A

Bromine holds significant value in medicinal chemistry due to its capacity to improve key pharmacokinetic properties, such as membrane permeability, target binding affinity, and metabolic stability—factors crucial for combating *Plasmodium* species effectively^47,48^. To harness the potential of selective C-H bromination of the (−)-premarineosin A scaffold, we computationally screened two bacterial flavin-dependent halogenases (FDHs). Molecular docking of **2** was performed with D3, an orphan FDH from *Saccharophagus degradans*^49^, and RebH, a FDH from the rebeccamycin biosynthetic pathway^50^. While RebH shows low substrate binding and minimal contact with the active site, the D3 halogenase revealed strong substrate binding (−8.6 kcal/mol), with (−)-premarineosin A oriented within the active site in a conformation that suggests the A-ring pyrrole would be most accessible for bromination (**Fig. 4a, b**). Reactions of D3 with (−)-premarineosin A *in vitro* resulted in a single brominated derivative, 1-bromo premarineosin A (**10**), with approximately 50% and 10% substrate conversion efficiency at small- and large-scale, respectively (**Fig. 4c, e**).

**Figure 4.**
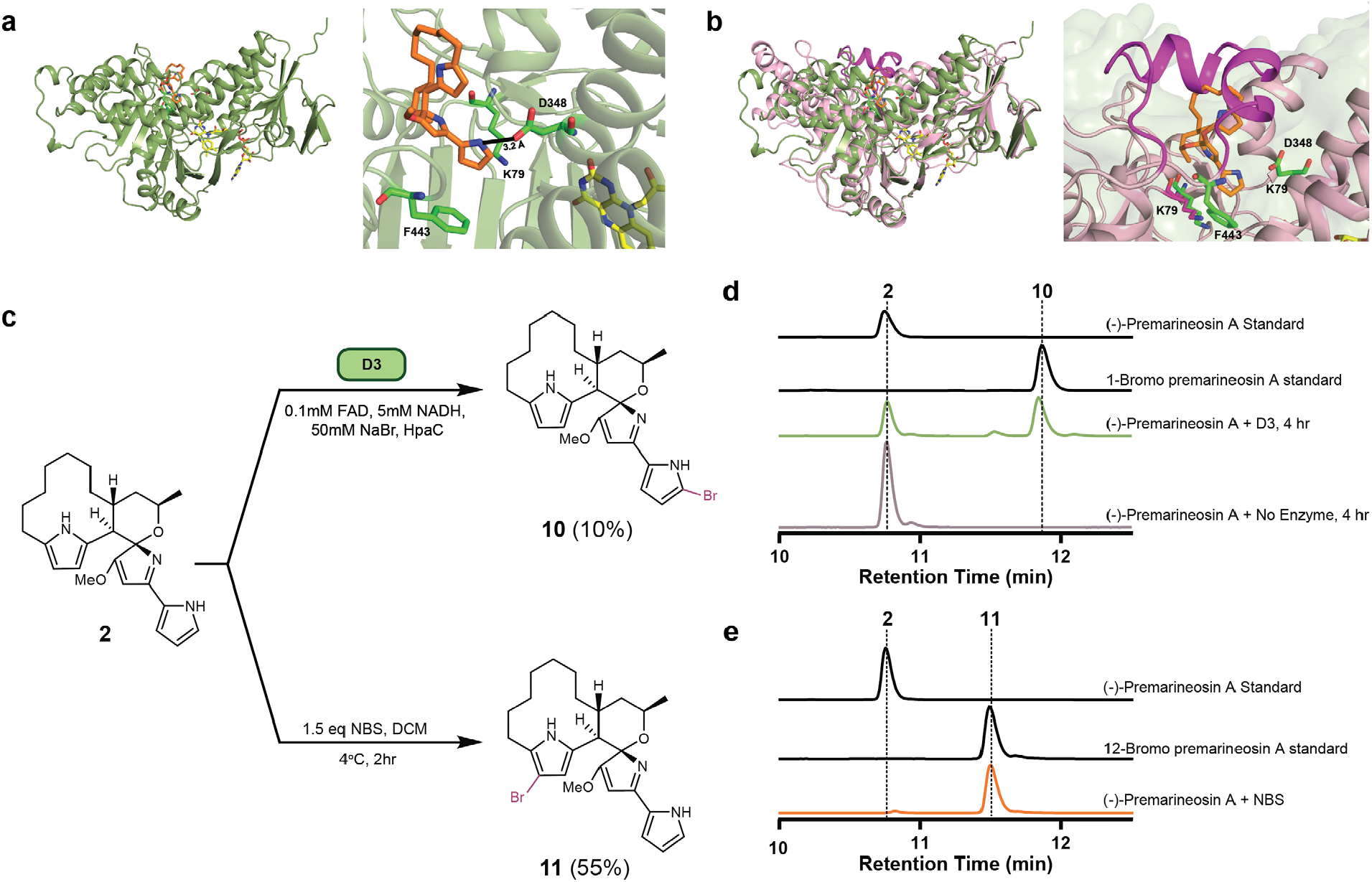
Site-selective biocatalytic and semi-synthetic strategies enabled the late-stage bromination of (−)-premarineosin A. **a**. In silico docking of (−)-premarineosin A (orange) into the AlphaFold-predicted structure of D3 (sage green) with FAD (yellow), showing favorable positioning for electrophilic aromatic substitution at the A pyrrole ring and displaying proximity to critical residues for catalysis (Lys79). **b**. Overlay of (−)-premarineosin A (orange) docked in D3 (sage green) with the crystal structure of RebH (PBD ID: 2OA1) (pink) showing the sterically hindering helix (magenta) that prevents binding in RebH, as opposed to the large, open active site cavity of D3, which accommodates (−)-premarineosin A. **c**. Biocatalysis (top) and semi-synthesis (bottom) of brominated premarineosin A analogs. **d**. HPLC traces of D3 and no enzyme control reactions. A peak corresponding to 1-bromo premarineosin A appears post-reaction. **e**. HPLC traces of the NBS reaction. A peak corresponding to 12-bromo premarineosin A appears post-reaction.

To evaluate product formation and regioselectivity, we compared this enzyme-catalyzed bromination with a synthetic approach. *N*-bromosuccinimide (NBS) was selected as the brominating agent because of its proven effectiveness in regioselective electrophilic aromatic bromination.^51^ Reaction with NBS at 4 °C in DCM for two hours nearly consumed (−)-premarineosin A, primarily producing a C-ring brominated product (55% isolated yield, 10mg reaction), 12-bromo premarineosin A (**11, Fig. 4c, e**).

The antiparasitic and cytotoxic activities of the brominated premarineosin A derivatives were also assessed *in vitro*. 1-Bromo premarineosin A (**10**) exhibited sub-micromolar antiparasitic activity, with EC_50_ values of 473 ± 113 nM (3D7) and 259 ± 24 nM (Dd2). On the other hand, 12-bromo premarineosin A (**11**) demonstrated high potency against 3D7 (EC_50_ = 2 nM) and Dd2 (EC_50_ = 22 ± 11 nM) with low cytotoxicity (**Fig. 2**). These findings highlight 12-bromo premarineosin A (**11**) as a lead compound, combining exceptional antimalarial potency and lower cytotoxicity with remarkable selectivity.

## Discussion

This work presents expanded efforts that highlight premarineosin A (**2**) as a promising scaffold for the development of antimalarial therapeutics. Overexpression of the cluster-situated regulator PmaD in *S. eitanensis* increased (−)-premarineosin A titers to 34.8 ± 1.6 mg/L, over 200-fold compared to *S. eitanensis* wild-type (0.2 ± 0.1 mg/L). While the difference in (−)-premarineosin A levels between the engineered FP10017 and FP10018 strains under optimized conditions was not statistically significant, we did observe that FP10018 produces nearly twice as much **2** per gram of dry cell weight in soybean meal media (**Fig. 1F**). As the accumulation of (−)-premarineosin A and 23-HUP might inhibit cell growth, we reason that further culture optimization or adaptive laboratory evolution can be employed to increase cell viability in the engineered FP10018 strain and enhance compound titers. Nevertheless, the engineered *S. eitanensis* FP10017 and FP10018 strains boast the highest production and isolation titers of (−)-premarineosin A reported to date. Interestingly, the strain integrating the empty vector (FP10016) also produced nearly 20-fold more (−)-premarineosin A than the wild-type in the soybean media. The increased production of (−)-premarineosin A in the engineered FP10016 strain could be a result of plasmid integration at the ΦC31 integrase (*attB*) site, which has been shown to alter the availability of malonyl-CoA and acetyl-CoA^52^ and induce a stress response^53^, stimulating secondary metabolite production.

Contradictory data regarding the stereochemistry of premarineosin-associated molecules have been reported, so we sought to definitively establish its absolute configuration. The stereocenters of the (−)-premarineosin A isolated in this work have the same relative configuration as previously reported structures of premarineosin A from both natural product isolation^6,12,20^ and total synthesis^26,27^ studies. However, the crystal structures of **2** and the gem-dimethyl-bridged dimer (**3**) support an absolute configuration of *8S, 9R, 21R, 23R*, which is enantiomeric in respect to the previously reported stereochemistry^6,12,26,27^ (**Fig. 1c, 3b**). Efforts toward the total synthesis of marineosin A by Shi et al.^27^ and Harran et al.^26^ followed the absolute stereochemistry proposed by Reynolds et al.^6,12^ based on the conclusion that marineosin A is biosynthesized from (23*S*)-HUP rather than (23*R*)-HUP. The optical rotation values for marineosin A obtained by total synthesis ([α]_D_^25^ = +62.6° (c 0.16, MeOH) by Shi et al.^27^, and [α]_D_^25^ = +138.7° (c 0.02, MeOH) by Harran et al.^26^) oppose the value originally reported by Fenical et al.^20^ for the natural product isolated from *Streptomyces* sp. CNQ-617 ([α] ;]_D_^24^ = −101.7° (c 0.06, MeOH)). Shi et al. noted additional differences in compound properties and suggested that their synthetic compound was not identical to the isolated marineosin A, but could instead be an isomer.^27^ Harran et al. instead postulated that the difference could be explained by an accidental interchange of the marineosin A and B optical rotation values reported by Fenical et al., claiming that, like their synthetic product, the original isolated marineosin A could be dextrorotatory.^26^ As the absolute configuration of the (−)-premarineosin A isolated in this study is enantiomeric respective to that reported by Reynolds, Shi, and Harran, we sought to confirm the absolute configuration of isolated marineosin A from *Streptomyces* sp. CNQ-617. Thus, marineosin A was isolated from this original producing strain and its structural identity was confirmed by NMR (**Supplementary Fig. 66**). Optical rotation analysis for marineosin A isolated in this work ([α]_D_^25^ = -105.8 ± 2.1° (c 0.2525, MeOH)) supported the polarity value (−) and was of similar magnitude to the value initially reported by Fenical et al.^20^ Together, these results suggest that the structure-confirmed dextrorotatory product obtained by total synthesis, (+)-marineosin A, is likely the enantiomer of the confirmed levorotatory marineosin A isomer isolated from *Streptomyces* sp. CNQ-617. Hence, the natural product marineosin A shares the same absolute stereochemistry at C8 (*S)*, C9 (*R*), C21 (*R*), and 23C (*R*) as the (−)-premarineosin A isolated from *S. eitanensis*.

Engineering *in vivo* production for sustainable and scalable access to **2** not only allowed us to resolve this stereochemical ambiguity but also enabled our expansion of the chemical space around this scaffold, culminating in the synthesis of nine new analogs. We first focused on the surprising discovery of pyrrole dimerization chemistry to diversify the C12 position of the (−)-premarineosin A C-ring. While electrophilic substitution has been utilized previously to functionalize the β-position of pyrroles^54,55^, to our knowledge this methodology has not been explored with any complex natural product scaffold. This reaction platform revealed an underexplored acid-catalyzed electrophilic aromatic substitution diversification strategy for pyrrolic alkaloids, highlighting (−)-premarineosin A as a synthetically pliable and chemically evolvable scaffold. Of the seven semi-synthetic premarineosin A derivatives (**3-9**), five show sub-micromolar potency against the chloroquine-resistant *P. falciparum* Dd2, with three having a lower EC_50_ than chloroquine (**Fig. 2**). However, they all have reduced bioactivity compared to **2**, suggesting that the C12 position plays a crucial role in the antimalarial activity of the premarineosin A scaffold (**Fig. 2, 5**). Remarkably, **3 – 9** are overall less cytotoxic than **2**, in agreement with the bioactivity of other C-ring functionalized prodiginines^56–58^. Hence, functionalizing the C12 position of the C-ring is an encouraging avenue to increase the selectivity of premarineosin A and improve its therapeutic potential. To explore this trend further, we carried out NBS-mediated bromination of (−)-premarineosin A. The resulting 12-bromo premarineosin A (**11**) derivative was substantially more potent (17-fold) than chloroquine against the multidrug-resistant *P. falciparum* Dd2 strain. While **11** did not outperform the outstanding potency and selectivity of the natural product (−)-premarineosin A against the Dd2 strain, **11** was twice as potent as **2** against the drug-sensitive *P. falciparum* 3D7 with improved selectivity relative to HEK293 mammalian cells (**Fig. 2, 5**). Bromination of the C12-position of (−)-premarineosin A decreased cytotoxicity while maintaining single-digit nanomolar potency against *P. falciparum* (EC_50_ < 5 nM), offering a unique opportunity to enhance drug efficacy, stability, and bioavailability^60,61^. Given that its potency exceeds both artesunate and chloroquine in the 3D7 strain, these results underscore **11** as a compelling lead compound for antimalarial drug development.

**Figure 5.**
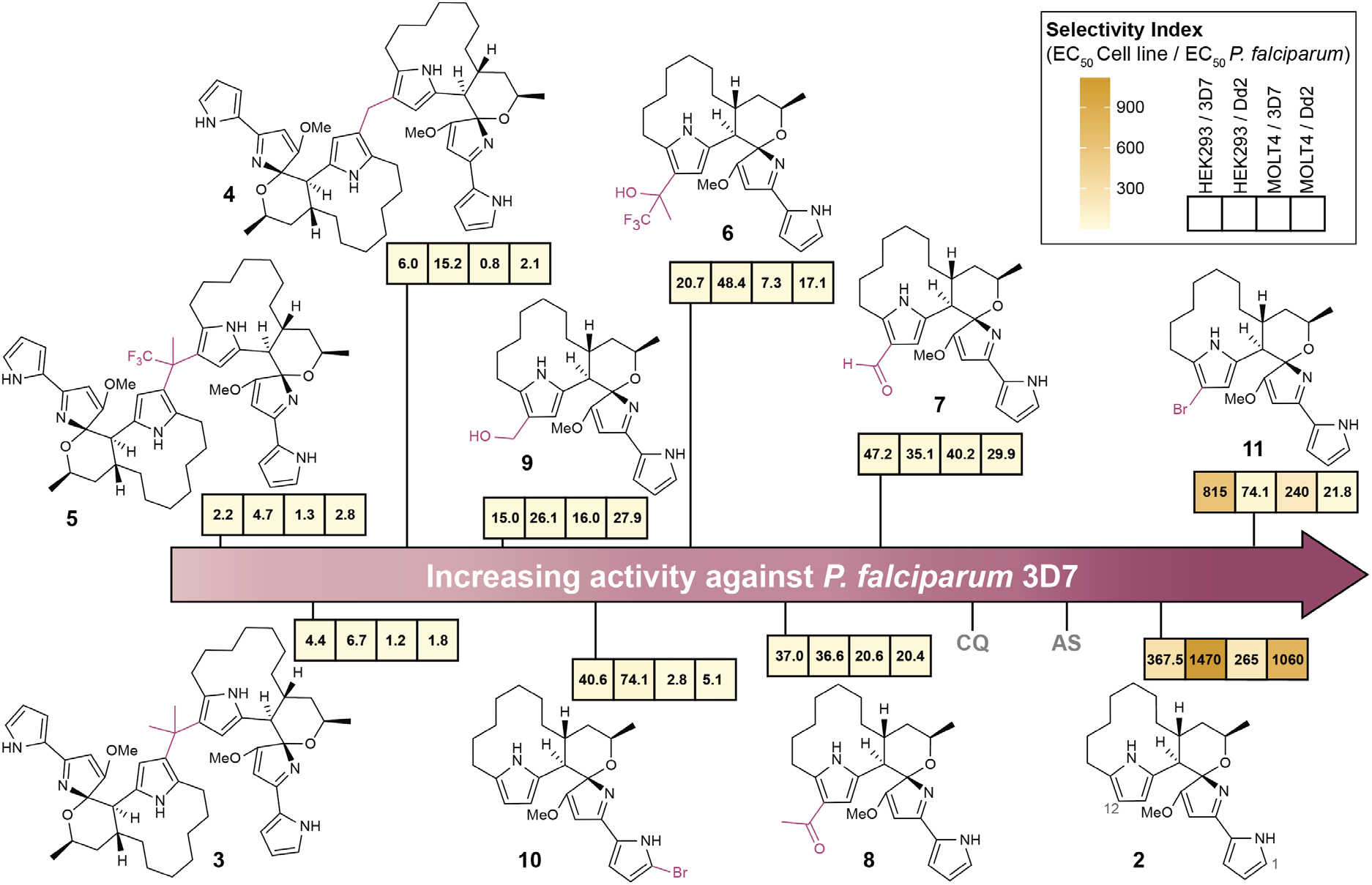
Diversifying the (−)-premarineosin A structure improved selectivity against *Plasmodium falciparum*. Compounds were tested against 3D7 (chloroquine-sensitive) and Dd2 (chloroquine-resistant) *P. falciparum* strains for antimalarial activity and the HEK293 embryonic kidney and MOLT4 T-cell leukemia cell lines for cytotoxicity to mammalian cells. Selectivity Index = EC_50_ Mammalian Cell Line / EC_50_ *P. falciparum*.

As a complement to synthetic chemistry methods, we next turned to biocatalysis to probe unexplored regions of the premarineosin A chemical space. To modify (−)-premarineosin A beyond the C12 position, we explored FDHs for late-stage C-H bromination of the A-ring pyrrole, which is beyond the reach of our semi-synthetic approach. While previous studies have used FDHs to halogenate simple pyrrole-containing precursors in early biosynthetic steps^59^, their application to fully assembled, highly complex natural products has remained largely unexplored. As such, our strategy of using the orphan FDH D3 from *S. degradans* to selectively brominate (−)-premarineosin A on the minimally substituted (A-ring) pyrrole holds significant promise for expanding the scope of enzymatic halogenation (**Fig. 3d**). This result not only highlights the catalytic flexibility of D3 but demonstrates the feasibility of using non-native halogenases to derivatize densely functionalized, late-stage scaffolds. As depicted in the AlphaFold-generated structure of D3 docked with (−)-premarineosin A (**Fig. 3a**), the enzyme’s unusually large and flexible active site pocket appears well-suited to readily accommodate the bulky (−)-premarineosin A scaffold in a conformation conducive for strong substrate binding and regio-selective bromination at the A-ring C1-position. The contrasting regioselectivity of the semi-synthetic and biosynthetic methods motivates further enzyme engineering studies to assess the efficiency and selectivity of C-H halogenation of the premarineosin scaffold. Although the resulting 1-bromo premarineosin A (**10**) derivative showed enhanced potency relative to chloroquine against the chloroquine-resistant *P. falciparum* Dd2 strain and reduced cytotoxicity compared to **2** and **11** against both cell lines, it exhibited decreased selectivity and antimalarial activity overall, suggesting the C1-position may play a critical role in the therapeutic efficacy of (−)-premarineosin A (**Fig. 2, 5**). Nevertheless, this work shows that even without prior engineering, naturally occurring and previously uncharacterized enzymes can be identified through computational modeling and repurposed to modify complex cyclic natural products. The ability of D3 to accept a bulky, non-native scaffold like (−)-premarineosin A and catalyze regioselective halogenation, despite being predicted to be a simple tryptophan halogenase, underscores the untapped potential of orphan biocatalysts. More broadly, this result emphasizes the value of pairing *in silico* screening and experimental validation to unveil enzymes that are inherently capable of functionalizing sites that are generally inaccessible via traditional synthetic methods.

Together, our results establish a tractable and modifiable framework for late-stage functionalization of (−)-premarineosin A and demonstrate an approach for strategic expansion of scaffold-level chemical space using both synthetic and biocatalytic approaches. In addition, the remarkable selectivity of 12-bromo premarineosin A (**11**) against *P. falciparum* warrants further investigation into its mechanism of action and *in vivo* efficacy studies. By combining metabolic engineering, biocatalysis, computational modeling, and semi-synthesis, we define a versatile platform for late-stage functionalization of complex pyrrolic alkaloids and create a roadmap for the systematic exploration of (−)-premarineosin A chemical space in the search for next-generation antimalarial agents.

## Materials and Methods

### Strains and culture conditions

All strains that were designed and/or cultured in this work are listed in **Supplementary Table 2**. Plasmid assembly, replication, and preservation were performed using high-efficiency *Escherichia coli* DH5α (NEB). *E. coli* S17-1 was used as the mobilization host for conjugation with *S. eitanensis*. All *E. coli* strains were cultivated in LB medium (10 g/L tryptone, 10 g/L NaCl, and 5 g/L yeast extract in ultrapure water) at 37 °C. The LB media was supplemented with 50 μg/mL kanamycin for plasmid maintenance and selection. All *Streptomyces* strains were cultivated in 2xYT (16 g/L tryptone, 10 g/L yeast extract, and 5 g/L NaCl in ultrapure water) or peanut meal & starch media (10 g/L glucose, 30 g/L starch, 5 g/L bacto peptone, 10 g/L peanut meal, 5 g/L yeast extract, 2 g/L CaCO_3_ in ultrapure water) for three days at 28 °C for seed culture. Strains were sporulated in OPAH (1 g/L oatmeal, 1 g/L pharmamedia, 1 g/L arabinose, 0.5 g/L humic acid, 0.5 mM KH_2_PO_4_, 0.5 mM CaCl_2_, 0.5 mM MgSO_4_, 1.9 mg/L Na_2_-EDTA·2H_2_O, 1.4 mg/L FeSO_4_·7H_2_O, 0.2 mg/L H_3_BO_3_, 0.05 mg/L MnSO_4_·H_2_O, 0.01 mg/L ZnSO_4_·7H_2_O, 0.01 mg/L Na_2_MoO_4_·2H_2_O, 0.01 mg/L CuSO_4_, and 0.01 mg/L CoCl_2_ in ultrapure water). Media was supplemented with 50 μg/mL kanamycin (Goldbio, K-120-25) and/or 25 μg/mL nalidixic acid (Cayman, 19807) as needed.

### Plasmid design and genome editing

All plasmids generated in this study are described in **Supplementary Table 3**. The *pmaD* and *pmaG* genes were amplified from wild-type *S. eitanensis* gDNA, then assembled via Gibson Assembly in plasmid pSET152k*-kasOp\** between the BamHI and EcoRI restriction sites (New England Biolabs, R0136S and R0101S), as previously described^22^. Synthetic ribosomal binding sites^45^ were used as overlapping regions for multi-gene vector design. Primers were designed with guidance from pyDNA^46^ (**Supplementary Table 4**). An integrative system, mediated by the *att*P site of the *Streptomyces* phage ΦC31^43^, was used for constitutive overexpression in *S. eitanensis*. Interspecies conjugation was performed to transfer DNA from *E. coli* S171 to S. eitanensis. Following incubation (12 h, 28 °C), the plates were overlaid with 1 mL sterilized water with 1.25 mg kanamycin and 0.5 mg nalidixic acid. After 3–5 days, recombinants were transferred to OPAH plates with 25 μg/mL nalidixic acid and 50 μg/mL kanamycin. The engineered antibiotic-resistant strains were whole genome sequenced. Cluster comparison and visualization were performed with clinker^47^ and DNAviewer^48^ to confirm genome integration (**Supplementary Fig. 2**).

### Genome extraction, sequencing, and assembly

High-quality genomic DNA was prepared using reagents from the MasterPure Complete DNA and RNA Purification Kit (Lucigen, MC85200). The wild-type and engineered *Streptomyces* strains were cultivated in 2xYT for three days at 28 °C. Cells were pelleted via centrifugation, resuspended in 480 μL EDTA (50 mM, pH 8, sterile filtered) and 120 μL lysozyme (10 mg/mL, DotScientific DSL38100-10) then incubated for 35 min at 37 °C. Post-incubation, the cells were centrifuged (1 min, 17,000 x g, RT) and resuspended in 200 μL MasterPure Tissue and Cell Lysis Solution treated with 1 μL Proteinase K (Qiagen RP107B-5). Cells were incubated for 15 min at 65 °C, incubated for 2 min at 95 °C, then cooled to RT. 30 μL of RNase A (10 mg/mL, Sigma Aldrich R5503-1G) was added prior to incubation (1.5 hr, 37 °C). The genomic DNA was precipitated following the MasterPure Kit Precipitation of Total Nucleic Acids protocol. Genomic DNA quality was assessed using gel electrophoresis. *S. eitanensis* bacterial genome sequencing was performed by Plasmidsaurus using Oxford Nanopore Technology with custom analysis and annotation.

### (−)-Premarineosin A production conditions

Spores from OPAH plates were inoculated into 50 mL of peanut meal & starch media (10 g/L glucose, 30 g/L starch, 5 g/L bacto peptone, 10 g/L peanut meal, 5 g/L yeast extract, 2 g/L CaCO_3_ in ultrapure water) and incubated (three days, 28 °C, 200 rpm). 10 mL of this pre-inoculum culture was used to inoculate 1 L of producing media: GICYE (10 g/L glucose, 30 g/L inulin, 5 g/L bacto peptone, 10 g/L corn gluten meal, 5 g/L yeast extract, 2 g/L CaCO_3_ in ultrapure water, adjusted to pH=7.0) or GISYE (10 g/L glucose, 30 g/L inulin, 5 g/L bacto peptone, 10 g/L soybean meal, 5 g/L yeast extract, 2 g/L CaCO_3_ in ultrapure water, adjusted to pH=7.0) in 2.8 L Fernbach flasks. Cultures were incubated for seven days at 22 °C and 170 rpm.

### Sample preparation and quantification of (−)-premarineosin A

Quantification of (−)-premarineosin A production in 1 L cultures was performed by extracting 1 mL biomass samples from three independent replicates. Biomass was collected and extracted every day for seven days to estimate production over time. Compound quantification is from extracts collected on day seven unless stated otherwise. On each collection day, 1mL was collected from each 1 L culture and the biomass was pelleted via centrifugation (4 min, 20800 rpm, RT). The cell pellet was resuspended in 1 mL of solvent (methanol or acetone) and 100 μL of glass beads. Supernatant was combined 50:50 with solvent and stored at -20 °C until quantification. Tubes were vortexed (2 hours, 4 °C), followed by centrifugation (4 min, 20800 rpm, RT). The supernatant was retained as the extract and was stored at -20 °C until quantification. Quantification of (−)-premarineosin A production was performed using analytical HPLC (Shimadzu) equipped with a PDA detector and analyzed with a Phenyl-Hexyl column (Luna 5 μM Phenyl-Hexyl 100 Å, LC Column 250 × 4.6mm, heated to 40 °C). Water + 0.1% Formic Acid (A) / Acetonitrile + 0.1% Formic Acid (B) (10% to 100% B) was used for the mobile phase at 2 ml/min. Calibration curves were prepared with known concentrations of pure (−)-premarineosin A and 23-HUP. Production titers were determined by comparing the calibration curve and sample peak areas (AUC) at 346 nm ((−)-premarineosin A) and 520 nm (23-HUP). (−)-Premarineosin A and 23-HUP were detected in culture supernatants at trace levels for all tested strains. Therefore, quantifications reported in this document correspond to their intracellular concentrations. Statistical analysis and visualization were performed with GraphPad Prism v. 10.4.2., with p-values calculated as the result of a two-tailed T-test.

### Biomass quantification

Daily 1 mL samples were collected from each of the three independent replicate cultures and vacuum filtered on a pre-dried membrane filter. The retained cells were dried in a microwave oven for 1.5 min^62^. As *S. eitanensis* is known to form small, spherical pellets in liquid culture, sampling was performed using sterile wide-bore pipette tips and serological pipettes.

### Isolation and purification of (−)-premarineosin A and 23-HUP

The total biomass (from 1 L culture) was separated via vacuum filtration. Cell pellets were broken by coating the cells with 750 mL of 100% acetone and then shaking overnight. The acetone was obtained by vacuum filtration, concentrated, and liquid-liquid extracted with ethyl acetate. The dried organic layer was concentrated with silica to be dry loaded for normal phase purification on a Biotage Isolera flash column system. Initial rounds of purification utilized a 40 g silica column with an ethyl acetate/hexane gradient (15-100%) with 1% acetic acid; (−)-premarineosin A eluted at 34-50% and 23-HUP eluted at 20-34%. (−)-Premarineosin A was further purified on a 40 g silica column with a (3% ammonia (7N in methanol) in DCM solution) / hexane gradient (15-100%); (−)-premarineosin A eluted at 20-30%. 23-HUP fraction(s) were further purified using preparative thin-layer chromatography with a 3% Ammonia (7N) Methanol in DCM mobile phase. Purity assessment of isolated compounds was performed using HPLC, NMR, and LC-MS/MS (**Supplementary Fig. 6**). All samples for bioactivity testing were >95% pure. Optical rotations were obtained using a Jasco P2000 polarimeter with a 100 mm cell.

### Marineosin A production, isolation, and purification conditions

Spores from modified A1 plates^20^ (10 g/L starch, 4 g/L yeast extract, 2 g/L peptone, 36 g/L Instant Ocean) were inoculated into 50 mL of peanut meal & starch media (10 g/L glucose, 30 g/L starch, 5 g/L bacto peptone, 10 g/L peanut meal, 5 g/L yeast extract, 2 g/L CaCO_3_ in ultrapure water) and incubated (three days, 28 °C, 200 rpm). 10 mL of this pre-inoculum culture was used to inoculate 1 L of producing media: GICYE (10 g/L glucose, 30 g/L inulin, 5 g/L bacto peptone, 10 g/L corn gluten meal, 5 g/L yeast extract, 2 g/L CaCO_3_ in ultrapure water, adjusted to pH=7.0) in 2.8 L Fernbach flasks. Cultures were incubated for seven days at 22 °C and 170 rpm. The total biomass (from 1 L culture) was separated via centrifugation. Cell pellets were broken by coating the cells with 750 mL of 100% acetone and then shaking overnight. The acetone extract was obtained by vacuum filtration, concentrated, and liquid-liquid extracted with ethyl acetate. The dried organic layer was concentrated with silica to be dry loaded for normal phase purification on a Biotage Isolera flash column system, utilizing a 40 g silica column with an ethyl acetate/hexane gradient (15-100%) with 1% acetic acid; marineosin A eluted at 34-50%. Marineosin A fraction(s) were further purified using preparative thin-layer chromatography with a 3% Ammonia (7N) Methanol in DCM mobile phase; marineosin A eluted as a strong UV-active band near the top of the plate. Purity assessment of isolated compounds was performed using HPLC, NMR (**Supplementary Fig. 67**), and LC-MS/MS. Optical rotations were obtained using a Jasco P2000 polarimeter with a 100 mm cell.

### (−)-Premarineosin A

^1^H NMR (599 MHz, Acetone) δ 13.65 (s, 1H), 12.05 (s, 1H), 9.80 (s, 1H), 7.65 (ddd, *J* = 3.3, 2.3, 1.3 Hz, 1H), 7.45 (s, 1H), 6.50 (dt, *J* = 4.0, 2.1 Hz, 1H), 6.26 (d, *J* = 1.7 Hz, 1H), 5.72 (t, *J* = 2.9 Hz, 1H), 5.52 (t, *J* = 2.9 Hz, 1H), 4.46 – 4.38 (m, 1H), 4.14 (s, 3H), 3.03 (d, *J* = 12.5 Hz, 1H), 2.69 (dp, *J* = 13.4, 4.9 Hz, 1H), 2.39 (dt, *J* = 14.8, 4.4 Hz, 1H), 2.28 (ddd, *J* = 14.8, 11.5, 3.3 Hz, 1H), 1.95 (dt, *J* = 13.8, 4.2 Hz, 1H), 1.92 – 1.83 (m, 2H), 1.70 (ddd, *J* = 14.6, 10.0, 5.2 Hz, 1H), 1.45 (d, *J* = 6.8 Hz, 3H), 1.36 (dtdd, *J* = 25.8, 18.0, 13.5, 7.1 Hz, 4H), 1.25 – 1.17 (m, 1H), 1.11 – 1.01 (m, 2H), 0.89 – 0.77 (m, 2H), 0.54 (p, *J* = 7.0 Hz, 1H). ^13^C NMR (151 MHz, Acetone) δ 181.77 (d, *J* = 5.5 Hz), 165.87 (d, *J* = 22.6 Hz), 135.05, 134.18, 127.55, 125.09, 121.64, 114.62, 110.64, 105.99, 97.24, 93.84, 71.23 (d, *J* = 4.3 Hz), 60.92, 46.32, 38.27, 33.56, 28.70, 28.19, 27.26, 25.87, 25.23 (2 carbons), 24.82, 20.92 (d, *J* = 4.3 Hz). LC-MS: calculated for C_25_H_33_N_3_O_2_ [M+H^+^] 408.265 m/z; found 408.265 m/z.

### Acid-catalyzed electrophilic aromatic substitution-based derivatization general procedure

(−)-Premarineosin A was dissolved in acetonitrile (MeCN) before trifluoroacetic acid (TFA) and a carbonyl-containing substrate was added at RT. The reaction was run until TLC (50% ethyl acetate in hexanes with 1% acetic acid) showed consumption of starting material (15 min - 6 h). After completion, the reaction was diluted with ethyl acetate and washed with 1M NaOH to quench the TFA. The aqueous layer was further extracted with ethyl acetate (x3) and organics were combined, dried over sodium sulfate, filtered, and concentrated. The material was purified through a combination of acidic and/or basic conditions. Acidic conditions: 50% ethyl acetate (with 1% acetic acid) and hexanes (with 1% acetic acid). Basic conditions: 20-50% of a (3% ammonia (7N in methanol) in DCM solution) in hexane gradient. (See SI for detailed information and NMR data.)

### D3 and RebH Docking Studies

The amino acid sequence of the flavin-dependent halogenase D3 was obtained from the Jared Lewis Lab (Indiana University) (UniProt Accession Number: Q21N77). The sequence was used as input for AlphaFold2 (version 2.2.0) to generate a predicted three-dimensional structure of D3. The default parameters and model configuration provided by the AlphaFold2 pipeline were utilized, with a focus on the most confident prediction based on the pLDDT score. Five models were generated, and the one with the highest overall pLDDT score was selected for subsequent studies. The crystal structure of the D3 homolog RebH (PDB ID: 2OA1) was retrieved from the Protein Data Bank. This structure was prepared for docking by removing water molecules, heteroatoms, and non-relevant ligands using PyMOL (version 2.5.0) and AutoDockTools (version 1.5.7). The substrate, premarineosin, was drawn and minimized using ChemDraw (version 21.0) and Chem3D (version 21.0). The minimized structure was exported in PDB format and converted to PDBQT format using AutoDockTools. Protonation states of premarineosin were assigned at pH 7.4, and torsional bonds were defined to enable flexible docking. Docking simulations were performed using AutoDock Vina (version 1.1.2). Both the AlphaFold-predicted D3 structure and the crystal structure of RebH were prepared for docking by adding polar hydrogens, assigning Kollman charges, and converting the files to PDBQT format using AutoDockTools. The binding site of RebH was identified based on the position of the co-crystallized substrate or ligand. For the AlphaFold model of D3, the putative binding site was identified using structural alignment with RebH and manual inspection of conserved active site residues. A grid box was defined to encompass the putative binding pocket with dimensions of 20 × 20 × 20 Å, centered on the active site. For both D3 and RebH, docking simulations were run with an exhaustiveness parameter of 8 to ensure thorough sampling of conformational space. The docking scores were recorded in kcal/mol, and the best-ranked poses were selected based on binding energy and alignment with the expected binding mode. The top docking poses were visualized using PyMOL to assess binding orientation and interactions with active site residues. Comparative analyses of the docking poses in D3 and RebH were conducted to infer similarities and differences in substrate binding.

### Expression and Purification of D3 Halogenase

For large-scale D3 halogenase (Q21N77_SACD2) production^63^, 10 mL culture tubes containing 5 mL LB with kanamycin were inoculated with BL-21(λDE3) E. coli cells harboring pET28b containing the D3 halogenase gene. The cultures were incubated overnight at 37 °C, 250 rpm. The next day, 750 mL TB supplemented with kanamycin was inoculated with the entire overnight culture. The inoculated expression cultures were incubated at 37 °C, 225 rpm until OD600 was between 0.6 and 0.8. The incubator was allowed to cool to 30 °C, and gene expression was induced with 100 μM IPTG, and the expression culture was incubated for 20 hours. Cells were harvested by centrifugation at 6000×g at 4°C for 15 min, and the cell pellet was stored at -80 °C until purification of the protein. Cells were lysed via sonication with a total processing time of 5 minutes with 1 minute on/off cycles. The cell lysate was centrifugated (60,000×g for 25 min), and the resulting clarified lysate was transferred to a fresh 50 mL centrifuge tube and added to pre-equilibrated Ni-NTA (equilibration buffer: 20 mM phosphate, 300 mM NaCl, 10 mM imidazole pH 7.4). Clarified lysate was incubated with resin for approximately 30 minutes, at which point it was transferred to uncapped spin columns, and the lysate was allowed to flow through. The resin was washed with at least 5 CV wash buffer (20 mM phosphate, 300 mM NaCl, 25 mM imidazole pH 7.4), at which point the spin column was transferred to a new centrifuge tube, and the resin was washed with elution buffer (20 mM phosphate, 300 mM NaCl, 250 mM imidazole, pH 7.4). Eluted protein was concentrated via diafiltration using Amicon spin filters Ultra 30K MWCO spin filters and the buffer was exchanged for storage buffer (25 mM HEPES and 10% glycerol, pH 7.4). For long-term storage, proteins were immediately frozen in liquid nitrogen and stored at – 80 °C until use.

### Expression and Purification of HpaC Flavin Reductase

For large-scale HpaC production, 10 mL culture tubes containing 5 mL LB with ampicillin were inoculated with BL-21(λDE3) pRare E. coli cells harboring pET21a containing the HpaC halogenase gene. The HpaC flavin reductase (phaC plasmid) was obtained from Prof. David Ballou (University of Michigan). The cultures were incubated overnight at 37 °C, 250 rpm. The next day, 750 mL TB with ampicillin was inoculated with the entire overnight culture. The inoculated expression cultures were incubated at 37 °C, 225 rpm until OD600 was between 0.6 and 0.8. The incubator was allowed to cool to 20 °C, and gene expression was induced with 100 μM IPTG, and the expression culture was incubated for 20 hours. Cells were harvested by centrifugation at 6000 ×g at 4°C for 15 min and the cell pellet was stored at -80 °C until purification of the protein. Cells were lysed via sonication with a total processing time of 5 minutes with 1 minute on/off cycles. The cell lysate was centrifugated (60,000 ×g for 25 min), and the resulting clarified lysate was transferred to a fresh 50 mL centrifuge tube and added to pre-equilibrated Ni-NTA (equilibration buffer: 20 mM HEPES, 300 mM NaCl, 50 μM FAD, 10% glycerol pH 7.4). Clarified lysate was incubated with resin for approximately 30 minutes, at which point it was transferred to uncapped spin columns, and the lysate was allowed to flow through. The resin was washed with at least 5 CV wash buffer (20 mM HEPES, 300 mM NaCl, 50 mM imidazole, 10% glycerol pH 7.4), at which point the spin column was transferred to a new centrifuge tube, and the resin was washed with elution buffer (20 mM HEPES, 300 mM NaCl, 500 mM imidazole, 10% glycerol pH 7.4). Eluted protein was concentrated via diafiltration using Amicon spin filters Ultra 30K MWCO spin filters and the buffer was exchanged for storage buffer (20 mM HEPES, 100 mM NaCl, 10% glycerol pH 7.4). For long-term storage, proteins were immediately frozen in liquid nitrogen and stored at – 80 °C until use.

### D3 Halogenase reaction conditions

Analytical reactions were performed using 20 μM D3, 45 μM HPAC flavin reductase, 100 μM FAD, 50 mM sodium bromide, and 400 μM of (−)-premarineosin A dissolved in DMSO was added and then diluted to a final volume of 250 μL with reaction buffer (10 mM HEPES, pH 7.4 containing 10% glycerol) and initiated by adding NADH (1 mM). Two control reactions were performed which included all contents except D3. Reactions were incubated at 30 °C for 4 h agitating at 600 rpm in a thermoshaker (Multithermoshaker, Benchmark) and quenched via addition of 750 μL of methanol, followed by vortexing at full speed S3 for 30 s. Quenched reactions were centrifuged at 17,000 ×g to remove insoluble material, and the supernatant was analyzed with chromatographic conditions identical to analytical reactions and were performed using an Agilent G6230B time-of-flight (TOF) mass spectrometer system operating in positive mode, monitoring a mass range of 200 to 1200 amu with ESI-MS, and UV (195−400 nm) detection. ESI conditions were set with the capillary temperature at 320 °C, source voltage at 3.5 kV, and a sheath gas flow rate of 11 L/min, and the first 1 min of flow was diverted to waste. Preparatory reactions were performed using 40 μM D3, 50 μM HPAC flavin reductase, 100 μM FAD, 50 mM sodium bromide, and 500 μM of (−)-premarineosin A dissolved in DMSO at a final volume of 5 mL in reaction buffer (10 mM HEPES, pH 7.4 containing 10% glycerol). The reaction was initiated upon the addition of 5 M NADH. The reaction was quenched via the addition of 20 mL of HPLC-grade methanol, vortexed on the highest setting for 20 s, then water bath sonicated for 30 s. The resulting mixture from all combined 5 mL reactions was then passed through a pad of Celite© and the filter cake was washed with an additional 20 mL of methanol. To purify the biocatalytic byproduct, the aqueous solution was extracted with DCM. The organics were combined and concentrated. The resulting material was further purified by preparative HPLC with a phenyl hexyl column (5 μm, 100 Å, 250 × 10 mm) using a gradient of 10-55% of acetonitrile and water, both modified with 0.1% formic acid, over 50 min at a flow rate of 5 mL/min. Products were analyzed using LC-MS/MS (**Supplementary Fig. 10**), HPLC, and NMR (**Supplementary Fig. 56-60**).

### NBS Chemical Bromination

To a solution of (−)-premarineosin A (13.8 mg, 5 mmol) in dichloromethane at 4 °C was added *N-*Bromo succinimide (10 mg, 10 mmol, DCM) drop wise. The mixture was stirred at 4 °C. After 2 h the reaction was quenched with sodium bicarbonate. Following a liquid-liquid extraction with DCM, the organic layer was extracted and dried. The resulting material was further purified by preparative HPLC with a phenyl hexyl column (5 μm, 100 Å, 250 × 10 mm) using a gradient of 10-55% of acetonitrile and water, both modified with 0.1% formic acid, over 50 min at a flow rate of 5 mL/min. Products were analyzed using LC-MS/MS (**Supplementary Fig. 10**), HPLC, and NMR (**Supplementary Fig. 61-66**).

### Mammalian Cell Lines and Cell Culture

Cells were obtained from the following sources: HEK293 (ATCC, cat # CRL-1573), MOLT-4 (ATCC, cat # CRL-1582). Cell lines were routinely tested for Mycoplasma contamination with the MycoAlert PLUS *Mycoplasma* Detection Kit (Lonza Bioscience, cat # LT07) according to manufacturer protocol. Cell number and cell viability were measured using the Countess Automated Cell Counter (Invitrogen) using Trypan blue stain (Invitrogen, cat # T10282). Human embryonic kidney HEK293 cells were maintained in DMEM with high glucose and GlutaMAX (Gibco, cat # 10566016) supplemented with 10% HyClone Characterized Fetal Bovine Serum (Cytiva, cat # SH30071.03), and 100U/mL Penicillin-Streptomycin (Gibco, cat # 15140122). Human T lymphoblast MOLT-4 cells were maintained in RPMI 1640 Medium (Gibco, cat # 11875093) supplemented with 10% HyClone Characterized Fetal Bovine Serum (Cytiva, cat # SH30071.03), 100U/mL Penicillin-Streptomycin (Gibco, cat # 15140122), and 10 mM HEPES (Gibco, 15630080). Cells were cultured in incubators maintained at 37^°^C with 5% CO_2_ and 95% relative humidity.

### CellTiter-Glo Cell Viability Assay (CTG) and Data Analysis

The CTG assay (Promega, cat # G7572) was used to quantify cellular ATP according to manufacturer protocols. The Multidrop Combi liquid dispenser (ThermoFisher) was used to dispense 4 μL of cells into white 1536-well microplates (Corning, cat # 7464) at a density of 1500 cells/well. Plated cells were incubated overnight. (−)-Premarineosin A and derivatives were transferred into respective wells of each plate at 25 nL/well with the mosquito (SPT Labtech) in 16-pt, 1:3 titrations for a final concentration of 125 uM – 8.71 pM for compounds with a high concentration of 20 mM, or 62.5 uM – 4.36 pM for compounds with a high concentration of 10 mM. Controls were transferred into respective wells of each plate at 25 nL/well with the mosquito with a 16-pt, 1:2 titration of Digitonin for a final concentration of 125 uM – 3.81 nM. The final concentration of DMSO was 0.58%. Cells were incubated for either 24, 48, and 72 h, after which point 3 μL/well of CTG was transferred to each plate with the BioRaptr 2.0 FRD (LetsGoRobotics). Plates were incubated for ten minutes in the dark at ambient temperature for ten minutes. Luminescence was measured using a ViewLux 1430Ultra HTS (PerkinElmer) with the following optical settings: exposure = 1 s, gain = high, speed = slow, binning = 2X. Dimethyl sulfoxide (“DMSO”; AMRESCO, cat # RGE-3070) was used as a vehicle control. Digitonin (Sigma-Aldrich, cat # D141) was prepared as a 20 mM stock solution in DMSO and stored at -30 °C for use as a cytotoxicity control. (−)-Premarineosin A and derivatives were prepared at either 10 mM or 20 mM stock solutions in DMSO. Data were normalized to 125 uM Digitonin as -100% inhibition in a row-wise manner across the plate. Normalization was performed in Excel (Microsoft) and Concentration Response Curves (CRCs) were plotted in GraphPad Prism (GraphPad Software, Inc.) with error bars representing the standard deviation of two replicate wells (**Supplementary Fig. 8**).

### *In vitro* antimalarial assay

The antimalarial growth inhibition assay was performed as described previously^64^. Briefly, the *Plasmodium falciparum* Dd2 and 3D7 parasites were diluted to 0.75% parasitemia with 2% hematocrit, and 50 μL diluted parasites were added to each well in a 96-well plate containing 50 μL of properly diluted drugs. Tested compounds were three-fold diluted in triplicate with concentrations ranging from 20 – 0.001 μM/10 – 0.0005 μM. The parasites were incubated with the drug at 37 °C under mixed gas (5% O_2_, 5% CO_2_, and 90% N_2_) conditions for 48 h. DNA was released from the cultured parasite and stained with SYBR green dye. The plate was placed in the dark with gentle agitation for 1 h, and signals were read in a FLUOstar OPTIMA reader (BMG Labtech, Germany). Data from the microplate reader were analyzed as described previously^64^ and plotted using Prism 9.0 software (GraphPad Software, Inc., San Diego, CA). Each *in vitro* experiment was performed in triplicate wells and repeated twice. Response Curves (CRCs) were plotted in GraphPad Prism (GraphPad Software, Inc.) with error bars representing the standard deviation of three replicate wells (**Supplementary Fig. 9**).

### Statistical Information

Statistical analysis was performed with Prism Graphpad version 10.4.2 unless otherwise described. Specific details on statistical tests are provided in figure legends and relevant methods.

## Data Availability

Crystal structure data has been deposited in the Cambridge Crystallographic Data Centre under deposition numbers 2455429 for (−)-premarineosin A and 2455430 for gem-dimethyl-bridged premarineosin A. Other datasets generated during and/or analyzed during this study are available from the corresponding authors upon reasonable request.

## Acknowledgements

We thank Prof. William Fenical (Scripps Institute of Oceanography, San Diego, CA) for helpful conversations and for gifting us the *Streptomyces* sp. CNQ-617 strain. We are grateful for support from NIH grant R35GM118101 and the Hans W. Vahlteich Professorship (to D.H.S.), start-up funding provided by the University of Michigan Life Sciences Institute (to F.P.), a National Science Foundation Graduate Research Fellowship #DGE2241144 (to C.M.M.), an NIH F-31 #5F31DA055451-03 (to N.R.H.), the University of Michigan College of Pharmacy Duellman Graduate Student Research Fund #PGG030232 (to S.A.), a Michigan Pioneer Fellowship (to B.J.C.), and an NIH F-31 #1F31AI186432-01 (to K.L.L.). This research was also supported in part by the Intramural Research Programs of the National Center for Advancing Translational Sciences, NIH under project 1ZIA TR000495-01 (to J.I.) and the Division of Intramural Research of the National Institute of Allergy and Infectious Diseases, NIH under project # ZIA AI000892-23 (to Xz.S). This work was also supported by the University of Michigan BioNMR Core Facility (U-M BioNMR). The U-M BioNMR Core is grateful for support from U-M including the College of Literature, Sciences and Arts, the Life Sciences Institute, the College of Pharmacy, and the U-M Biosciences Initiative.

## Author Contributions

C.M.M. performed experiments, analyzed data, and wrote the manuscript. M.M. designed experiments, performed experiments, analyzed data, and wrote the manuscript. N.R.H. performed experiments, analyzed data, and wrote the manuscript. S.A. performed experiments, analyzed data. B.J.C. analyzed data. L.VH. performed experiments. A.A.T., E.N.O., K.L., P.D., F.Q., performed experiments, analyzed data. H.M.S. performed experiments. J.C.L. supervised the project. J.I. analyzed data, supervised the project, designed experiments, provided feedback on the manuscript. X.S. supervised the project, designed experiments, provided feedback on the manuscript. F.P. conceived the project, designed experiments, performed experiments, analyzed data, supervised the project, and wrote the manuscript. D.H.S. conceived the project, designed experiments, supervised the project, and wrote the manuscript. All authors approved the final version of the manuscript.

## Materials & Correspondence

Correspondence should be addressed to Filipa Pereira, Ph.D. and David H. Sherman, Ph.D.

## References

1. Geneva: World Health Organization. World Malaria Report 2023. (2023).

2. Dereje, N. et al. Resurgence of malaria and artemisinin resistance in Africa requires a concerted response. Nat. Med. 1–2 (2025) doi:10.1038/s41591-024-03439-z.

3. CDC. Malaria’s Impact Worldwide. Malaria https://www.cdc.gov/malaria/php/impact/index.html (2024).

4. Nayak, S. et al. Population genomics and transcriptomics of Plasmodium falciparum in Cambodia and Vietnam uncover key components of the artemisinin resistance genetic background. Nat. Commun. 15, 10625 (2024).

5. Hu, D. X., Withall, D. M., Challis, G. L. & Thomson, R. J. Structure, chemical synthesis, and biosynthesis of prodiginine natural products. Chem. Rev. 116, 7818 (2016).

6. Kancharla, P., Lu, W., Salem, S. M., Kelly, J. X. & Reynolds, K. A. Stereospecific synthesis of 23-hydroxyundecylprodiginines and analogues and conversion to antimalarial premarineosins via a Rieske oxygenase catalyzed bicyclization. J. Org. Chem. 79, 11674– 11689 (2014).

7. Kancharla, P., Kelly, J. X. & Reynolds, K. A. Synthesis and structure–activity relationships of tambjamines and B-ring functionalized prodiginines as potent antimalarials. J. Med. Chem. 58, 7286 (2015).

8. Kumar, A. et al. Optimization of B-ring-functionalized antimalarial tambjamines and prodiginines. J. Med. Chem. 67, 19755–19776 (2024).

9. Papireddy, K. et al. Antimalarial activity of natural and synthetic prodiginines. J. Med. Chem. 54, 5296–5306 (2011).

10. Kancharla, P. et al. Discovery and optimization of tambjamines as a novel class of antileishmanial agents. J. Med. Chem. 67, 8323–8345 (2024).

11. Kancharla, P. et al. Total synthesis and antimalarial activity of 2-(p-Hydroxybenzyl)-prodigiosins, isoheptylprodigiosin, and geometric isomers of tambjamine MYP1 isolated from marine bacteria. J. Med. Chem. 64, 8739–8754 (2021).

12. Salem, S. M. et al. Elucidation of final steps of the marineosins biosynthetic pathway through identification and characterization of the corresponding gene cluster. J. Am. Chem. Soc. 136, 4565–4574 (2014).

13. Anwar, M. M., Albanese, C., Hamdy, N. M. & Sultan, A. S. Rise of the natural red pigment ‘prodigiosin’ as an immunomodulator in cancer. Cancer Cell Int. 22, 419 (2022).

14. Williamson, N. R. et al. Anticancer and immunosuppressive properties of bacterial prodiginines. Future Microbiol. 2, 605–618 (2007).

15. Klein, A. S. et al. Preparation of cyclic prodiginines by mutasynthesis in Pseudomonas putida KT2440. ChemBioChem 19, 1545–1552 (2018).

16. Alihosseini, F., Ju, K.-S., Lango, J., Hammock, B. D. & Sun, G. Antibacterial colorants: Characterization of prodiginines and their applications on textile materials. Biotechnol. Prog. 24, 742–747 (2008).

17. You, Z. et al. Insights into the anti-infective properties of prodiginines. Appl. Microbiol. Biotechnol. 103, 2873–2887 (2019).

18. de Rond, T. et al. Oxidative cyclization of prodigiosin by an alkylglycerol monooxygenase-like enzyme. Nat. Chem. Biol. 13, 1155–1157 (2017).

19. Jones, B. T., Hu, D. X., Savoie, B. M. & Thomson, R. J. Elimination of butylcycloheptylprodigiosin as a known natural product inspired by an evolutionary hypothesis for cyclic prodigiosin biosynthesis. J. Nat. Prod. 76, (2013).

20. Boonlarppradab, C., Kauffman, C. A., Jensen, P. R. & Fenical, W. Marineosins A and B, cytotoxic spiroaminals from a marine-derived actinomycete. Org. Lett. 10, 5505–5508 (2008).

21. Barnes, E. C., Kumar, R. & Davis, R. A. The use of isolated natural products as scaffolds for the generation of chemically diverse screening libraries for drug discovery. Nat. Prod. Rep. 33, 372–381 (2016).

22. Majhi, S. & Das, D. Chemical derivatization of natural products: Semisynthesis and pharmacological aspects-A decade update. Tetrahedron 78, 131801 (2021).

23. Camp, D., Davis, R. A., Evans-Illidge, E. A. & Quinn, R. J. Guiding principles for natural product drug discovery. Future Med. Chem. 4, 1067–1084 (2012).

24. Camp, D., Garavelas, A. & Campitelli, M. Analysis of physicochemical properties for drugs of natural origin. J. Nat. Prod. 78, 1370–1382 (2015).

25. Pascolutti, M. & Quinn, R. J. Natural products as lead structures: chemical transformations to create lead-like libraries. Drug Discov. Today 19, 215–221 (2014).

26. Feng, Z., Allred, T. K., Hurlow, E. E. & Harran, P. G. Anomalous chromophore disruption enables an eight-step synthesis and stereochemical reassignment of (+)-marineosin A. J. Am. Chem. Soc. 141, 2274–2278 (2019).

27. Xu, B., Li, G., Li, J. & Shi, Y. Total synthesis of the proposed structure of marineosin A. Org. Lett. 18, 2028–2031 (2016).

28. Hong, B., Luo, T. & Lei, X. Late-stage diversification of natural products. ACS Cent. Sci. 6, 622–635 (2020).

29. Huo, T. et al. Late-stage modification of bioactive compounds: Improving druggability through efficient molecular editing. Acta Pharm. Sin. B 14, 1030–1076 (2024).

30. Pereira, F. et al. Optimized production of concanamycins using a rational metabolic engineering strategy. Metab. Eng. 88, 63–76 (2025).

31. Yang, D., Eun, H., Prabowo, C. P. S., Cho, S. & Lee, S. Y. Metabolic and cellular engineering for the production of natural products. Curr. Opin. Biotechnol. 77, 102760 (2022).

32. Liu, P., Zhu, H., Zheng, G., Jiang, W. & Lu, Y. Metabolic engineering of Streptomyces coelicolor for enhanced prodigiosins (RED) production. Sci. China Life Sci. 60, 948–957 (2017).

33. Pan, X. et al. Improving prodigiosin production by transcription factor engineering and promoter engineering in Serratia marcescens. Front. Microbiol. 13, (2022).

34. Alzahrani, N. H., El-Bondkly, A. A. M., El-Gendy, M.M.A.A. & El-Bondkly, A.M. Enhancement of undecylprodigiosin production from marine endophytic recombinant strain Streptomyces sp. ALAA-R20 through low-cost induction strategy. J. Appl. Genet. 62, 165– 182 (2021).

35. Bhatia, S. K. et al. Medium engineering for enhanced production of undecylprodigiosin antibiotic in Streptomyces coelicolor using oil palm biomass hydrolysate as a carbon source. Bioresour. Technol. 217, 141–149 (2016).

36. Stankovic, N. et al. Streptomyces sp. JS520 produces exceptionally high quantities of undecylprodigiosin with antibacterial, antioxidative, and UV-protective properties. Appl. Microbiol. Biotechnol. 96, 1217–1231 (2012).

37. Zdouc, M. M. et al. MIBiG 4.0: advancing biosynthetic gene cluster curation through global collaboration. Nucleic Acids Res. 53, D678–D690 (2025).

38. Takano, E. et al. Transcriptional regulation of the redD transcriptional activator gene accounts for growth-phase-dependent production of the antibiotic undecylprodigiosin in Streptomyces coelicolor A3(2). Mol. Microbiol. 6, 2797–2804 (1992).

39. Malpartida, F., Niemi, J., Navarrete, R. & Hopwood, D. A. Cloning and expression in a heterologous host of the complete set of genes for biosynthesis of the Streptomyces coelicolor antibiotic undecylprodigiosin. Gene 93, 91–99 (1990).

40. Malpartida, F. & Hopwood, D. A. Molecular cloning of the whole biosynthetic pathway of a Streptomyces antibiotic and its expression in a heterologous host. Nature 309, 462–464 (1984).

41. Ferraiuolo, S. B., Cammarota, M., Schiraldi, C. & Restaino, O. F. Streptomycetes as platform for biotechnological production processes of drugs. Appl. Microbiol. Biotechnol. 105, 551 (2021).

42. Evangelista-Martínez, Z. et al. Antibacterial activity of Streptomyces sp. Y15 against pathogenic bacteria and evaluation of culture media for antibiotic production. TIP Rev. Esp. Cienc. Quím. Biol.biológicas 25, (2022).

43. Gilchrist, C. L. M. & Chooi, Y.-H. clinker & clustermap.js: automatic generation of gene cluster comparison figures. Bioinformatics 37, 2473–2475 (2021).

44. Fan, Y. et al. Methylene-bridged dimeric natural products involving one-carbon unit in biosynthesis. Nat. Prod. Rep. 39, 1305–1324 (2022).

45. Li, L. et al. Exploring diversity through dimerization in natural products by a rational tandem mass-based molecular network strategy. Org. Lett. 25, 4016–4021 (2023).

46. White, J. D. The condensation of acetone with 2,5-disubstituted pyrroles. Chem. Commun. (London) 711–712 (1966) doi:10.1039/C19660000711.

47. Sleebs, B. E. et al. Transition state mimetics of the Plasmodium export element are potent inhibitors of plasmepsin V from P. falciparum and P. vivax. J. Med. Chem. 57, 7644–7662 (2014).

48. Jiang, X. et al. Structural basis for blocking sugar uptake into the malaria parasite Plasmodium falciparum. Cell 183, 258-268.e12 (2020).

49. Lewis, J. C. Identifying and engineering flavin dependent halogenases for selective biocatalysis. Acc. Chem. Res. 57, 2067–2079 (2024).

50. Yeh, E., Garneau, S. & Walsh, C. T. Robust in vitro activity of RebF and RebH, a two-component reductase/halogenase, generating 7-chlorotryptophan during rebeccamycin biosynthesis. Proc. Natl. Acad. Sci. U.S.A. 102, 3960–3965 (2005).

51. Li, H.-J. et al. Regioselective electrophilic aromatic bromination: Theoretical analysis and experimental verification. Molecules 19, 3401–3416 (2014).

52. Talà, A. et al. Pirin: A novel redox-sensitive modulator of primary and secondary metabolism in Streptomyces. Metab. Eng. 48, 254–268 (2018).

53. Fayed, B., Younger, E., Taylor, G. & Smith, M. C. M. A novel Streptomyces spp. integration vector derived from the S. venezuelae phage, SV1. BMC Biotechnol. 14, 51 (2014).

54. Hunjan, M. K. et al. Recent advances in functionalization of pyrroles and their translational potential. Chem. Rec. 21, 715–780 (2021).

55. Tsuchimoto, T. Selective synthesis of β-alkylpyrroles. Chem. Eur. J. 17, 4064–4075 (2011).

56. Jenkins, S., D. Incarvito, C., Parr, J. & H. Wasserman, H. Structural studies of C-ring substituted unnatural analogues of prodigiosin. CrystEngComm 11, 242–245 (2009).

57. Regourd, J., Al-Sheikh Ali, A. & Thompson, A. Synthesis and anti-cancer activity of C-ring-functionalized prodigiosin analogues. J. Med. Chem. 50, 1528–1536 (2007).

58. Rastogi, S. et al. Synthetic prodigiosenes and the influence of C-ring substitution on DNA cleavage, transmembrane chloride transport and basicity. Org. Biomol. Chem. 11, 3834– 3845 (2013).

59. Peh, G. et al. Site-selective chlorination of pyrrolic heterocycles by flavin dependent enzyme PrnC. Commun. Chem. 7, 1–9 (2024).

60. Mendez, L., Henriquez, G., Sirimulla, S. & Narayan, M. Looking back, looking forward at halogen bonding in drug discovery. Molecules 22, 1397 (2017).

61. Wilcken, R., Zimmermann, M. O., Lange, A., Joerger, A. C. & Boeckler, F. M. Principles and applications of halogen bonding in medicinal chemistry and chemical biology. J. Med. Chem. 56, 1363–1388 (2013).

62. Borodina, I. et al. Antibiotic overproduction in Streptomyces coelicolor A3(2) mediated by phosphofructokinase deletion *. J. Biol. Chem. 283, 25186–25199 (2008).

63. Snodgrass, H. M. Methods and applications for the identification and directed evolution of biocatalysts for chemical synthesis. (The University of Chicago, United States --Illinois, 2023).

64. Liu, S., Mu, J., Jiang, H. & Su, X. Effects of Plasmodium falciparum mixed infections on in vitro antimalarial drug tests and genotyping. Am. J. Trop. Med. Hyg. 79, 178–184 (2008)

