## Supplementary Information for "Metabolic engineering and late-stage functionalization expand the chemical space of the antimalarial premarineosin A"

**Supplementary Methods 2. Synthetic Chemistry Methods** .......... **5-9**
**Supplementary Figure 18-23.** NMR Spectra for gem-dimethyl-bridged premarineosin A (**3**) ... 25-30
**Supplementary Figure 24-29.** NMR Spectra for methylene-bridged premarineosin A (**4**) .......... 31-36
**Supplementary Figure 30-35.** NMR Spectra for trifluoromethyl-bridged premarineosin A (**5**) .. 37-42
**Supplementary Figure 36-41.** NMR Spectra for 12-trifluoropropanol premarineosin A (**6**) .......... 43-48
**Supplementary Figure 42-46.** NMR Spectra for 12-formyl premarineosin A (**7**) .......... 49-53

**Supplementary Figure 47-50.** NMR Spectra for 12-acetyl premarineosin A (**8**) ....................... 54-57
**Supplementary Figure 51-55.** NMR Spectra for 12-hydroxymethyl premarineosin A (**9**).......... 58-62
**Supplementary Figure 56-60.** NMR Spectra for 1-bromo premarineosin A (**10**)....................... 63-67
**Supplementary Figure 61-66.** NMR Spectra for 12-bromo premarineosin A (**11**)..................... 68-73
**Supplementary Figure 67.** $^1$H NMR Spectrum of marineosin A ..................................................... 74
**Supplementary X-Ray Crystallography Data**.................................................................................. **75-87**
**Supplementary Data 1.** X-Ray Crystal Structure Determination for (-)-premarineosin A .......... 75-80
**Supplementary Data 2.** X-Ray Crystal Structure Determination for Gem-dimethyl-bridged premarineosin A ..................................................................................................................... 81-87
**Supplementary References**............................................................................................................ **88**

#### Supplementary Methods.

**Supplementary Methods 1.** Untargeted Metabolomics Methods.

**Sample preparation for untargeted metabolomics.** Intracellular metabolite extracts were used for untargeted metabolomics. The culture medium was extracted as a control and run alongside experimental samples to account for sampling and laboratory contamination. Reagent blanks (MeOH) were prepared and analyzed to account for extraneous signals occurring from chemicals and analytical systems. Three biological replicates were prepared for each strain to demonstrate reproducibility.

**UHPLC-QTOF-MS/MS profiling of crude extracts.** Metabolite extracts were analyzed using ultra-high-performance liquid chromatography coupled with quadrupole time-of-flight mass spectrometry (UHPLC-LCMS) performed with an Agilent 1290 Infinity II UHPLC coupled to an Agilent 6545 ESI-Q-TOF-MS. The samples were injected, and data was collected using auto-MS/MS in positive mode. The chromatographic separation was carried out on a Phenomenex Kinetex Phenyl-Hexyl column (1.7 µm, 2.1 × 100 mm). Isocratic elution was performed with 90% solvent A (water + 0.1% formic acid) for 1 min, followed by a 9 min linear gradient to 100% solvent B (95% acetonitrile + 5% water + 0.1% formic acid). Electrospray ionization was performed at a capillary temperature of 320 °C, with a source voltage of 3.5 kV, and a sheath gas flow rate of 11 L/min. Ion fragmentation utilized ramped collision energy (5 x m/z/100 + 10 eV) for up to nine selected precursor ions per cycle. Purine $C_5H_4N_4$ $[M + H]^+$ (*m/z* 121.0508) and hexakis(1H,1H,3H-tetrafluoropropoxy)-phosphazene $C_{18}H_{18}F_{24}N_3O_6P_3$ $[M + H]^+$ (*m/z* 922.0098) served as internal lock standards, and these ions were added to the static exclusion list. This exclusion list was refined by identifying high-intensity contaminant ions (>1000 counts) present in blank solvent samples.

**Metabolomic data processing.** The raw mass spectrometry data was converted from vendor-specific instrument files to mzML format using the msConvert software from ProteoWizard[1] with standard peak detection settings. Subsequent data processing was conducted in MZmine3[2] using an untargeted LC-MS/MS preprocessing workflow from the Processing Wizard. HPLC retention time was selected between 0.4 and 9 min, allowing for a maximum of 15 peaks in the chromatogram and requiring at least four consecutive scans. The full width at half maximum (FWHM) for features was set at approximately 0.045 min. Retention time (RT) tolerance was defined as 0.08 min for intra-sample variations and 0.4 min for sample-to-sample differences. The noise levels were set at $5.0 \times 10^2$ for MS1 and $2.0 \times 10^2$ for MS2. The minimum feature height was established at $1.0 \times 10^3$, with a scan-to-scan m/z tolerance of 0.005 m/z (20 ppm), an intra-sample m/z tolerance of 0.0015 m/z (3 ppm), and a sample-to-sample m/z tolerance of 0.004 m/z (8 ppm). The output, including the feature quantification table, peak list, and metadata table, was processed through feature-based molecular networking on the Global Natural Products Social Molecular Networking (GNPS)[3] platform. MS/MS spectra of molecular features were tentatively matched to the GNPS MS/MS Spectral Libraries if they had a cosine similarity score greater than 0.7 and shared at least 4 peaks. The resulting molecular network was visualized using Cytoscape 3.9.1.[4]

**Supplementary Methods 2.** Synthetic Chemistry Methods.

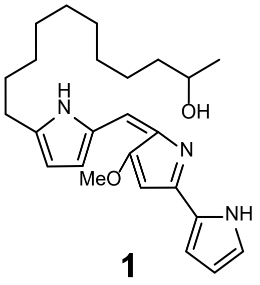

**1**

**23-Hydroxyundecylprodiginine (1).** $^1$H NMR (599 MHz, Acetone) δ 7.34 (s, 1H), 7.20 (d, J = 3.6 Hz, 1H), 7.12 (s, 1H), 7.02 (s, 1H), 6.60 (s, 1H), 6.40 (dd, J = 3.8, 2.5 Hz, 1H), 6.28 (d, J = 3.8 Hz, 1H), 4.09 (s, 3H), 3.68 (h, J = 6.1 Hz, 1H), 2.89 (t, J = 7.7 Hz, 1H), 1.78 (p, J = 7.5 Hz, 2H), 1.43 – 1.27 (m, 16H), 1.09 (d, J = 6.1 Hz, 3H). LC-MS: calculated for $C_{25}H_{35}N_3O_2$ [M+H$^+$] 410.2763 m/z; found 410.2812 m/z.

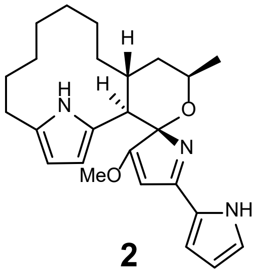

**2**

**(-)-Premarineosin A* (2)**. $^1$H NMR (599 MHz, Acetone) δ 13.65 (s, 1H), 12.05 (s, 1H), 9.80 (s, 1H), 7.65 (ddd, *J* = 3.3, 2.3, 1.3 Hz, 1H), 7.45 (s, 1H), 6.50 (dt, *J* = 4.0, 2.1 Hz, 1H), 6.26 (d, *J* = 1.7 Hz, 1H), 5.72 (t, *J* = 2.9 Hz, 1H), 5.52 (t, *J* = 2.9 Hz, 1H), 4.46 – 4.38 (m, 1H), 4.14 (s, 3H), 3.03 (d, *J* = 12.5 Hz, 1H), 2.69 (dp, *J* = 13.4, 4.9 Hz, 1H), 2.39 (dt, *J* = 14.8, 4.4 Hz, 1H), 2.28 (ddd, *J* = 14.8, 11.5, 3.3 Hz, 1H), 1.95 (dt, *J* = 13.8, 4.2 Hz, 1H), 1.92 – 1.83 (m, 2H), 1.70 (ddd, *J* = 14.6, 10.0, 5.2 Hz, 1H), 1.45 (d, *J* = 6.8 Hz, 3H), 1.36 (dtdd, *J* = 25.8, 18.0, 13.5, 7.1 Hz, 4H), 1.25 – 1.17 (m, 1H), 1.11 – 1.01 (m, 2H), 0.89 – 0.77 (m, 2H), 0.54 (p, *J* = 7.0 Hz, 1H). $^{13}$C NMR (151 MHz, Acetone) δ 181.77 (d, *J* = 5.5 Hz), 165.87 (d, *J* = 22.6 Hz), 135.05, 134.18, 127.55, 125.09, 121.64, 114.62, 110.64, 105.99, 97.24, 93.84, 71.23 (d, *J* = 4.3 Hz), 60.92, 46.32, 38.27, 33.56, 28.70, 28.19, 27.26, 25.87, 25.23 (2 carbons), 24.82, 20.92 (d, *J* = 4.3 Hz). LC-MS: calculated for $C_{25}H_{33}N_3O_2$ [M+H$^+$] 408.2651 m/z; found 408.2673 m/z.

*all nitrogens in (-)-premarineosin A are bound to a hydrogen if last treated with acid.

**Crystallization method for (-)-premarineosin A.** Compound was dissolved in minimal amounts of ethyl acetate. Vapor diffusion with hexanes was performed over the course of a day to give colorless, prism-shaped crystals.

**(-)-Premarineosin A derivatization general procedure.** (-)-Premarineosin A was dissolved in MeCN before TFA and a carbonyl was added at RT. Reaction was run until TLC (50% ethyl acetate in hexanes with 1% acetic acid) showed consumption of starting material (15 min - 6 h). After completion, the reaction was diluted with ethyl acetate and washed with 1M NaOH to quench TFA. The aqueous layer was further extracted with ethyl acetate (x3) and organics were combined, dried over sodium sulfate, filtered, and concentrated. The material then was purified through a combination of acid and/or base conditions. Acidic conditions: 50% ethyl acetate (with 1% acetic acid) and hexanes (with 1% acetic acid). Basic conditions: 20-50% of a (3% ammonia (7 M in methanol) in DCM solution) in hexanes gradient.

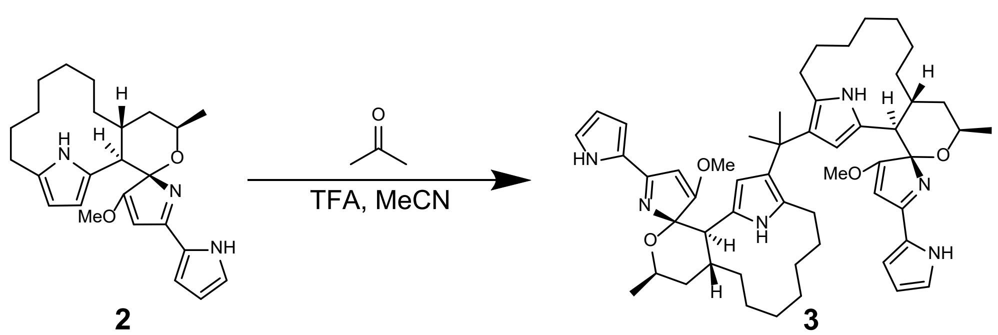

**2** **3**

**Gem-dimethyl-bridged premarineosin A (3)**. Amounts: (-)-premarineosin A (15.0 mg, 36.8 µmol, 1 equiv), MeCN (1.50 mL), acetone (100 µL, 1.36 mmol, 37 equiv), TFA (113 µL, 1.47 mmol, 40 equiv). Reaction time was 2 h. Starting material was recovered (2.0 mg, 4.9 µmol, 13%) and product (8.0 mg, 9.7 µmol, 26%). Reaction is also successful when ran neat in acetone with same equivalence of TFA and equivalent product yields. $^{1}$H NMR (599 MHz, Acetone) δ 6.95 (s, 2H), 6.45 (s, 2H), 6.13 (s, 2H), 5.53 (d, J = 3.1 Hz, 2H), 5.34 (s, 2H), 4.26 (t, J = 6.7 Hz, 2H), 3.76 (s, 6H), 2.64 (d, J = 11.9 Hz, 2H), 2.51 (s, 2H), 1.80 (d, J = 13.1 Hz, 2H), 1.64 (tt, J = 18.0, 9.7 Hz, 4H), 1.50 (d, J = 6.9 Hz, 8H), 1.39 (t, J = 12.8 Hz, 4H), 1.29 (s, 6H), 1.27 – 1.10 (m, 10H), 1.01 (tt, J = 9.8, 5.0 Hz, 2H), 0.93 (d, J = 13.8 Hz, 2H), 0.84 (s, 2H), 0.78 – 0.69 (m, 2H), 0.62 (d, J = 10.4 Hz, 2H). 13C NMR (151 MHz, Acetone) δ 181.70, 164.97, 129.10, 128.55, 127.53, 124.92, 122.99, 113.43, 109.94, 106.58, 102.40, 94.80, 70.64, 58.81, 48.28, 39.89, 33.45 – 32.90 (m, 3 different carbon signals), 30.10, 27.83, 25.86, 25.72, 25.51 (2 distinct carbon signals), 24.98, 22.98. LC-MS: calculated for $C_{53}H_{70}N_6O_4$ [M+H$^+$] 855.5537 m/z; found 855.5551 m/z.

**Crystallization method for gem-dimethyl-bridged (-)-premarineosin A (3).** Compound was dissolved in minimal amounts of ethyl acetate. Vapor diffusion with hexanes was performed over the course of two weeks to give orange, irregular crystals.

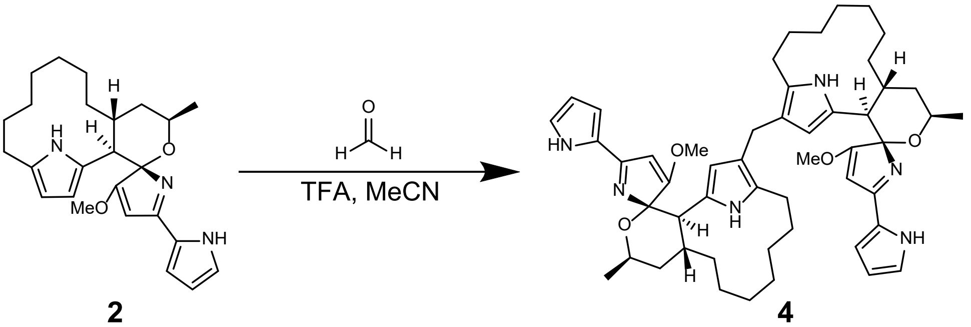

**2** **4**

**Methylene-bridged premarineosin A (4).** Amounts: (-)-premarineosin A (15.0 mg, 36.8 µmol, 1 equiv), MeCN (1.50 mL), formaldehyde (151 µL, 2.02 mmol, 55 equiv, 37% in water), TFA (113 µL, 1.47 mmol, 40 equiv). Reaction time was 10-20 minutes. Product yield: 5 mg (36.8 µmol, 20%). $^{1}$H NMR (600 MHz, Acetone) δ 7.01 (s, 2H), 6.56 (s, 2H), 6.17 (t, *J* = 3.2 Hz, 2H), 5.47 – 5.44 (m, 2H), 5.43 (s, 2H), 4.26 (p, *J* = 6.6 Hz, 2H), 3.76 (s, 6H), 3.11 (s, 2H), 2.62 (d, *J* = 11.9 Hz, 2H), 2.46 (d, *J* = 12.5 Hz, 2H), 2.37 (ddd, *J* = 14.7, 11.0, 3.5 Hz, 2H), 2.09 (dt, *J* = 13.1, 3.9 Hz, 2H), 1.86 – 1.78 (m, 2H), 1.68 – 1.54 (m, 6H), 1.49 (d, *J* = 6.9 Hz, 6H), 1.42 – 1.32 (m, 6H), 1.24 (t, *J* = 7.4 Hz, 2H), 1.20 – 1.13 (m, 2H), 1.05 (ddd, *J* = 14.3, 10.0, 5.1 Hz, 2H), 0.85 (dp, *J* = 21.8, 8.1, 7.3 Hz, 4H), 0.72 (q, *J* = 7.8, 7.1 Hz, 2H), 0.61 – 0.54 (m, 2H). $^{13}$C NMR (151 MHz, Acetone) δ 182.10, 165.29, 128.86, 127.08, 126.69, 122.91, 119.20, 113.85, 110.89, 110.16, 102.21, 94.78, 70.61, 58.98, 47.85, 39.69, 32.24, 30.36, 28.17, 26.28, 25.83, 25.45, 22.94, 22.86. LC-MS: calculated for $C_{51}H_{66}N_6O_4$ [M+H$^+$] 827.5224 m/z; found 827.5216 m/z.

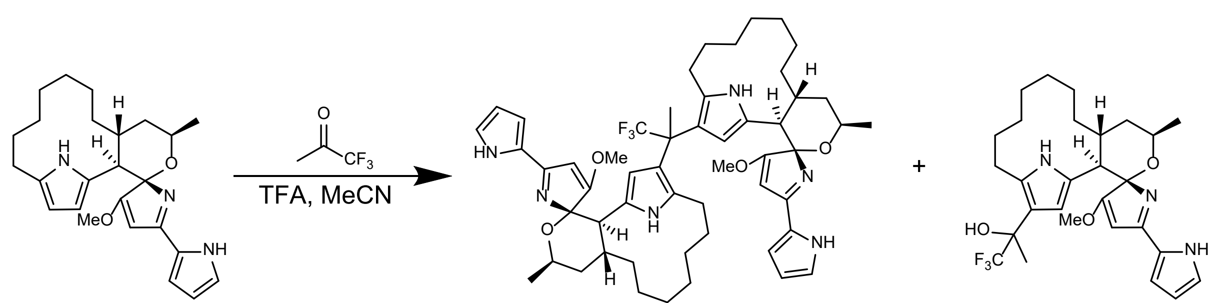

2 5 6

**Trifluoromethyl-bridged premarineosin A (5).** Amounts: (-)-premarineosin A (15.0 mg, 36.8 µmol, 1 equiv), MeCN (1.50 mL), trifluoroacetone (329 µL, 3.68 mmol, 100 equiv), TFA (170 µL, 2.21 mmol, 60 equiv). Reaction time was 4 h. Starting material was recovered (3.0 mg, 7.4 µmol, 20%) as well as two products: dimer **5** (2.0 mg, 2.2 µmol 6%) and monomer **6** (10 mg, 19 µmol, 52%). If the reaction is ran longer (6-8 h) a higher conversion rate to the dimer can be achieved. $^{1}$H NMR (600 MHz, Acetone) δ 7.53 (d, *J* = 27.8 Hz, 2H), 7.30 – 7.25 (m, 2H), 6.44 (dt, *J* = 23.3, 3.1 Hz, 2H), 6.11 (d, *J* = 24.1 Hz, 2H), 5.74 – 5.65 (m, 2H), 4.38 (ddp, *J* = 10.3, 7.0, 4.0, 3.2 Hz, 2H), 4.05 (d, *J* = 19.3 Hz, 6H), 2.91 – 2.78 (m, 4H), 2.02 – 1.93 (m, 2H), 1.92 – 1.85 (m, 2H), 1.65 (ddd, *J* = 13.3, 10.4, 5.5 Hz, 2H), 1.62 – 1.55 (m, 2H), 1.53 (s, 3H), 1.49 (t, *J* = 7.3 Hz, 6H), 1.46 – 1.24 (m, 8H), 1.20 (tt, *J* = 12.1, 6.5 Hz, 4H), 1.01 (dtq, *J* = 18.2, 9.5, 5.2, 4.2 Hz, 4H), 0.94 – 0.75 (m, 6H), 0.57 (q, *J* = 13.6, 12.1 Hz, 2H). $^{13}$C NMR (151 MHz, Acetone) δ 180.95, 168.81, 165.61 (d, *J* = 7.4 Hz), 134.19, 131.77 (d, *J* = 319.0 Hz), 125.84, 123.08, 122.84, 122.28, 120.40 (d, *J* = 250.1 Hz), 113.58 (d, *J* = 19.9 Hz), 108.83 (d, *J* = 223.9 Hz), 97.71, 93.95 (d, *J* = 8.0 Hz), 71.44 (d, *J* = 6.5 Hz), 60.42 (d, *J* = 20.3 Hz), 46.88 (d, *J* = 10.5 Hz), 44.69 (t, *J* = 25.4 Hz), 38.63 (d, *J* = 7.7 Hz), 33.56, 27.57 (d, *J* = 25.6 Hz), 25.90, 25.71, 25.50, 25.42, 25.13 – 24.49 (m), 24.33, 21.17 (d, *J* = 20.1 Hz). LC-MS: calculated for $C_{53}H_{67}F_3N_6O_4$ [M+H$^+$] 909.5254 m/z; found 909.5257 m/z.

**12-trifluoropropanol premarineosin A (6).** $^{1}$H NMR (600 MHz, CDCl3) δ 10.27 – 10.24 (m, 1H), 7.47 (d, *J* = 1.9 Hz, 1H), 7.02 (d, *J* = 4.1 Hz, 1H), 6.38 (dd, *J* = 4.0, 2.2 Hz, 1H), 5.71 (dd, *J* = 14.2, 2.7 Hz, 1H), 5.54 (d, *J* = 7.8 Hz, 1H), 4.43 (t, *J* = 6.7 Hz, 1H), 4.01 (d, *J* = 6.4 Hz, 3H), 2.92 (dd, *J* = 12.5, 3.1 Hz, 1H), 2.88 – 2.76 (m, 2H), 2.47 – 2.40 (m, 1H), 1.96 (d, *J* = 12.6, 1H), 1.91 (dt, *J* = 13.8, 3.5 Hz, 2H), 1.73 – 1.65 (m, 2H), 1.61 (d, *J* = 14.9 Hz, 3H), 1.56 (s, 1H), 1.52 (d, *J* = 6.9 Hz, 3H), 1.42 (dd, *J* = 9.1, 4.2 Hz, 2H), 1.29 (s, 1H), 1.14 (dt, *J* = 14.6, 7.1 Hz, 1H), 0.92 (ddtd, *J* = 42.7, 18.4, 12.4, 5.8 Hz, 3H), 0.56 (d, *J* = 11.0 Hz, 1H). $^{13}$C NMR (151 MHz, $CDCl_3$) δ 180.10, 179.41, 164.64 (d, *J* = 2.7 Hz), 134.31, 132.27 (d, *J* = 52.9 Hz), 125.08, 123.54, 121.65, 115.85 (d, *J* = 95.5 Hz), 113.16, 109.19 (d, *J* = 105.1 Hz), 96.55, 92.82 (d, *J* = 6.3 Hz), 71.51 (d, *J* = 14.2 Hz), 59.85, 45.52 (d, *J* = 31.0 Hz), 37.84 (d, *J* = 9.2 Hz), 32.07, 31.37, 29.85, 27.28, 27.21 – 26.78 (m), 25.05 – 24.66 (m), 24.66 – 24.20 (m), 24.05 (d, *J* = 13.3 Hz), 20.73 (d, *J* = 5.2 Hz). LC-MS: calculated for $C_{28}H_{36}F_3N_3O_3$ [M+H$^+$] 450.2757 m/z; found 450.2798 m/z.

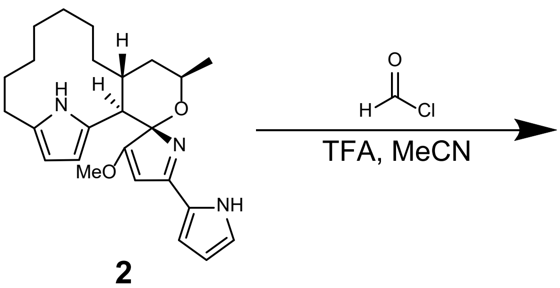

**12-formyl premarineosin A (7).** Amounts: (-)-premarineosin A (31.0 mg, 76.1 μmol), MeCN (3.10 mL), formyl chloride* (327 μL, 6.08 mmol, 80 equiv), TFA (352 μL, 4.56 mmol, 60 equiv). Reaction time was 4 h. Product yield: 32.0 mg (73.5 μmol, 97%). *Formyl chloride was made fresh by using the following procedure: To a flame dried flask and under $N_2$ thionyl chloride was added (19.0 g, 11.6 mL, 22.8 Eq, 160 mmol). Formic acid (322 mg, 264 μL, 1 Eq, 7.00 mmol) and DMF (132 mg, 140 μL, 0.258 Eq, 1.81 mmol) were added sequentially before the reaction was heated to 70 °C for 1.5 h. There should be a color change from clear to yellow to indicate the reaction is/has progressed. The excess oxalyl chloride was removed by rotoevaporation with KOH solution in the trap. Product obtained should be yellow and was used immediately in next set of reactions. $^1$H NMR (600 MHz, Acetone) δ 9.67 (d, *J* = 1.6 Hz, 1H), 7.06 (s, 1H), 6.62 (s, 1H), 6.19 (s, 1H), 6.10 (s, 1H), 5.55 (s, 1H), 4.31 (p, *J* = 6.5 Hz, 1H), 3.82 (s, 3H), 3.01 (ddd, *J* = 14.6, 10.5, 3.6 Hz, 1H), 2.77 (d, *J* = 12.0 Hz, 1H), 2.52 (d, *J* = 13.8 Hz, 1H), 2.37 – 2.31 (m, 1H), 1.92 – 1.84 (m, 2H), 1.69 (td, *J* = 12.5, 5.8 Hz, 1H), 1.62 – 1.57 (m, 1H), 1.51 (d, *J* = 6.8 Hz, 4H), 1.40 (t, *J* = 7.5 Hz, 1H), 1.36 (s, 1H), 1.33 (d, *J* = 12.2 Hz, 1H), 1.28 (d, *J* = 12.6 Hz, 1H), 1.12 (tt, *J* = 9.9, 5.1 Hz, 1H), 0.97 (dq, *J* = 29.4, 10.1, 8.6 Hz, 2H), 0.79 (p, *J* = 7.2, 6.5 Hz, 1H), 0.37 (dp, *J* = 19.5, 6.9 Hz, 1H). $^{13}$C NMR (151 MHz, Acetone) δ 184.48, 181.93, 165.69, 142.84, 131.18, 128.63, 123.48, 122.43 (d, *J* = 3.9 Hz), 114.58, 110.60, 108.89, 101.62, 95.00, 70.83, 59.40, 47.48, 39.15, 31.95, 30.45, 29.95, 27.88, 26.16 – 24.96 (m), 22.71. LC-MS: calculated for $C_{26}H_{33}N_3O_3$ [M+H$^+$] 436.2600 m/z; found 436.2636 m/z.

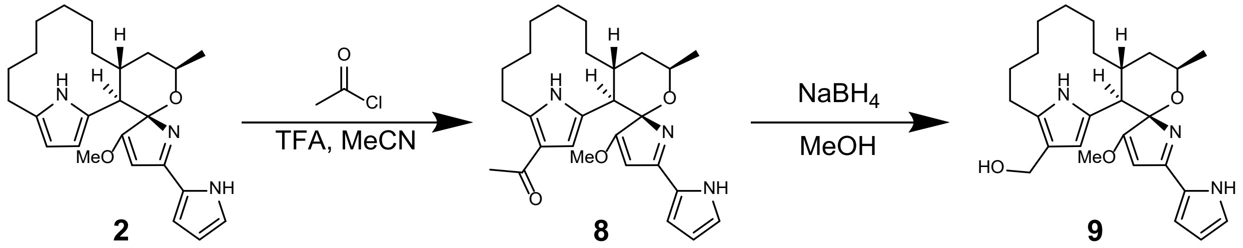

**12-acetyl premarineosin A (8).** Amounts: (-)-premarineosin A (18.0 mg, 44.2 μmol, 1 equiv), MeCN (1.80 mL), acetyl chloride (314 μL, 4.42 mmol, 100 equiv), TFA (2.04 μL, 2.65 mmol, 60 equiv). Reaction time was 4 h. Product yield: 7 mg (16 μmol, 35%). $^1$H NMR (599 MHz, Acetone) δ 7.10 (s, 1H), 6.67 (s, 1H), 6.21 (s, 1H), 6.18 (s, 1H), 5.60 (s, 1H), 4.32 (q, *J* = 6.9 Hz, 1H), 3.86 (s, 3H), 3.32 (s, 1H), 2.79 (d, *J* = 12.0 Hz, 1H), 2.55 (s, 1H), 2.18 (s, 3H), 1.91 – 1.85 (m, 1H), 1.78 (d, *J* = 7.4 Hz, 1H), 1.69 (td, *J* = 12.4, 5.8 Hz, 1H), 1.52 (d, *J* = 6.9 Hz, 3H), 1.45 – 1.32 (m, 3H), 1.32 – 1.24 (m, 3H), 1.18 – 1.08 (m, 2H), 1.03 – 0.93 (m, 2H), 0.76 (dq, *J* = 15.2, 7.2 Hz, 1H), 0.35 (hept, *J* = 6.2 Hz, 1H). $^{13}$C NMR (151 MHz, Acetone) δ 193.83, 181.79, 165.63, 139.17, 128.16 (d, *J* = 30.4 Hz), 124.41, 120.48, 117.31 – 113.69 (m), 111.85, 110.75, 101.30, 94.86, 70.77, 59.38, 47.27, 38.99, 31.74, 30.59, 30.35, 29.84, 28.49, 28.28, 27.66 (d, *J* = 6.9 Hz), 26.06, 25.73 (d, *J* = 15.3 Hz), 25.23 (d, *J* = 59.9 Hz), 22.54. LC-MS: calculated for $C_{27}H_{35}N_3O_3$ [M+H$^+$] 450.2757 m/z; found 450.2798 m/z.

**12-hydroxymethyl premarineosin A (9).** To a solution of **12-formyl premarineosin** (19.0 mg, 1 Eq, 43.6 µmol) in MeOH (690 mg, 872 µL, 494 Eq, 21.5 mmol) at 0 °C was added $NaBH_4$ (3.30 mg, 2.00 Eq, 87.2 µmol). The reaction was run until TLC showed consumption of starting material [50% (3% ammonia (7 M in methanol) in DCM solution) in hexanes]. The reaction was quenched by addition of 1M HCl at 0 °C until acidified and extracted with ethyl acetate (x3). The organics combined, dried over sodium sulfate, filtered and concentrated. The crude material was purified by pTLC using 50% (3% ammonia (7 M in methanol) in DCM solution) in hexanes to yield the product **9** (13.0 mg, 29.7 µmol, 68.1 %) as a pale yellow solid. $^1$H NMR (600 MHz, Acetone) δ 7.02 (s, 1H), 6.61 (d, *J* = 3.7 Hz, 1H), 6.18 (d, *J* = 3.5 Hz, 1H), 5.68 (t, *J* = 2.0 Hz, 1H), 5.49 (s, 1H), 4.29 – 4.21 (m, 3H), 3.78 (s, 3H), 2.69 (d, *J* = 11.8 Hz, 1H), 2.46 (ddt, *J* = 14.7, 10.8, 5.5 Hz, 2H), 2.16 (dt, *J* = 15.1, 4.6 Hz, 1H), 1.86 – 1.80 (m, 1H), 1.72 (d, *J* = 9.8 Hz, 1H), 1.65 (td, *J* = 12.5, 5.9 Hz, 1H), 1.58 (dt, *J* = 12.2, 6.2 Hz, 1H), 1.49 (d, *J* = 6.8 Hz, 3H), 1.46 – 1.34 (m, 3H), 1.29 (q, *J* = 8.0 Hz, 2H), 1.18 (s, 1H), 1.06 (ddd, *J* = 14.1, 9.9, 5.0 Hz, 1H), 0.89 (qd, *J* = 9.9, 6.1, 5.3 Hz, 1H), 0.82 (t, *J* = 9.2 Hz, 1H), 0.75 (qd, *J* = 10.1, 6.0, 4.9 Hz, 1H), 0.56 – 0.48 (m, 1H). $^{13}$C NMR (151 MHz, Acetone) δ 182.13, 165.50, 129.56, 128.90, 127.67, 123.07, 119.52, 114.05, 110.25, 102.07, 94.84, 70.58, 59.07, 57.33, 47.64, 39.62, 32.22, 30.36 (d, *J* = 3.0 Hz), 30.22, 27.94, 26.19, 25.86 – 25.47 (m), 25.39, 22.91. LC-MS: calculated for $C_{26}H_{35}N_3O_3$ [M+H$^+$] 438.2757 m/z; found 438.2754 m/z.

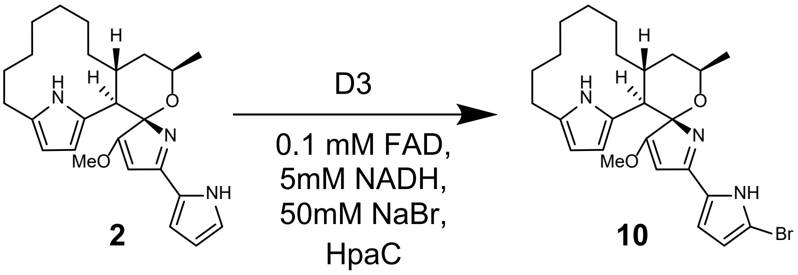

**1-bromo premarineosin A (10).** Preparatory reactions were performed using 40 µM D3, 50 µM HPAC flavin reductase, 100 µM FAD, 50 mM sodium bromide, and 500 µM of (-)-premarineosin A dissolved in DMSO at a final volume of 5mL in reaction buffer (10 mM HEPES, pH 7.4 containing 10% glycerol). The reaction was initiated upon the addition of 5M NADH. The reaction was quenched via the addition of 20 mL of HPLC-grade methanol, vortexed on the highest setting for 20 s, then water bath sonicated for 30 s. The resulting mixture from all combined 5mL reactions was then passed through a pad of Celite© and the filter cake was washed with an additional 20 mL of methanol. To purify the biocatalytic byproduct, the aqueous solution was extracted with DCM. The organics were combined and concentrated. The resulting material was further purified by preparative HPLC with a phenyl hexyl column (5 µm, 100 Å, 250 x 10 mm) using a gradient of 10-55% of acetonitrile and water, both modified with 0.1% formic acid, over 50 minutes at a flow rate of 5 mL/min. $^1$H NMR (600 MHz, Acetone) $^1$H NMR (600 MHz, Acetone): δ 9.49 (br s, 1H), 6.89 (br d, *J* = 2.0 Hz, 1H), 6.17 (br d, *J* = 2.5 Hz, 1H), 5.74 (br s, 1H), 5.68 – 5.65 (m, 1H), 5.52 – 5.48 (m, 1H), 4.33 (br p, *J* = 6.4 Hz, 1H), 3.91 (s, 3H), 2.93 (d, *J* = 12.3 Hz, 1H), 2.66 – 2.58 (m, 1H), 2.46 (dt, *J* = 15.1, 4.6 Hz, 1H), 2.31 (t, *J* = 14.0, 2.6 Hz, 1H), 1.91 (dt, *J* = 13.6, 4.0 Hz, 1H), 1.80 (br q, *J* = 11 Hz, 1H), 1.70 – 1.63 (m, 2H), 1.50 (d, *J* = 6.7 Hz, 1H), 1.48 – 1.43 (m, 1H), 1.43 – 1.30 (m, 3H), 1.21 – 1.13 (m, 1H), 1.09 – 1.00 (m, 2H), 0.84 – 0.75 (m, 2H), 0.61 – 0.50 (m, 1H).$^{13}$C NMR (151 MHz, Acetone) δ 177.65, 161.77, 133.17, 130.58, 127.36, 124.88, 117.10, 109.87, 105.89, 97.07, 93.38, 70.46, 59.42, 46.91, 38.94, 33.30, 29.01, 28.80, 27.38, 25.84, 25.66, 25.28, 22.06. LC-MS: calculated for $C_{25}H_{32}BrN_3O_2$ [M+H$^+$] 486.1756 m/z; found 486.1771 m/z.

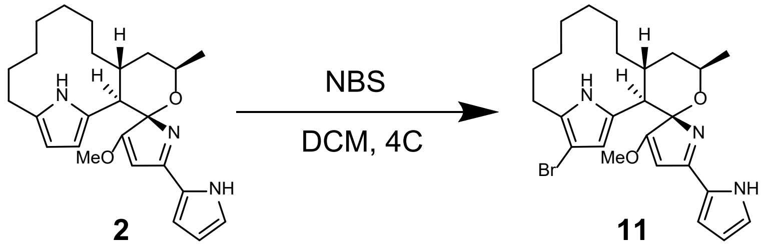

**12-bromo premarineosin A (11).** To a solution of (-)-premarineosin A (13.8mg, 5mmol) in dichloromethane at 4 °C was added *N*-Bromo succinimide (10 mg, 10mmol, DCM) drop wise. The mixture was stirred at 4 °C. After 2 h the reaction was quenched with sodium bicarbonate. Following a liquid-liquid extraction with DCM, the organic layer was extracted and dried. The resulting material was further purified by preparative HPLC with a phenyl hexyl column (5 µm, 100 Å, 250 x 10 mm) using a gradient of 10-55% of acetonitrile and water, both modified with 0.1% formic acid, over 50 min at a flow rate of 5 mL/min. $^{1}H$ NMR (600 MHz, $CDCl_3$, formate adduct): δ 14.68 (br s, 1H), 10.56 (br s, 1H), 13.41 (br s, 1H), 8.84 (s, 1H), 7.48 (br s, 1H), 7.07 (br d, *J* = 2.90 Hz, 1H), 5.77 (br s, 1H), 5.58 (br s, 1H), 4.41 (br p, *J* = 6.8 Hz, 1H), 4.05 (s, 3H), 2.93 (d, *J* = 12.6 Hz, 1H), 2.80 – 2.70 (m, 1H), 2.49 (ddd, *J* = 14.7, 10.9, 3.6 Hz, 1H), 2.32 (dt, *J* = 14.7, 4.6 Hz, 1H), 1.92 (dt, *J* = 13.8, 4.0 Hz, 1H), 1.72 – 1.64 (m, 1H), 1.52 (d, *J* = 6.9 Hz, 3H), 1.42 – 1.32 (m, 3H), 1.30 – 1.22 (m, 1H), 1.17 – 1.09 (m, 1H), 1.05 – 0.97 (m, 1H), 0.96 – 0.83 (m, 2H), 0.53 – 0.44 (m, 1H). $^{13}C$ NMR (151 MHz, $CDCl_3$) δ 179.34, 170.28, 164.40, 134.62, 132.06, 125.88, 124.67, 121.12, 113.53, 111.85, 96.27, 92.75, 71.37, 60.03, 45.48, 37.74, 32.37, 27.36, 27.18, 26.31, 25.17, 24.47, 24.33, 20.87. LC-MS: calculated for $C_{25}H_{32}BrN_3O_2$ [M+H$^+$] 486.1756 m/z; found 486.1811 m/z.

#### Supplementary Tables.

**Supplementary Table 1.** (-)-Premarineosin A biosynthetic genes in *S. eitanensis* and their homology to the *Streptomyces sp. CNQ-617 mar* BGC genes.

| ***S. eitanensis*** **Protein** | **Description** | ***S.*** **sp. CNQ-617 NCBI Accession No.** | **Similarity (%)** | **Identity (%)** |
|---|---|---|---|---|
| PmaD | transcriptional regulator, SARP family | AHF22840.1 | 77% | 65% |
| PmaX | beta-ketoacyl synthase | AHF22841.1 | 68% | 60% |
| PmaW | acyl-CoA dehydrogenase | AHF22842.1 | 83% | 74% |
| PmaY | unknown | AHF22843.1 | 68% | 57% |
| PmaZ | LuxR family DNA binding response regulator | AHF22844.1 | 74% | 55% |
| PmaV | unknown | AHF22845.1 | 64% | 56% |
| PmaU | 4'-phosphopantetheinyl transferase | AHF22846.1 | 69% | 58% |
| PmaT | unknown | AHF22847.1 | 76% | 63% |
| PmaR | Beta-ketoacyl synthase | AHF22848.1 | 86% | 79% |
| PmaQ | PP-binding | AHF22849.1 | 82% | 67% |
| PmaP | 3-oxoacyl-(acyl carrier protein) synthase III | AHF22850.1 | 84% | 76% |
| PmaO | Putative acyl carrier protein | AHF22851.1 | 85% | 69% |
| PmaN | PP-binding, aminotransferase I/II, Amino-7-oxononanoate synthase | AHF22852.1 | 77% | 68% |
| PmaM | AMP-binding, AMP-dependent synthetase and ligase | AHF22853.1 | 76% | 67% |
| PmaL | NRPS-PKS hybrid | AHF22854.1 | 70% | 61% |
| PmaK | RmlD_sub_binding, NAD-dependent epimerase/dehydratase | AHF22855.1 | 81% | 73% |
| PmaJ | thioesterase | AHF22856.1 | 64% | 56% |
| PmaI | methyltransferase | AHF22857.1 | 81% | 70% |
| PmaH | PPDK_N | AHF22858.1 | 83% | 75% |
| PmaG | NONE | AHF22859.1 | 85% | 77% |

**Supplementary Table 2.** Strains used in this work.

| Strain | Genotype | Reference |
|---|---|---|
| ***Streptomyces* Strains** | | |
| WT | *Streptomyces eitanensis* wild-type strain | Sherman Lab Collection |
| FP10016 | *Streptomyces eitanensis*::*kas*Op*, $Kan^R$ | This Work |
| FP10017 | *Streptomyces eitanensis*::*kas*Op*-*pmaD*, $Kan^R$ | This Work |
| FP10018 | *Streptomyces eitanensis*::*kas*Op*-*pmaD-pmaG,* $Kan^R$ | This Work |
| | *Streptomyces sp. CNQ-617* | Prof. William Fenical[5] |
| ***E. coli* Strains** | | |
| DH5α | *fhuA2Δ(argF-lacZ)U169 phoA glnV44 Φ80Δ(lacZ)M15 gyrA96 recA1 relA1 endA1 thi-1 hsdR17* | New England Biolabs |
| S17-1 | RP4 derivative integrated in chromosome | Laboratory Stock |

**Supplementary Table 3.** Plasmids used in this work.

| Plasmid | Relevant Characteristics | Reference |
|---|---|---|
| pSET152 | Integrative *E. coli-Streptomyces* shuttle vector, attP (ΦC31 integrase), $Apr^R$ | Laboratory Stock[6] (Bierman et al., 1992) |
| pSET152k-*kasOp** | Derived from pSET152k[7], expression driven by *kasO** promoter, $Kan^R$ | Laboratory Stock[8] (Pereira et al., 2025) |
| pSET152k-*pmaD* | Derived from pSET152k-*kas*Op*, *expressing pmaD*, $Kan^R$ | This Work |
| pSET152k-*pmaDG* | Derived from pSET152k-*kas*Op*, expressing *pmaD and pmaG*, $Kan^R$ | This Work |

**Supplementary Table 4.** Primers used in this work.

| Plasmid | Name | Sequence (synRBS) |
|---|---|---|
| pSET152k-*pmaD* | pmaD_fwd | TCTAAGTAAGGAGTGTCCATATGGCTATCCATCTTCTC |
| | pmaD_rev | GACGACAAAACTTTAGAGATTCACACGAGGCCGGCGTGCG |
| pSET152k-pmaDG | pmaD_fwd | see above |
| | pmaD-pmaG_rev | ATGGACACTCCTTACTCGAATCACACGAGGCCGGCGTGCG |
| | pmaD-pmaG_fwd | TTCGAGTAAGGAGTGTCCATATGATCCCCGACCTGTGG |
| | pmaG_rev | AGACGACAAAACTTTAGATCACCCTGCCGCCGCAC |

#### Supplementary Figures.

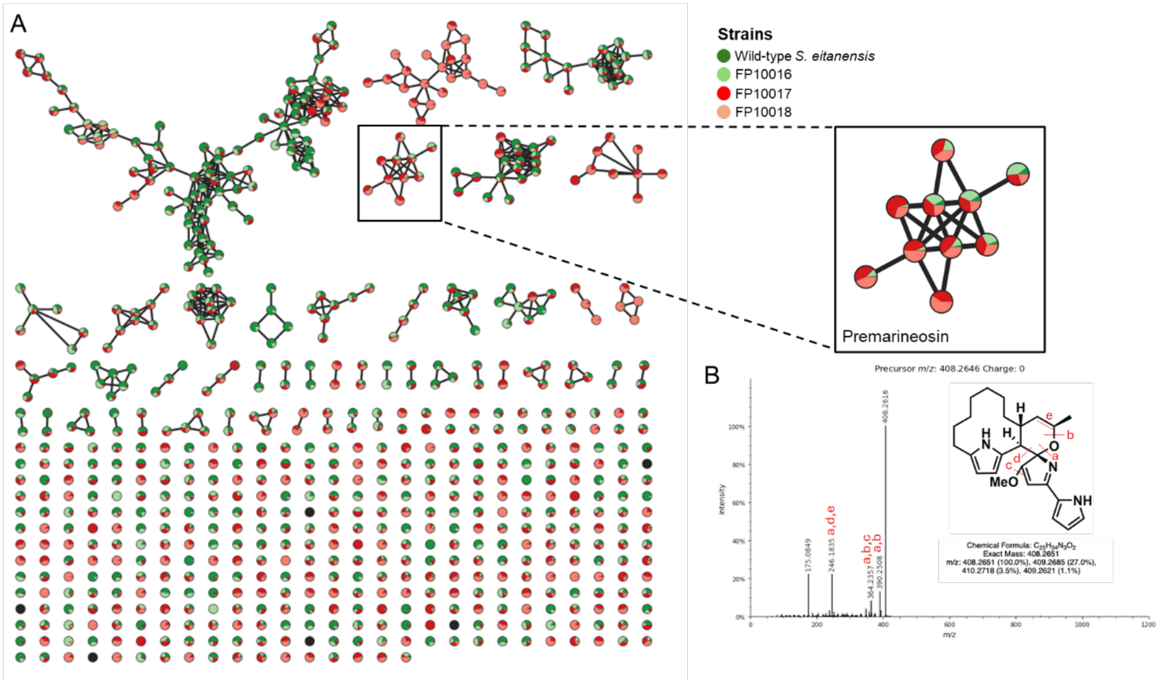

**Supplementary Figure 1. Untargeted metabolic analysis of *S. eitanensis* revealed production of a compound with the same molecular weight and fragmentation pattern as premarineosin A.** (**A**) Feature based analysis of untargeted metabolomics data from wild-type, vector control strain FP10016, and engineered strains FP10017 and FP10018 at day seven of strain cultivation in GICYE. (**B**) MS/MS fragmentation spectra for premarineosin A (408.26 m/z) with noted major fragments.

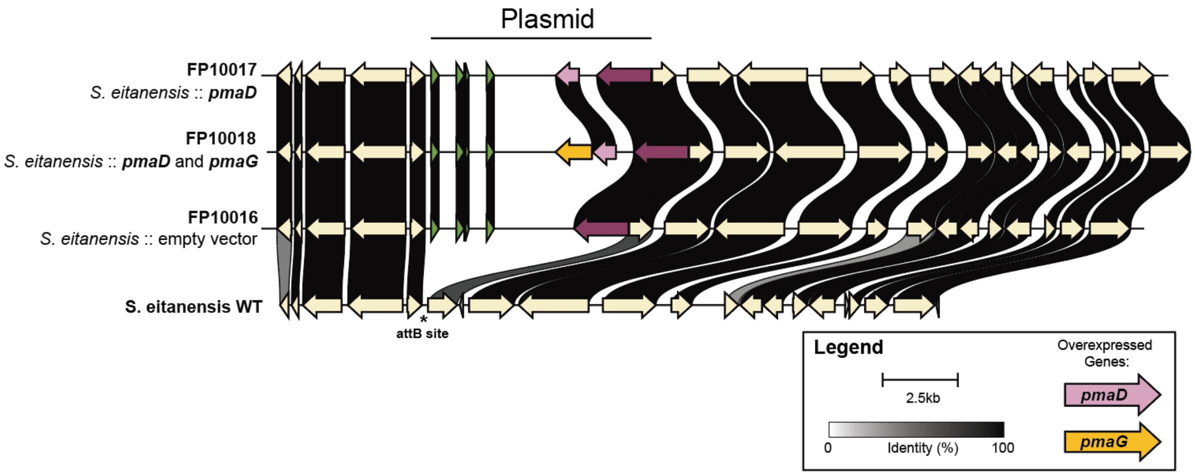

**Supplementary Figure 2. Integration of plasmids containing *pmaD* and *pmaG* in the *Streptomyces eitanensis* genome.** Comparative analysis of wild-type, FP10016, FP10017, and FP10018 genomes at the *attB* integration site using clinker.[9]

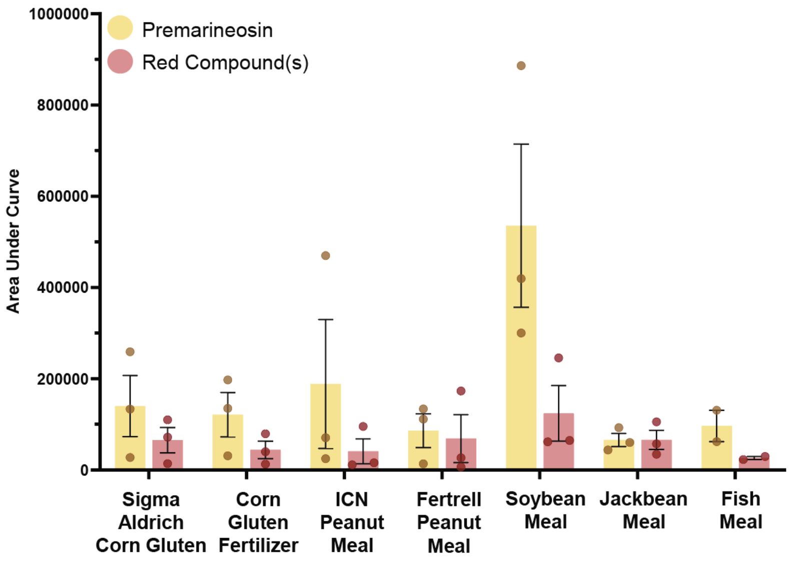

**Supplementary Figure 3. Effect of crude complex protein source on yield of (-)-premarineosin A in 1 mL extracts from 50 mL cultures of FP10018.** Dots represent three independent culture replicates. Error bars represent standard deviation (n=3).

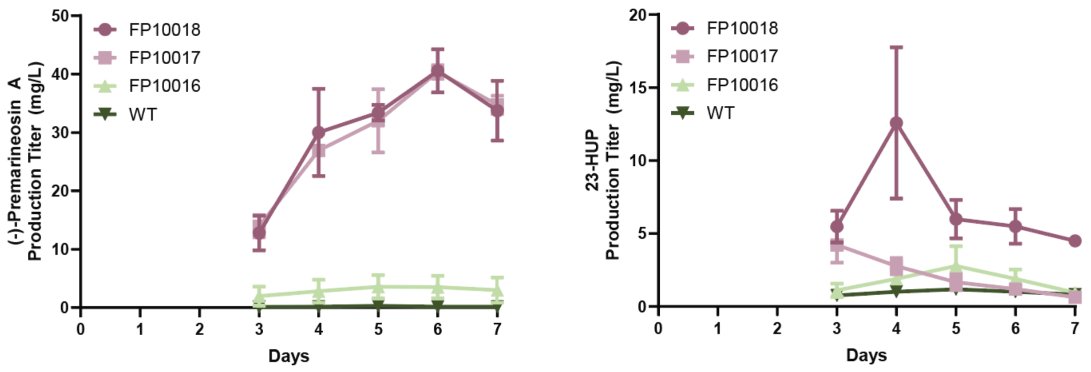

**Supplementary Figure 4. (-)-Premarineosin A and 23-HUP production (mg/L) increase over time.** Titers quantified by AUC (HPLC) against standard curves of each compound. Dots represent three independent culture replicates. Error bars indicate standard deviation (n=3). Production titers were below the limit of detection on days 1 and 2.

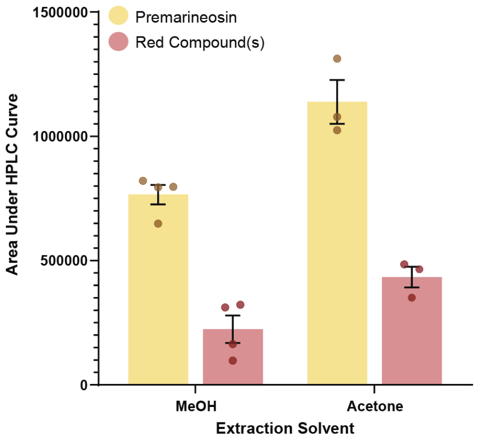

**Supplementary Figure 5. Extracting with acetone improves (-)-premarineosin A titers.** Production quantified by AUC (HPLC) from 1 mL extracts of FP10018 cultures grown in soybean meal culture media. Error bars represent standard error (n=3).

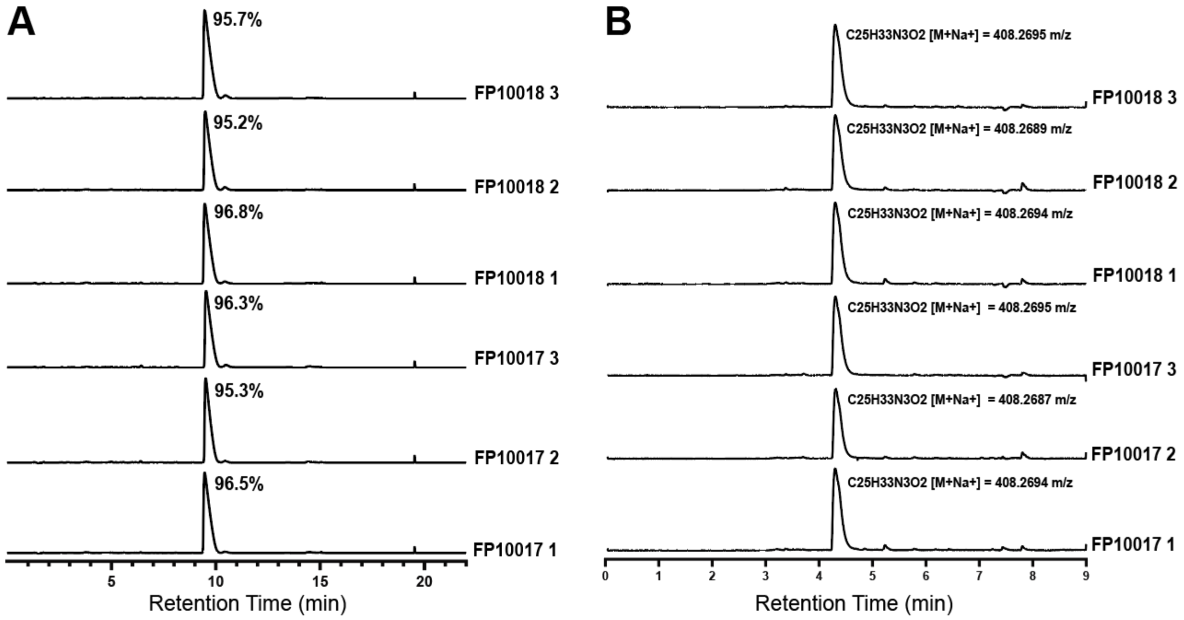

**Supplementary Figure 6. Purity analysis of replicate isolated yields of (-)-premarineosin A. (A)** HPLC traces for each replicate; premarineosin elutes at 9.5 min. The percentage represents degree of purity based on AUC at 346nm. **(B)** ESI EIC MS traces for each replicate; premarineosin elutes at 4.4 min.

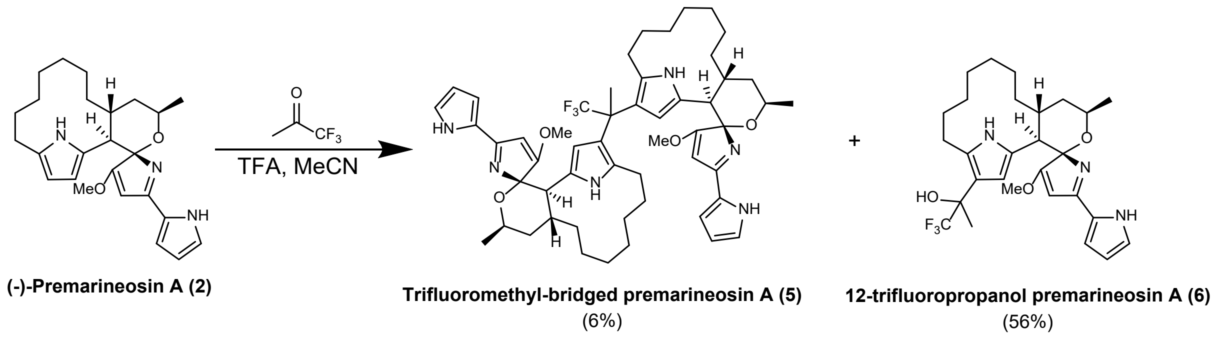

**Supplementary Figure 7. Formation of the 12-trifluoropropanol premarineosin A monomer.**

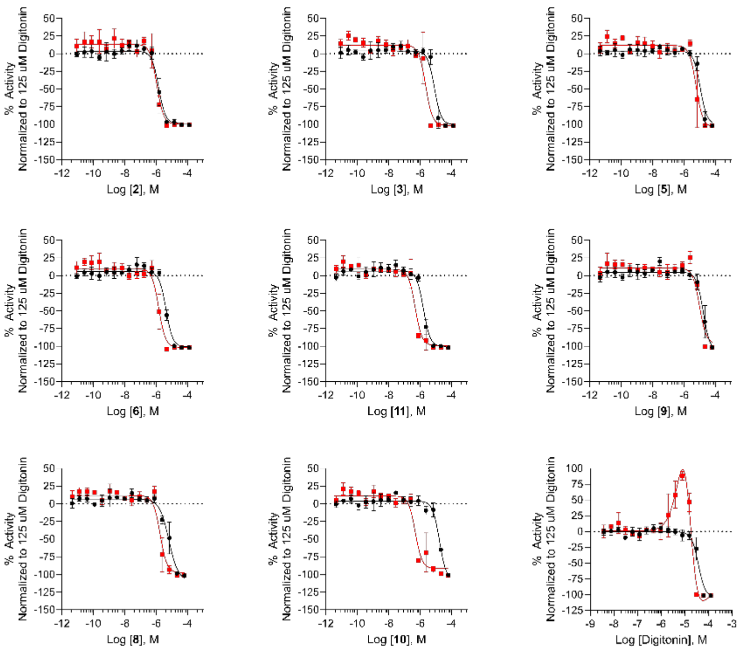

**Supplementary Figure 8. Mammalian cell cytotoxicity (72 h treatment) in HEK293 embryonic kidney (black symbols) and MOLT4 T-cell leukemia (red symbols) cell lines for two replicates per cell line (open and closed symbols).** Data was normalized to 125 µM Digitonin as -100% inhibition. Error bars represent standard deviation (n=2).

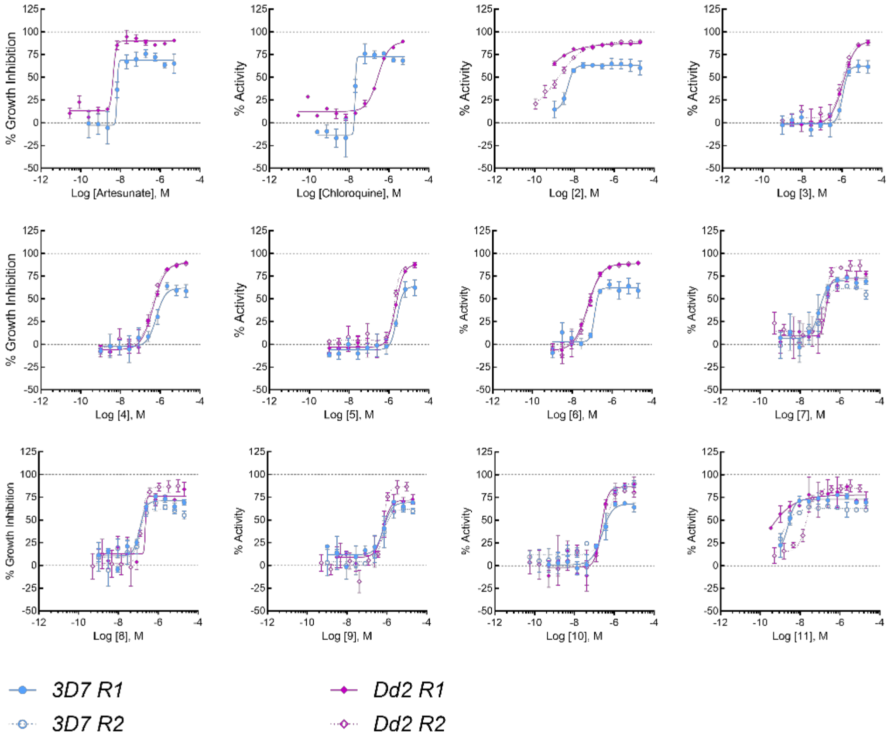

**Supplementary Figure 9. Activity of artesunate, chloroquine, (-)-premarineosin A, and derivatives 3-11 against *P. falciparum* 3D7 (chloroquine sensitive) and Dd2 (chloroquine resistant) parasite lines in 48 h assays.** Each *in vitro* experiment was performed in triplicate wells. Chloroquine and artesunate were tested in a single replicate experiment for each strain. Compounds **2-6** were tested in two independent experiments in Dd2 and one experiment in 3D7, and compounds **7-11** were tested in two independent experiments against both Dd2 and 3D7. Error bars represent standard deviation (n=3).

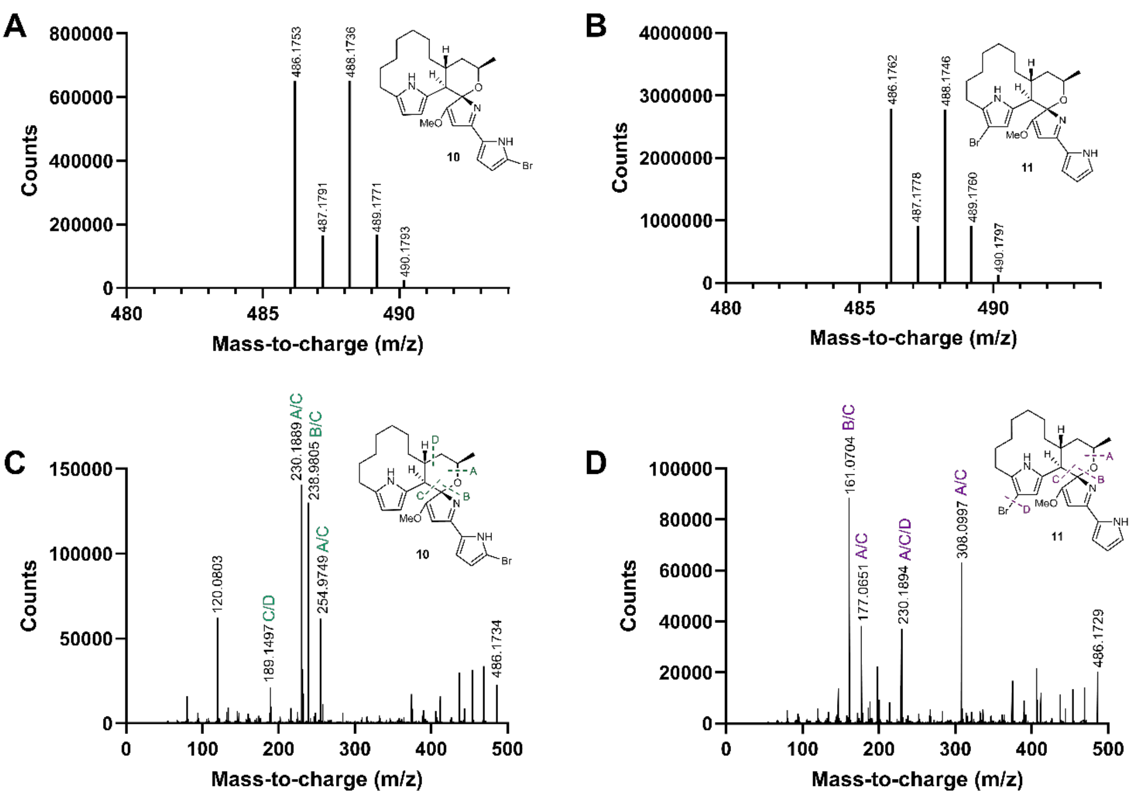

**Supplementary Figure 10. LC-MS/MS characterization of 1-bromo and 12-bromo premarineosin A.** Isotope pattern of **10** (A) and **11** (B), and MS2 fragmentation patterns for **10** (C) and **11** (D).

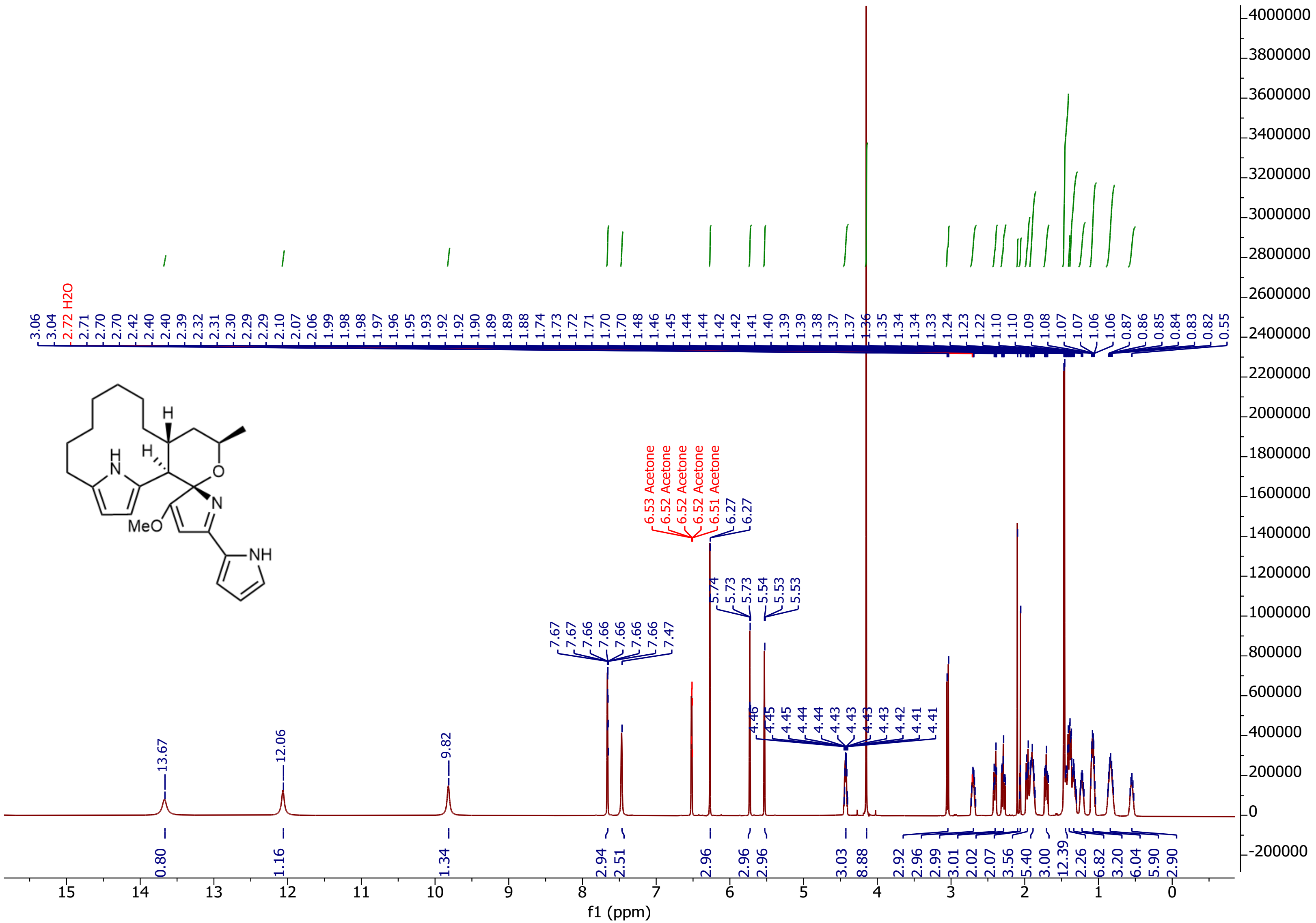

**Supplementary Figure 11.** $^{1}H$ NMR Spectrum of (-)-premarineosin A (**2**) in Acetone-$D_6$.

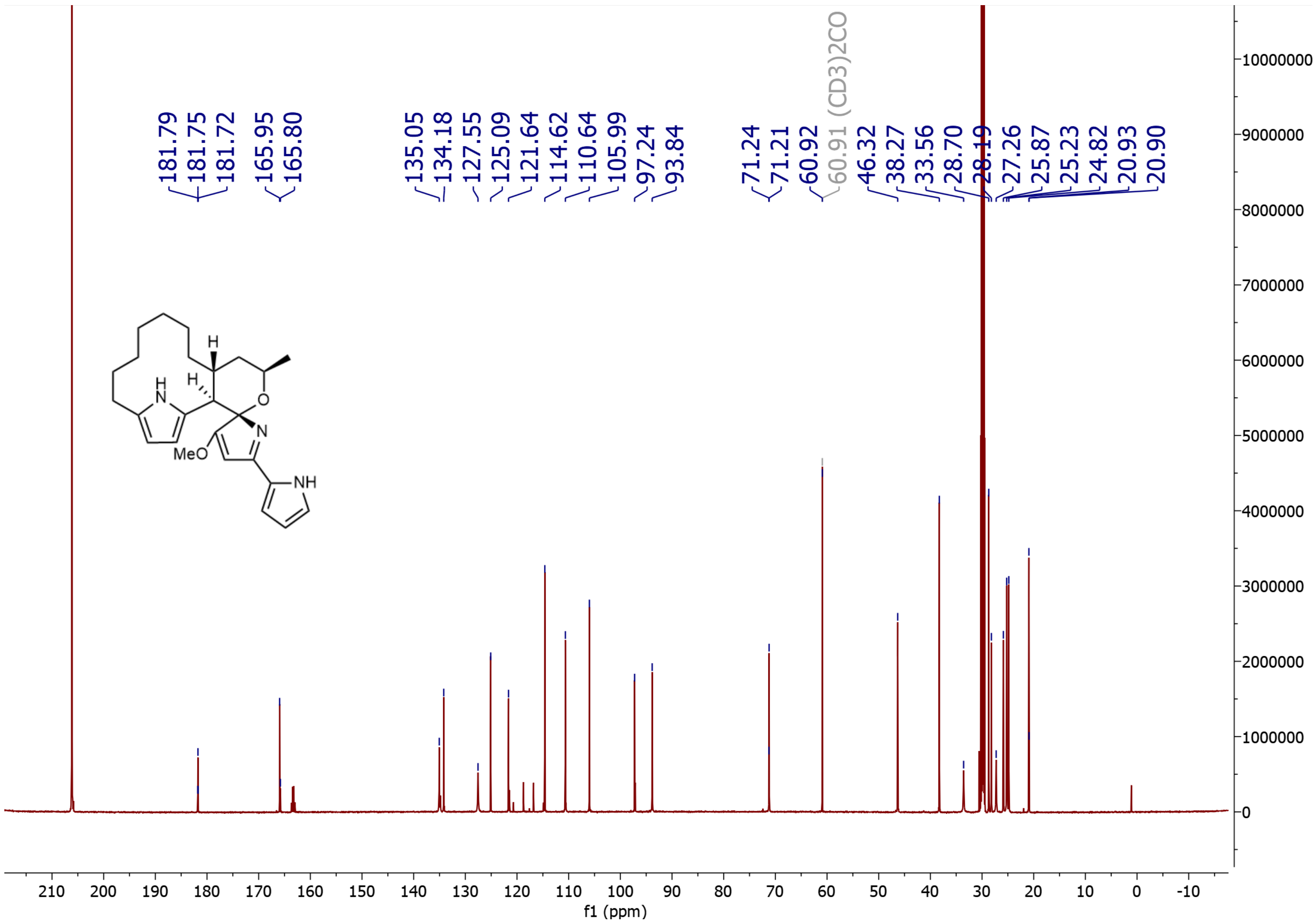

**Supplementary Figure 12.** $^{13}C$ NMR Spectrum of (-)-premarineosin A (**2**) in Acetone-$D_6$.

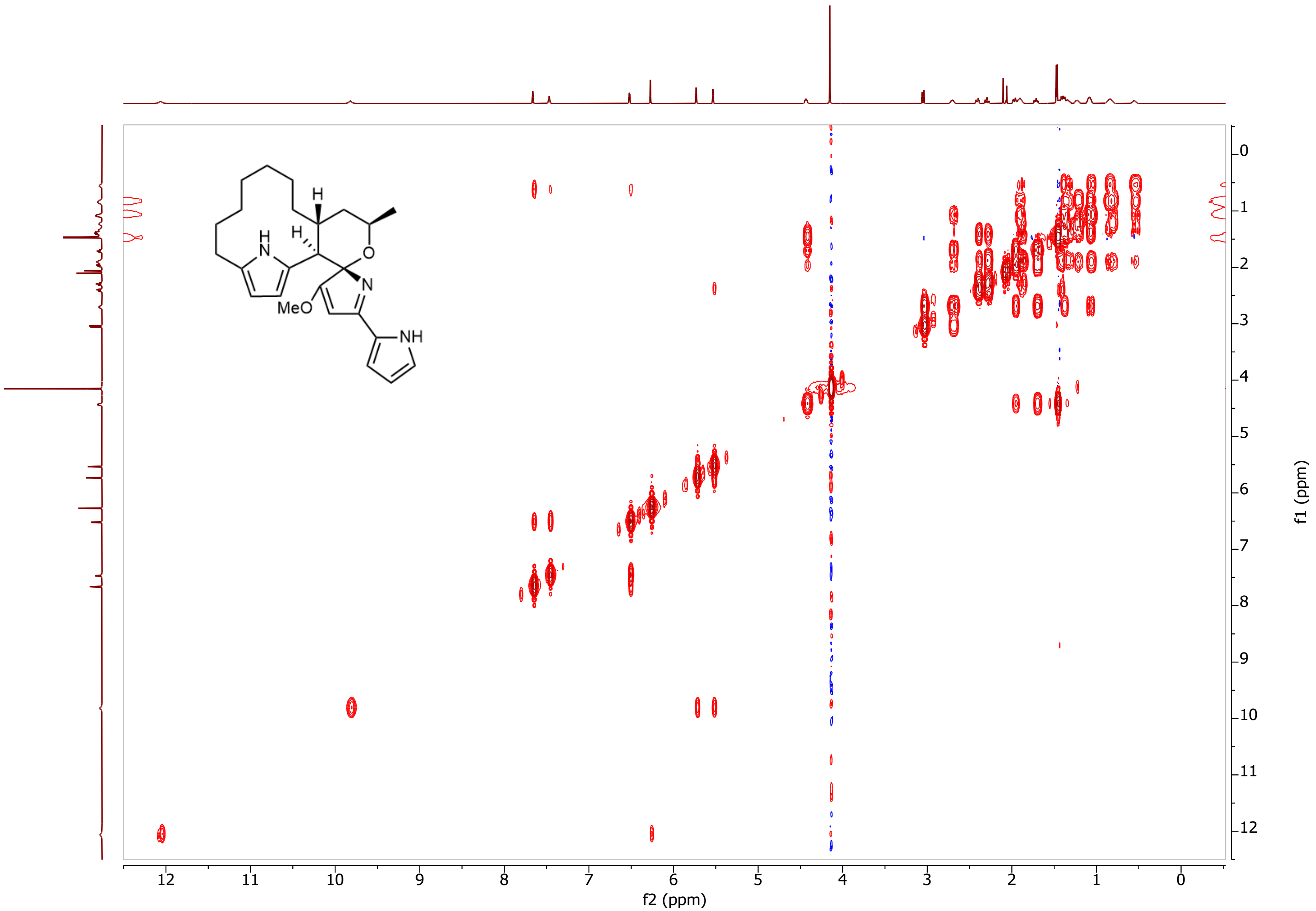

**Supplementary Figure 13.** $^{1}H$-$^{1}H$ COSY NMR Spectrum of (-)-premarineosin A (**2**) in Acetone-$D_6$.

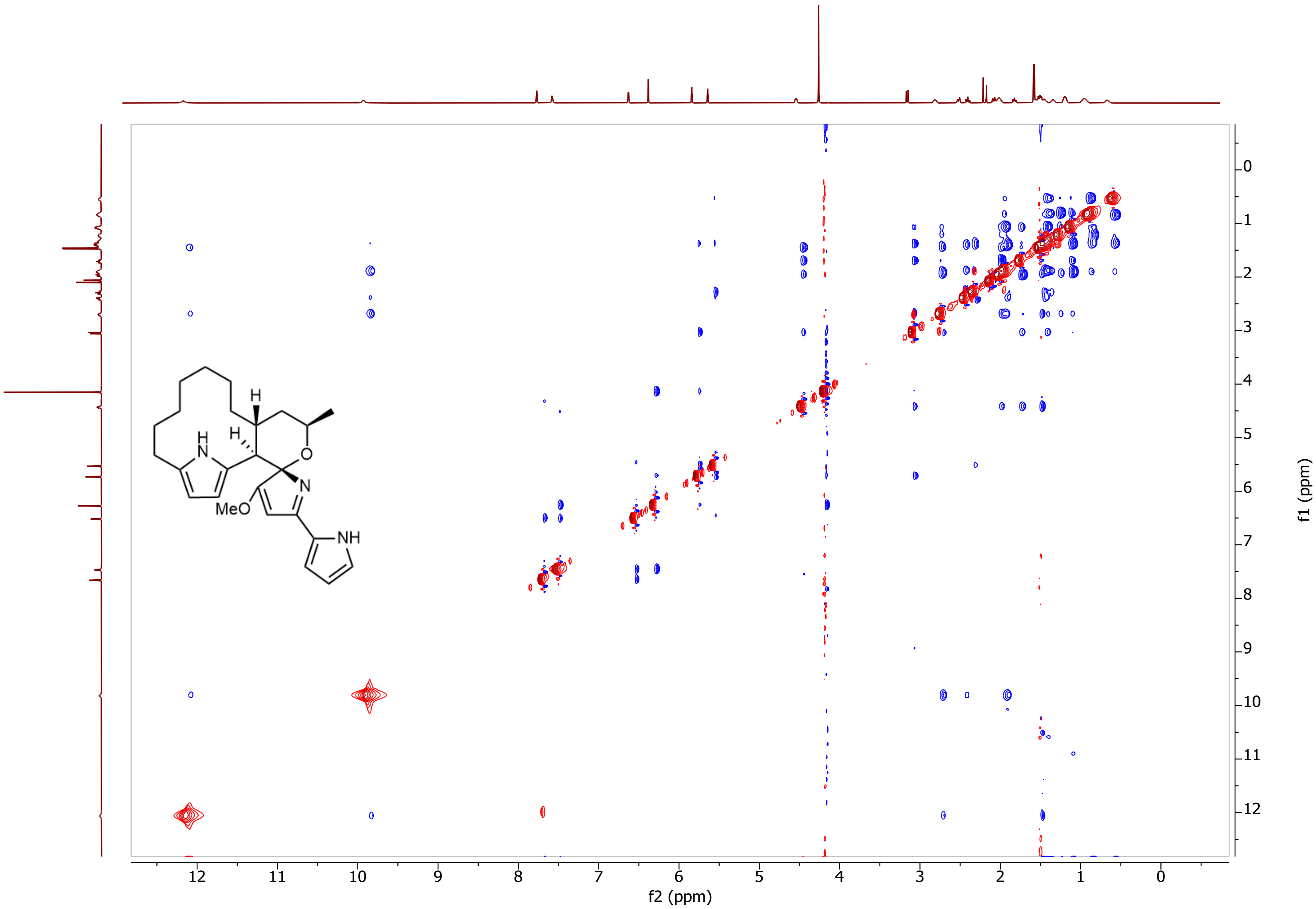

**Supplementary Figure 14.** $^{1}H$-$^{1}H$ NOESY NMR Spectrum of (-)-premarineosin A (**2**) in Acetone-$D_6$.

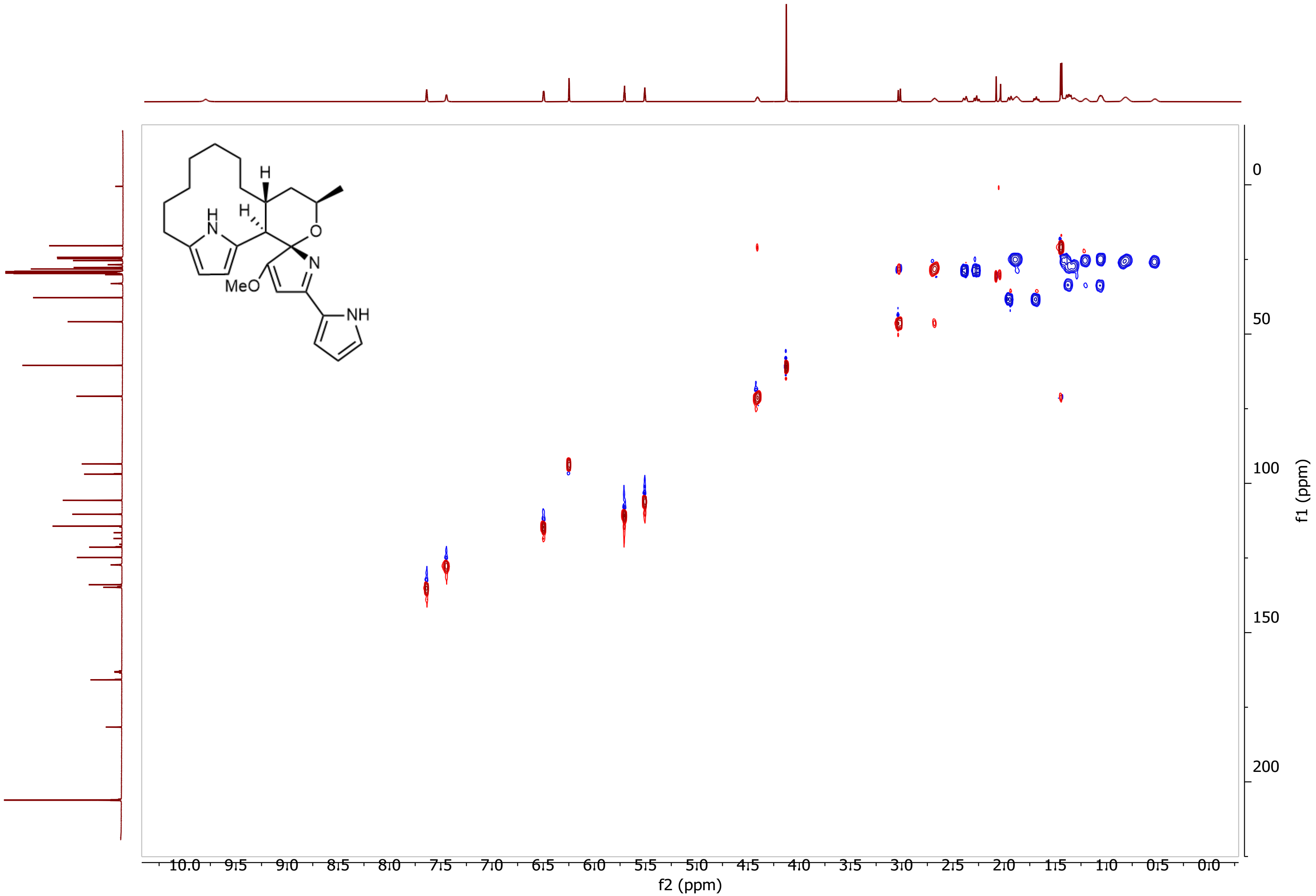

**Supplementary Figure 15.** $^1$H-$^{13}$C HSQC NMR Spectrum of (-)-premarineosin A (**2**) in Acetone-$D_6$.

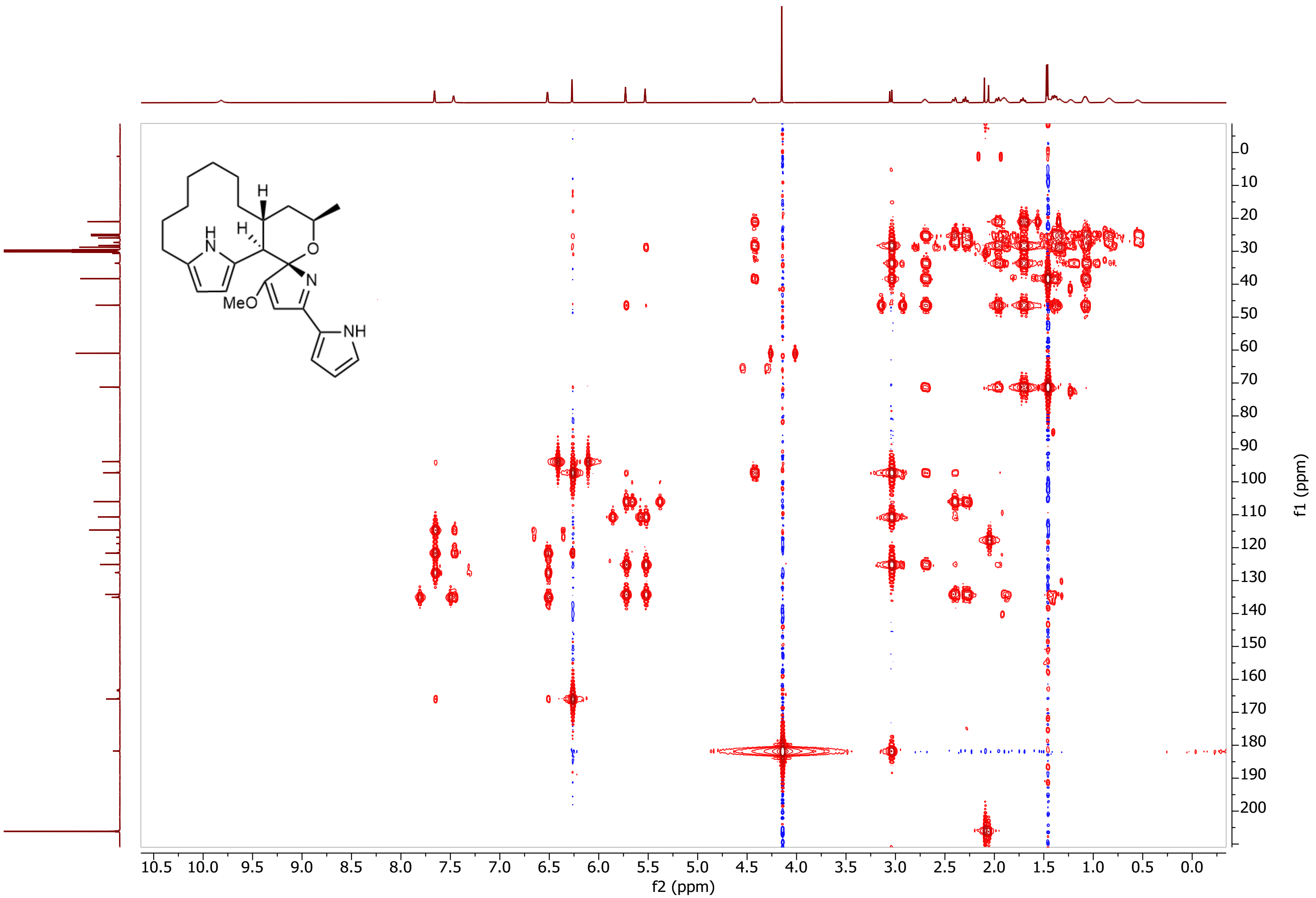

**Supplementary Figure 16.** $^{1}H$-$^{13}C$ HMBC NMR Spectrum of (-)-premarineosin A (**2**) in Acetone-$D_6$.

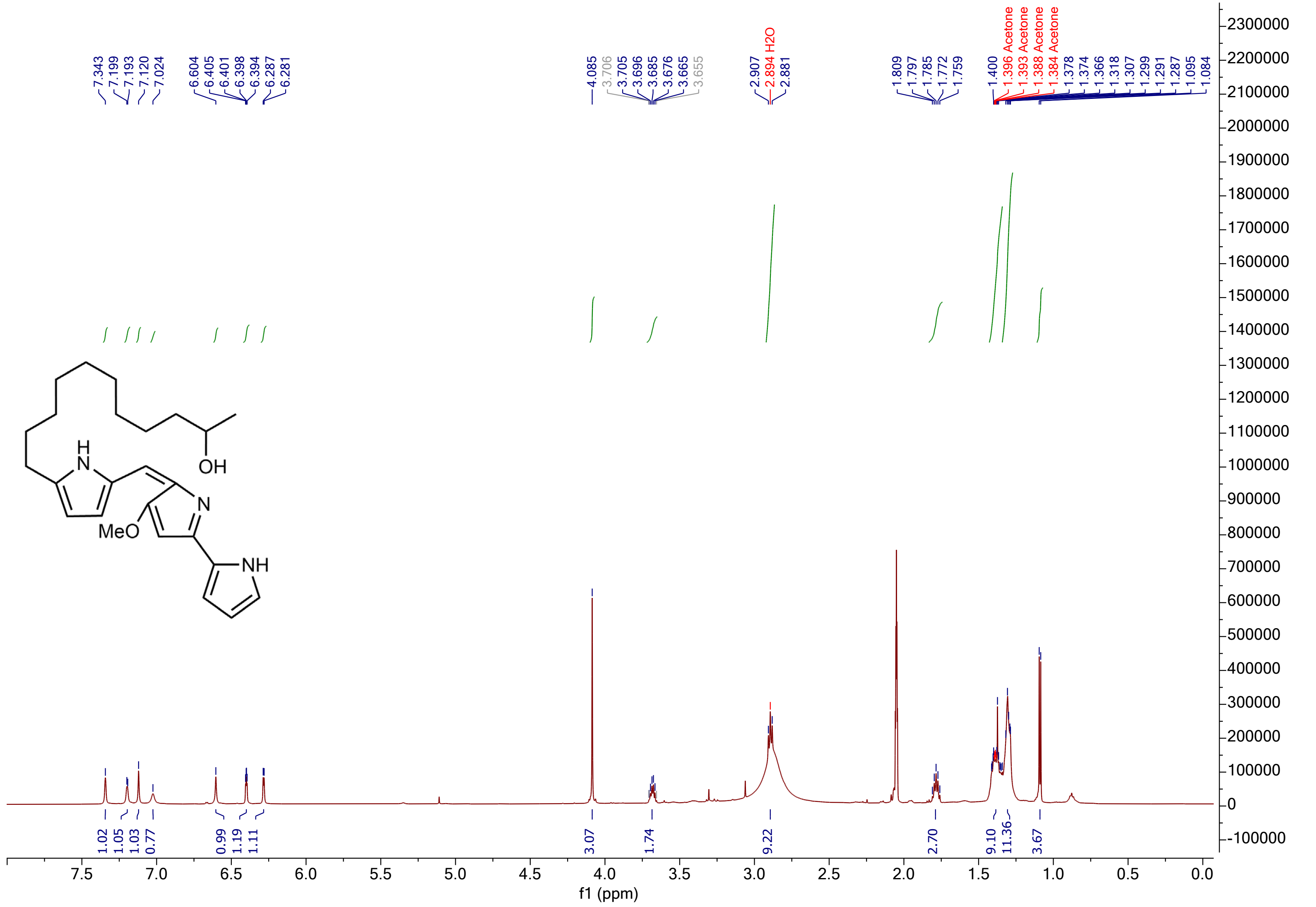

**Supplementary Figure 17.** $^{1}$H NMR Spectrum of 23-hydroxyundecylprodiginine (**1**) in Acetone-$D_6$.

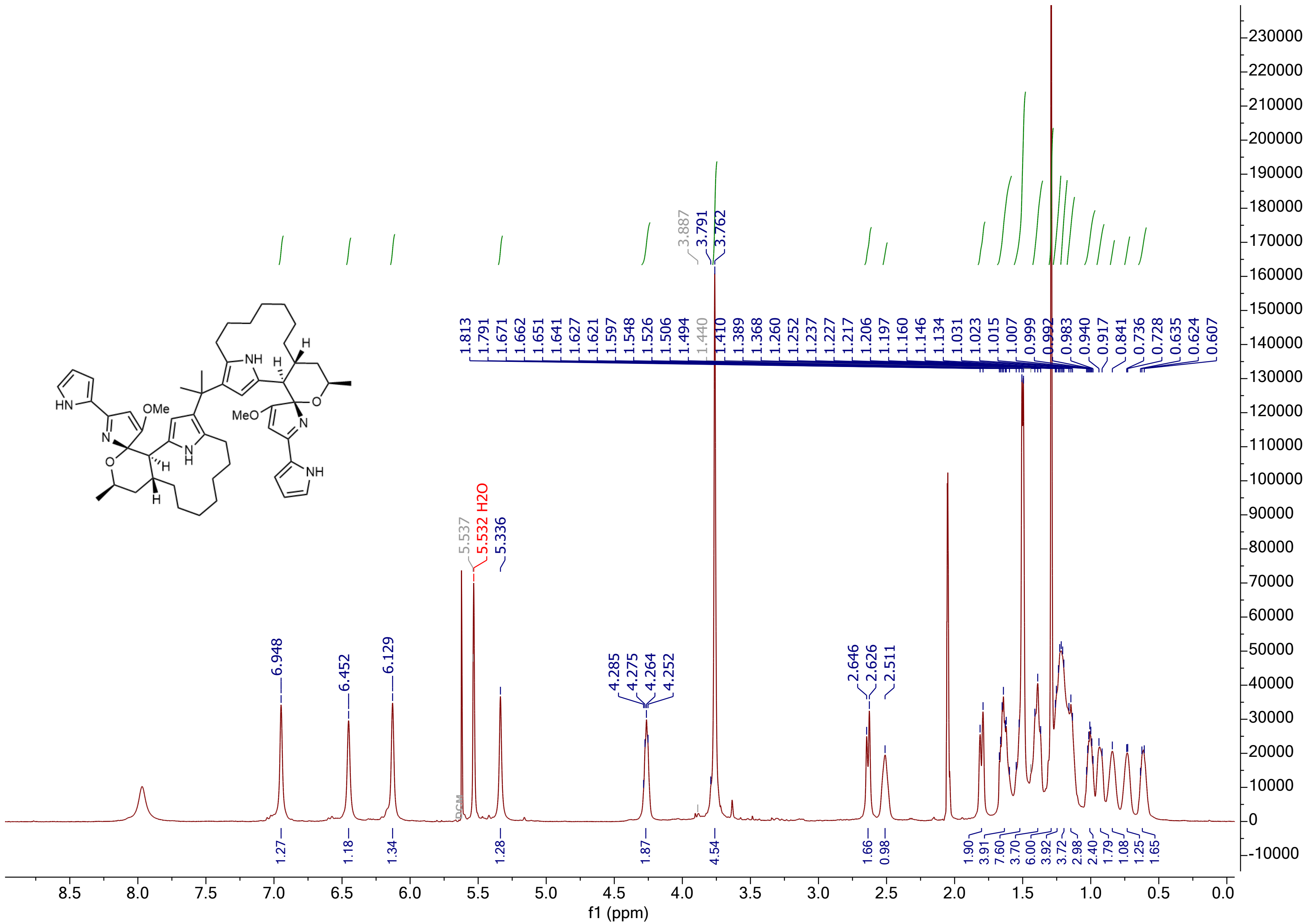

**Supplementary Figure 18.** $^{1}$H NMR Spectrum of gem-dimethyl-bridged premarineosin A (**3**) in Acetone-D$_6$.

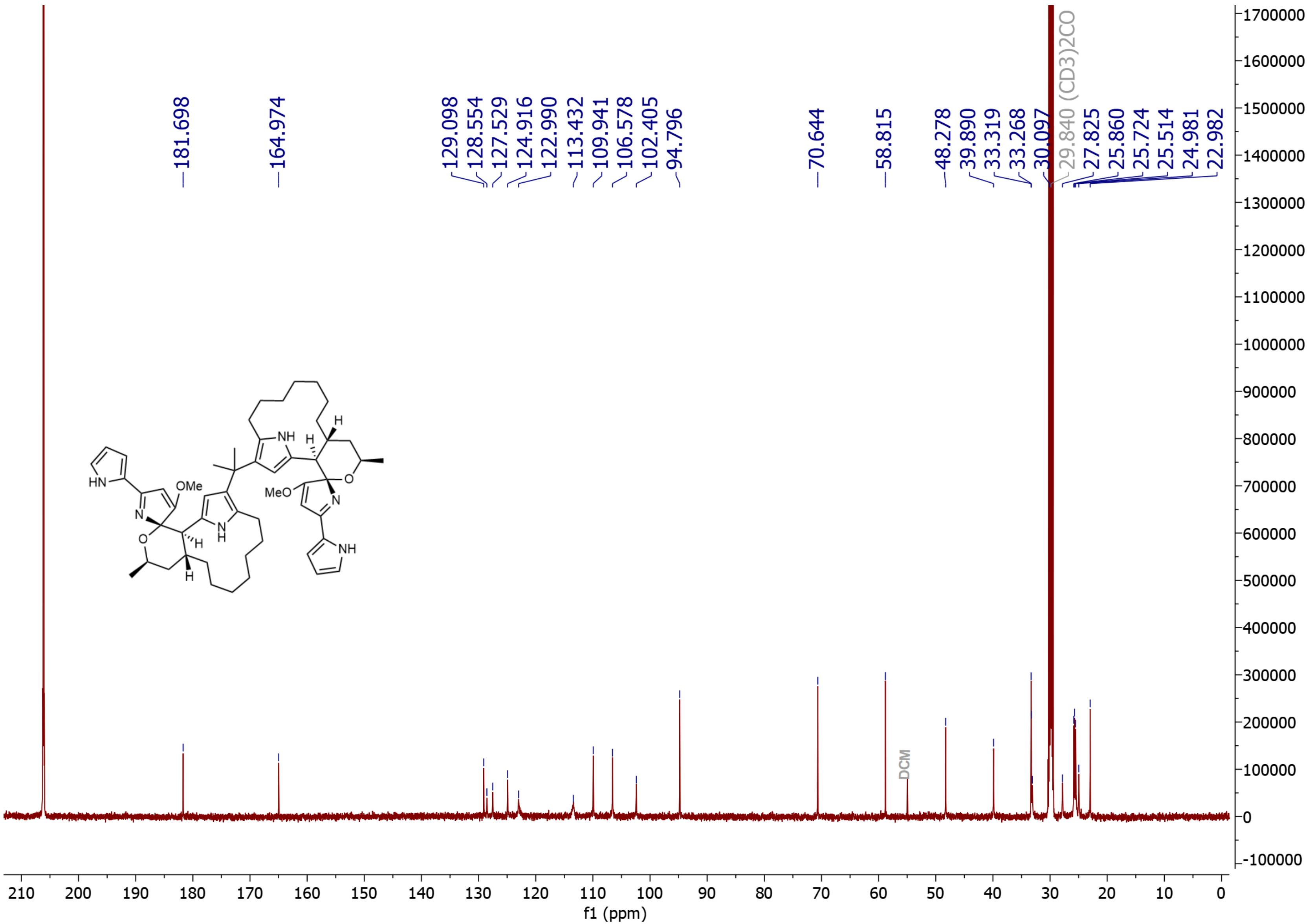

**Supplementary Figure 19.** $^{13}C$ NMR Spectrum of gem-dimethyl-bridged premarineosin A (**3**) in Acetone-$D_6$.

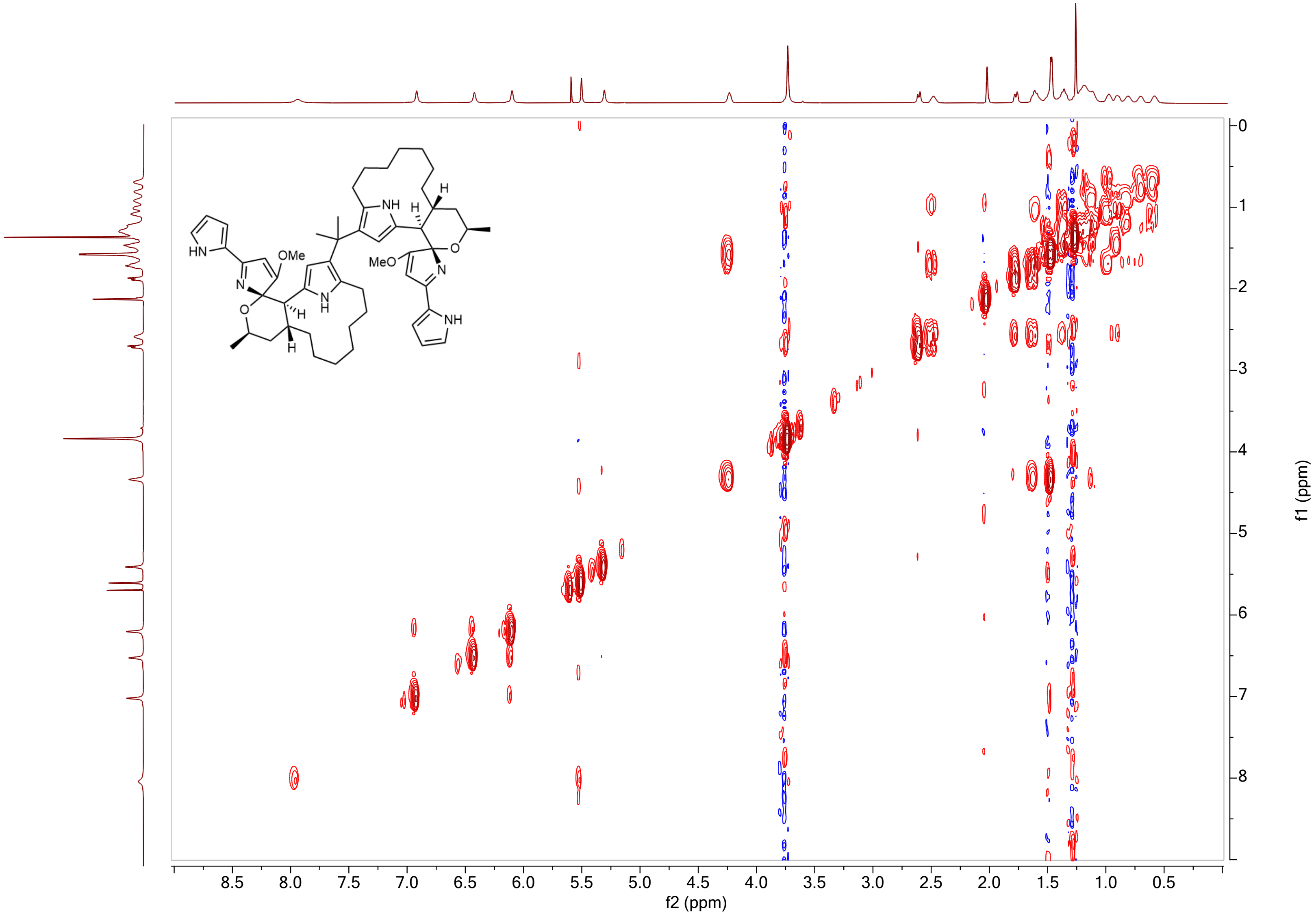

**Supplementary Figure 20.** $^1H$-$^1H$ COSY NMR Spectrum of gem-dimethyl-bridged premarineosin A (**3**) in Acetone-$D_6$.

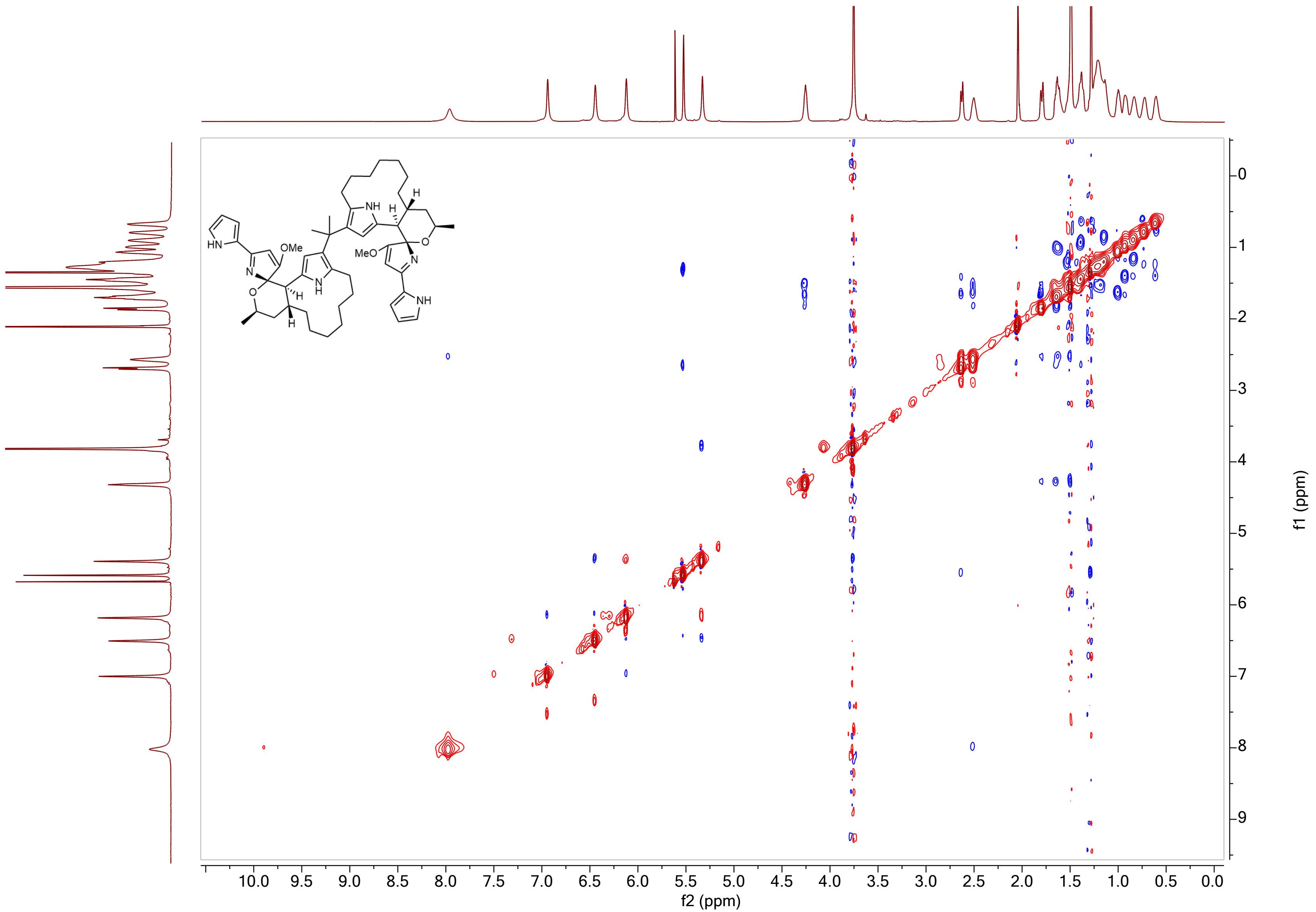

**Supplementary Figure 21.** $^{1}H$-$^{1}H$ NOESY NMR Spectrum of gem-dimethyl-bridged premarineosin A (**3**) in Acetone-$D_6$.

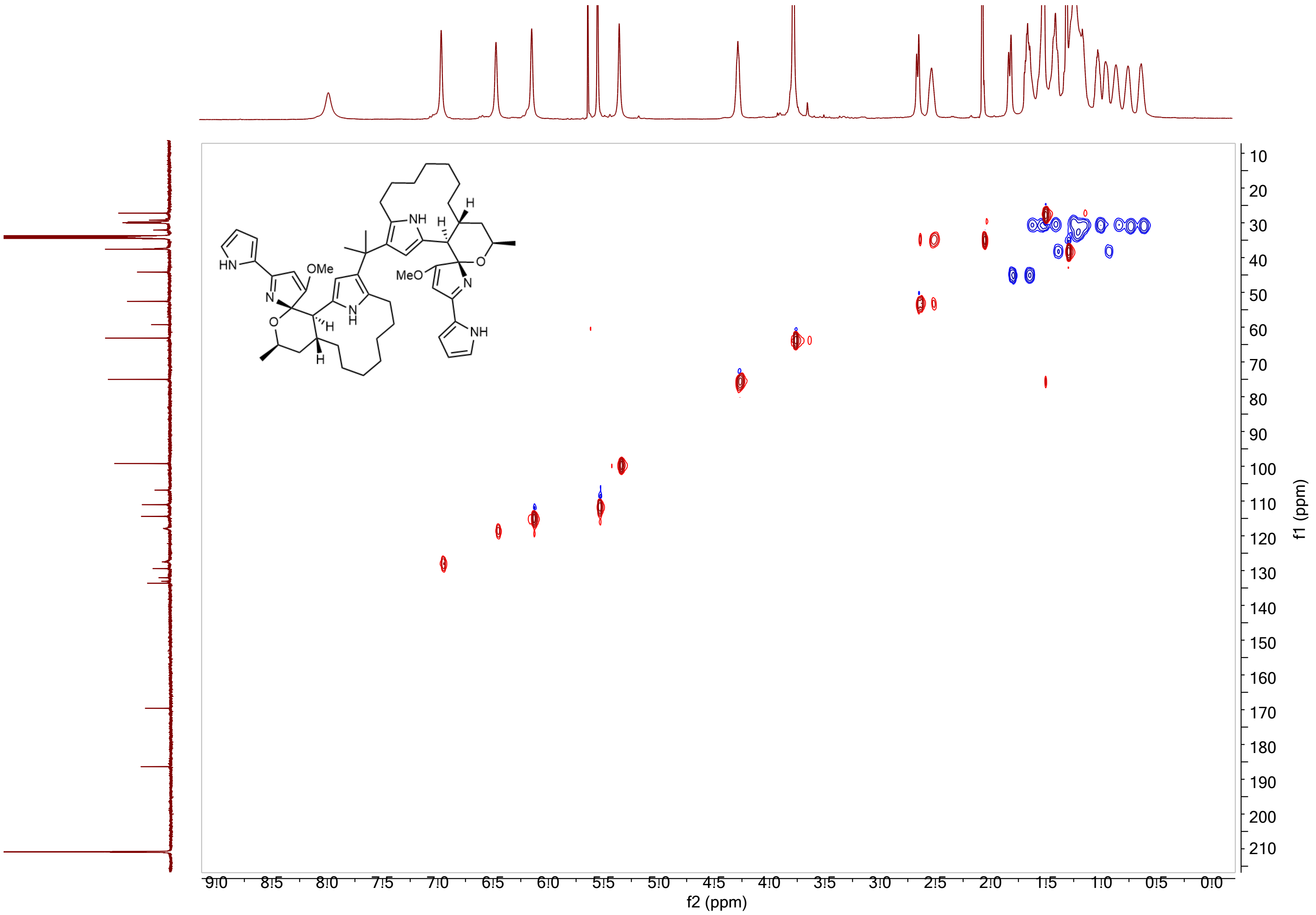

**Supplementary Figure 22.** $^{1}H$-$^{13}C$ HSQC NMR Spectrum of gem-dimethyl-bridged premarineosin A (**3**) in Acetone-$D_6$.

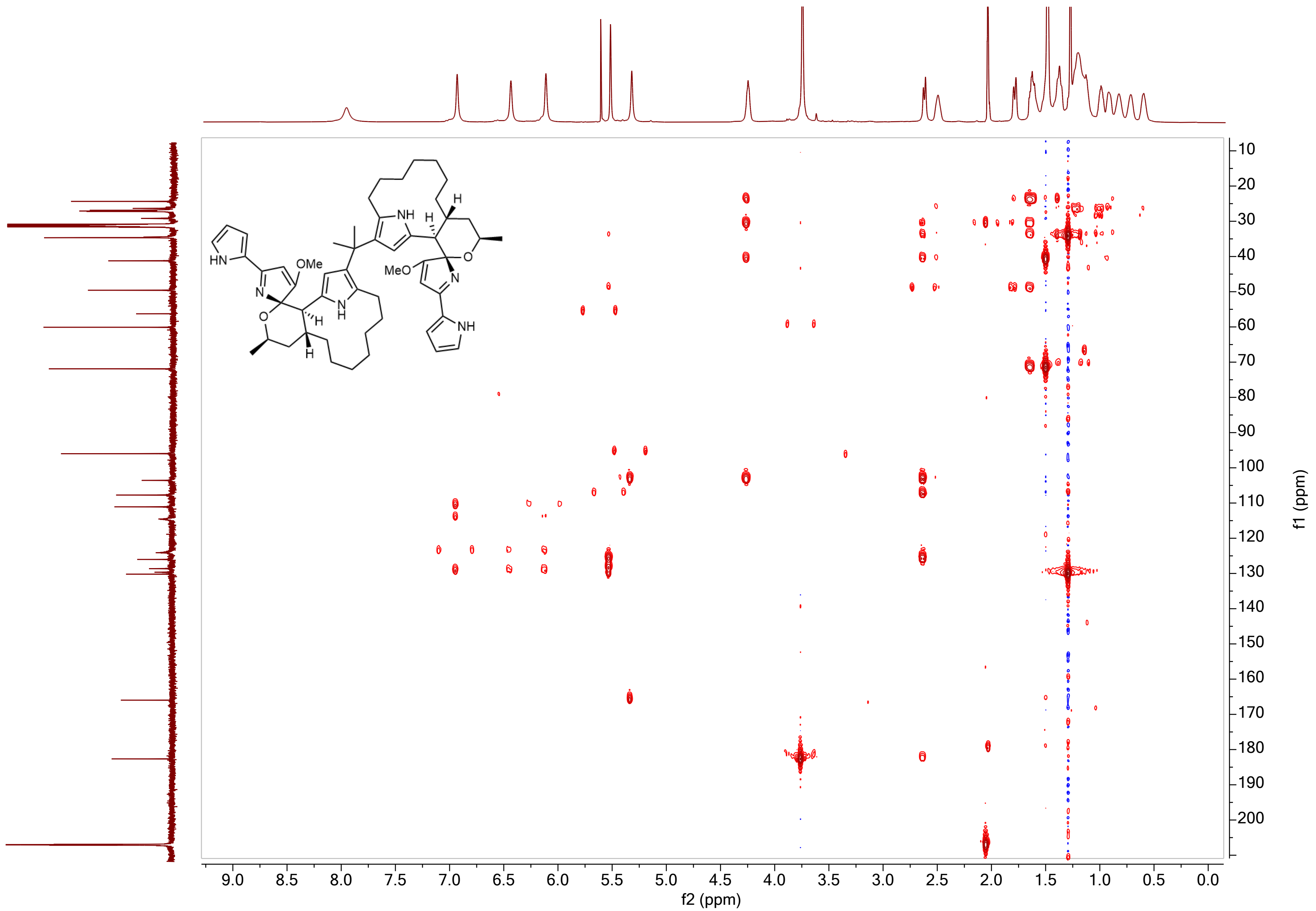

**Supplementary Figure 23.** $^{1}H$-$^{13}C$ HMBC NMR Spectrum of gem-dimethyl-bridged premarineosin A (**3**) in Acetone-$D_6$.

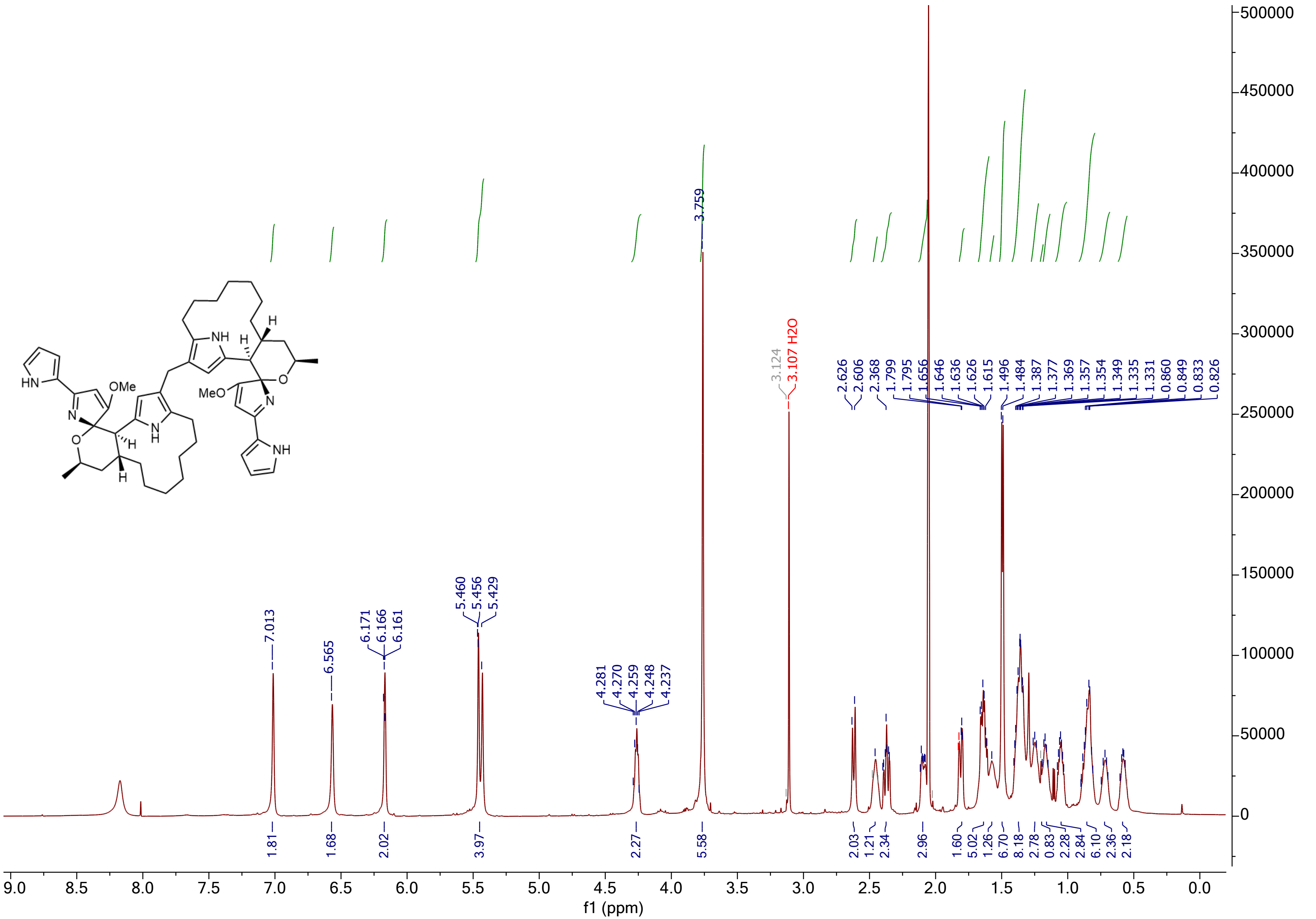

**Supplementary Figure 24.** $^{1}$H NMR Spectrum of methylene-bridged premarineosin A (**4**) in Acetone-$D_6$.

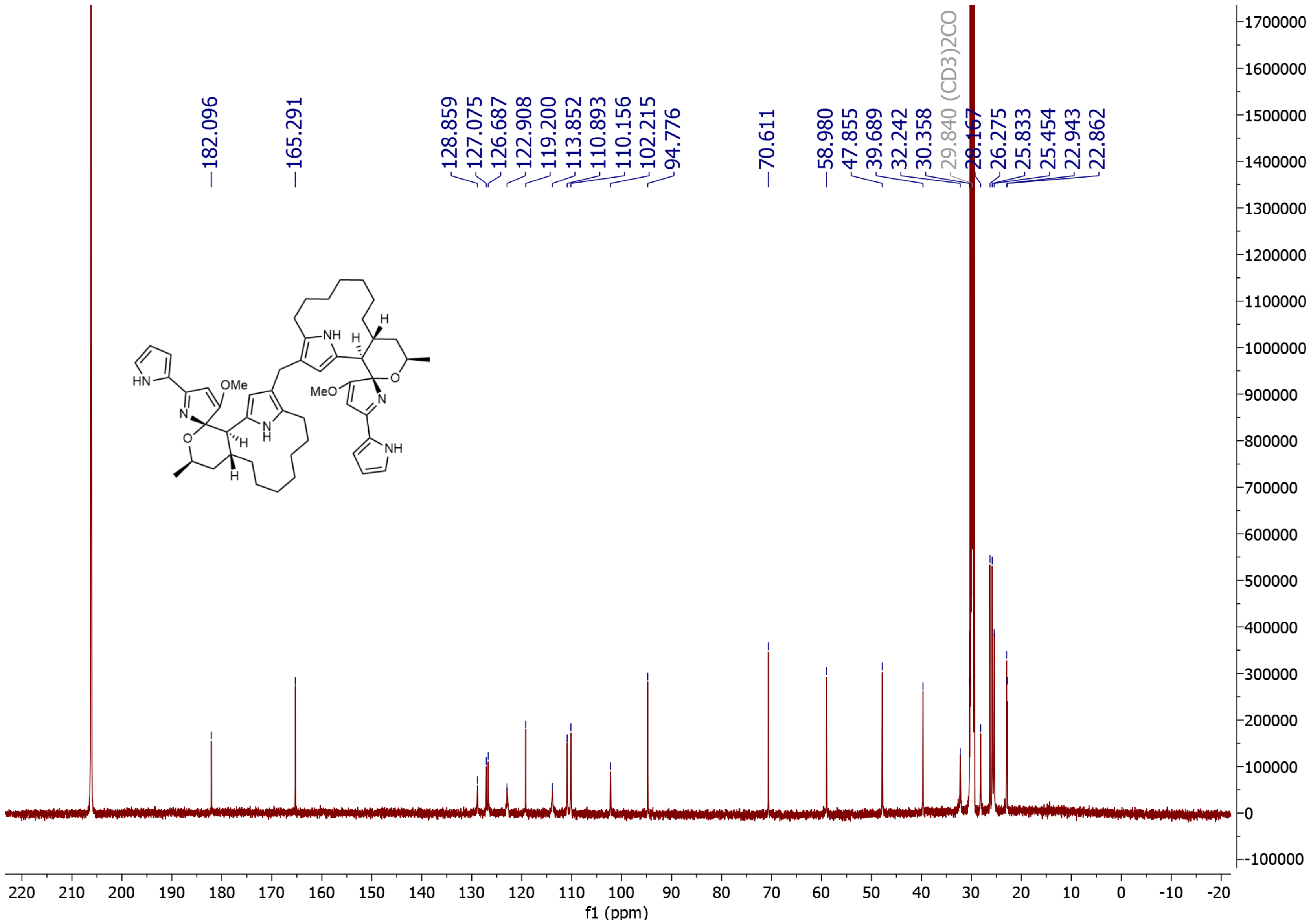

**Supplementary Figure 25.** $^{13}C$ NMR Spectrum of methylene-bridged premarineosin A (**4**) in Acetone-$D_6$.

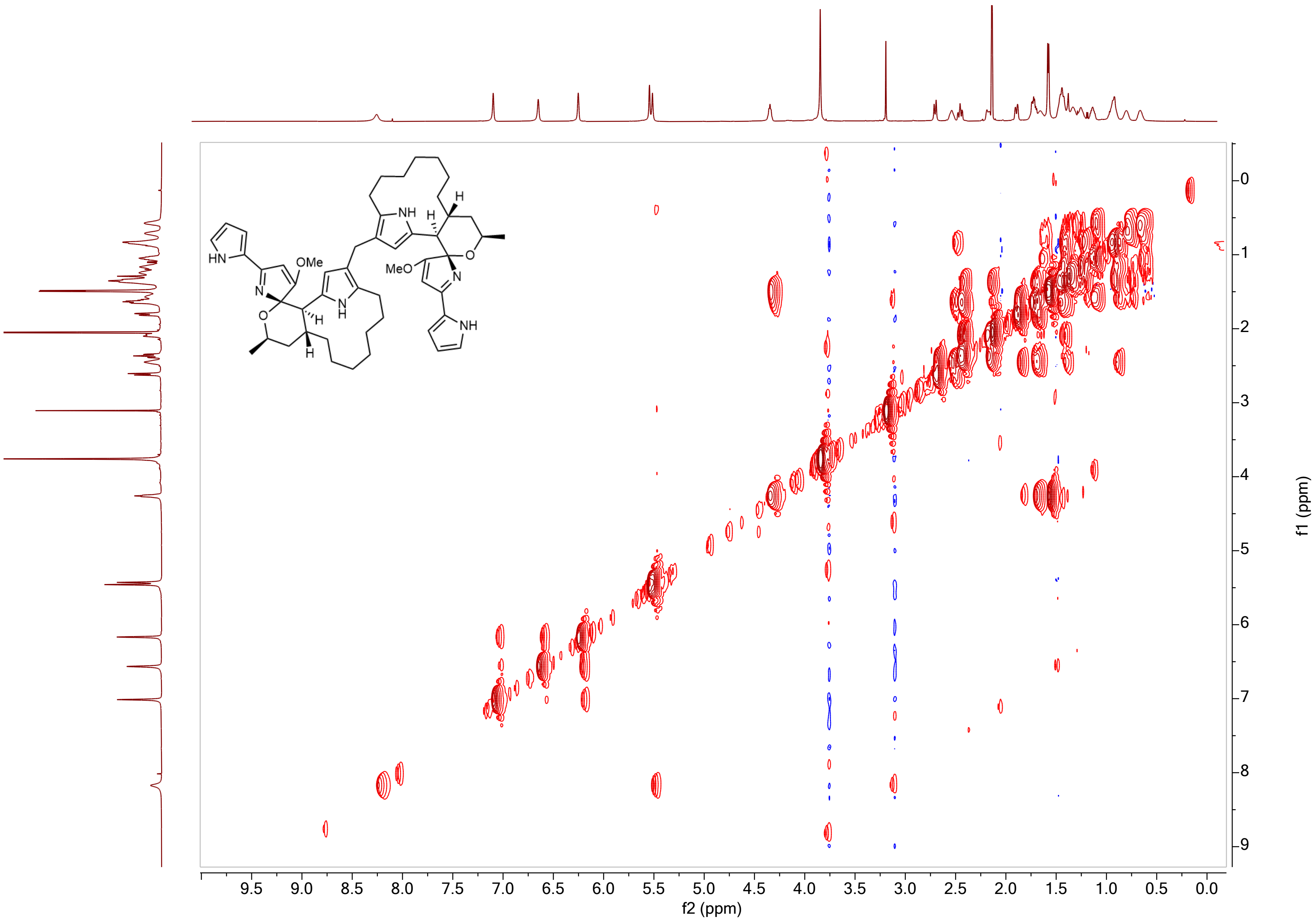

**Supplementary Figure 26.** $^{1}H$-$^{1}H$ COSY NMR Spectrum of methylene-bridged premarineosin A (**4**) in Acetone-$D_6$.

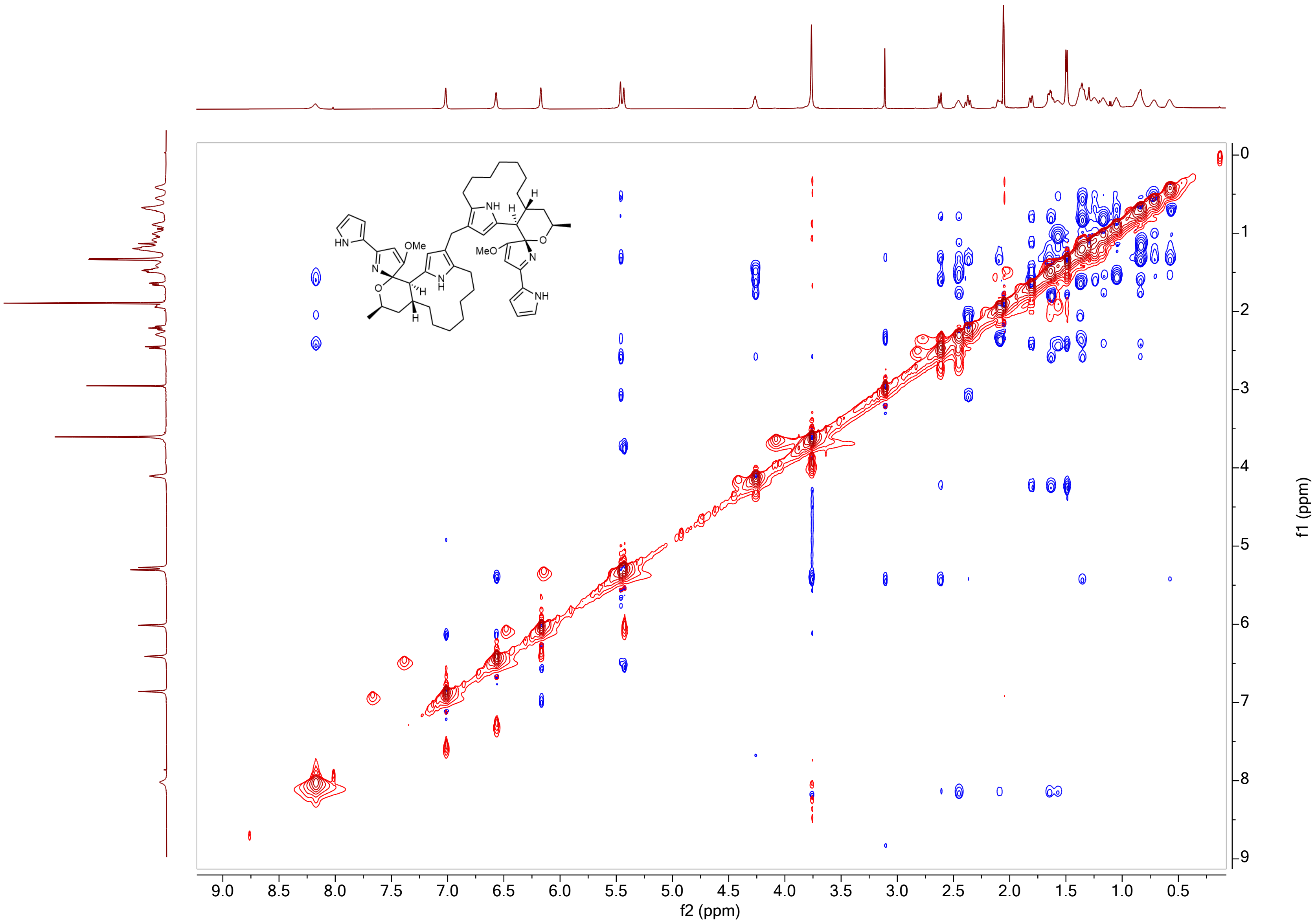

**Supplementary Figure 27.** $^1$H-$^1$H NOESY NMR Spectrum of methylene-bridged premarineosin A (**4**) in Acetone-D$_6$.

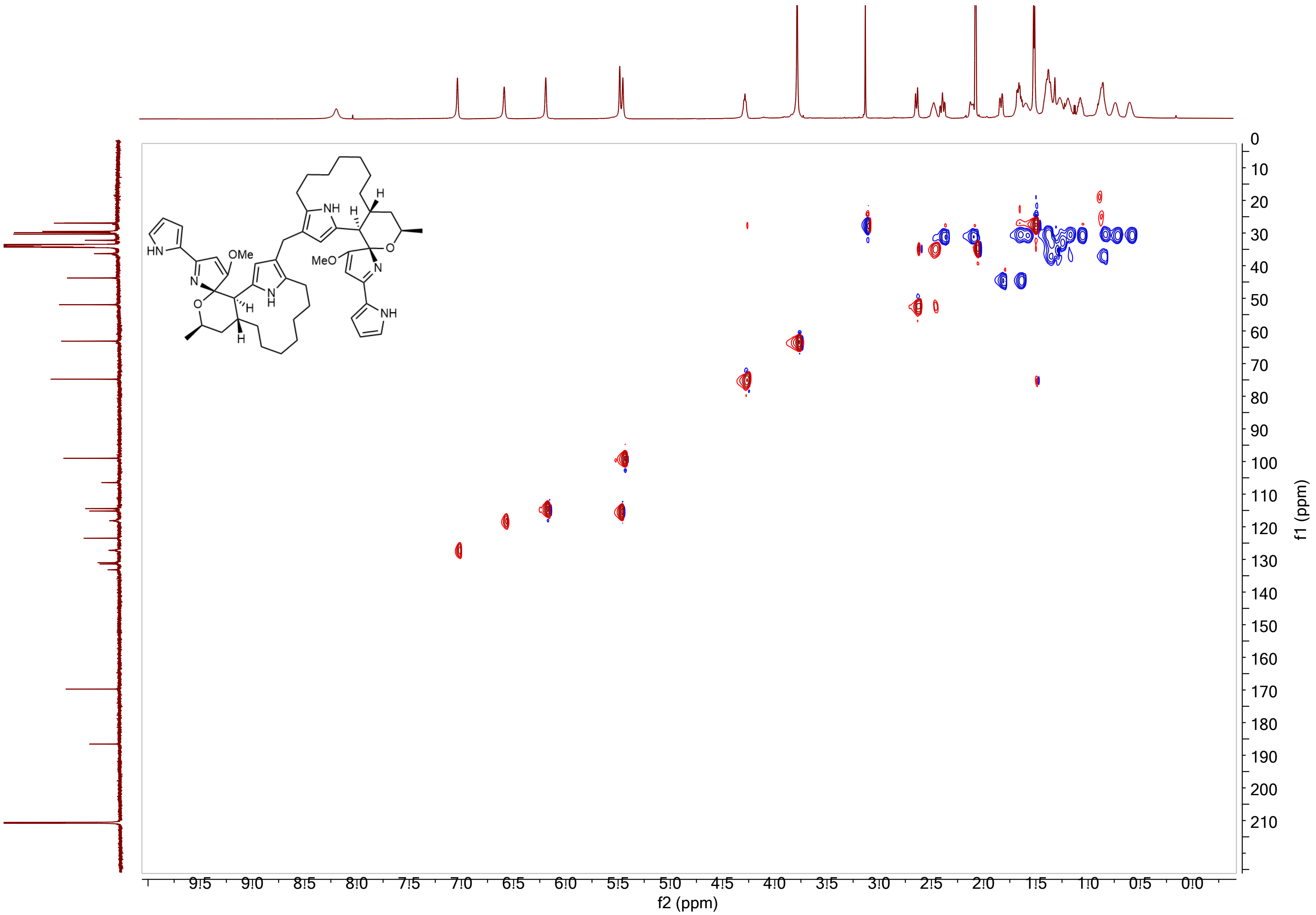

**Supplementary Figure 28.** $^{1}H$-$^{13}C$ HSQC NMR Spectrum of methylene-bridged premarineosin A (**4**) in Acetone-$D_6$.

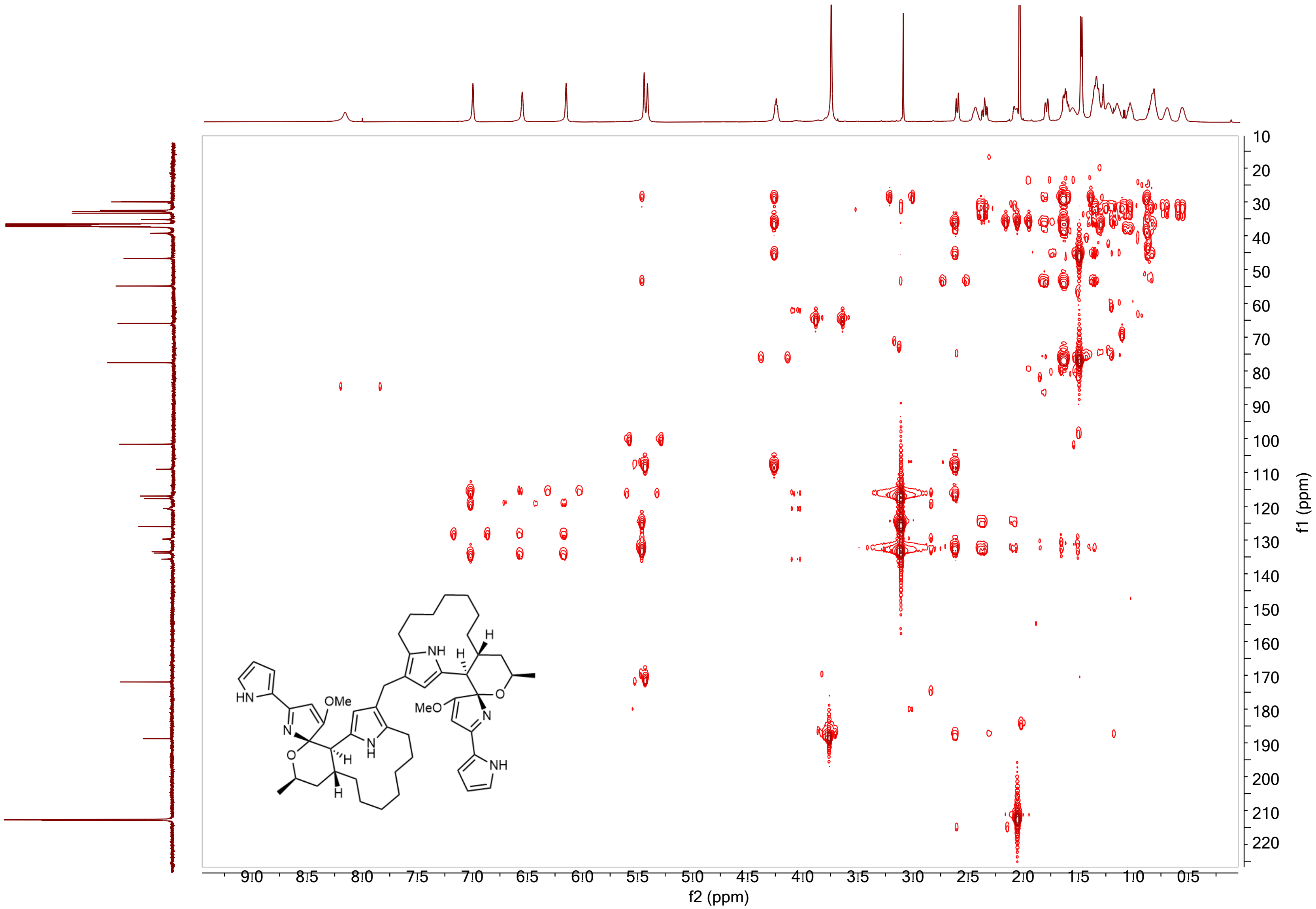

**Supplementary Figure 29.** $^{1}H$-$^{13}C$ HMBC NMR Spectrum of methylene-bridged premarineosin A (**4**) in Acetone-$D_6$.

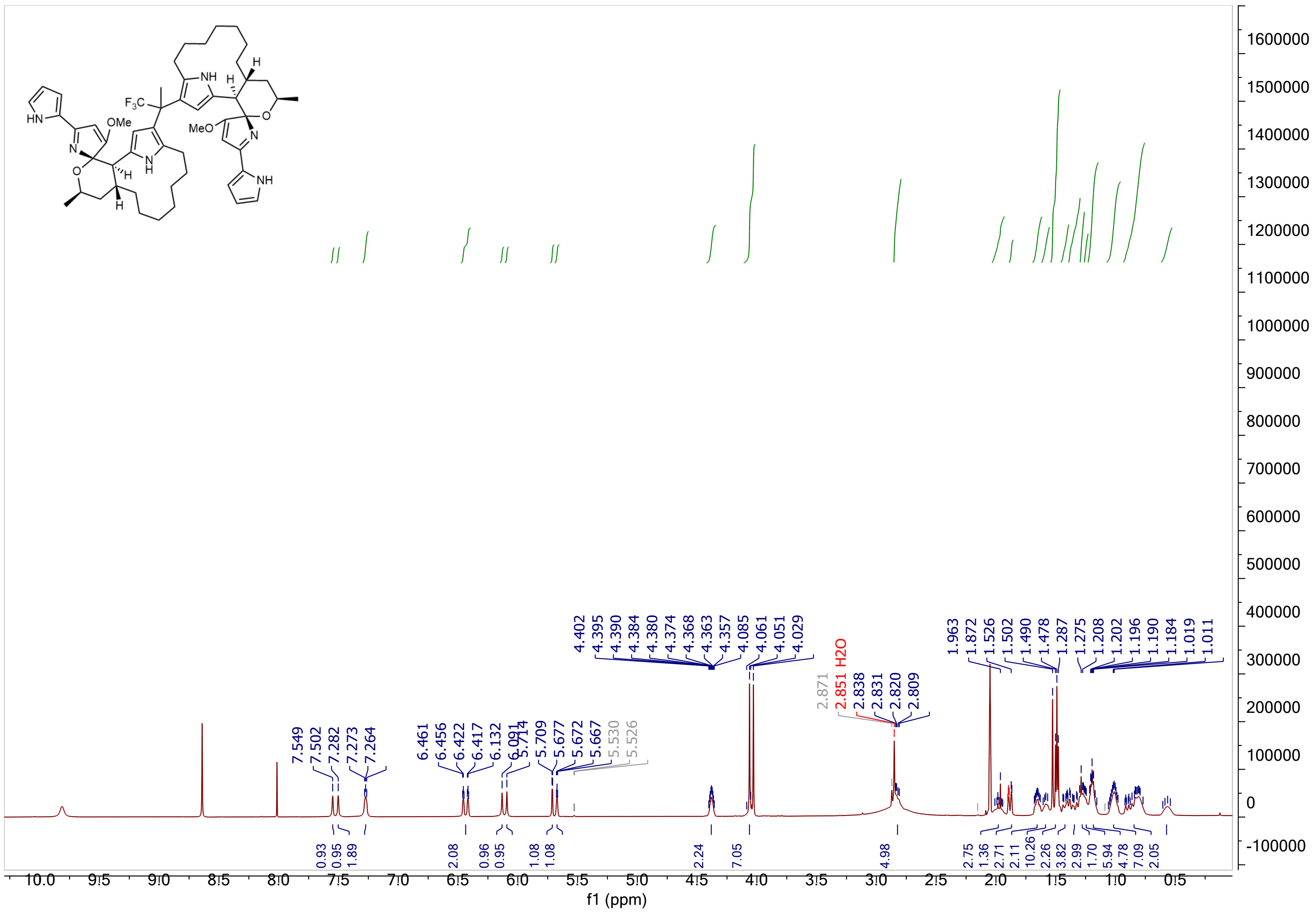

**Supplementary Figure 30.** $^{1}H$ NMR Spectrum of trifluoromethyl-bridged premarineosin A (**5**) in Acetone-D$_6$.

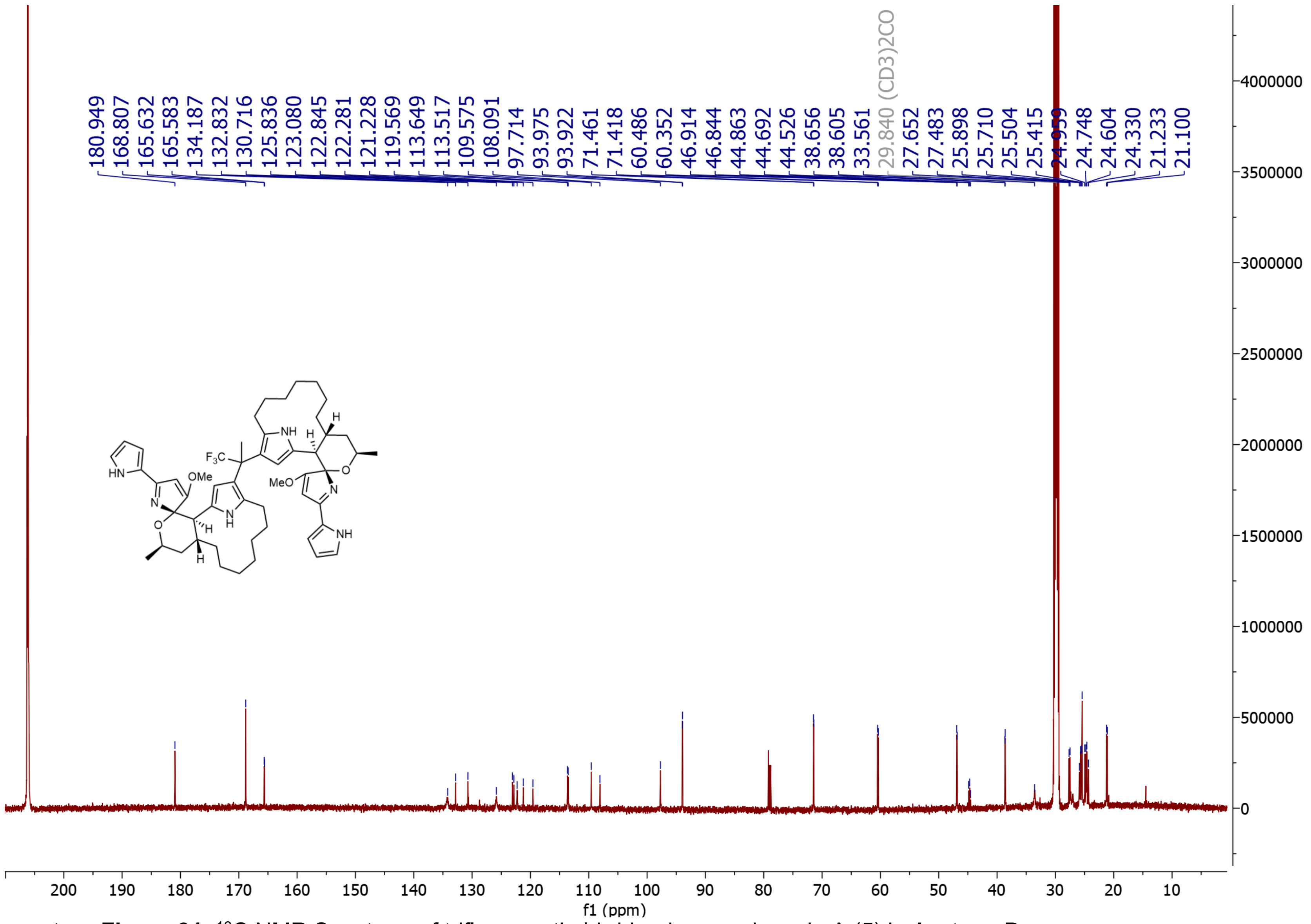

**Supplementary Figure 31.** $^{13}C$ NMR Spectrum of trifluoromethyl-bridged premarineosin A (**5**) in Acetone-$D_6$.

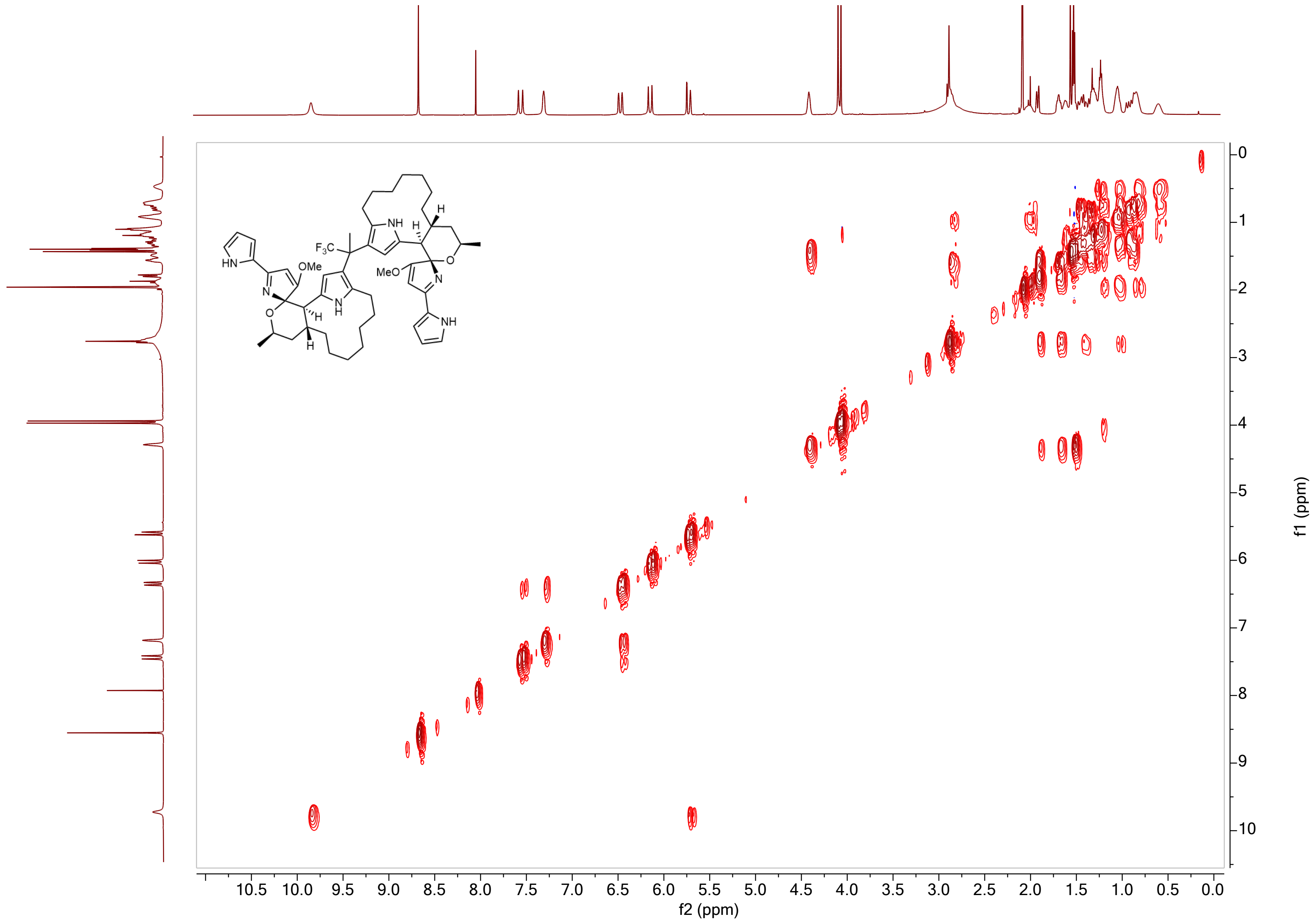

**Supplementary Figure 32.** $^{1}H$-$^{1}H$ COSY NMR Spectrum of trifluoromethyl-bridged premarineosin A (**5**) in Acetone-$D_6$.

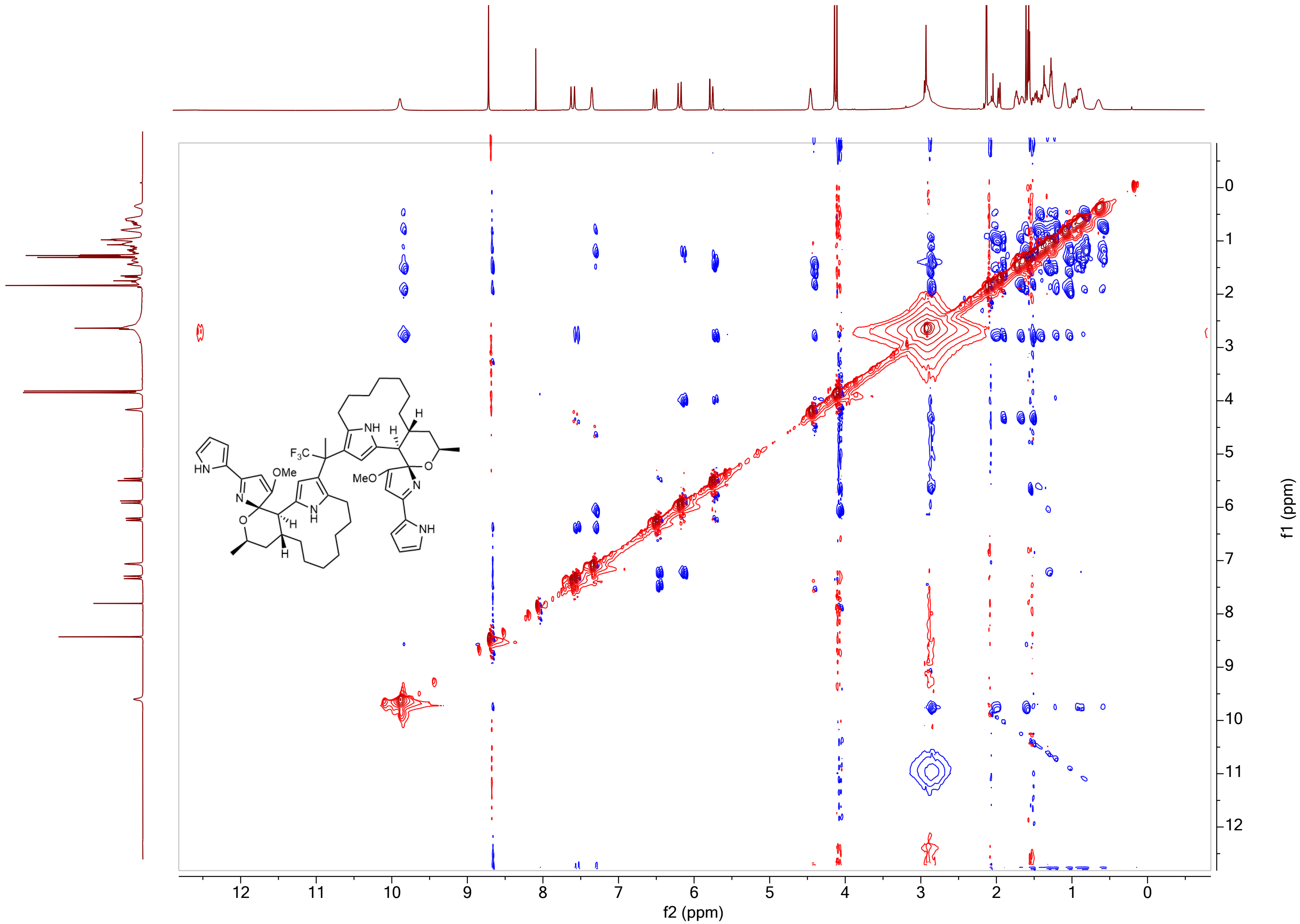

**Supplementary Figure 33.** ¹H-¹H NOESY NMR Spectrum of trifluoromethyl-bridged premarineosin A (**5**) in Acetone-$D_6$.

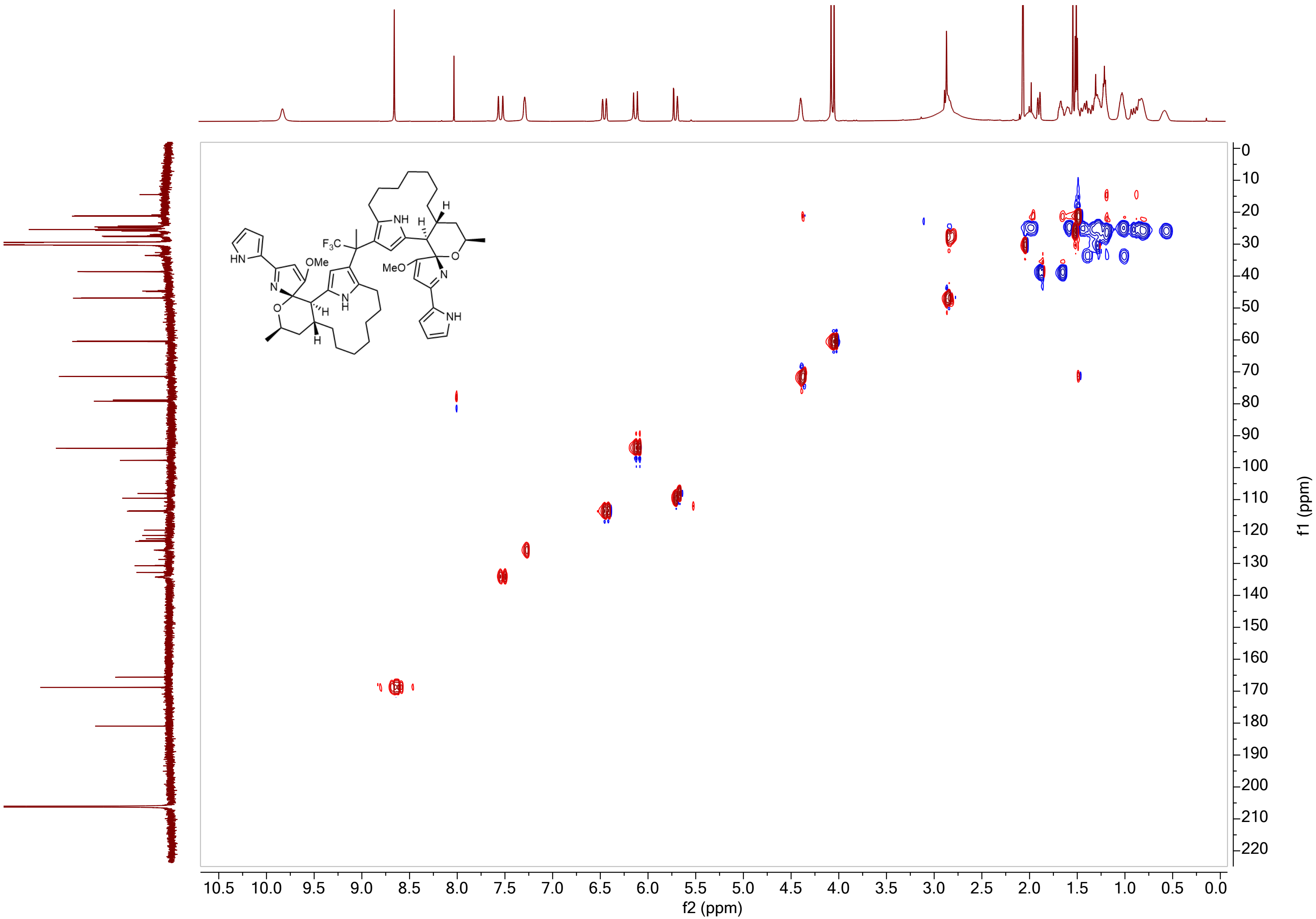

**Supplementary Figure 34.** $^{1}H$-$^{13}C$ HSQC NMR Spectrum of trifluoromethyl-bridged premarineosin A (**5**) in Acetone-$D_6$.

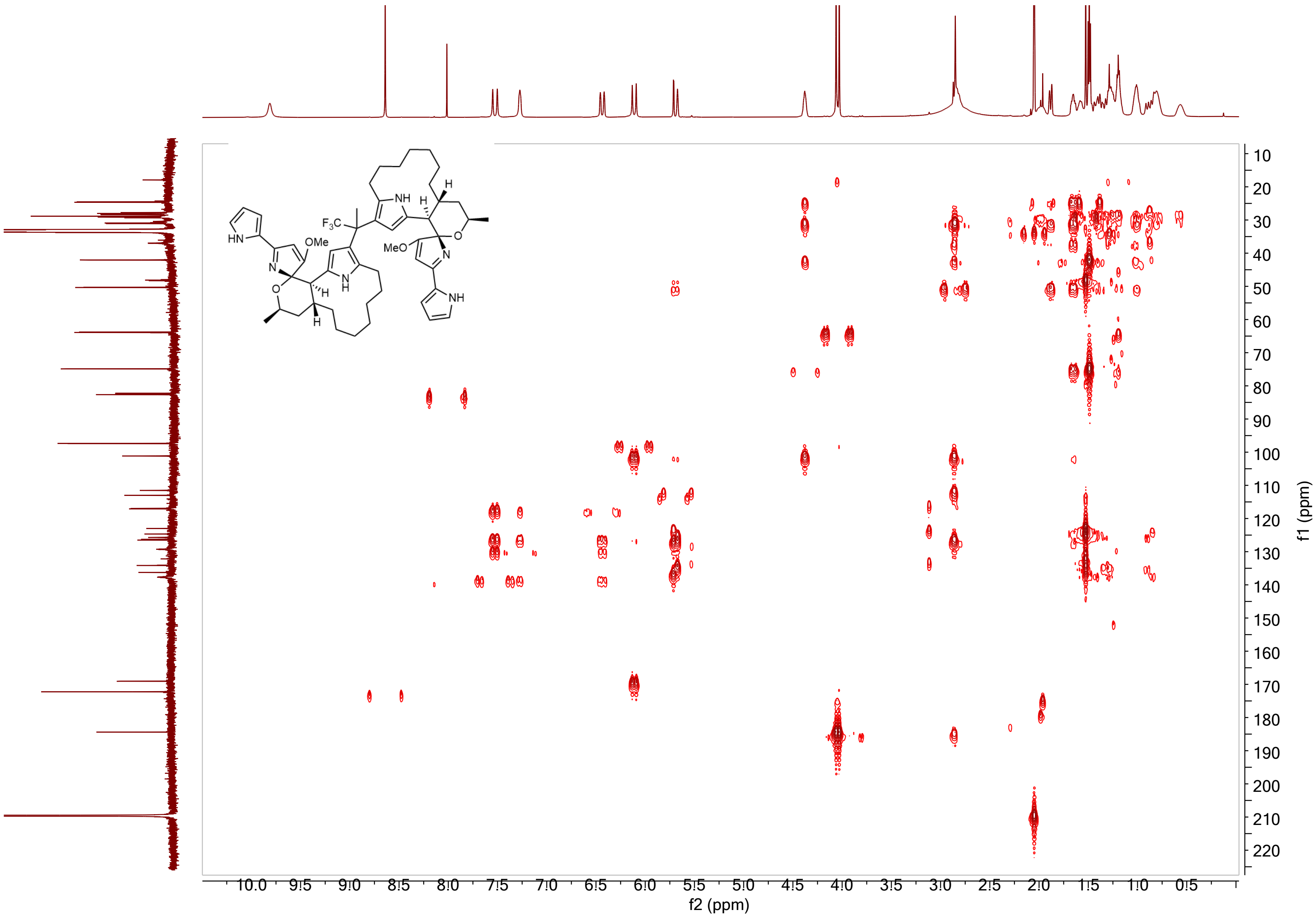

**Supplementary Figure 35.** $^{1}H$-$^{13}C$ HMBC NMR Spectrum of trifluoromethyl-bridged premarineosin A (**5**) in Acetone-$D_6$.

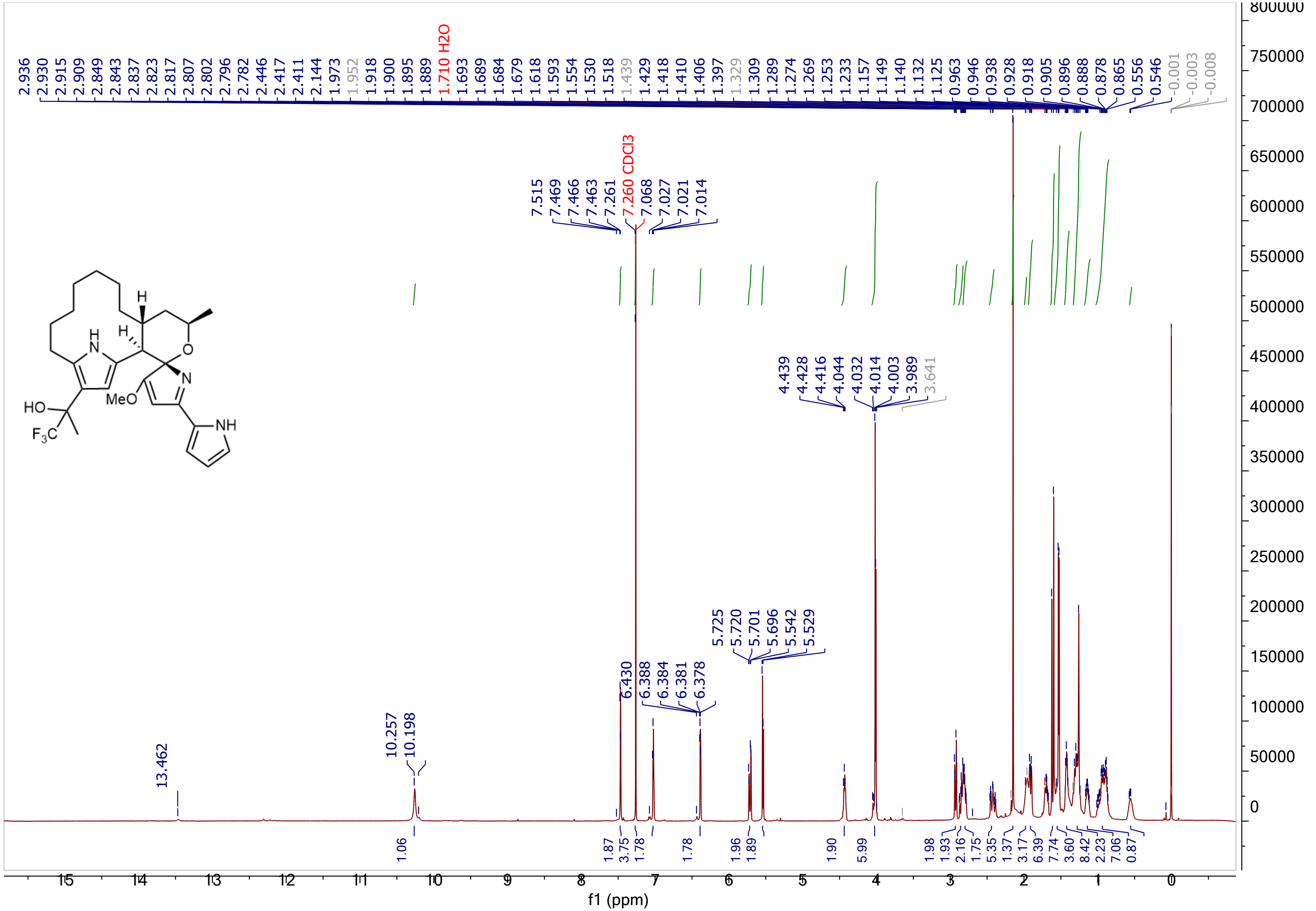

**Supplementary Figure 36.** [1]H NMR Spectrum of 12-trifluoropropanol premarineosin A (**6**) in Chloroform-D.

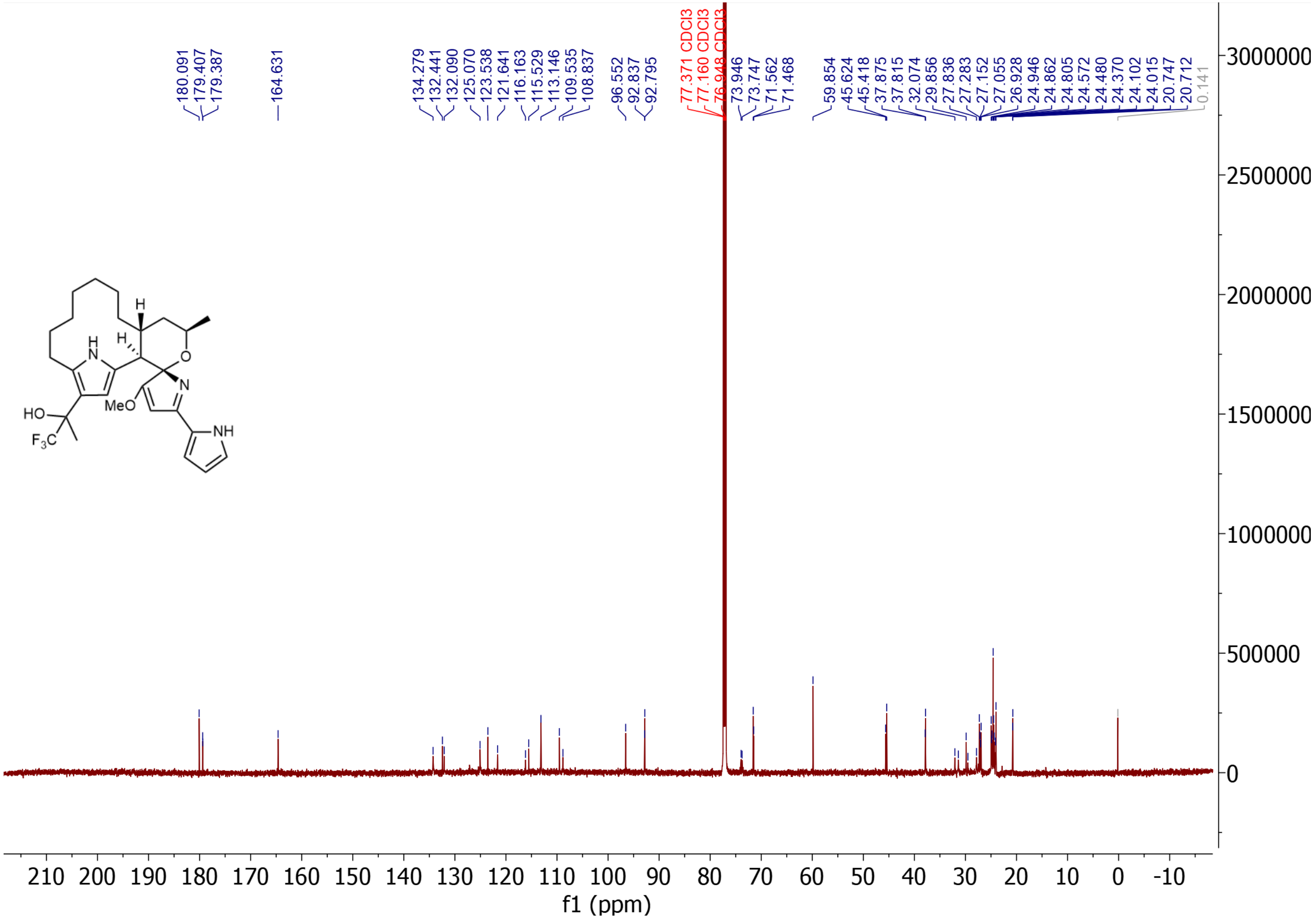

**Supplementary Figure 37.** $^{13}$C NMR Spectrum of 12-trifluoropropanol premarineosin A (**6**) in Chloroform-D.

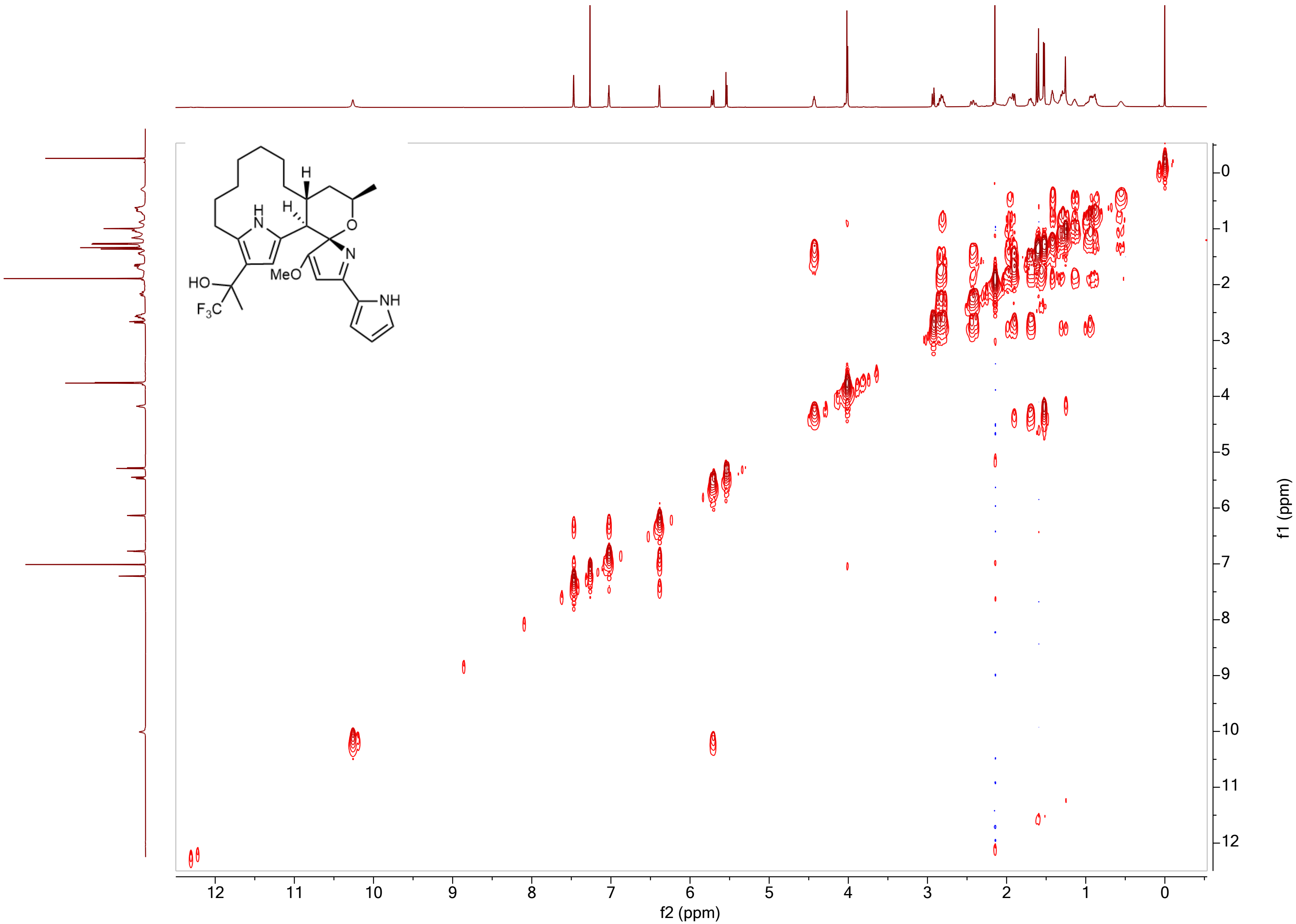

**Supplementary Figure 38.** $^1$H-$^1$H COSY NMR Spectrum of 12-trifluoropropanol premarineosin A (**6**) in Chloroform-D.

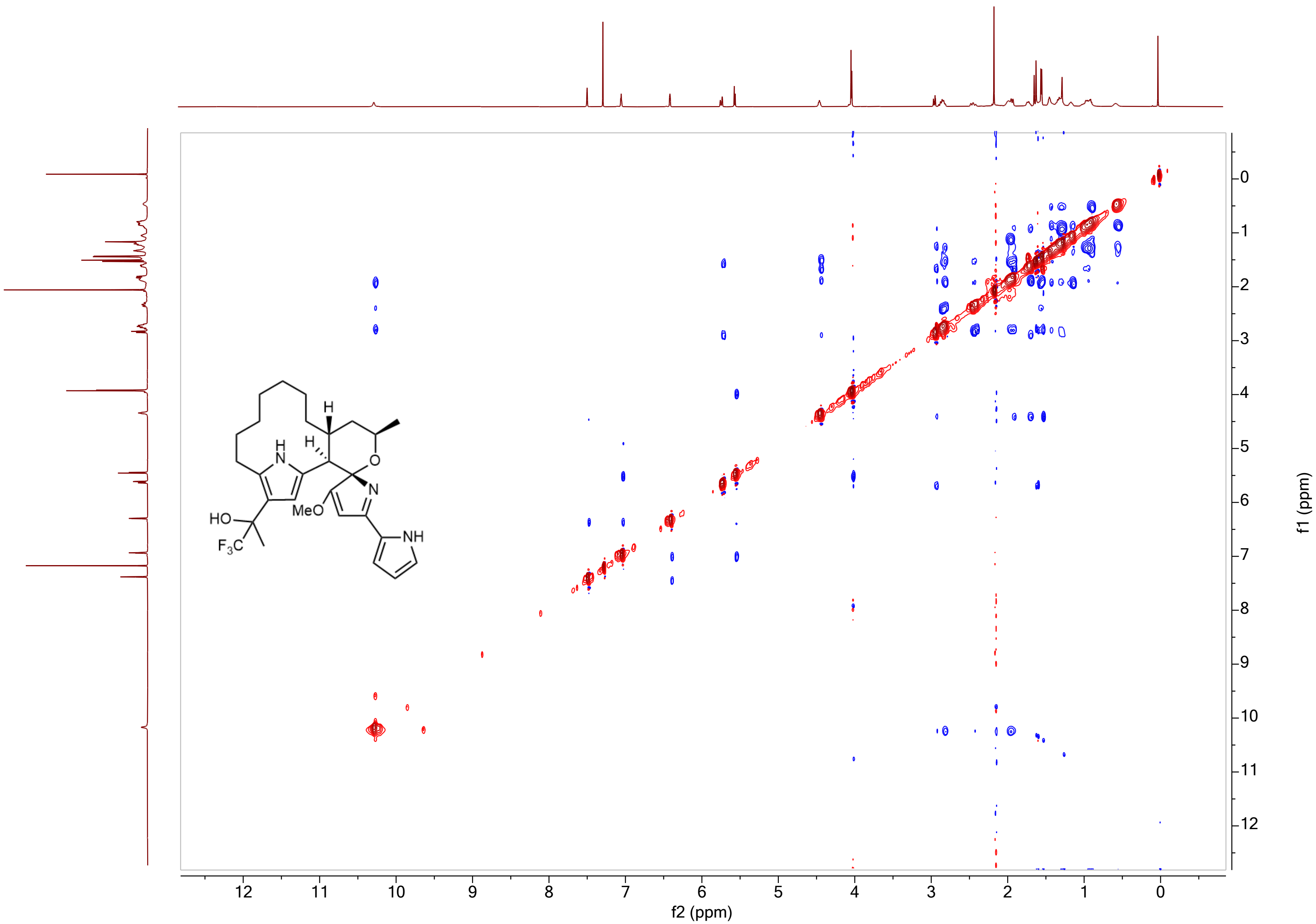

**Supplementary Figure 39.** $^{1}H$-$^{1}H$ NOESY NMR Spectrum of 12-trifluoropropanol premarineosin A (**6**) in Chloroform-D.

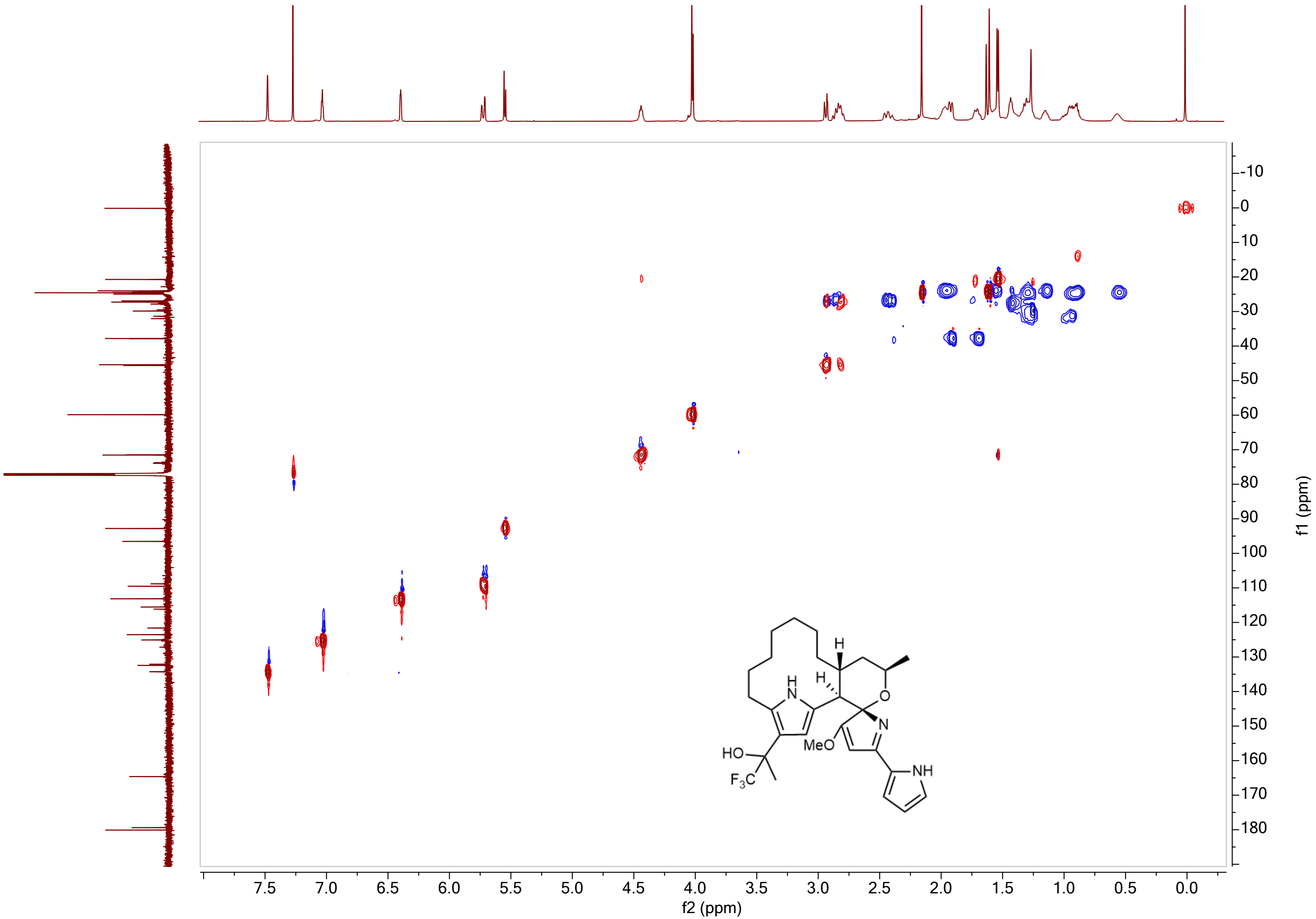

**Supplementary Figure 40.** $^{1}H$-$^{13}C$ HSQC NMR Spectrum of 12-trifluoropropanol premarineosin A (**6**) in Chloroform-D.

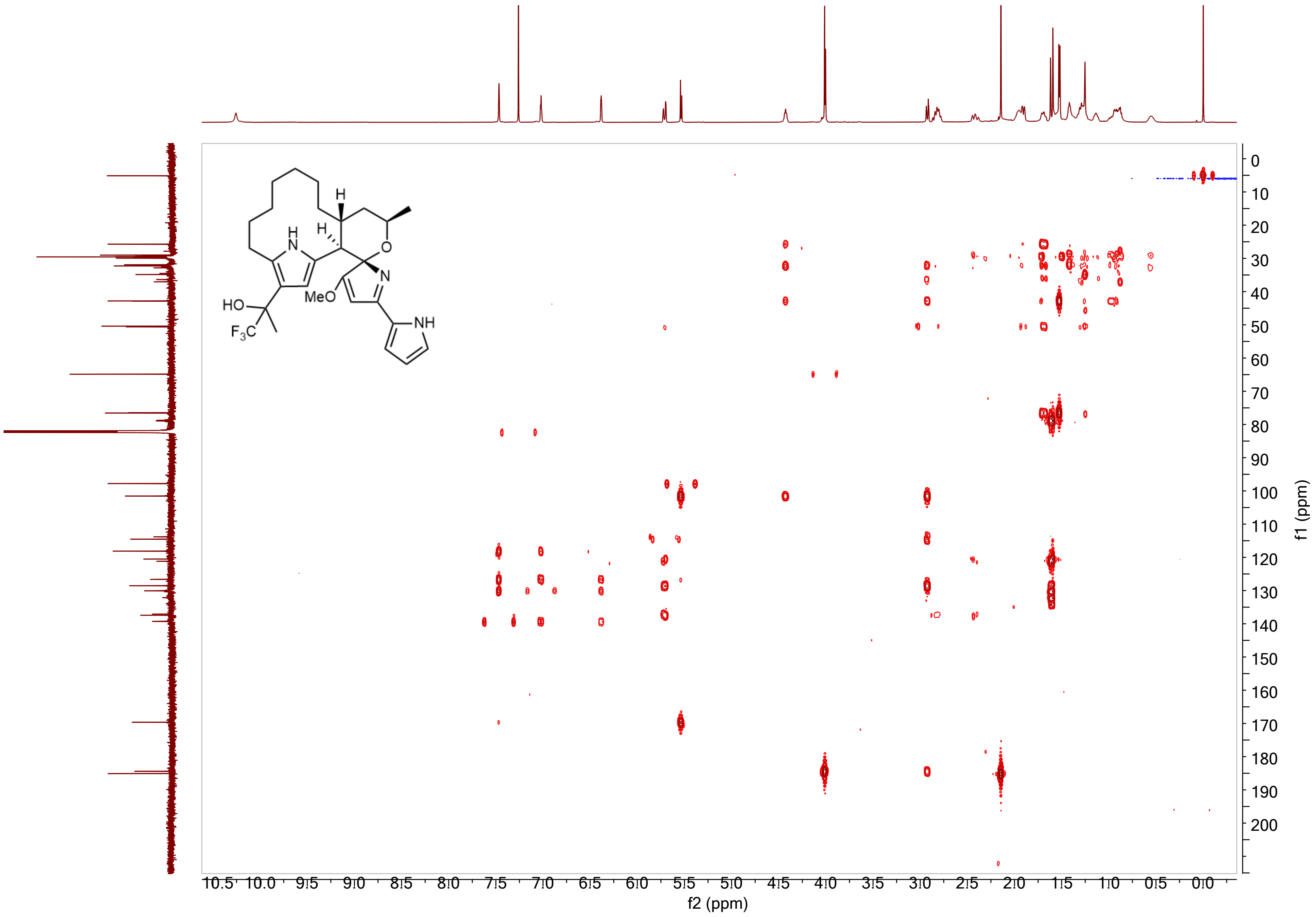

**Supplementary Figure 41.** $^{1}H$-$^{13}C$ HMBC NMR Spectrum of 12-trifluoropropanol premarineosin A (**6**) in Chloroform-D.

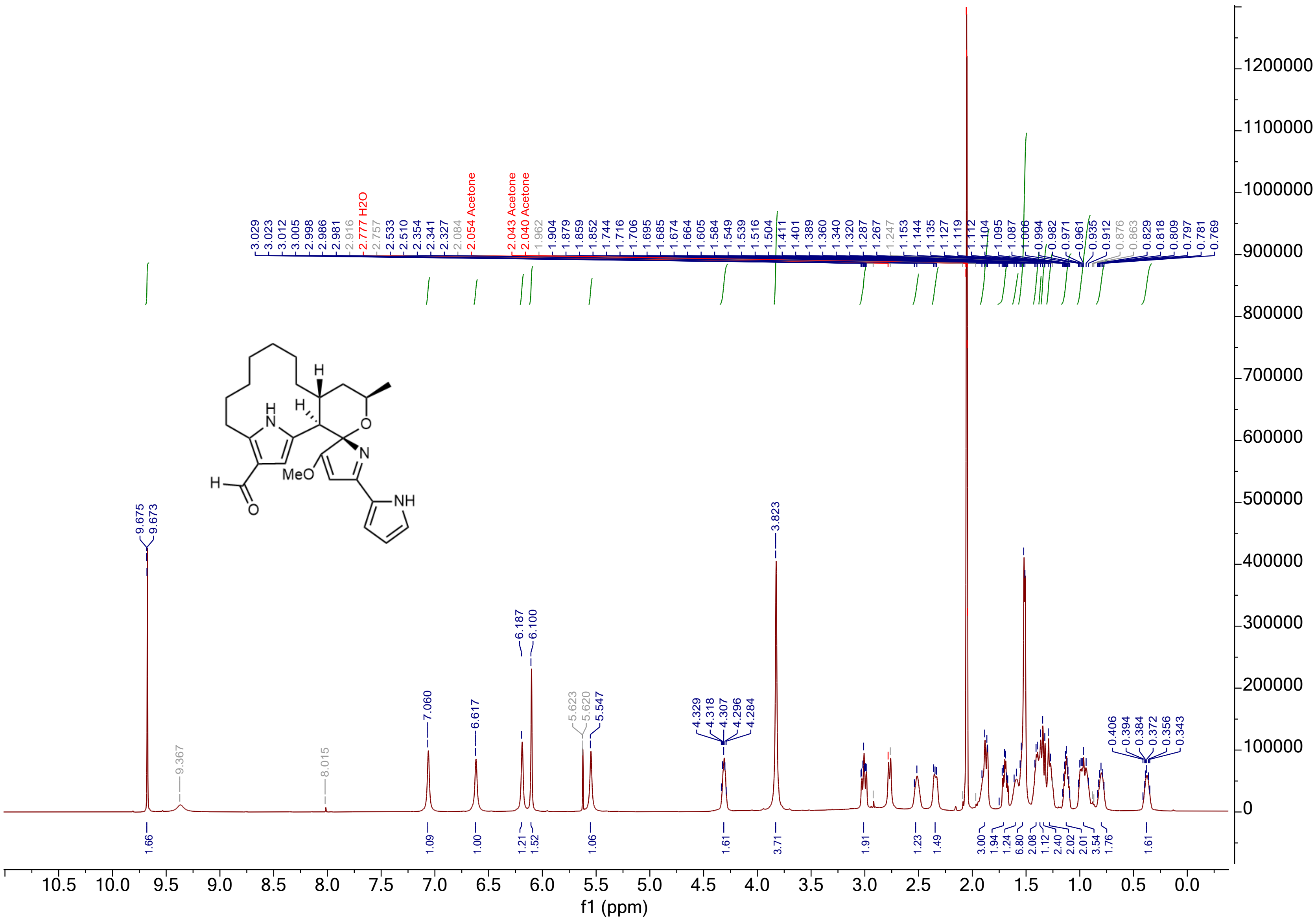

**Supplementary Figure 42.** $^{1}$H NMR Spectrum of 12-formyl premarineosin A (**7**) in Acetone-$D_6$.

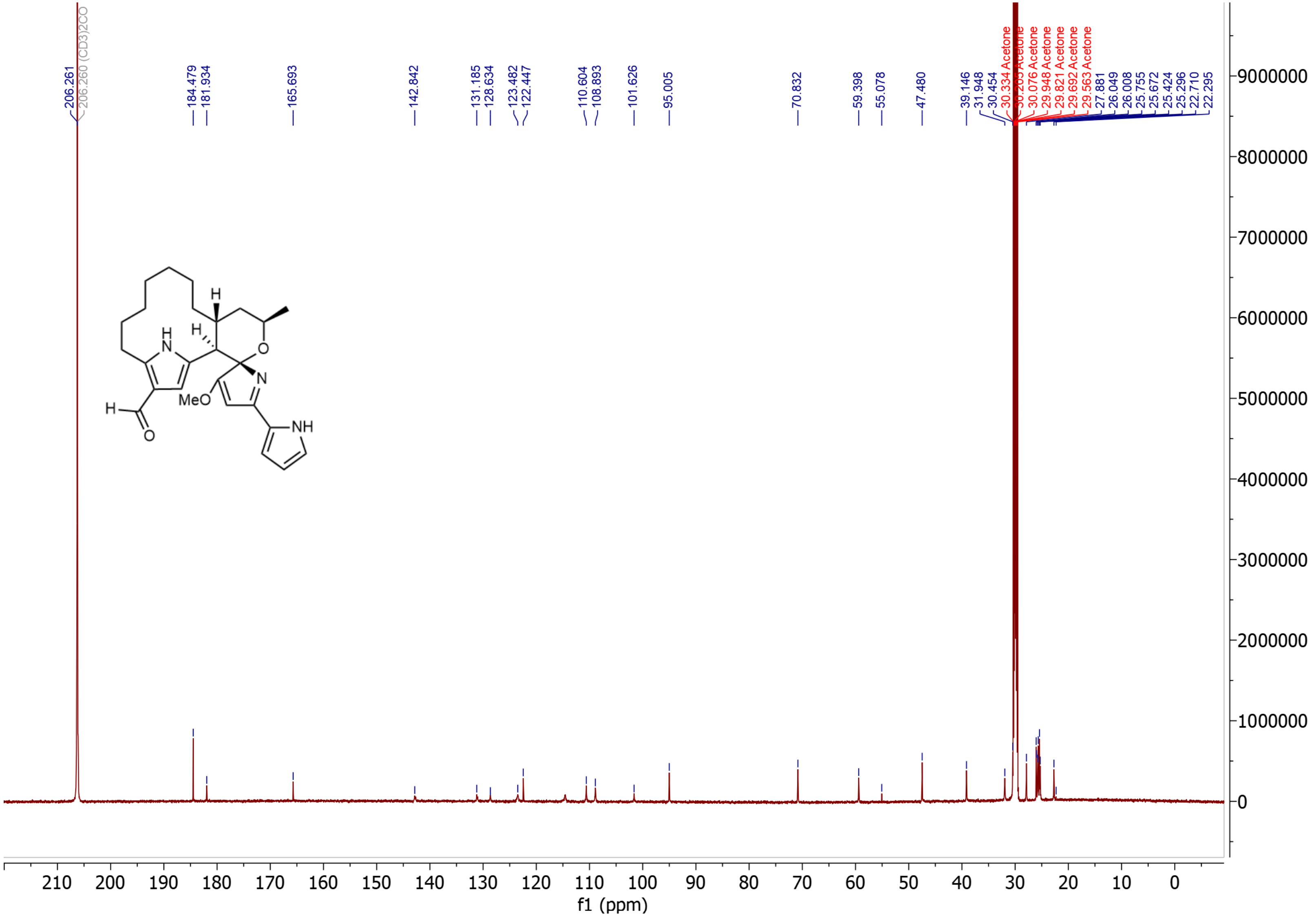

**Supplementary Figure 43.** $^{13}C$ NMR Spectrum of 12-formyl premarineosin A (**7**) in Acetone-$D_6$.

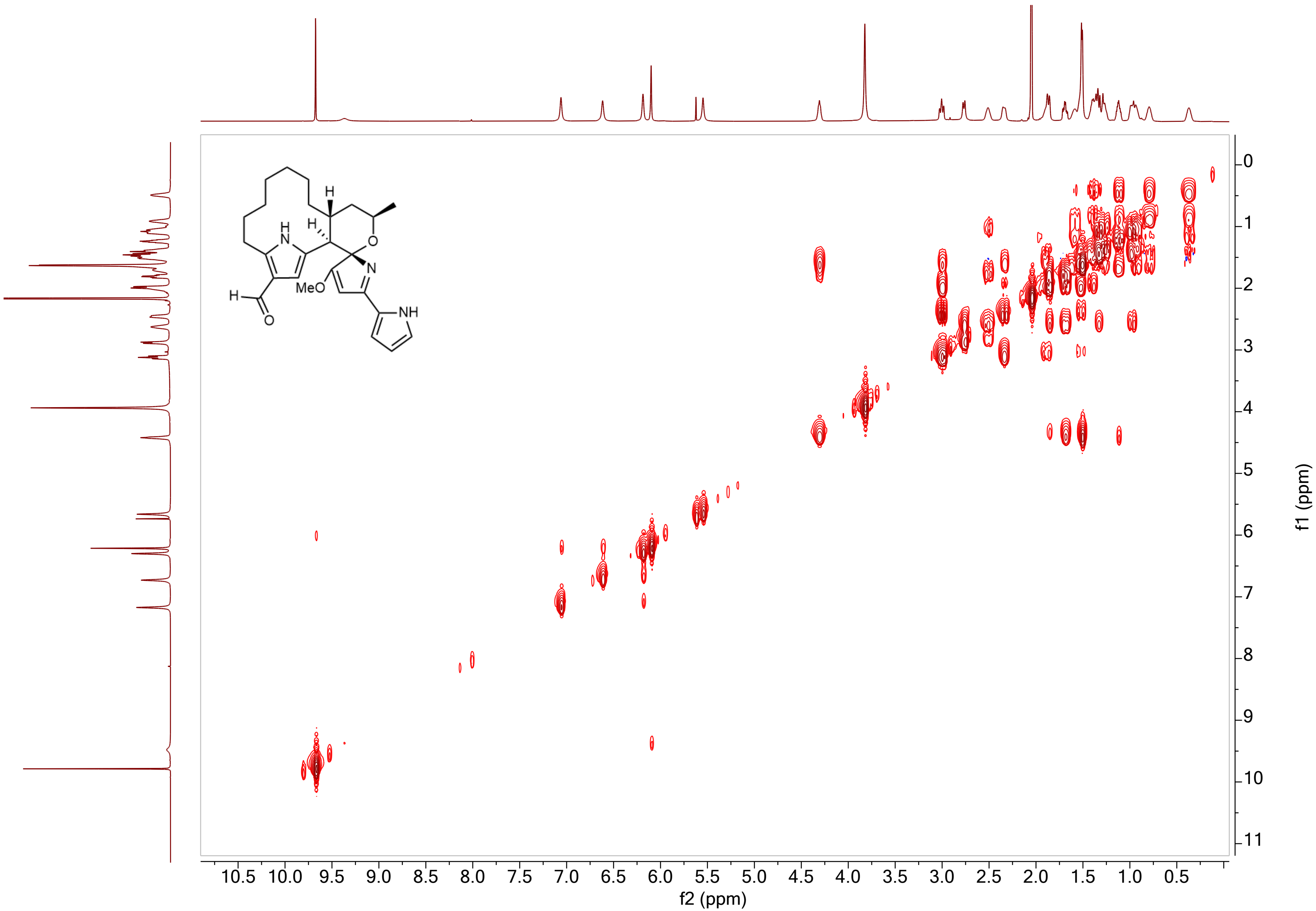

**Supplementary Figure 44.** 1H-1H COSY NMR Spectrum of 12-formyl premarineosin A (**7**) in Acetone-D6.

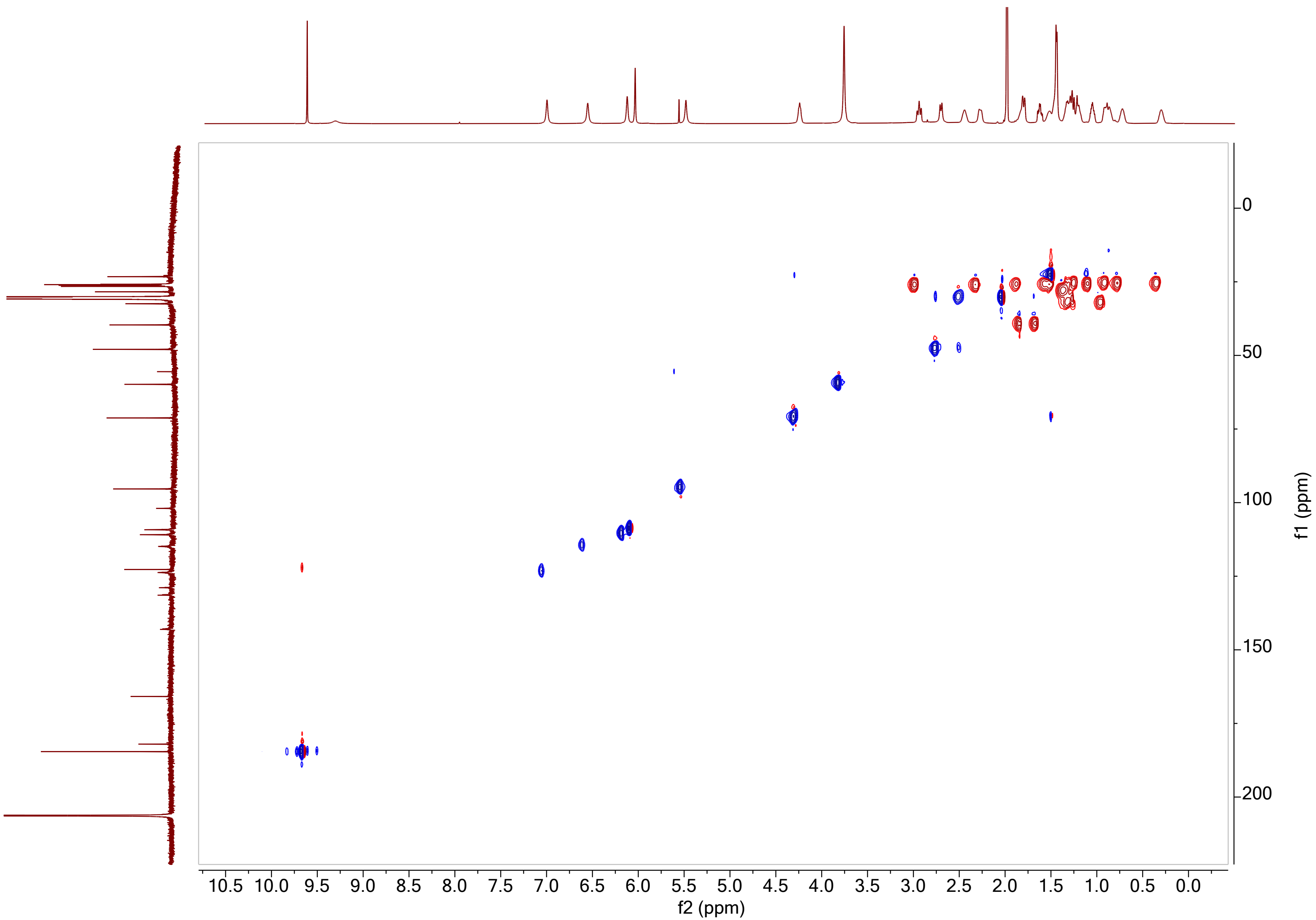

**Supplementary Figure 45.** $^{1}H$-$^{13}C$ HSQC NMR Spectrum of 12-formyl premarineosin A (**7**) in Acetone-$D_6$.

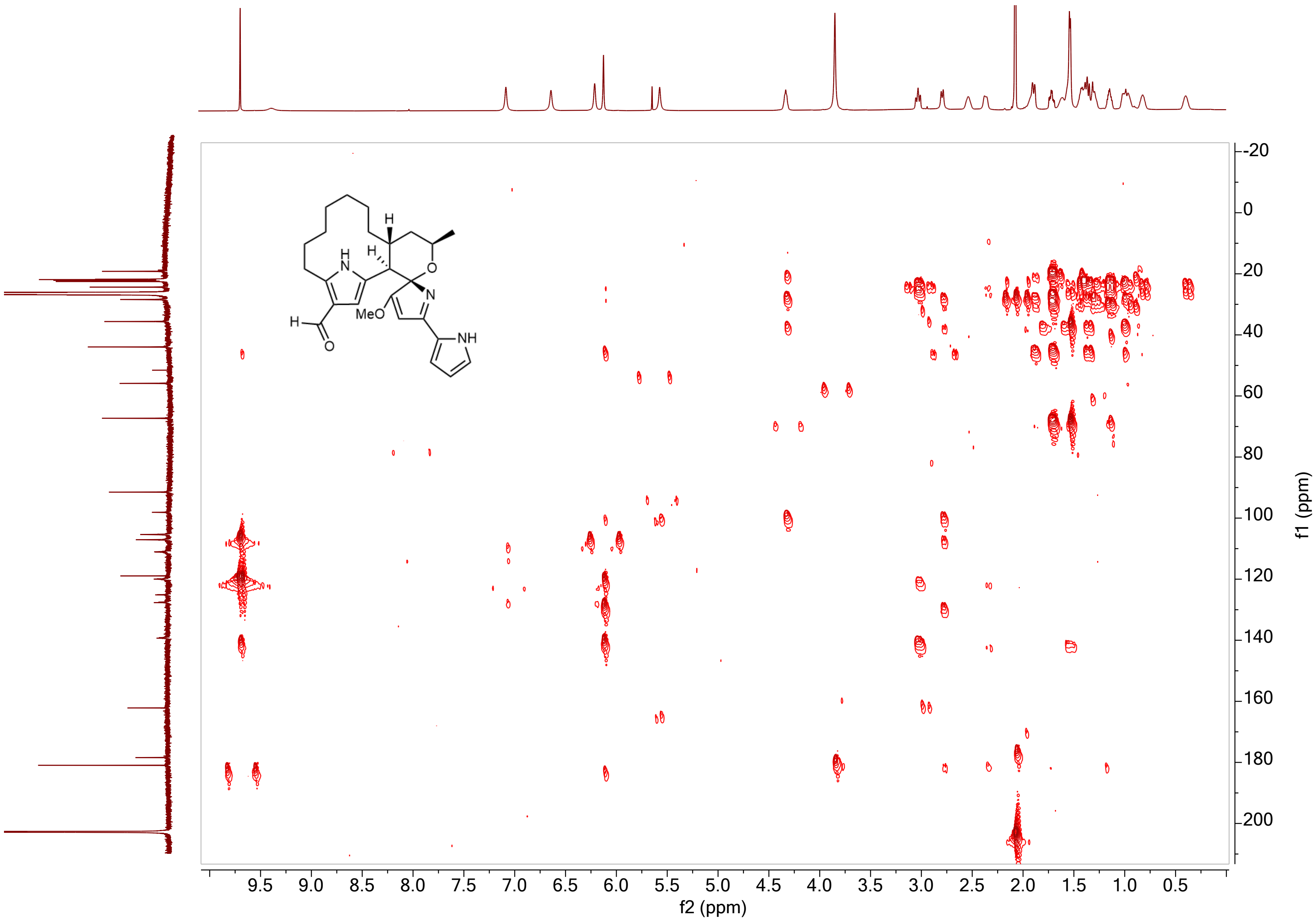

**Supplementary Figure 46.** $^{1}H$-$^{13}C$ HMBC NMR Spectrum of 12-formyl premarineosin A (**7**) in Acetone-$D_6$.

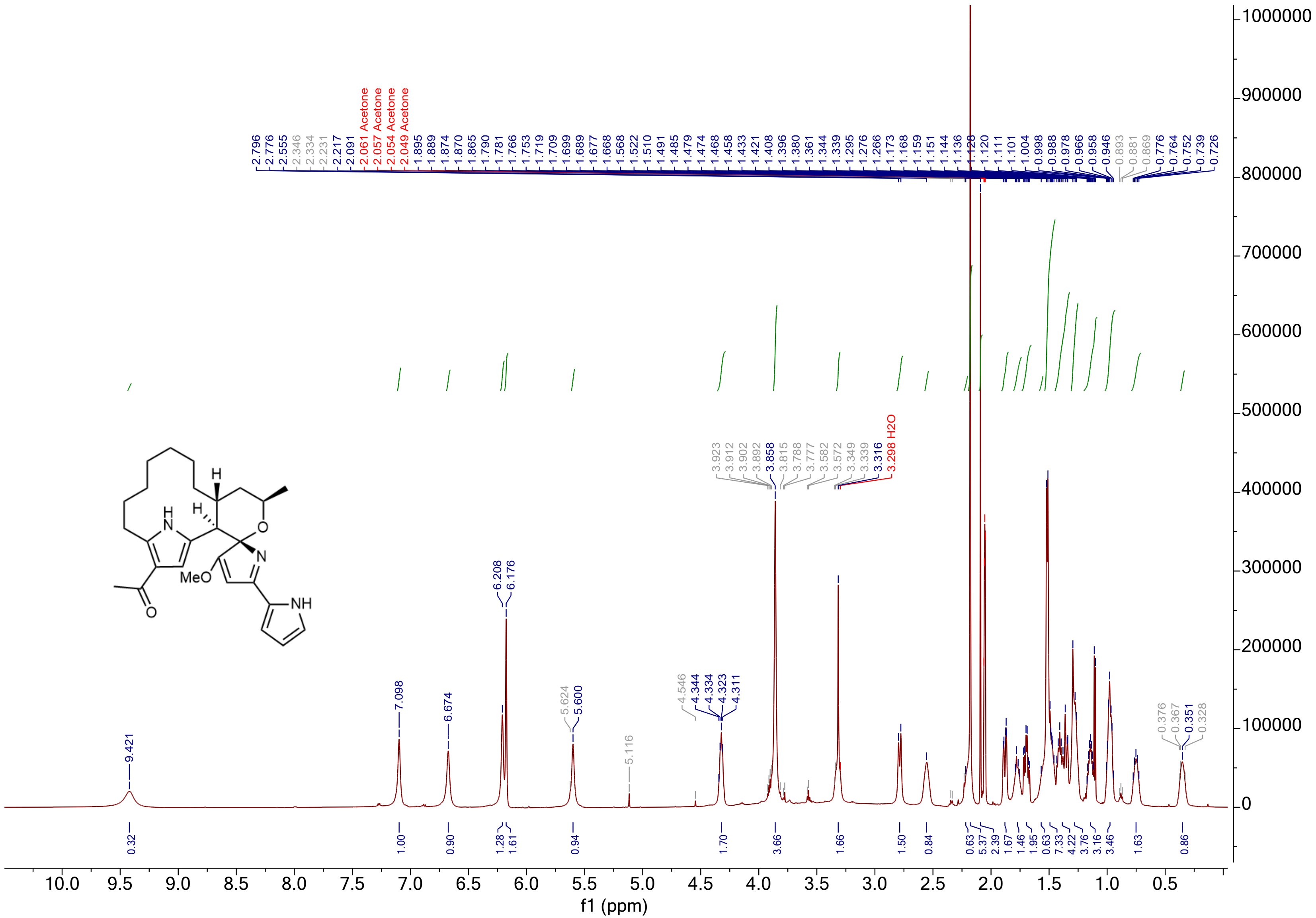

**Supplementary Figure 47.** $^1$H NMR Spectrum of 12-acetyl premarineosin A (**8**) in Acetone-$D_6$.

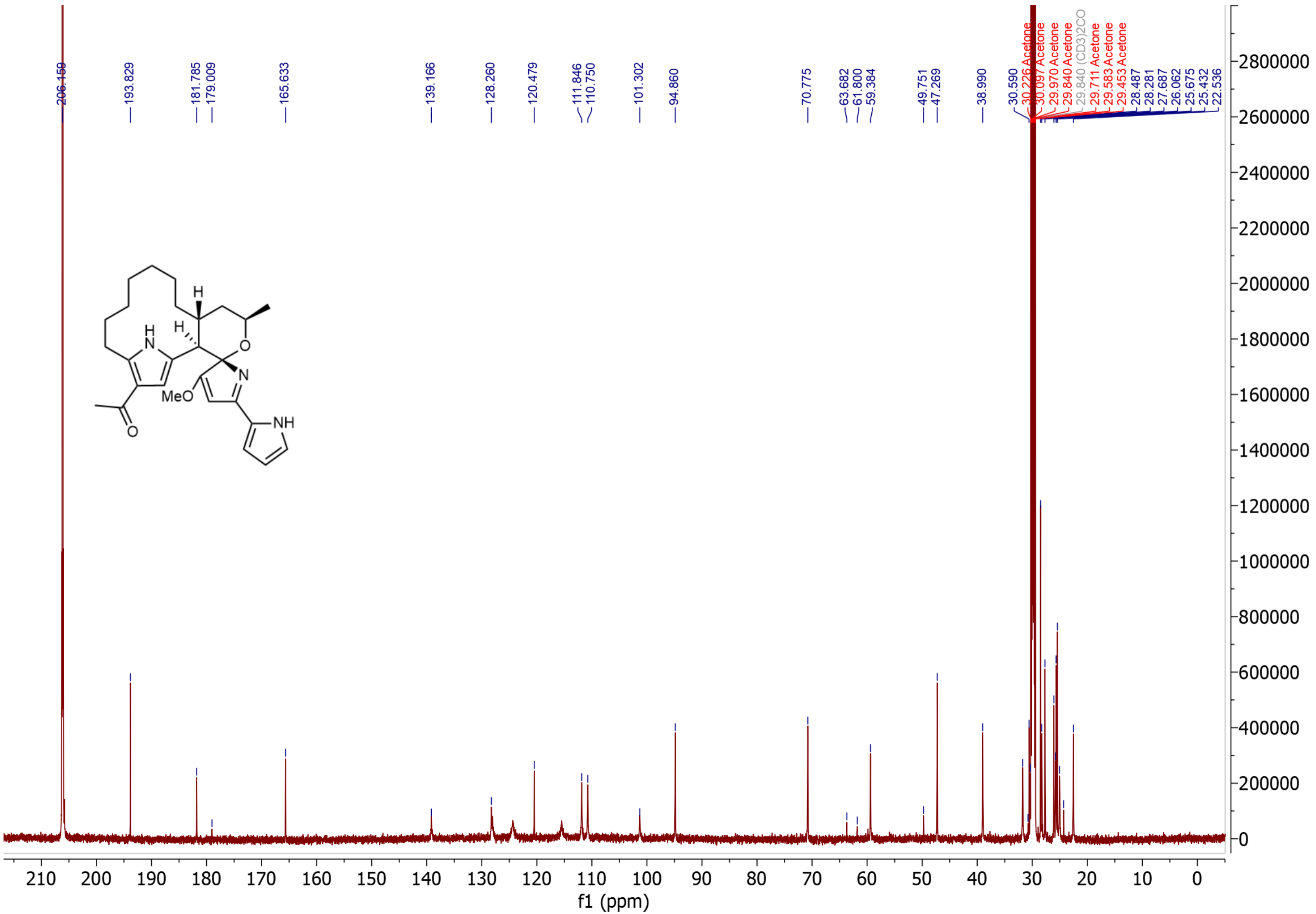

**Supplementary Figure 48.** $^{13}C$ NMR Spectrum of 12-acetyl premarineosin A (**8**) in Acetone-$D_6$.

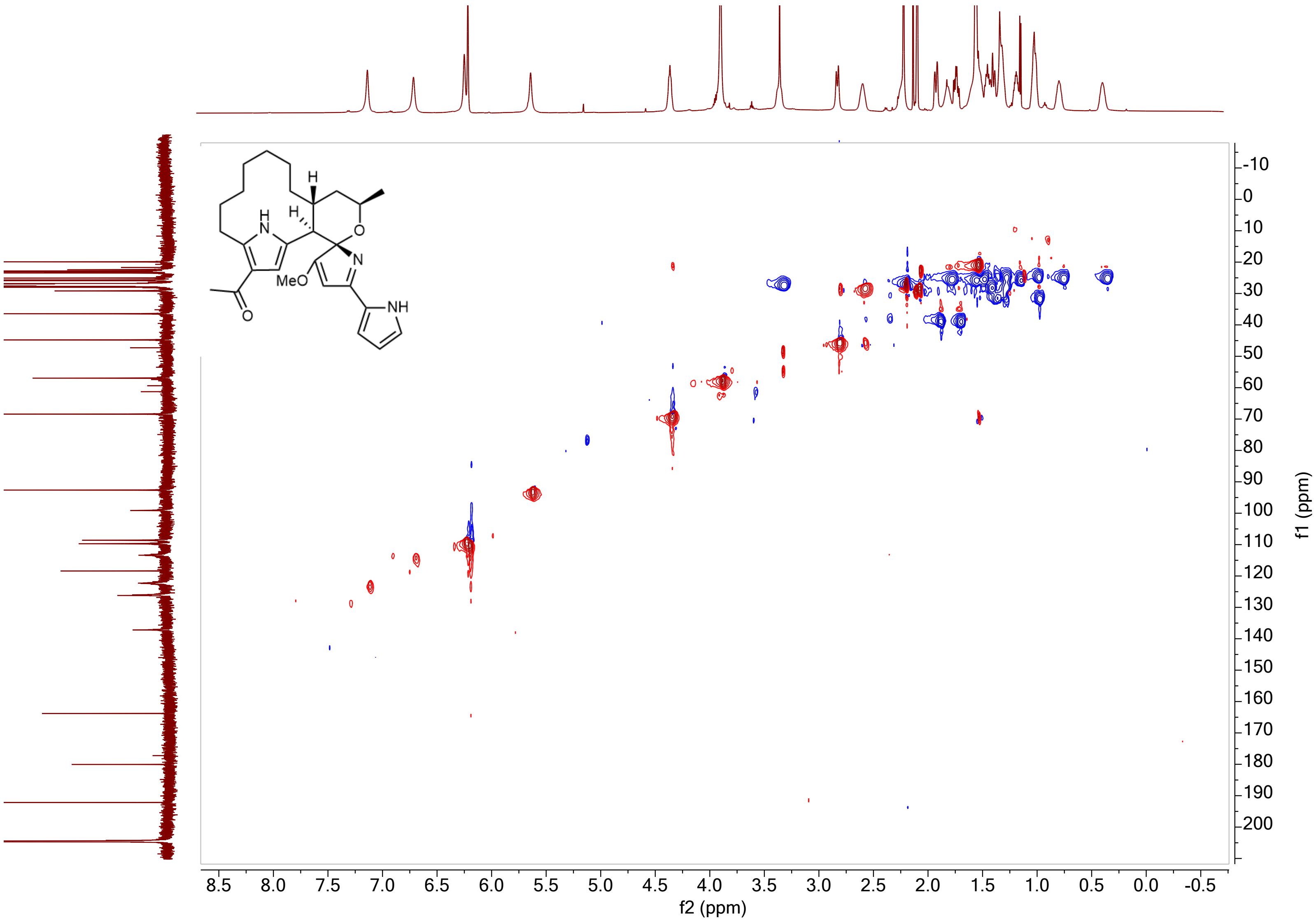

**Supplementary Figure 49.** $^{1}H$-$^{13}C$ HSQC NMR Spectrum of 12-acetyl premarineosin A (**8**) in Acetone-$D_6$.

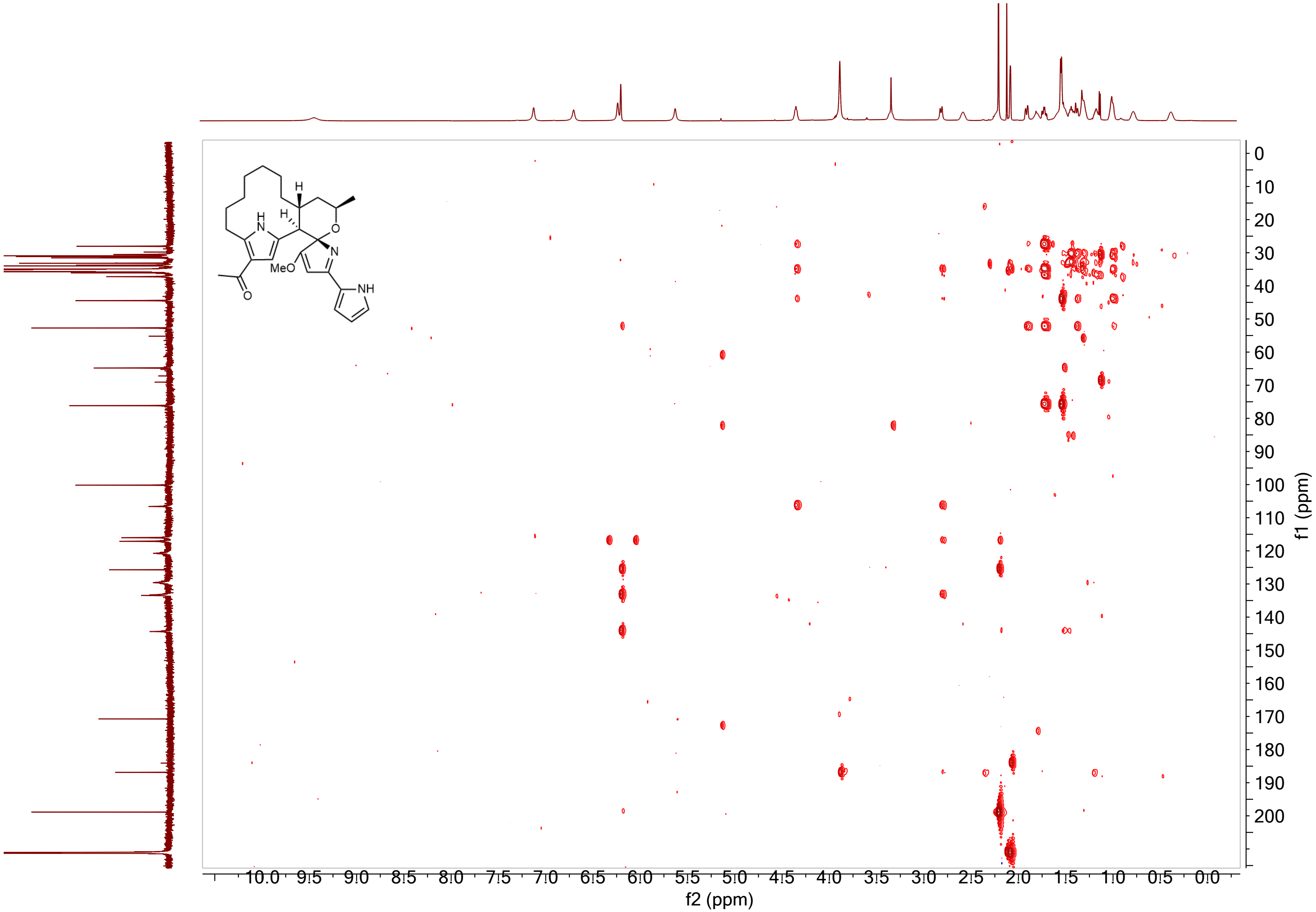

**Supplementary Figure 50.** $^{1}H$-$^{13}C$ HMBC NMR Spectrum of 12-acetyl premarineosin A (**8**) in Acetone-$D_6$.

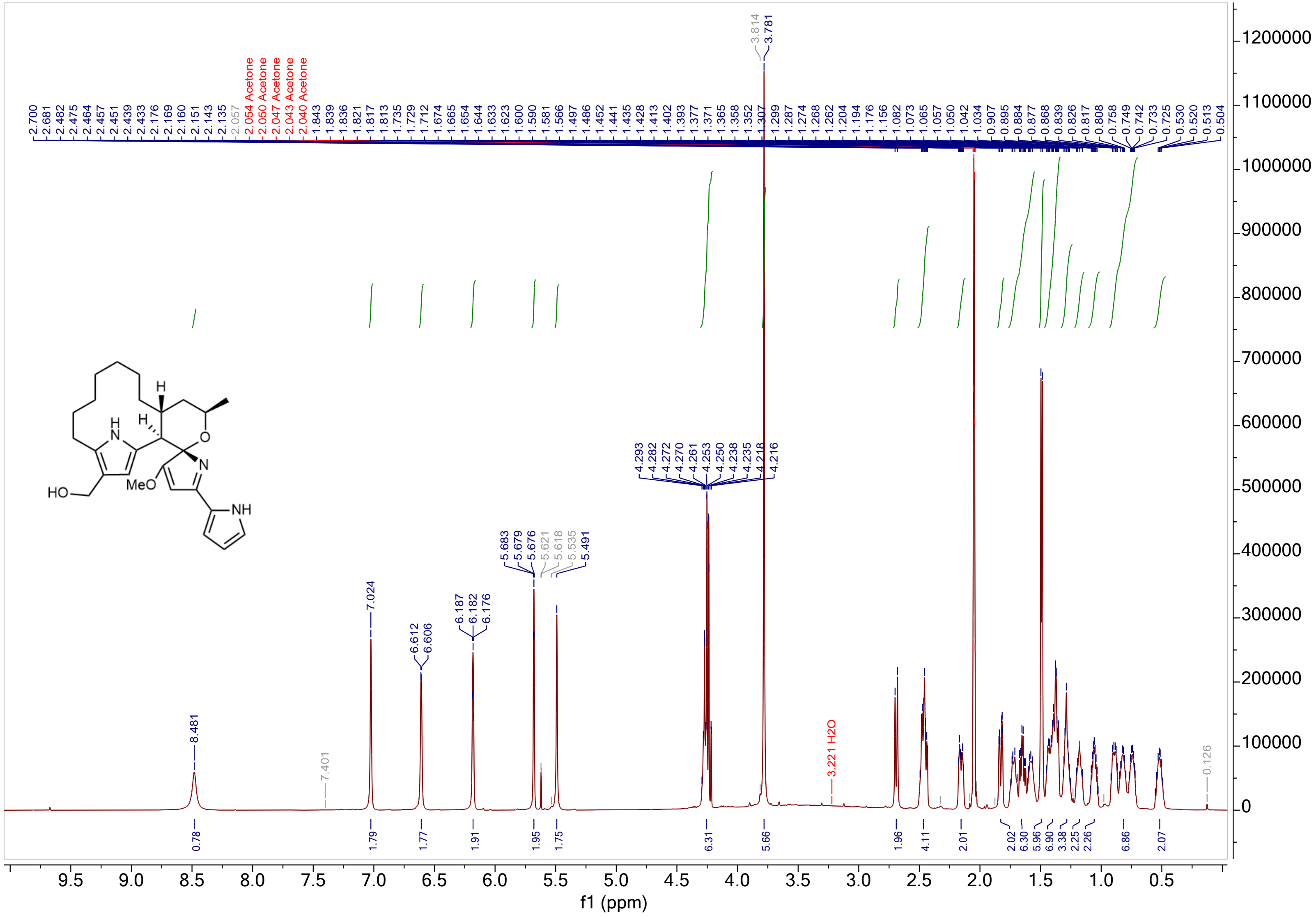

**Supplementary Figure 51.** $^{1}$H NMR Spectrum of 12-hydroxymethyl premarineosin A (**9**) in Acetone-D$_6$.

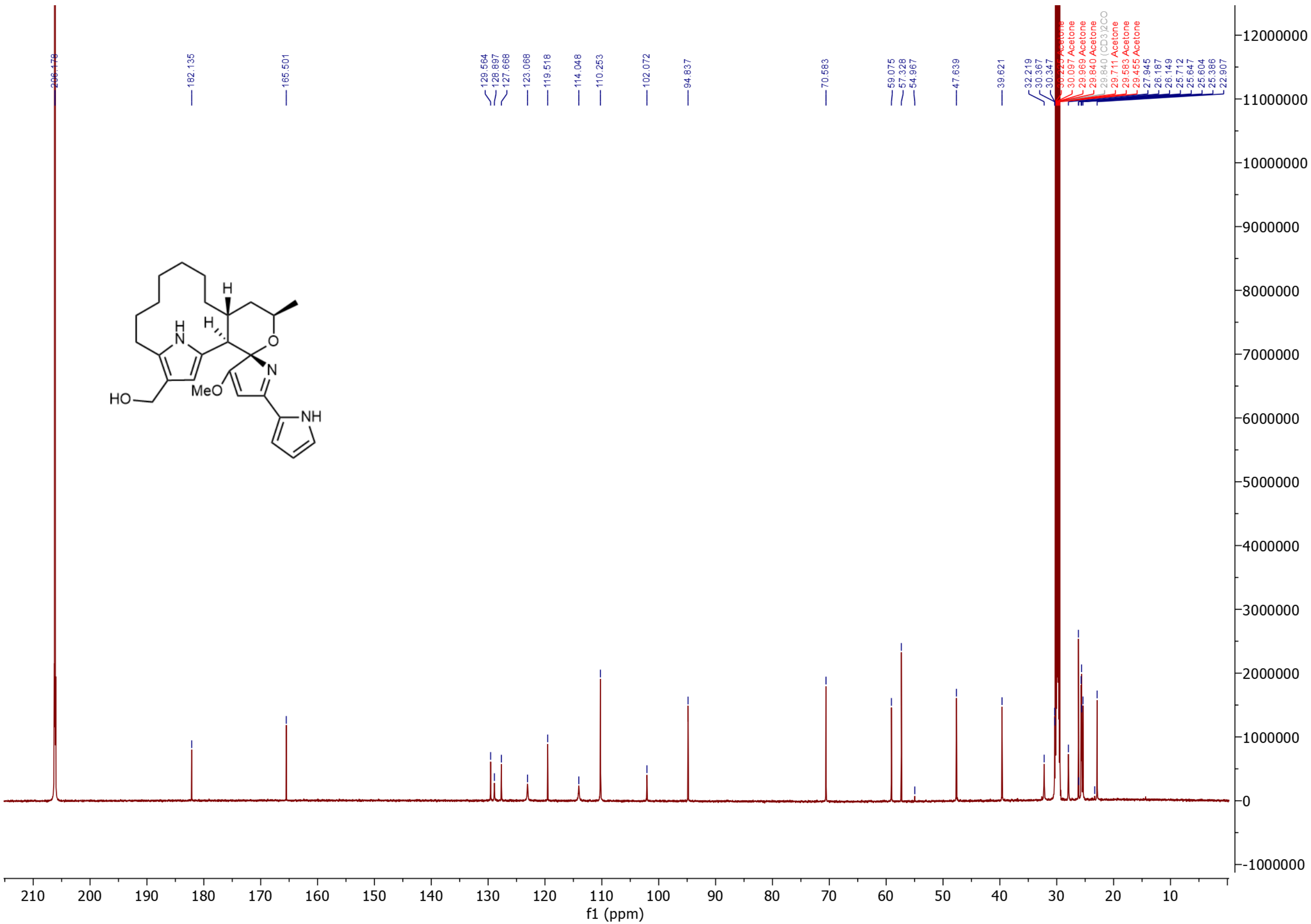

**Supplementary Figure 52.** $^{13}C$ NMR Spectrum of 12-hydroxymethyl premarineosin A (**9**) in Acetone-$D_6$.

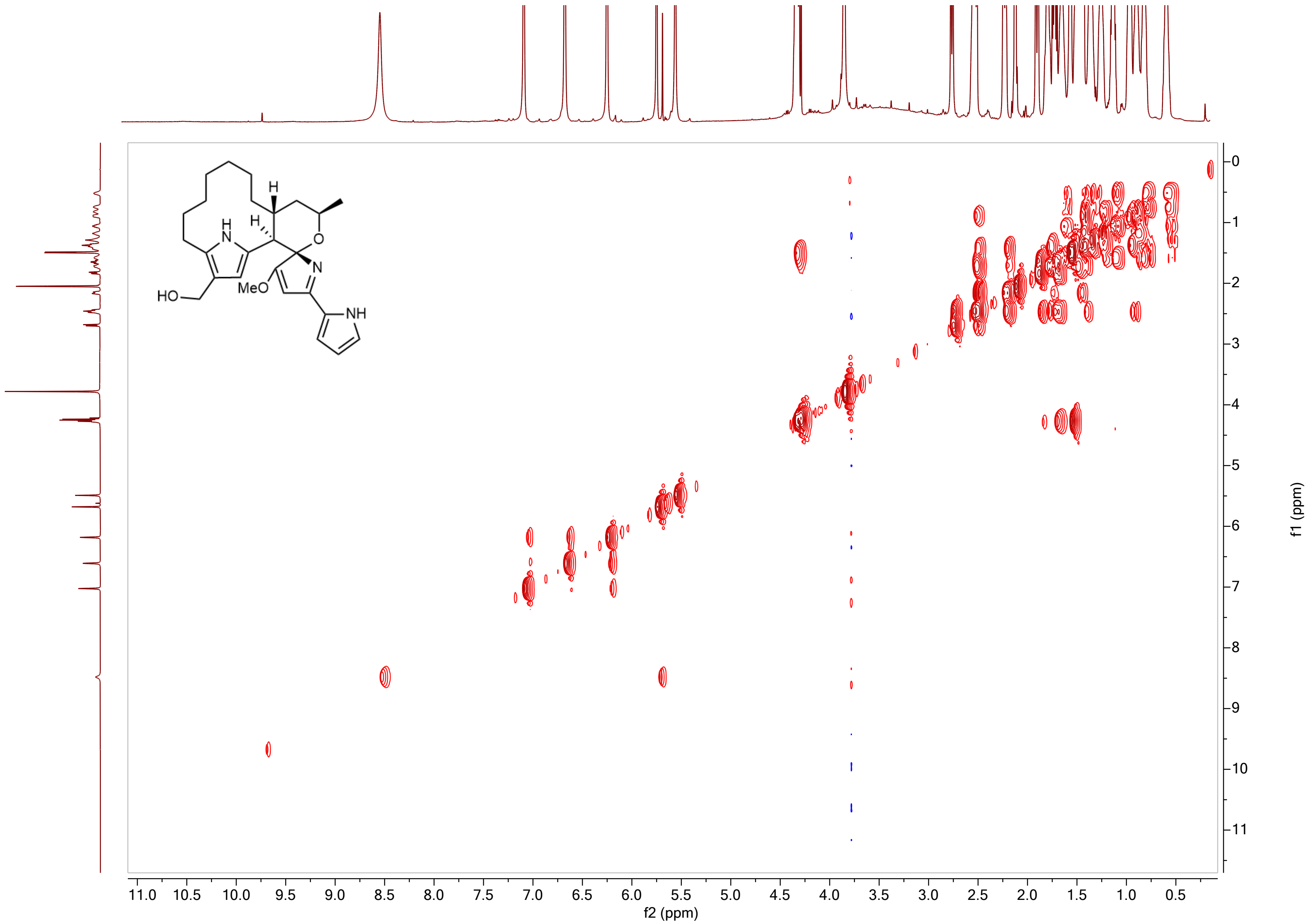

**Supplementary Figure 53.** $^1H$-$^1H$ COSY NMR Spectrum of 12-hydroxymethyl premarineosin A (**9**) in Acetone-$D_6$.

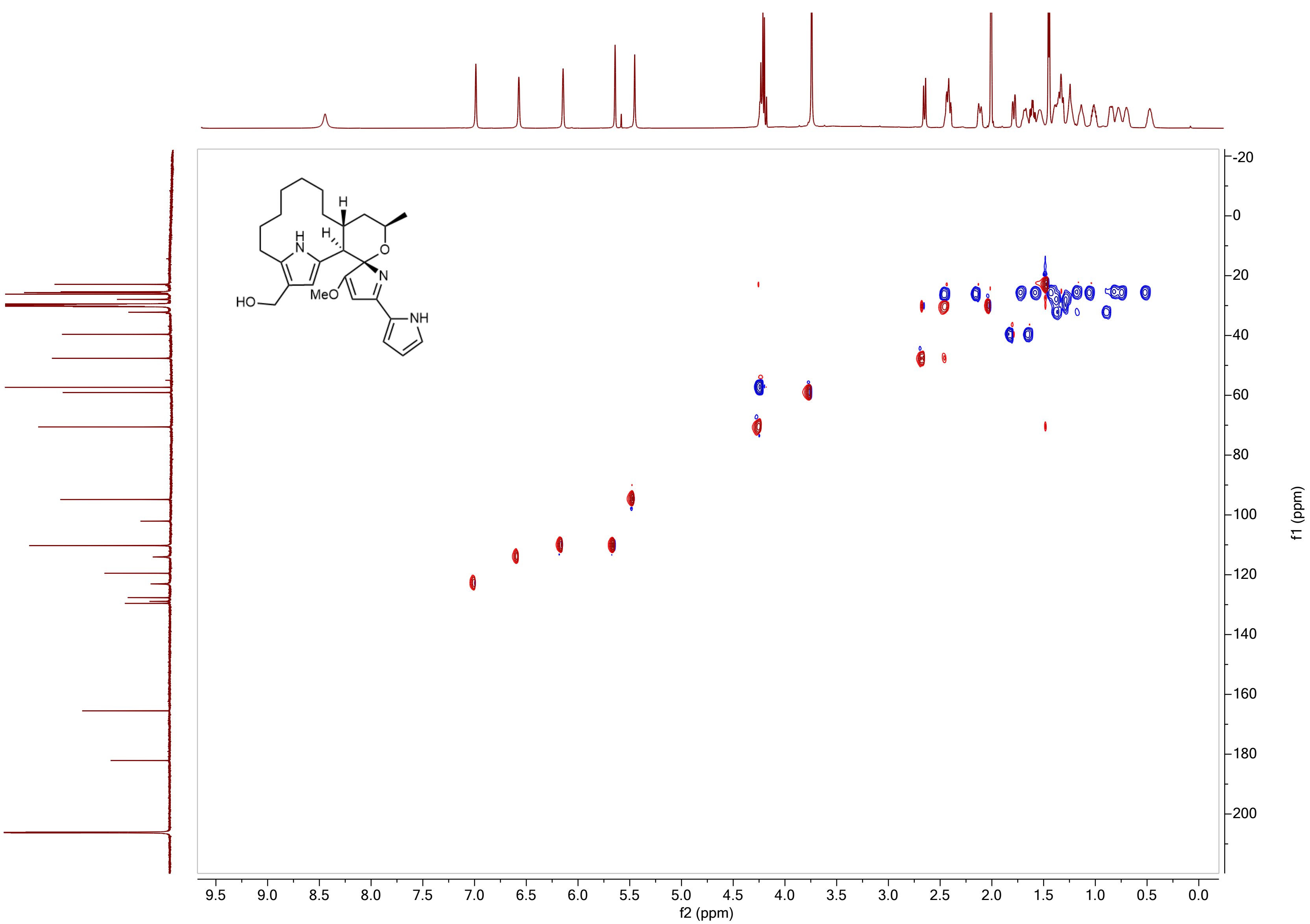

**Supplementary Figure 54.** $^{1}H$-$^{13}C$ HSQC NMR Spectrum of 12-hydroxymethyl premarineosin A (**9**) in Acetone-$D_6$.

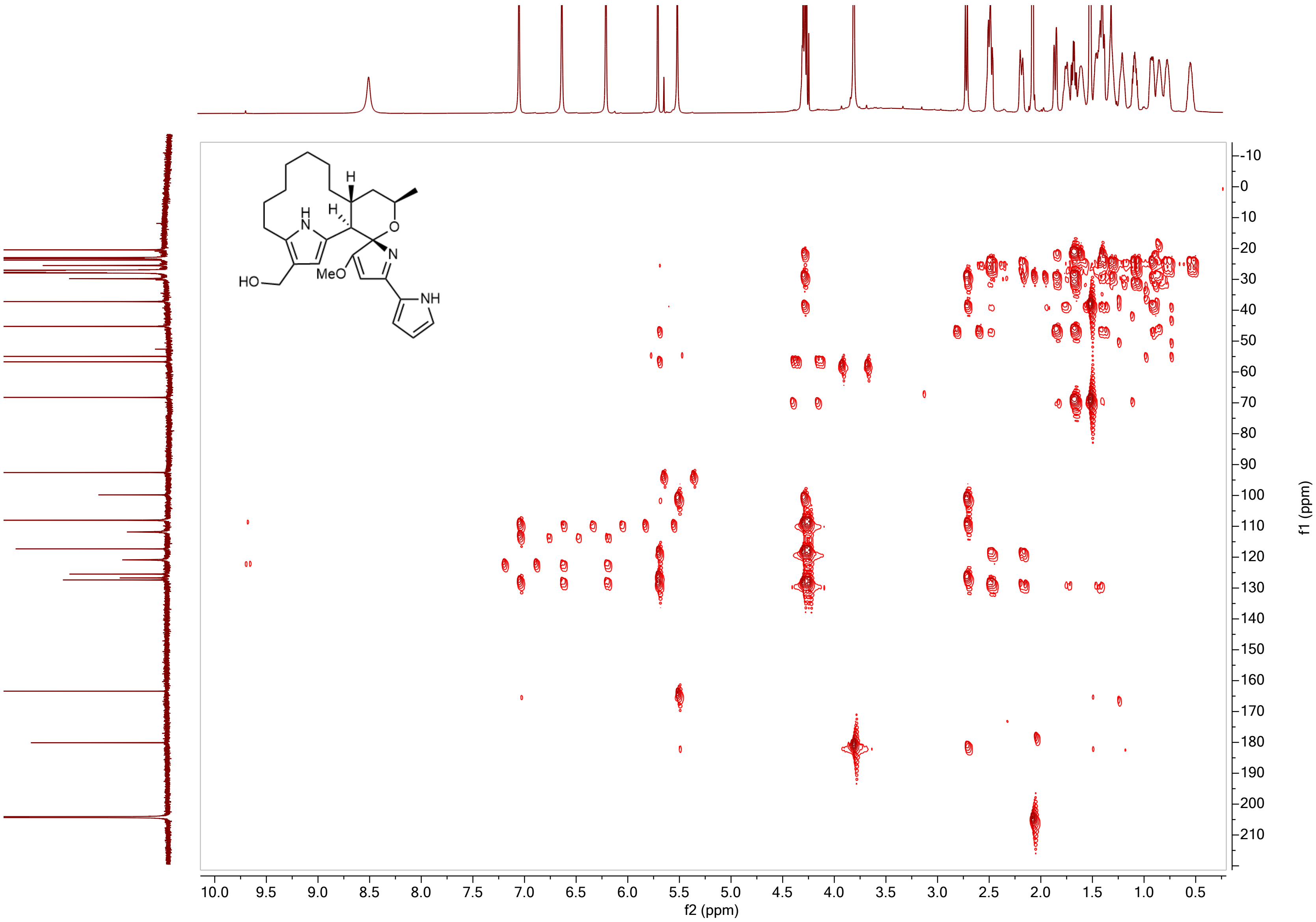

**Supplementary Figure 55.** $^{1}H$-$^{13}C$ HMBC NMR Spectrum of 12-hydroxymethyl premarineosin A (**9**) in Acetone-$D_6$.

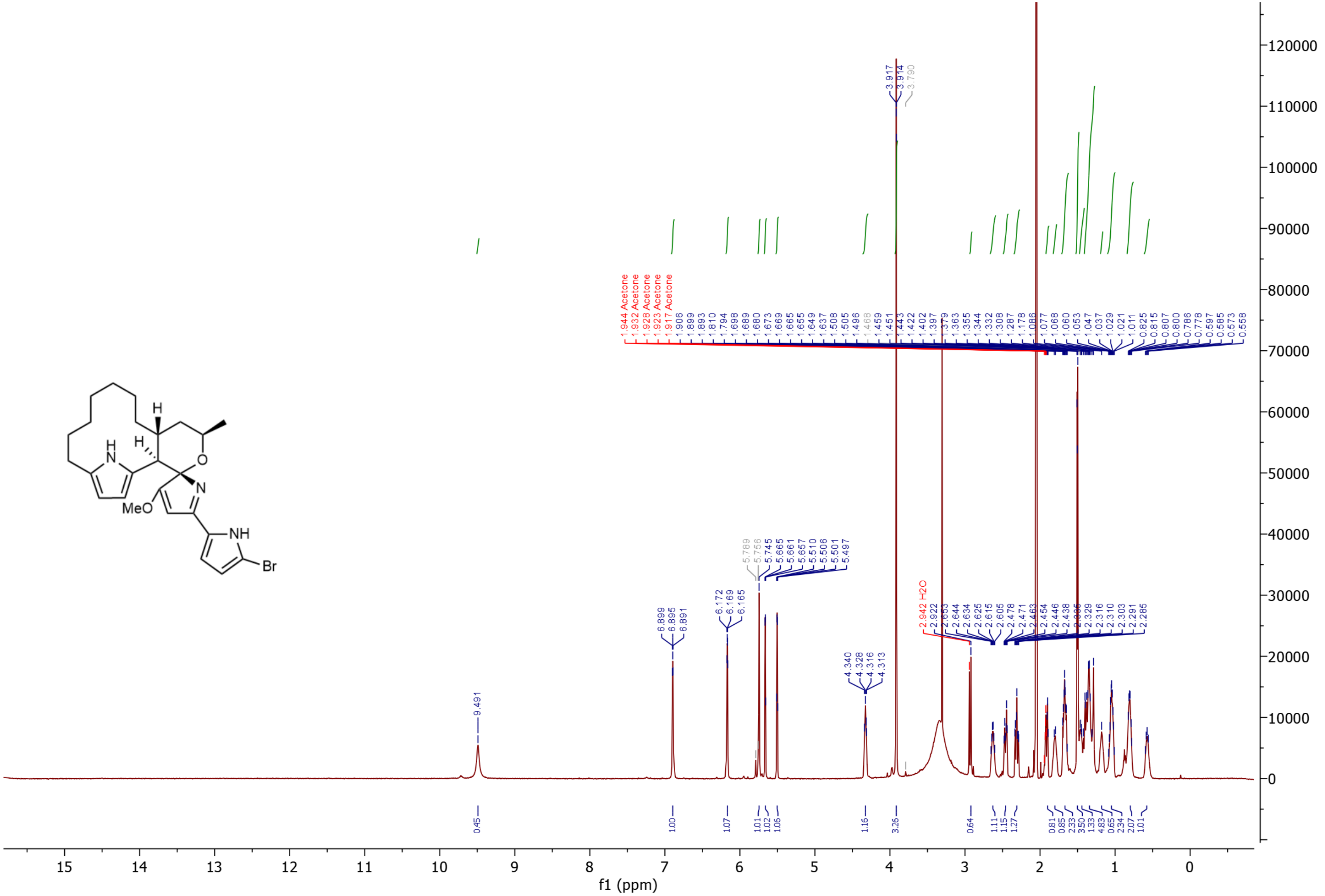

**Supplementary Figure 56.** $^{1}$H NMR Spectrum of 1-bromo-premarineosin (**10**) in Acetone-$D_6$.

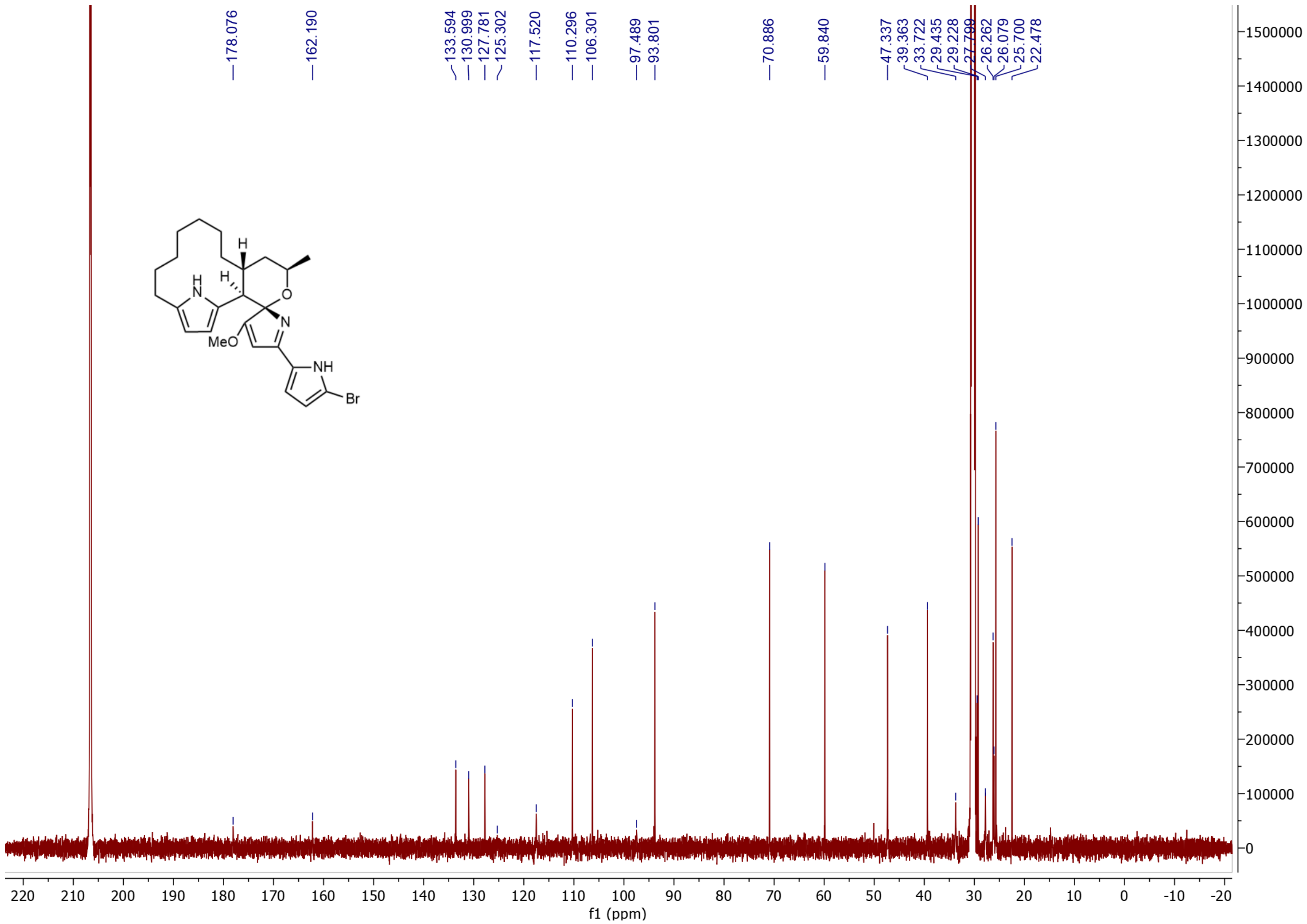

**Supplementary Figure 57.** $^{1}$C NMR Spectrum of 1-bromo-premarineosin (**10**) in Acetone-$D_6$.

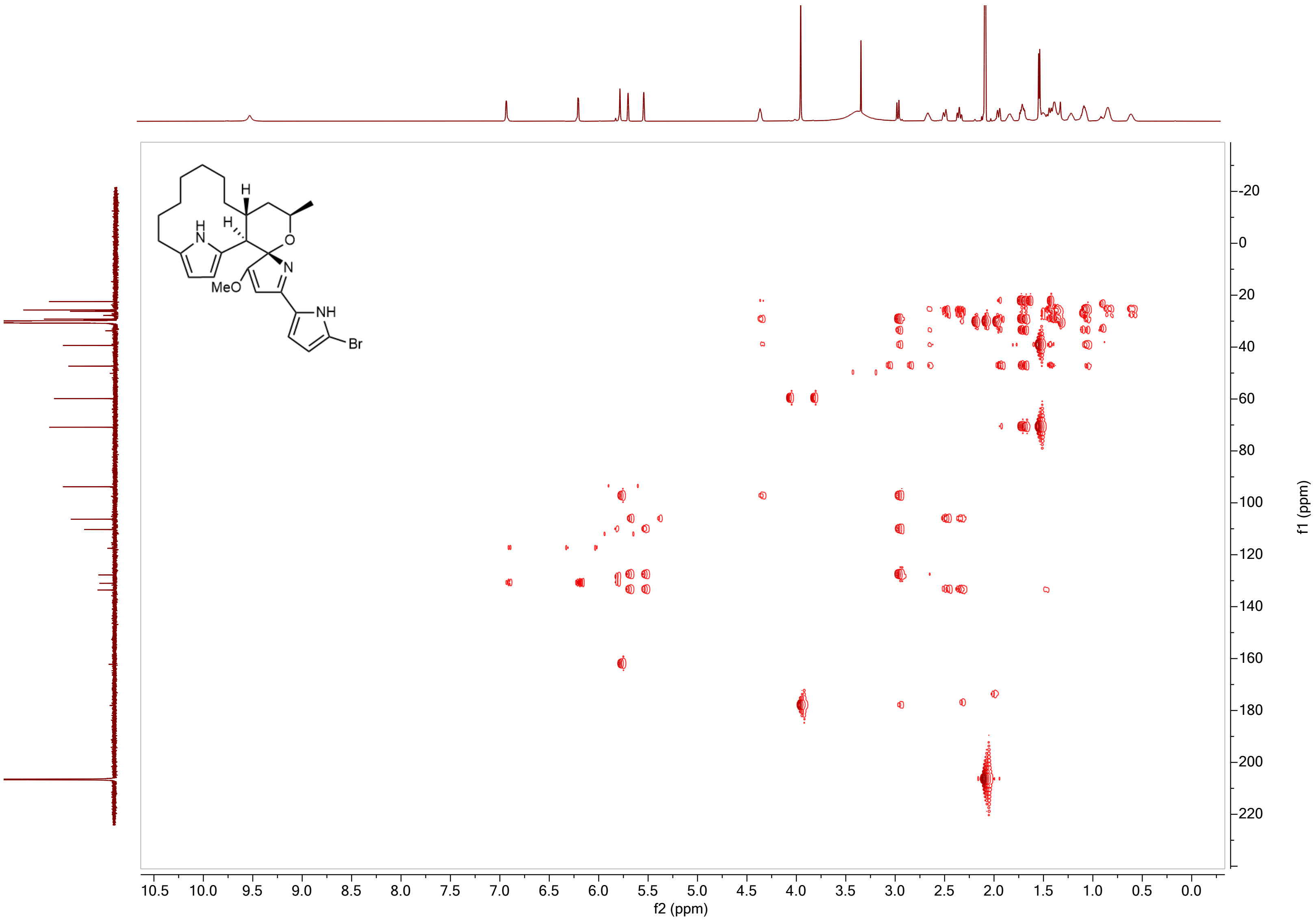

**Supplementary Figure 58.** $^{1}H$-$^{13}C$ HMBC NMR Spectrum of 1-bromo-premarineosin (**10**) in Acetone-$D_6$.

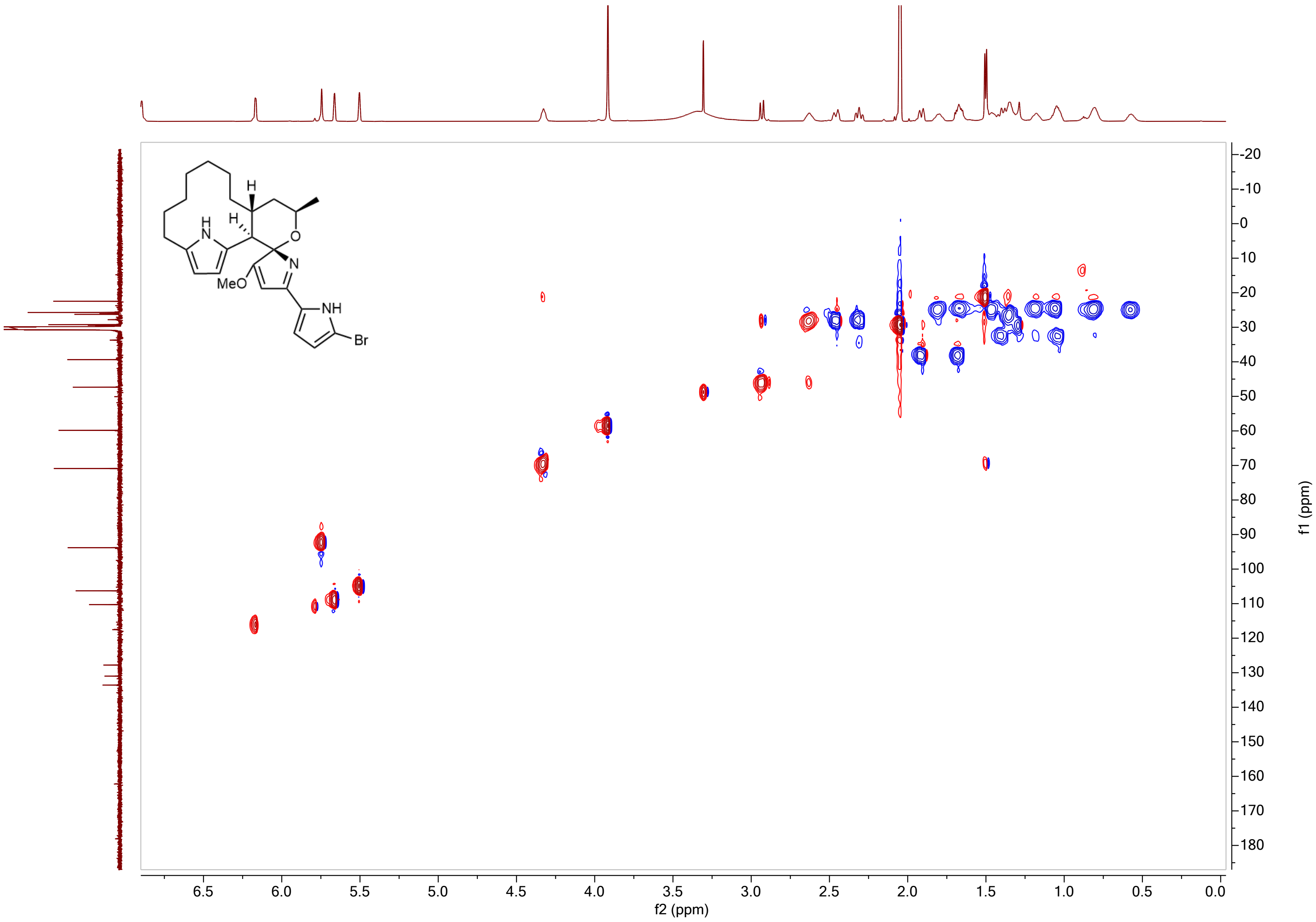

**Supplementary Figure 59.** $^{1}H$-$^{13}C$ HSQC NMR Spectrum of 1-bromo-premarineosin (**10**) in Acetone-$D_6$.

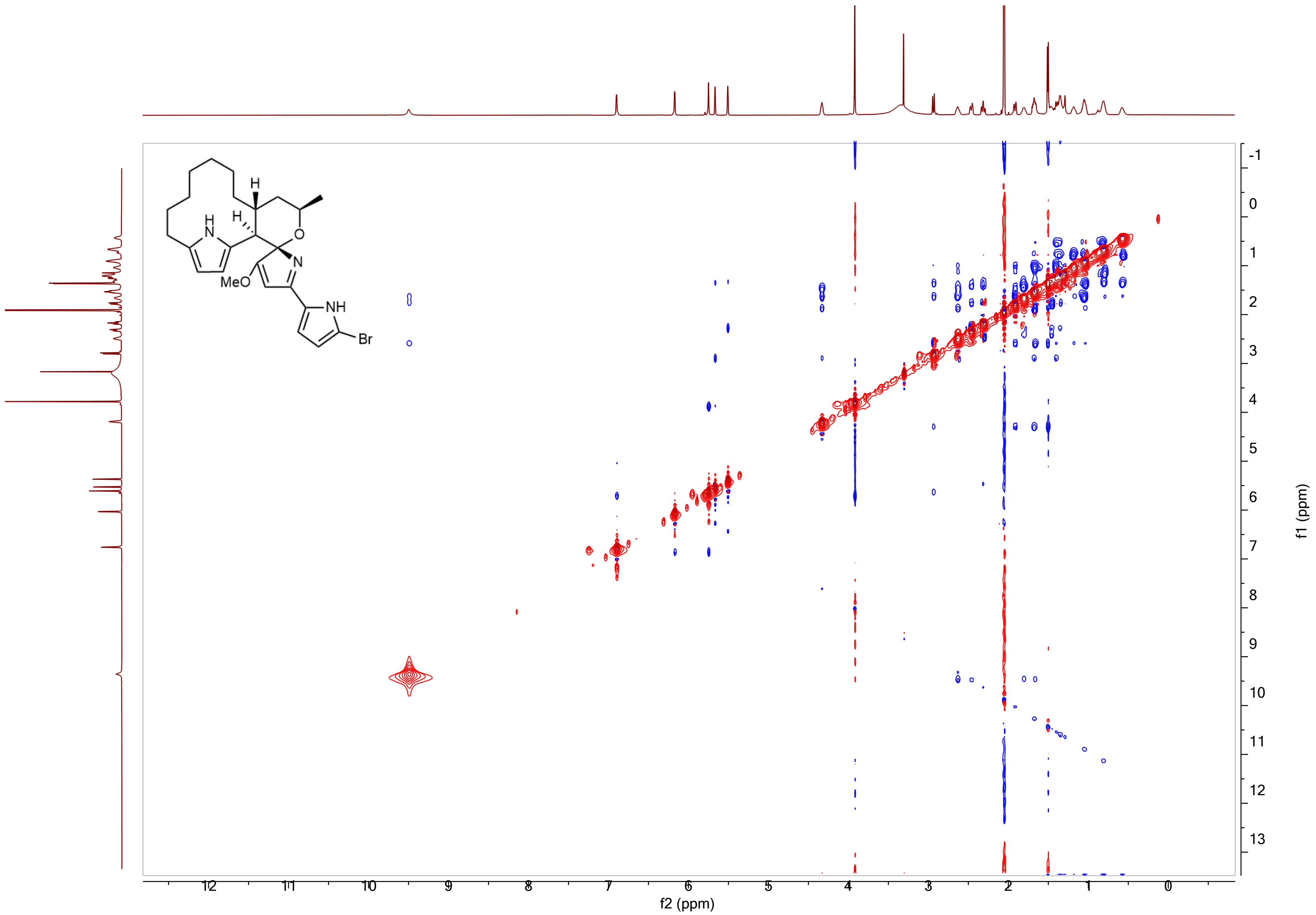

**Supplementary Figure 60.** $^{1}$H-$^{13}$C NOESY Spectrum of 1-bromo-premarineosin (**10**) in Acetone-D$_6$.

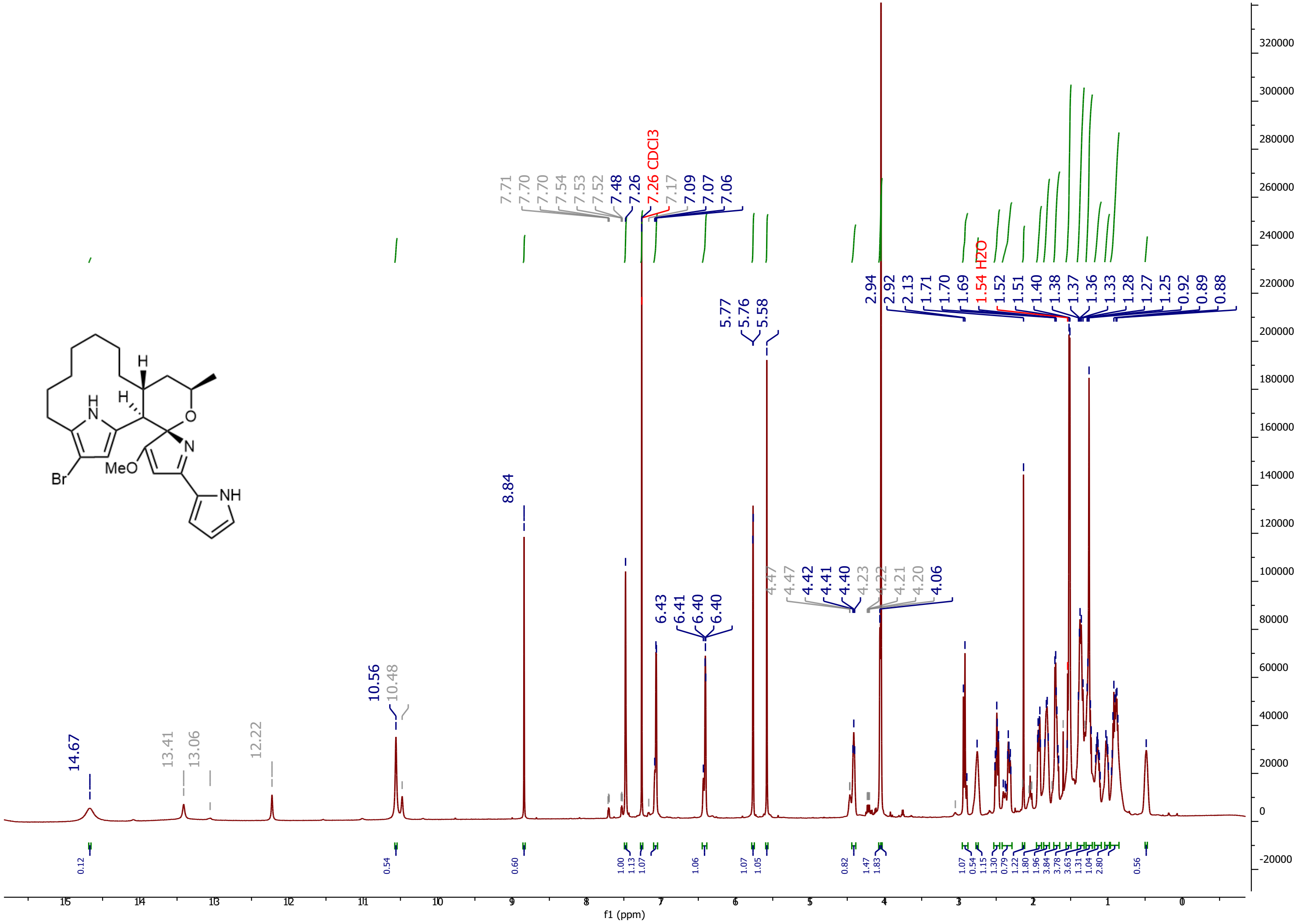

**Supplementary Figure 61.** $^1$H NMR Spectrum of 12-bromo-premarineosin (**11**) in chloroform-D.

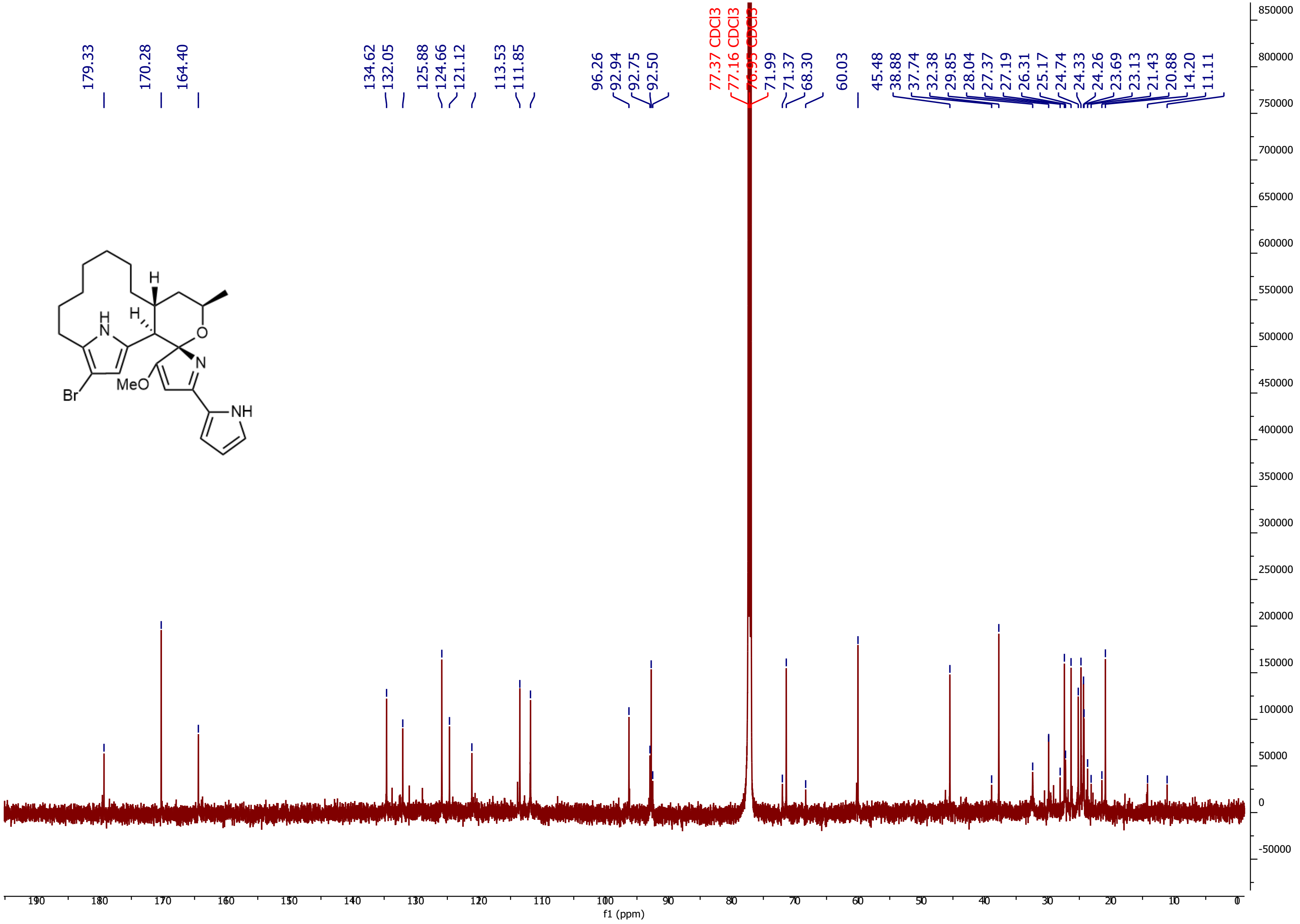

**Supplementary Figure 62.** 1C NMR Spectrum of 12-bromo-premarineosin (**11**) in chloroform-D.

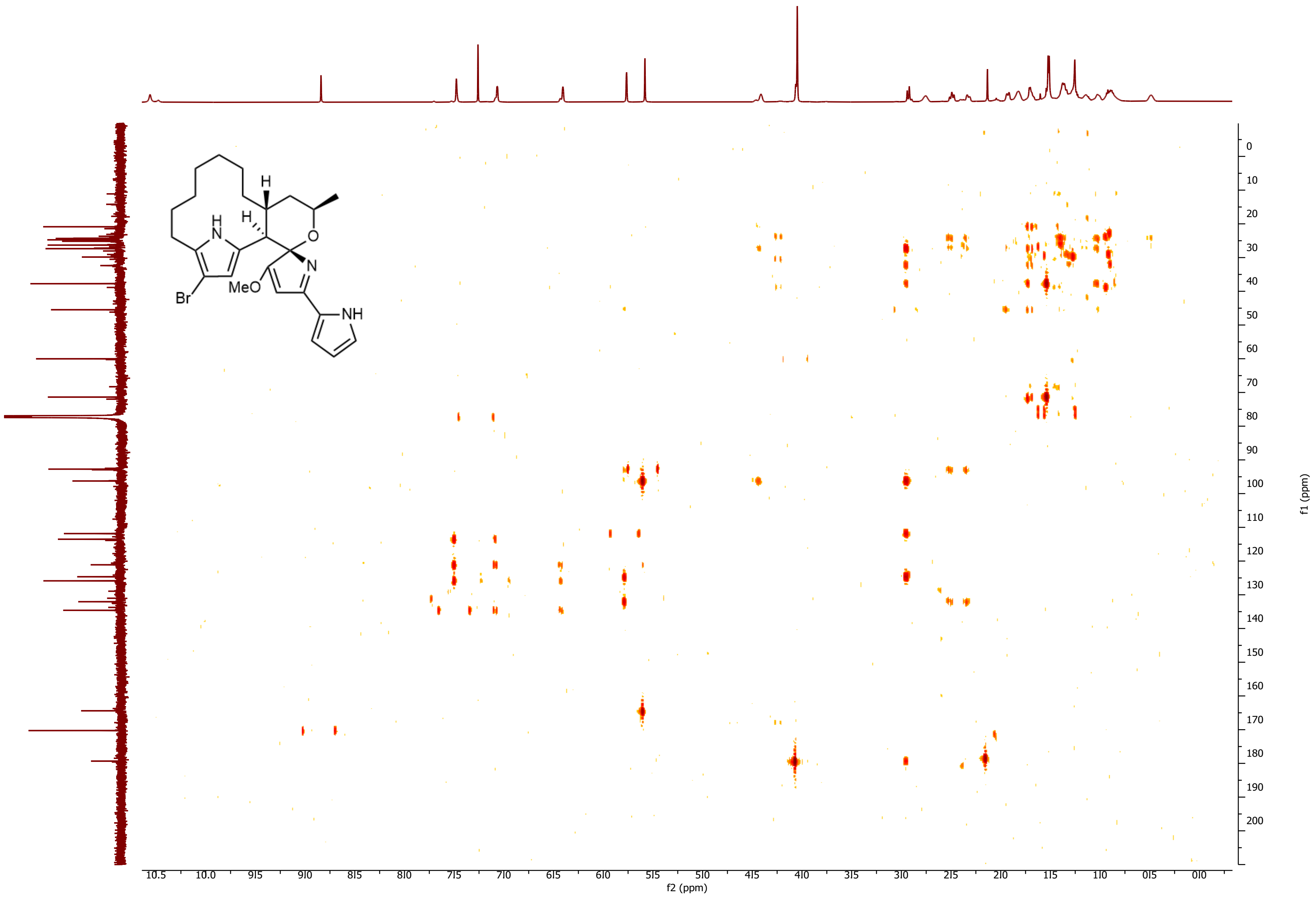

**Supplementary Figure 63.** $^{1}H$-$^{13}C$ HMBC Spectrum of 12-bromo-premarineosin (**11**) in chloroform-D

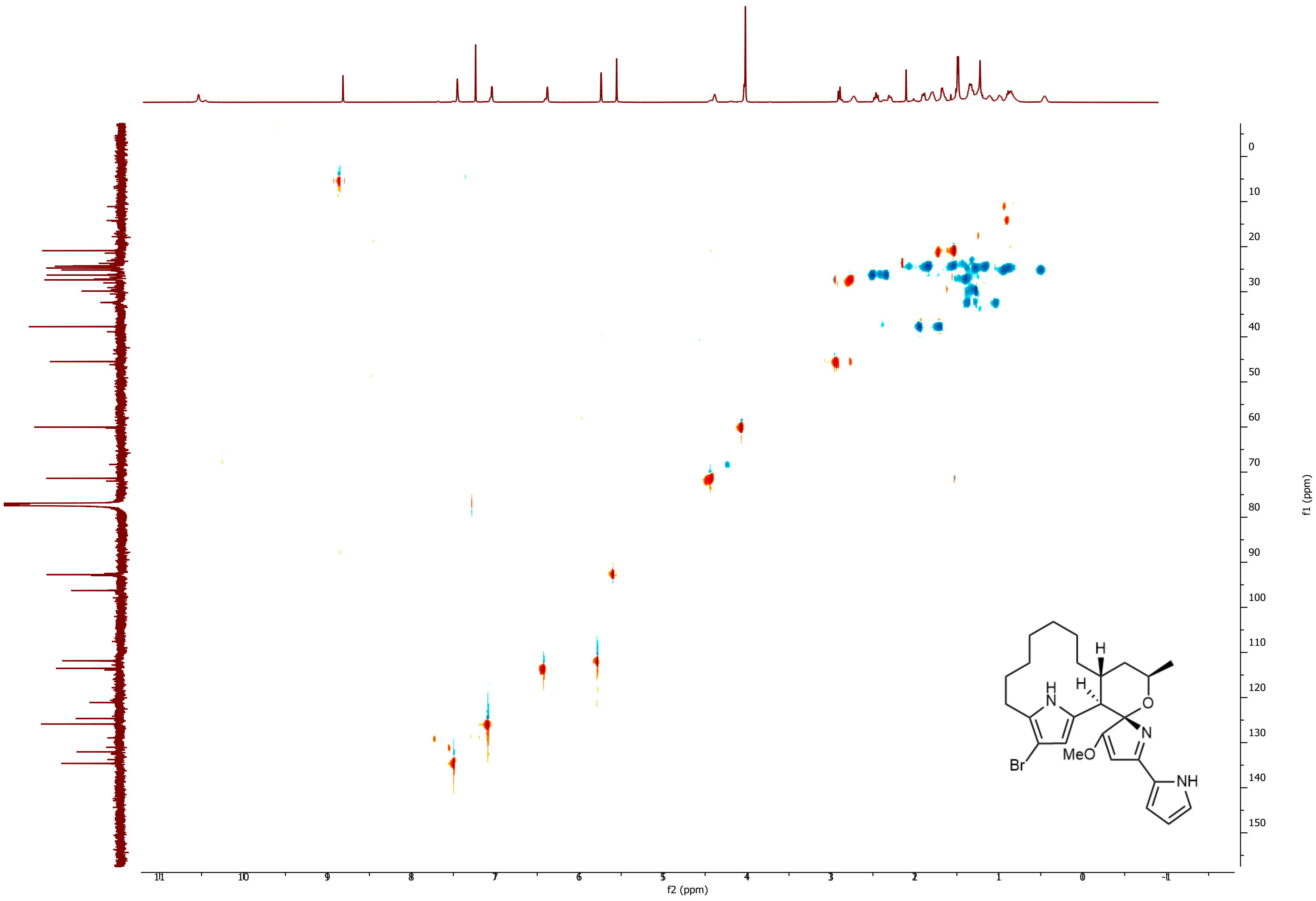

**Supplementary Figure 64.** $^1H$-$^{13}C$ HSQC NMR Spectrum of 12-bromo-premarineosin (**11**) in chloroform-D.

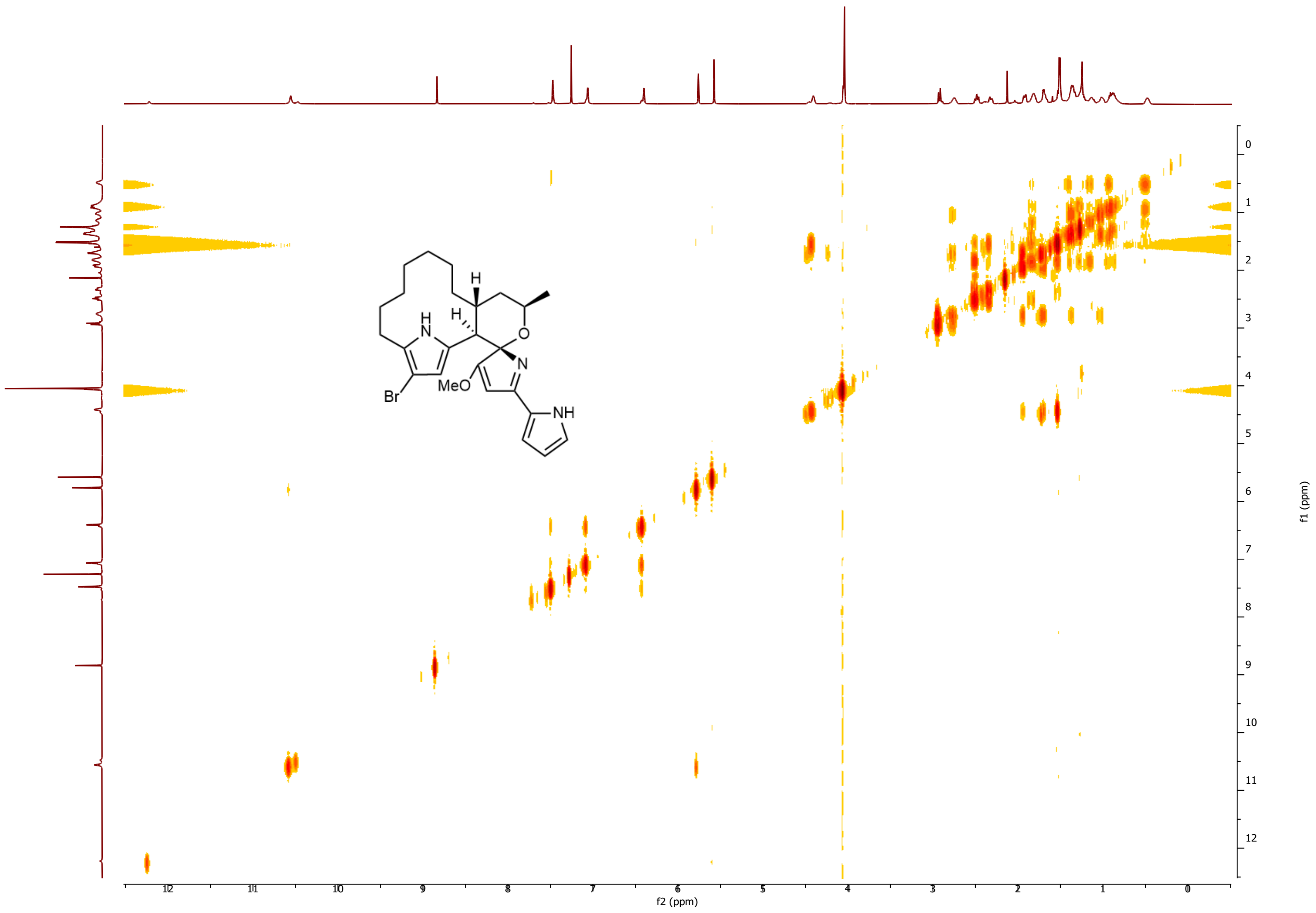

**Supplementary Figure 65.** $^{1}H$-$^{1}H$ COSY NMR Spectrum of 12-bromo-premarineosin (**11**) in chloroform-D.

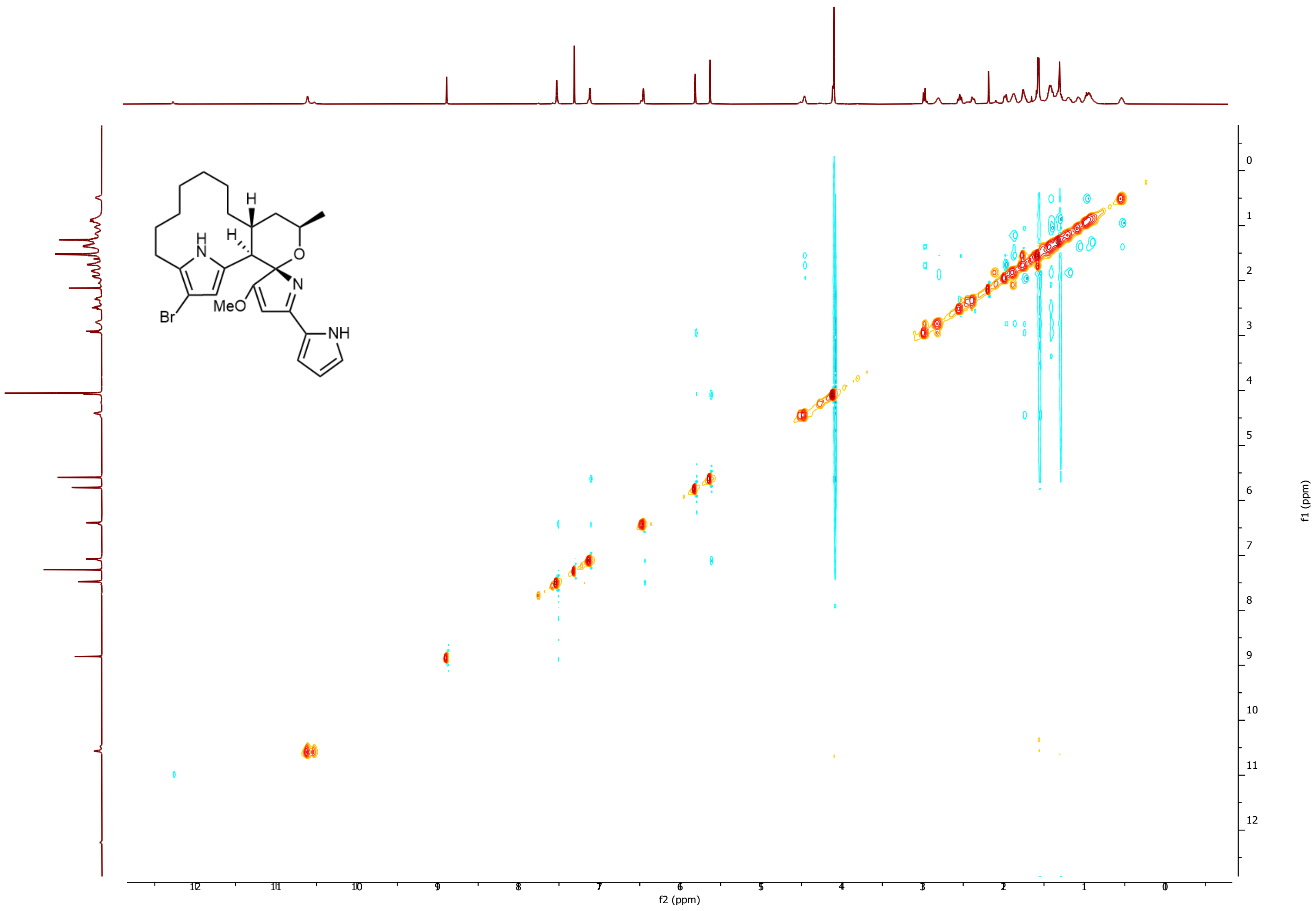

**Supplementary Figure 66.** $^1$H-$^{13}$C NOESY Spectrum of 12-bromo-premarineosin (**11**) in chloroform-D.

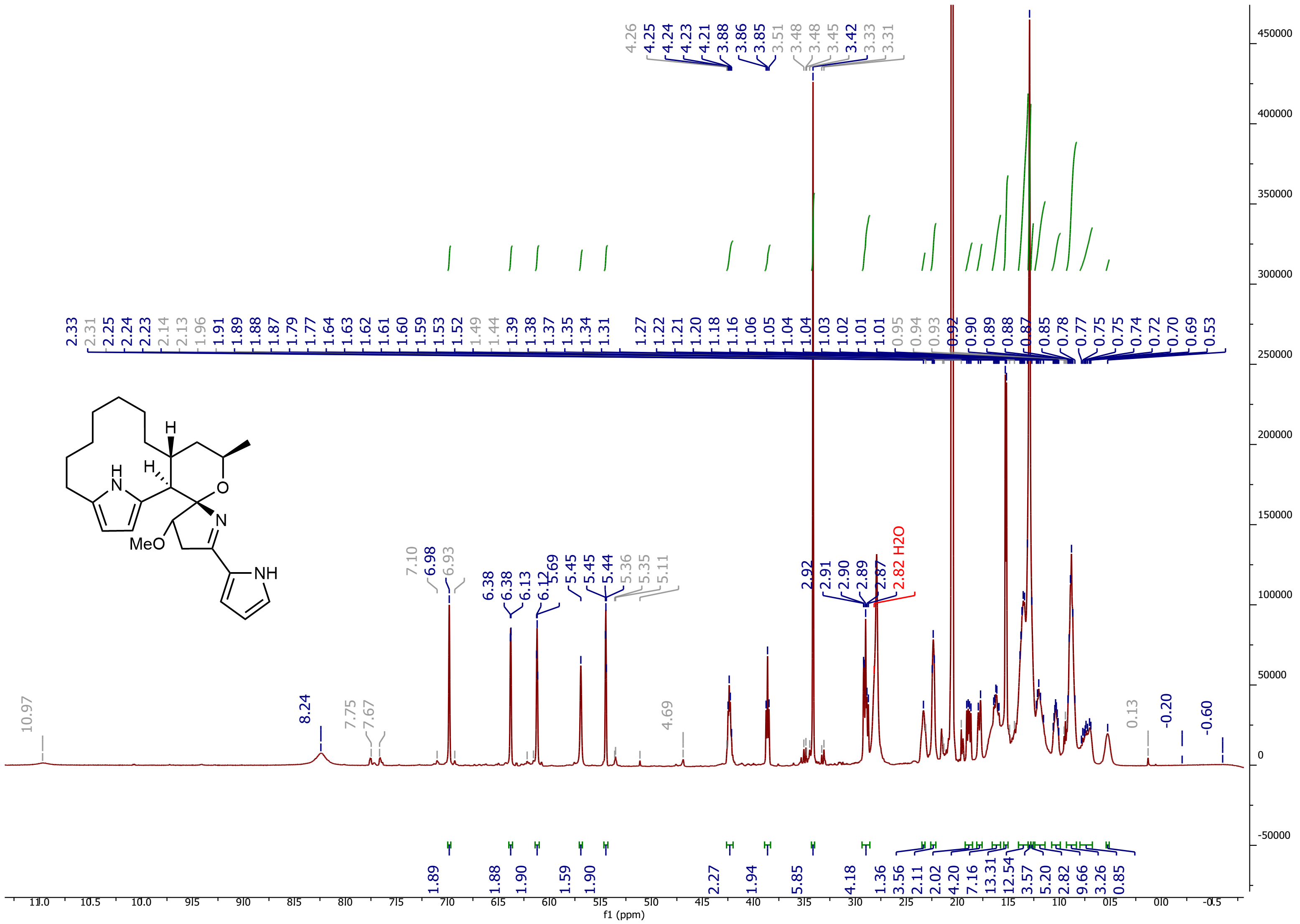

**Supplementary Figure 67.** $^{1}$H NMR Spectrum of marineosin A in Acetone-$D_6$.

### Supplementary Data.

#### Supplementary Data 1. X-Ray Crystal Structure Determination for (-)-premarineosin A.

**Experimental.** Single colourless prism-shaped crystals of **2** were used as supplied. A suitable crystal with dimensions 0.21 × 0.11 × 0.09 mm$^3$ was selected and mounted on a dtrek-CrysAlisPro-abstract goniometer imported rigaku-d*trek images diffractometer. The crystal was kept at a steady *T* = 85 K during data collection. The structure was solved with the ShelXT 2018/2[10] solution program using dual methods and by using Olex2 1.5-alpha[11] as the graphical interface. The model was refined with ShelXL 2019/3[12] using full matrix least squares minimisation on $\boldsymbol{F}^2$.

**Crystal Data.** $C_{25}H_{35}N_3O_3$, $M_r$ = 425.56, orthorhombic, $P2_12_12_1$ (No. 19), a = 8.14850(10) Å, b = 12.6238(2) Å, c = 21.7326(3) Å, *a* = *b* = *g* = 90˚, *V* = 2235.53(5) Å$^3$, *T* = 85 K, *Z* = 4, *Z'* = 1, *m*(Cu K$_a$) = 0.662, 33092 reflections measured, 4146 unique ($R_{int}$ = 0.0795) which were used in all calculations. The final $wR_2$ was 0.0971 (all data) and $R_1$ was 0.0373 (I≥2 *s*(I)).

| | |
|---|---|
| Formula | $C_{25}H_{35}N_3O_3$ |
| $D_{calc.}$/ g cm$^{-3}$ | 1.264 |
| *m*/mm$^{-1}$ | 0.662 |
| Formula Weight | 425.56 |
| Colour | colourless |
| Shape | prism-shaped |
| Size/mm$^3$ | 0.21×0.11×0.09 |
| *T*/K | 85 |
| Crystal System | orthorhombic |
| Flack Parameter | 0.09(13) |
| Hooft Parameter | 0.13(10) |
| Space Group | $P2_12_12_1$ |
| *a*/Å | 8.14850(10) |
| *b*/Å | 12.6238(2) |
| *c*/Å | 21.7326(3) |
| *a*/˚ | 90 |
| *b*/˚ | 90 |
| *g*/˚ | 90 |
| V/Å$^3$ | 2235.53(5) |
| *Z* | 4 |
| *Z'* | 1 |
| Wavelength/Å | 1.54184 |
| Radiation type | Cu K$_a$ |
| $Q_{min}$/˚ | 4.050 |
| $Q_{max}$/˚ | 69.810 |
| Measured Refl's. | 33092 |
| Indep't Refl's | 4146 |
| Refl's I≥2 *s*(I) | 3937 |
| $R_{int}$ | 0.0795 |
| Parameters | 318 |
| Restraints | 12 |
| Largest Peak | 0.204 |
| Deepest Hole | -0.193 |
| GooF | 1.088 |
| $wR_2$ (all data) | 0.0971 |
| $wR_2$ | 0.0888 |
| $R_1$ (all data) | 0.0418 |
| $R_1$ | 0.0373 |

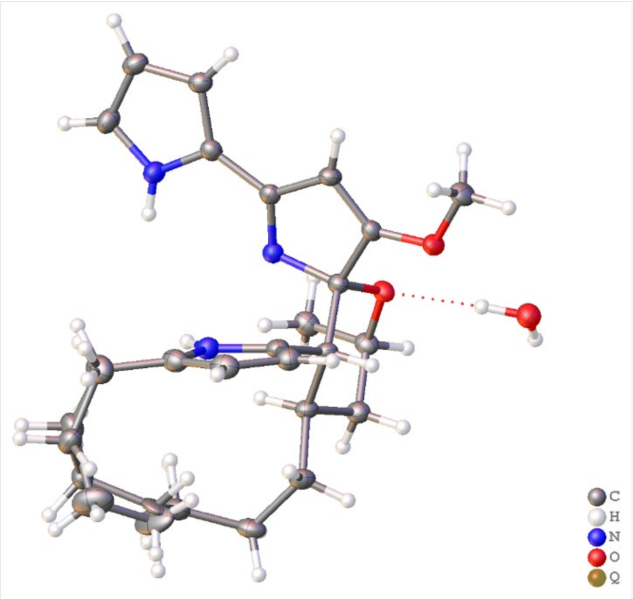

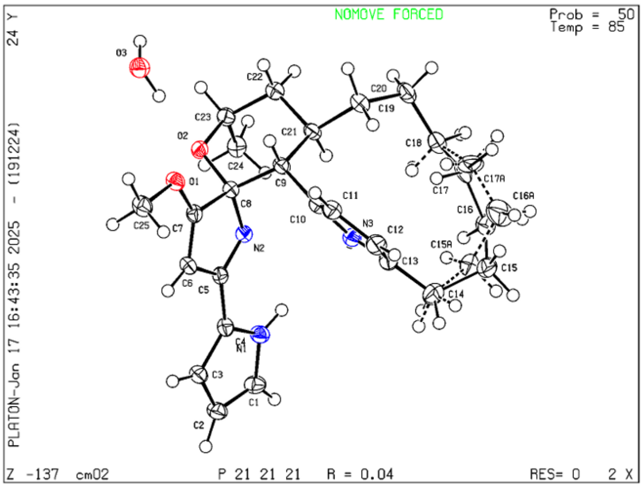

##### Structure Quality Indicators

| | | | | | |
|---|---|---|---|---|---|
| **Reflections:** | d min (CuKα) $2\Theta=139.6°$ 0.82 | I/σ(I) 27.8 | Rint m=8.01 7.95% | Full 135.4° 99% to 139.6° 100 | |
| **Refinement:** | Shift -0.001 | Max Peak 0.2 | Min Peak -0.2 | GooF 1.088 | Hooft .13(10) |

A colorless prism-shaped-shaped crystal with dimensions 0.21 × 0.11 × 0.09 $mm^3$ was mounted. Data were collected using a dtrek-CrysAlisPro-abstract goniometer imported rigaku-d*trek images diffractometer operating at $T$ = 85 K.

Data were measured using $w$ scans with Cu $K_a$ radiation. The diffraction pattern was indexed and the total number of runs and images was based on the strategy calculation from the program DTREK_VERSION=d*TREK version 9.9.9.4 W9RSSI – Aug 22 2012. The maximum resolution that was achieved was $Q$ = 69.810° (0.82 Å).

The unit cell was refined using CrysAlisPro 1.171.43.130a[13] on 18942 reflections, 57% of the observed reflections.

Data reduction, scaling and absorption corrections were performed using CrysAlisPro 1.171.43.130a[13]. The final completeness is 100.00 % out to 69.810° in Q. A multi-scan absorption correction was performed using CrysAlisPro 1.171.43.130a[13]. Empirical absorption correction using spherical harmonics, implemented in SCALE3 ABSPACK scaling algorithm.. The absorption coefficient $m$ of this material is 0.662 $mm^{-1}$ at this wavelength ($l$ = 1.54184Å) and the minimum and maximum transmissions are 0.814 and 1.000.

The structure was solved and the space group $P2_12_12_1$ (# 19) determined by the ShelXT 2018/2[10] structure solution program using using dual methods and refined by full matrix least squares minimization on $\boldsymbol{F}^2$ using version 2019/3 of ShelXL 2019/3[12]. All non-hydrogen atoms were refined anisotropically. Hydrogen atom positions were calculated geometrically and refined using the riding model. Most hydrogen atom positions were calculated geometrically and refined using the riding model, but some hydrogen atoms were refined freely.

*_exptl_absorpt_process_details*: CrysAlisPro 1.171.43.130a[13] using spherical harmonics, implemented in SCALE3 ABSPACK scaling algorithm.

There is a single formula unit in the asymmetric unit, which is represented by the reported sum formula. In other words: Z is 4 and Z' is 1. The moiety formula is C25 H33 N3 O2, H2 O.

The Flack parameter was refined to 0.09(13). Determination of absolute structure using Bayesian statistics on Bijvoet differences using the Olex2 results in 0.13(10). The chiral atoms in this structure are: C8(S), C9(R), C21(R), C23(R). Note: The Flack parameter is used to determine chirality of the crystal studied, the value should be near 0, a value of 1 means that the stereochemistry is wrong and the model should be inverted. A value of 0.5 means that the crystal consists of a racemic mixture of the two enantiomers.

Fractional Atomic Coordinates (×$10^4$) and Equivalent Isotropic Displacement Parameters (Å$^2$×$10^3$) for **2**. $U_{eq}$ is defined as 1/3 of the trace of the orthogonalised $U_{ij}$.

| Atom | x | y | z | $U_{eq}$ |
|---|---|---|---|---|
| O1 | 7086(2) | 6707.0(14) | 4084.1(8) | 23.2(4) |
| O2 | 5883(2) | 5983.1(13) | 2941.7(8) | 20.6(4) |
| O3 | 9377(3) | 5605.3(15) | 3034.9(9) | 28.6(4) |
| N1 | 364(3) | 7831.4(17) | 3170.5(10) | 23.4(5) |
| N2 | 3222(3) | 6581.2(16) | 3272.2(9) | 19.2(4) |
| N3 | 1647(3) | 5099.1(17) | 4234.5(10) | 22.0(5) |
| C1 | -745(3) | 8630(2) | 3131.9(12) | 26.5(6) |
| C2 | -99(3) | 9511(2) | 3418.6(12) | 25.9(6) |
| C3 | 1470(3) | 9223(2) | 3642.3(12) | 22.8(5) |
| C4 | 1732(3) | 8176(2) | 3478.5(11) | 20.4(5) |
| C5 | 3151(3) | 7493.0(19) | 3548.7(11) | 19.1(5) |
| C6 | 4598(3) | 7767.2(19) | 3920.4(11) | 20.3(5) |
| C7 | 5599(3) | 6930.4(19) | 3861.4(11) | 19.3(5) |
| C8 | 4792(3) | 6104.9(19) | 3453.3(11) | 19.1(5) |
| C9 | 4516(3) | 5044.9(19) | 3791.7(11) | 20.0(5) |
| C10 | 3327(3) | 5099.9(19) | 4318.9(12) | 21.4(5) |
| C11 | 3605(3) | 4934(2) | 4933.4(12) | 24.5(6) |
| C12 | 2053(4) | 4815(2) | 5224.1(12) | 28.2(6) |
| C13 | 861(4) | 4910(2) | 4786.5(12) | 26.3(6) |
| C14 | -953(4) | 4776(2) | 4830.7(14) | 35.2(7) |
| C15 | -1675(6) | 3660(3) | 4799(2) | 30.3(13) |
| C15A | -1627(9) | 3888(5) | 4417(4) | 31(2) |
| C16 | -1368(6) | 3094(4) | 4183(2) | 31.1(14) |
| C16A | -1119(15) | 2716(6) | 4448(5) | 50(3) |
| C17 | 380(8) | 2633(9) | 4142(7) | 35(2) |
| C17A | 370(20) | 2421(19) | 4142(13) | 55(5) |
| C18 | 898(4) | 2604(2) | 3475.3(14) | 32.1(7) |
| C19 | 2687(4) | 2303(2) | 3372.3(13) | 27.5(6) |
| C20 | 3924(3) | 3071(2) | 3661.4(12) | 24.9(6) |
| C21 | 4030(3) | 4155.1(19) | 3336.7(12) | 20.6(5) |
| C22 | 5272(3) | 4107(2) | 2808.6(12) | 23.9(6) |
| C23 | 5477(3) | 5171.4(19) | 2488.9(11) | 22.2(5) |
| C24 | 4038(3) | 5481(2) | 2082.0(12) | 25.8(6) |
| C25 | 7782(3) | 7494(2) | 4488.7(13) | 27.0(6) |

Anisotropic Displacement Parameters (×$10^4$) for **2**. The anisotropic displacement factor exponent takes the form: $-2p^2[h^2a^{*2} \times U_{11} + ... +2hka^* \times b^* \times U_{12}]$

| Atom | $U_{11}$ | $U_{22}$ | $U_{33}$ | $U_{23}$ | $U_{13}$ | $U_{12}$ |
|---|---|---|---|---|---|---|
| O1 | 20.0(9) | 19.4(8) | 30.3(9) | -2.9(7) | -5.9(8) | -0.7(7) |
| O2 | 20.6(9) | 19.4(9) | 21.8(8) | -3.5(7) | 2.3(7) | -3.0(7) |
| O3 | 25.0(10) | 25.7(10) | 35.0(10) | -3.3(8) | -1.1(9) | -1.0(8) |
| N1 | 21.3(11) | 20.5(10) | 28.5(11) | -1.3(9) | -2.5(9) | 2.2(9) |
| N2 | 18.4(10) | 17.6(10) | 21.6(10) | 0.7(8) | 0.2(8) | 0.0(8) |
| N3 | 22.4(11) | 21.0(10) | 22.6(10) | 0.6(9) | 0.6(9) | 1.0(9) |
| C1 | 22.8(13) | 26.8(14) | 29.8(14) | 0.6(11) | -2.4(11) | 5.9(11) |
| C2 | 28.5(14) | 20.1(12) | 28.9(13) | 1.7(11) | 2.3(11) | 5.0(11) |
| C3 | 25.5(14) | 18.1(12) | 24.8(12) | 1.0(10) | 1.1(11) | 0.6(10) |
| C4 | 21.2(12) | 19.0(12) | 21.1(12) | 2.3(10) | -0.5(10) | -1.9(10) |
| C5 | 19.0(12) | 17.5(12) | 20.8(12) | 2.6(9) | 1.3(10) | -1.4(10) |
| C6 | 22.9(13) | 15.2(11) | 22.7(12) | -0.6(10) | -1.1(10) | -1.1(10) |
| C7 | 18.2(12) | 18.5(12) | 21.3(11) | 0.7(10) | -1.3(10) | -2.6(10) |
| C8 | 19.1(12) | 16.8(11) | 21.5(12) | -1.1(9) | 1.6(10) | -0.8(9) |
| C9 | 22.0(13) | 15.1(11) | 23.0(12) | 1.1(10) | -0.8(10) | 0.7(10) |
| C10 | 22.8(13) | 14.7(11) | 26.6(12) | -1.5(10) | 0.8(11) | 1.1(10) |

| Atom | $U_{11}$ | $U_{22}$ | $U_{33}$ | $U_{23}$ | $U_{13}$ | $U_{12}$ |
|---|---|---|---|---|---|---|
| C11 | 26.6(14) | 22.2(13) | 24.6(13) | -1.6(11) | -0.1(10) | 0.2(11) |
| C12 | 35.9(16) | 25.8(14) | 23.0(12) | -1.5(11) | 6.3(11) | -0.8(12) |
| C13 | 29.1(14) | 21.9(13) | 27.8(13) | -2.3(11) | 7.4(11) | -1.4(11) |
| C14 | 32.4(16) | 35.4(17) | 37.7(16) | -5.5(13) | 11.2(13) | -3.2(13) |
| C15 | 25(2) | 28(2) | 38(3) | 7(2) | 5(2) | 3(2) |
| C15A | 23(4) | 31(5) | 39(5) | -1(4) | 5(4) | -5(3) |
| C16 | 21(2) | 25(3) | 47(3) | -3(3) | -5(2) | -3(2) |
| C16A | 73(8) | 37(6) | 40(6) | 11(4) | 15(5) | 4(5) |
| C17 | 37(4) | 15(4) | 54(4) | -12(3) | 1(3) | -13(3) |
| C17A | 66(8) | 33(9) | 67(7) | 0(7) | 47(7) | 13(6) |
| C18 | 32.7(16) | 20.2(13) | 43.3(17) | 0.6(12) | 1.3(13) | -4.1(12) |
| C19 | 37.8(16) | 17.1(12) | 27.7(13) | -2.9(11) | 3.1(12) | -4.5(11) |
| C20 | 29.5(14) | 18.3(12) | 27.1(13) | 0.9(10) | -1.5(11) | 0.7(11) |
| C21 | 23.2(13) | 15.1(12) | 23.5(12) | -1.7(10) | 0.0(11) | 0.6(10) |
| C22 | 25.4(14) | 20.3(12) | 26.0(13) | -1.4(10) | 2.6(11) | 0.0(10) |
| C23 | 24.7(14) | 21.1(12) | 20.7(11) | -5.1(10) | 2.2(10) | -2.5(11) |
| C24 | 28.5(14) | 26.6(13) | 22.3(12) | -0.5(11) | 0.2(11) | -4.3(11) |
| C25 | 26.9(15) | 24.4(13) | 29.6(14) | -5.1(11) | -7.0(11) | -3.9(11) |

Bond Lengths in Å for **2.**

| Atom | Atom | Length/Å |
|---|---|---|
| O1 | C7 | 1.335(3) |
| O1 | C25 | 1.443(3) |
| O2 | C8 | 1.432(3) |
| O2 | C23 | 1.459(3) |
| N1 | C1 | 1.357(3) |
| N1 | C4 | 1.372(3) |
| N2 | C5 | 1.300(3) |
| N2 | C8 | 1.467(3) |
| N3 | C10 | 1.382(3) |
| N3 | C13 | 1.381(3) |
| C1 | C2 | 1.379(4) |
| C2 | C3 | 1.416(4) |
| C3 | C4 | 1.385(4) |
| C4 | C5 | 1.450(3) |
| C5 | C6 | 1.471(3) |
| C6 | C7 | 1.341(4) |
| C7 | C8 | 1.518(3) |
| C8 | C9 | 1.543(3) |
| C9 | C10 | 1.502(3) |

| Atom | Atom | Length/Å |
|---|---|---|
| C9 | C21 | 1.548(3) |
| C10 | C11 | 1.371(4) |
| C11 | C12 | 1.421(4) |
| C12 | C13 | 1.365(4) |
| C13 | C14 | 1.491(4) |
| C14 | C15 | 1.528(3) |
| C14 | C15A | 1.538(3) |
| C15 | C16 | 1.537(3) |
| C15A | C16A | 1.538(3) |
| C16 | C17 | 1.541(3) |
| C16A | C17A | 1.430(17) |
| C17 | C18 | 1.509(16) |
| C17A | C18 | 1.53(3) |
| C18 | C19 | 1.523(4) |
| C19 | C20 | 1.533(4) |
| C20 | C21 | 1.542(3) |
| C21 | C22 | 1.531(4) |
| C22 | C23 | 1.522(3) |
| C23 | C24 | 1.520(4) |

Bond Angles in ° for **2**.

| Atom | Atom | Atom | Angle/° |
|---|---|---|---|
| C7 | O1 | C25 | 115.6(2) |
| C8 | O2 | C23 | 117.28(18) |
| C1 | N1 | C4 | 109.6(2) |
| C5 | N2 | C8 | 106.1(2) |
| C13 | N3 | C10 | 110.1(2) |
| N1 | C1 | C2 | 108.5(2) |
| C1 | C2 | C3 | 107.0(2) |
| C4 | C3 | C2 | 107.2(2) |
| N1 | C4 | C3 | 107.6(2) |
| N1 | C4 | C5 | 120.7(2) |
| C3 | C4 | C5 | 131.5(2) |
| N2 | C5 | C4 | 120.9(2) |
| N2 | C5 | C6 | 115.3(2) |
| C4 | C5 | C6 | 123.8(2) |
| C7 | C6 | C5 | 104.5(2) |

| Atom | Atom | Atom | Angle/° |
|---|---|---|---|
| N3 | C10 | C9 | 122.5(2) |
| C11 | C10 | N3 | 107.0(2) |
| C11 | C10 | C9 | 129.1(2) |
| C10 | C11 | C12 | 107.6(2) |
| C13 | C12 | C11 | 108.3(2) |
| N3 | C13 | C14 | 122.4(3) |
| C12 | C13 | N3 | 106.9(2) |
| C12 | C13 | C14 | 130.6(3) |
| C13 | C14 | C15 | 118.9(3) |
| C13 | C14 | C15A | 113.5(4) |
| C14 | C15 | C16 | 113.9(4) |
| C14 | C15A | C16A | 125.4(7) |
| C15 | C16 | C17 | 112.2(6) |
| C17A | C16A | C15A | 117.2(13) |
| C18 | C17 | C16 | 108.9(8) |

| Atom | Atom | Atom | Angle/° |
|---|---|---|---|
| O1 | C7 | C6 | 133.2(2) |
| O1 | C7 | C8 | 117.4(2) |
| C6 | C7 | C8 | 109.5(2) |
| O2 | C8 | N2 | 112.2(2) |
| O2 | C8 | C7 | 104.97(19) |
| O2 | C8 | C9 | 111.56(19) |
| N2 | C8 | C7 | 104.66(19) |
| N2 | C8 | C9 | 110.9(2) |
| C7 | C8 | C9 | 112.3(2) |
| C8 | C9 | C21 | 111.23(19) |
| C10 | C9 | C8 | 114.7(2) |
| C10 | C9 | C21 | 110.9(2) |
| C16A | C17A | C18 | 129.9(19) |
| C17 | C18 | C19 | 114.5(3) |
| C19 | C18 | C17A | 111.9(8) |
| C18 | C19 | C20 | 114.3(2) |
| C19 | C20 | C21 | 114.2(2) |
| C20 | C21 | C9 | 111.5(2) |
| C22 | C21 | C9 | 109.8(2) |
| C22 | C21 | C20 | 110.2(2) |
| C23 | C22 | C21 | 112.3(2) |
| O2 | C23 | C22 | 109.70(19) |
| O2 | C23 | C24 | 112.8(2) |
| C24 | C23 | C22 | 114.1(2) |

Torsion Angles in ° for **2**.

| Atom | Atom | Atom | Atom | Angle/° |
|---|---|---|---|---|
| O1 | C7 | C8 | O2 | -60.4(3) |
| O1 | C7 | C8 | N2 | -178.7(2) |
| O1 | C7 | C8 | C9 | 61.0(3) |
| O2 | C8 | C9 | C10 | -177.6(2) |
| O2 | C8 | C9 | C21 | -50.8(3) |
| N1 | C1 | C2 | C3 | -0.3(3) |
| N1 | C4 | C5 | N2 | 6.7(4) |
| N1 | C4 | C5 | C6 | -174.1(2) |
| N2 | C5 | C6 | C7 | -0.8(3) |
| N2 | C8 | C9 | C10 | -51.8(3) |
| N2 | C8 | C9 | C21 | 75.0(2) |
| N3 | C10 | C11 | C12 | 0.9(3) |
| N3 | C13 | C14 | C15 | 93.1(4) |
| N3 | C13 | C14 | C15A | 56.3(5) |
| C1 | N1 | C4 | C3 | 0.4(3) |
| C1 | N1 | C4 | C5 | -176.1(2) |
| C1 | C2 | C3 | C4 | 0.5(3) |
| C2 | C3 | C4 | N1 | -0.6(3) |
| C2 | C3 | C4 | C5 | 175.4(3) |
| C3 | C4 | C5 | N2 | -168.8(3) |
| C3 | C4 | C5 | C6 | 10.4(4) |
| C4 | N1 | C1 | C2 | -0.1(3) |
| C4 | C5 | C6 | C7 | 179.9(2) |
| C5 | N2 | C8 | O2 | -115.1(2) |
| C5 | N2 | C8 | C7 | -1.8(2) |
| C5 | N2 | C8 | C9 | 119.5(2) |
| C13 | N3 | C10 | C11 | -1.4(3) |
| C13 | C14 | C15 | C16 | -63.3(5) |
| C13 | C14 | C15A | C16A | 58.9(9) |
| C14 | C15 | C16 | C17 | 79.3(8) |
| C14 | C15A | C16A | C17A | -81.7(18) |
| C15 | C16 | C17 | C18 | -151.5(5) |
| C15A | C16A | C17A | C18 | -54(3) |
| C16 | C17 | C18 | C19 | 172.2(5) |
| C16A | C17A | C18 | C19 | 171.4(18) |
| C17 | C18 | C19 | C20 | -61.5(6) |
| C17A | C18 | C19 | C20 | -72.1(10) |
| C18 | C19 | C20 | C21 | -71.1(3) |
| C5 | C6 | C7 | O1 | 179.6(3) |
| C5 | C6 | C7 | C8 | -0.4(3) |
| C6 | C7 | C8 | O2 | 119.6(2) |
| C6 | C7 | C8 | N2 | 1.4(3) |
| C6 | C7 | C8 | C9 | -119.0(2) |
| C7 | C8 | C9 | C10 | 64.9(3) |
| C7 | C8 | C9 | C21 | -168.3(2) |
| C8 | O2 | C23 | C22 | -54.8(3) |
| C8 | O2 | C23 | C24 | 73.5(3) |
| C8 | N2 | C5 | C4 | -179.0(2) |
| C8 | N2 | C5 | C6 | 1.7(3) |
| C8 | C9 | C10 | N3 | 79.3(3) |
| C8 | C9 | C10 | C11 | -116.0(3) |
| C8 | C9 | C21 | C20 | 174.2(2) |
| C8 | C9 | C21 | C22 | 51.9(3) |
| C9 | C10 | C11 | C12 | -165.7(2) |
| C9 | C21 | C22 | C23 | -54.5(3) |
| C10 | N3 | C13 | C12 | 1.4(3) |
| C10 | N3 | C13 | C14 | -174.9(2) |
| C10 | C9 | C21 | C20 | -56.9(3) |
| C10 | C9 | C21 | C22 | -179.3(2) |
| C10 | C11 | C12 | C13 | -0.1(3) |
| C11 | C12 | C13 | N3 | -0.8(3) |
| C11 | C12 | C13 | C14 | 175.0(3) |
| C12 | C13 | C14 | C15 | -82.2(4) |
| C13 | N3 | C10 | C9 | 166.2(2) |
| C19 | C20 | C21 | C9 | 150.5(2) |
| C19 | C20 | C21 | C22 | -87.4(3) |
| C20 | C21 | C22 | C23 | -177.6(2) |
| C21 | C9 | C10 | N3 | -47.7(3) |
| C21 | C9 | C10 | C11 | 117.0(3) |
| C21 | C22 | C23 | O2 | 54.0(3) |
| C21 | C22 | C23 | C24 | -73.6(3) |
| C23 | O2 | C8 | N2 | -71.3(2) |
| C23 | O2 | C8 | C7 | 175.62(19) |
| C23 | O2 | C8 | C9 | 53.7(3) |
| C25 | O1 | C7 | C6 | 2.2(4) |
| C25 | O1 | C7 | C8 | -177.7(2) |

Hydrogen Fractional Atomic Coordinates (×$10^4$) and Equivalent Isotropic Displacement Parameters (Å$^2$×$10^3$) for **2**. $U_{eq}$ is defined as 1/3 of the trace of the orthogonalised $U_{ij}$.

| Atom | x | y | z | *U_eq* |
|---|---|---|---|---|
| H3B | 8250(60) | 5750(40) | 3040(20) | 88(16) |
| H3C | 9540(50) | 5240(30) | 2658(11) | 71(13) |
| H1 | 225.6 | 7189.8 | 3020.87 | 28 |
| H3 | 1149.39 | 5203.97 | 3880.29 | 26 |
| H1A | -1790.08 | 8589.96 | 2939.7 | 32 |
| H2 | -609.92 | 10183.97 | 3458.49 | 31 |
| H3A | 2206.62 | 9664.69 | 3863.52 | 27 |
| H6 | 4781.98 | 8395.89 | 4150.61 | 24 |
| H9 | 5599.3 | 4837.18 | 3970.87 | 24 |
| H11 | 4645.47 | 4903.83 | 5129.56 | 29 |
| H12 | 1876.44 | 4690.05 | 5650.11 | 34 |
| H14C | -1489.26 | 5451.2 | 4716.13 | 42 |
| H14D | -1246.72 | 4618.97 | 5263.24 | 42 |
| H14A | -1456.16 | 5196.23 | 4495.61 | 42 |
| H14B | -1312.37 | 5096.63 | 5223.86 | 42 |
| H15A | -2873.56 | 3702.66 | 4870.31 | 36 |
| H15B | -1197.05 | 3228.57 | 5134.89 | 36 |
| H15C | -1414.76 | 4116.44 | 3988.54 | 37 |
| H15D | -2832.93 | 3898.66 | 4470.78 | 37 |
| H16A | -1536.43 | 3601.77 | 3842.23 | 37 |
| H16B | -2175.48 | 2513.66 | 4134.78 | 37 |
| H16C | -1011.73 | 2517.94 | 4887.43 | 60 |
| H16D | -2022.99 | 2286.58 | 4274.06 | 60 |
| H17A | 1149.66 | 3077.99 | 4380.94 | 42 |
| H17B | 398.06 | 1908.17 | 4315.84 | 42 |
| H17C | 452.64 | 1645.37 | 4202.58 | 66 |
| H17D | 1253.21 | 2735.9 | 4393.55 | 66 |
| H18A | 702.78 | 3310.58 | 3291.93 | 38 |
| H18B | 194.7 | 2089.99 | 3254.2 | 38 |
| H18C | 194.56 | 2177.83 | 3198.33 | 38 |
| H18D | 741.63 | 3359.85 | 3369.34 | 38 |
| H19A | 2895.03 | 2265.99 | 2924.06 | 33 |
| H19B | 2875.16 | 1587.33 | 3544.66 | 33 |
| H20A | 5023.23 | 2737.87 | 3655.45 | 30 |
| H20B | 3618.7 | 3187.48 | 4097.15 | 30 |
| H21 | 2927.33 | 4326.7 | 3161.01 | 25 |
| H22A | 6347.9 | 3874.94 | 2972.18 | 29 |
| H22B | 4904.84 | 3573.9 | 2504.17 | 29 |
| H23 | 6452.18 | 5103.27 | 2212.36 | 27 |
| H24A | 3021.92 | 5457.81 | 2322.92 | 39 |
| H24B | 3956.8 | 4984.29 | 1736.64 | 39 |
| H24C | 4207.02 | 6199.6 | 1923.99 | 39 |
| H25A | 7823.58 | 8178.2 | 4276.23 | 40 |
| H25B | 8895.13 | 7280.42 | 4606.19 | 40 |
| H25C | 7100.64 | 7557.74 | 4858.37 | 40 |

Atomic Occupancies for all atoms that are not fully occupied in **2**.

| Atom | Occupancy | Atom | Occupancy | Atom | Occupancy |
|---|---|---|---|---|---|
| H14C | 0.379(8) | H15D | 0.379(8) | H17B | 0.621(8) |
| H14D | 0.379(8) | C16 | 0.621(8) | C17A | 0.379(8) |
| H14A | 0.621(8) | H16A | 0.621(8) | H17C | 0.379(8) |
| H14B | 0.621(8) | H16B | 0.621(8) | H17D | 0.379(8) |
| C15 | 0.621(8) | C16A | 0.379(8) | H18A | 0.621(8) |
| H15A | 0.621(8) | H16C | 0.379(8) | H18B | 0.621(8) |
| H15B | 0.621(8) | H16D | 0.379(8) | H18C | 0.379(8) |
| C15A | 0.379(8) | C17 | 0.621(8) | H18D | 0.379(8) |
| H15C | 0.379(8) | H17A | 0.621(8) | | |

#### Supplementary Data 2. X-Ray Crystal Structure Determination for Gem-dimethyl-bridged premarineosin A.

**Experimental.** Single orange irregular-shaped crystals of **3** were used as supplied. A suitable crystal with dimensions 0.20 × 0.18 × 0.12 mm$^3$ was selected and mounted on a dtrek-CrysAlisPro-abstract goniometer imported rigaku-d*trek images diffractometer. The crystal was kept at a steady *T* = 85 K during data collection. The structure was solved with the ShelXT 2018/2[10] solution program using dual methods and by using Olex2 1.5-alpha[11] as the graphical interface. The model was refined with ShelXL 2019/3[12] using full matrix least squares minimisation on $\boldsymbol{F}^2$.

**Crystal Data.** $C_{57}H_{72}F_6N_6O_8$, $M_r$ = 1083.20, orthorhombic, $C222_1$ (No. 20), a = 15.27900(10) Å, b = 20.9910(2) Å, c = 17.29930(10) Å, *a* = *b* = *g* = 90˚, *V* = 5548.26(7) Å$^3$, *T* = 85 K, *Z* = 4, *Z'* = 0.5, *m*(Cu K$_a$) = 0.838, 42424 reflections measured, 5150 unique ($R_{int}$ = 0.0425) which were used in all calculations. The final $wR_2$ was 0.1663 (all data) and $R_1$ was 0.0614 (I≥2 *s*(I)).

| | |
|---|---|
| Formula | $C_{57}H_{72}F_6N_6O_8$ |
| $D_{calc.}$/ g cm$^{-3}$ | 1.297 |
| *m*/mm$^{-1}$ | 0.838 |
| Formula Weight | 1083.20 |
| Colour | orange |
| Shape | irregular-shaped |
| Size/mm$^3$ | 0.20×0.18×0.12 |
| *T*/K | 85 |
| Crystal System | orthorhombic |
| Flack Parameter | 0.05(6) |
| Hooft Parameter | 0.04(5) |
| Space Group | $C222_1$ |
| *a*/Å | 15.27900(10) |
| *b*/Å | 20.9910(2) |
| *c*/Å | 17.29930(10) |
| *a*/˚ | 90 |
| *b*/˚ | 90 |
| *g*/˚ | 90 |
| V/Å$^3$ | 5548.26(7) |
| *Z* | 4 |
| *Z'* | 0.5 |
| Wavelength/Å | 1.54184 |
| Radiation type | Cu K$_a$ |
| $Q_{min}$/˚ | 3.578 |
| $Q_{max}$/˚ | 69.305 |
| Measured Refl's. | 42424 |
| Indep't Refl's | 5150 |
| Refl's I≥2 *s*(I) | 4984 |
| $R_{int}$ | 0.0425 |
| Parameters | 402 |
| Restraints | 157 |
| Largest Peak | 0.529 |
| Deepest Hole | -0.386 |
| GooF | 1.069 |
| $wR_2$ (all data) | 0.1663 |
| $wR_2$ | 0.1631 |
| $R_1$ (all data) | 0.0630 |
| $R_1$ | 0.0614 |

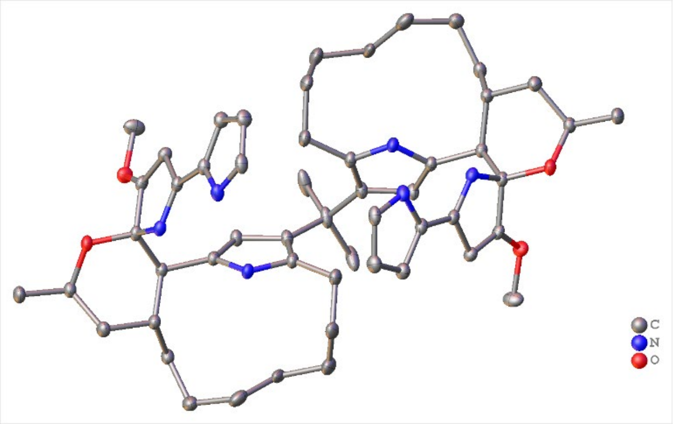

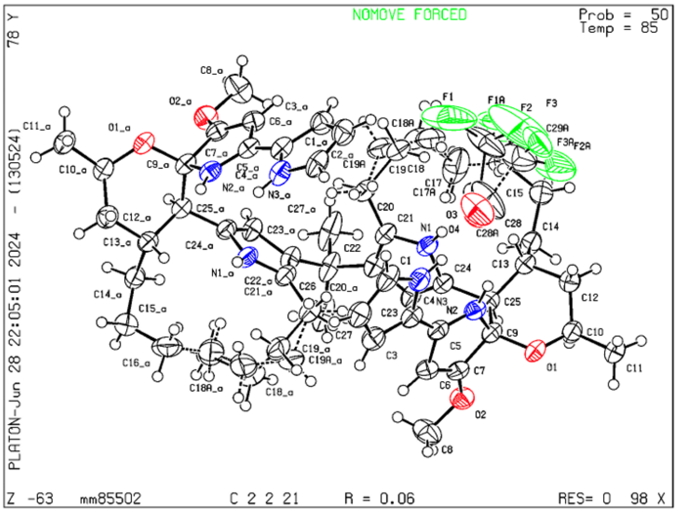

##### Structure Quality Indicators

| Reflections: | d min (CuKα) 2Θ=138.6° 0.82 | I/σ(I) 48.7 | Rint m=8.25 4.25% | Full 135.4° 100 | |
|---|---|---|---|---|---|
| **Refinement:** | Shift -0.001 | Max Peak 0.5 | Min Peak -0.4 | GooF 1.069 | Hooft .04(5) |

An orange irregular-shaped-shaped crystal with dimensions 0.20 × 0.18 × 0.12 $mm^3$ was mounted. Data were collected using a dtrek-CrysAlisPro-abstract goniometer imported rigaku-d*trek images diffractometer operating at $T$ = 85 K.

Data were measured using *w* scans with Cu $K_a$ radiation. The diffraction pattern was indexed and the total number of runs and images was based on the strategy calculation from the program DTREK_VERSION=d*TREK version 9.9.9.4 W9RSSI – Aug 22 2012. The maximum resolution that was achieved was $Q$ = 69.305° (0.82 Å).

The unit cell was refined using CrysAlisPro 1.171.43.127a[13] on 26227 reflections, 62% of the observed reflections.

Data reduction, scaling and absorption corrections were performed using CrysAlisPro 1.171.43.127a[13]. The final completeness is 100.00 % out to 69.305° in $Q$. A multi-scan absorption correction was performed using CrysAlisPro 1.171.43.127a[13]. Empirical absorption correction using spherical harmonics, implemented in SCALE3 ABSPACK scaling algorithm.. The absorption coefficient *m* of this material is 0.838 $mm^{-1}$ at this wavelength ($l$ = 1.54184Å) and the minimum and maximum transmissions are 0.972 and 1.000.

The structure was solved and the space group $C222_1$ (# 20) determined by the ShelXT 2018/2[10] structure solution program using using dual methods and refined by full matrix least squares minimization on $F^2$ using version 2019/3 of ShelXL 2019/3[12]. All non-hydrogen atoms were refined anisotropically. Hydrogen atom positions were calculated geometrically and refined using the riding model. Hydrogen atom positions were calculated geometrically and refined using the riding model.

*_refine_special_details*: A trifluoroacetate was found disordered at two positions. It was modelled as such with the implementation of some restraints.

*_exptl_absorpt_process_details*: CrysAlisPro 1.171.43.127a[13] using spherical harmonics, implemented in SCALE3 ABSPACK scaling algorithm.

The value of Z' is 0.5. This means that only half of the formula unit is present in the asymmetric unit, with the other half consisting of symmetry equivalent atoms. The moiety formula is C53 H72 N6 O4, C4 F6 O4.

The Flack parameter was refined to 0.05(6). Determination of absolute structure using Bayesian statistics on Bijvoet differences using the Olex2 results in 0.04(5). The chiral atoms in this structure are: C9(S), C10(R), C13(R), C25(R). Note: The Flack parameter is used to determine chirality of the crystal studied, the value should be near 0, a value of 1 means that the stereochemistry is wrong and the model should be inverted. A value of 0.5 means that the crystal consists of a racemic mixture of the two enantiomers.

Fractional Atomic Coordinates (×104) and Equivalent Isotropic Displacement Parameters (Å2×103) for 3. Ueq is defined as 1/3 of the trace of the orthogonalised Uij.

| Atom | x | y | z | $U_{eq}$ |
|---|---|---|---|---|
| O1 | 2633.1(16) | 5361.9(12) | 1239.3(13) | 42.8(6) |
| O2 | 1469.0(17) | 4381.1(13) | 1833.2(16) | 47.4(6) |
| N1 | 2151(2) | 5798.2(15) | 3788.8(16) | 41.9(7) |
| N2 | 3311.9(18) | 5256.4(14) | 2466.1(16) | 38.4(6) |
| N3 | 4852.7(19) | 4936.6(16) | 3502.1(17) | 45.4(7) |
| C1 | 5425(2) | 4610(2) | 3944(2) | 51.0(9) |
| C2 | 5162(3) | 3986(2) | 3998(2) | 53.2(9) |
| C3 | 4391(3) | 3932.2(19) | 3585(2) | 47.9(8) |
| C4 | 4194(2) | 4530.8(19) | 3265.6(19) | 41.5(8) |
| C5 | 3483(2) | 4692.0(17) | 2781.6(18) | 38.4(7) |
| C6 | 2814(2) | 4235.9(17) | 2569(2) | 40.4(7) |
| C7 | 2227(2) | 4561.0(17) | 2142.6(19) | 38.6(7) |
| C8 | 1213(3) | 3733(2) | 2002(3) | 63.8(12) |
| C9 | 2483(2) | 5246.5(17) | 2028.5(18) | 37.4(7) |
| C10 | 2253(3) | 5943.1(18) | 917(2) | 43.7(8) |
| C11 | 2652(3) | 6003(2) | 117(2) | 52.6(9) |
| C12 | 2432(3) | 6503.0(18) | 1440(2) | 47.7(8) |
| C13 | 2020(2) | 6407.0(17) | 2242(2) | 40.9(7) |
| C14 | 1202(3) | 6814.8(19) | 2387(2) | 46.7(8) |
| C15 | 1389(4) | 7469(2) | 2728(3) | 59.6(11) |
| C16 | 1801(4) | 7463(2) | 3532(3) | 72.1(14) |
| C17 | 1226(14) | 7146(7) | 4159(7) | 63(5) |
| C17A | 1360(20) | 7100(17) | 4191(14) | 106(15) |
| C18 | 1562(13) | 7072(6) | 4998(7) | 74(4) |
| C18A | 2090(30) | 7106(9) | 4796(16) | 99(10) |
| C19 | 2386(10) | 6663(4) | 5124(6) | 57(3) |
| C19A | 1890(20) | 6618(7) | 5401(11) | 73(7) |
| C20 | 2218(3) | 5939(2) | 5203(2) | 51.4(9) |
| C21 | 1805(2) | 5653.8(18) | 4497.7(19) | 42.5(8) |
| C22 | 1060(2) | 5299.7(19) | 4383.1(19) | 43.0(8) |
| C23 | 941(2) | 5252.5(18) | 3562.3(19) | 40.9(7) |
| C24 | 1616(2) | 5561.9(16) | 3215.3(18) | 38.4(7) |
| C25 | 1795(2) | 5698.4(16) | 2381.9(18) | 37.1(7) |
| C26 | 486(4) | 5000 | 5000 | 54.6(15) |
| C27 | -102(3) | 4482(3) | 4642(2) | 78.9(18) |
| C29 | 4742(3) | 7121.6(14) | 3908(3) | 106(4) |
| F1 | 4977(4) | 7077(3) | 4659(3) | 250(9) |
| F2 | 4023(3) | 7496.0(16) | 3898(4) | 140(4) |
| F3 | 5384(3) | 7467.0(15) | 3560(5) | 245(8) |
| O3 | 3840.0(14) | 6333.3(13) | 3347.6(19) | 61.1(8) |
| O4 | 5308.8(15) | 6175.0(16) | 3364(3) | 83.2(12) |
| C28 | 4613.8(15) | 6458.1(10) | 3494.7(17) | 74(3) |
| C29A | 4736(5) | 7216(2) | 3306(4) | 94(4) |
| F1A | 4486(7) | 7505(5) | 3969(5) | 121(5) |
| F2A | 4232(7) | 7477(4) | 2741(6) | 148(6) |
| F3A | 5559(5) | 7423(4) | 3169(6) | 107(4) |
| C28A | 4641(6) | 6468(2) | 3319(8) | 97(6) |

Anisotropic Displacement Parameters (×10[4]) for **3**. The anisotropic displacement factor exponent takes the form: $-2p^2[h^2a^{*2} \times U_{11} + ... +2hka^* \times b^* \times U_{12}]$.

| Atom | $U_{11}$ | $U_{22}$ | $U_{33}$ | $U_{23}$ | $U_{13}$ | $U_{12}$ |
|---|---|---|---|---|---|---|
| O1 | 47.3(13) | 53.4(13) | 27.7(11) | -2.9(9) | 1.3(10) | 0.2(11) |
| O2 | 41.9(13) | 50.8(14) | 49.6(14) | -2.6(11) | -12.5(11) | -5.7(11) |
| N1 | 49.9(16) | 46.7(15) | 29.3(13) | -5.4(11) | -1.2(12) | -4.6(13) |
| N2 | 35.3(13) | 47.0(15) | 32.7(12) | -5.5(12) | 0.4(11) | -6.2(12) |

| Atom | $U_{11}$ | $U_{22}$ | $U_{33}$ | $U_{23}$ | $U_{13}$ | $U_{12}$ |
|---|---|---|---|---|---|---|
| N3 | 42.7(16) | 58.8(18) | 34.8(14) | 1.6(13) | -1.6(12) | -10.1(14) |
| C1 | 38.2(18) | 82(3) | 33.1(16) | 0.7(17) | -4.8(14) | -0.4(18) |
| C2 | 48(2) | 66(2) | 46.3(19) | 3.3(18) | -9.5(17) | 3.5(18) |
| C3 | 45.1(19) | 55(2) | 44.0(19) | -1.9(16) | -4.9(15) | 2.5(16) |
| C4 | 34.8(16) | 57(2) | 32.3(15) | -4.2(14) | 1.1(13) | -4.1(14) |
| C5 | 37.3(16) | 48.4(18) | 29.5(14) | -5.9(13) | 2.4(12) | -4.1(14) |
| C6 | 38.2(17) | 42.6(16) | 40.2(17) | -5.1(14) | -3.6(14) | -3.0(14) |
| C7 | 35.4(16) | 46.9(17) | 33.4(15) | -8.5(13) | -4.3(13) | -2.6(14) |
| C8 | 52(2) | 51(2) | 88(3) | 4(2) | -22(2) | -14.9(18) |
| C9 | 36.3(16) | 47.5(17) | 28.3(15) | -3.9(13) | 0.5(12) | -4.4(14) |
| C10 | 48.5(19) | 50.2(18) | 32.5(16) | 1.0(14) | 0.4(14) | -1.9(15) |
| C11 | 67(2) | 57(2) | 33.5(17) | 0.6(16) | 6.6(17) | -4.2(19) |
| C12 | 55(2) | 49.2(19) | 39.3(18) | 0.2(15) | 5.8(16) | -5.8(16) |
| C13 | 46.7(18) | 43.1(17) | 33.0(16) | -2.1(13) | 2.4(14) | -0.8(14) |
| C14 | 53(2) | 53(2) | 34.1(16) | 2.0(15) | -0.1(15) | 8.4(17) |
| C15 | 82(3) | 47(2) | 50(2) | -0.8(17) | 6(2) | 8(2) |
| C16 | 93(4) | 55(2) | 68(3) | -26(2) | -12(3) | 11(2) |
| C17 | 117(10) | 37(5) | 35(7) | -10(4) | -20(7) | 12(6) |
| C17A | 170(30) | 91(18) | 58(16) | 8(11) | 29(15) | 76(17) |
| C18 | 118(10) | 62(5) | 42(5) | -16(4) | -11(6) | 2(6) |
| C18A | 190(30) | 47(7) | 62(12) | -14(8) | -43(16) | 4(13) |
| C19 | 79(7) | 56(4) | 36(4) | -7(3) | -11(4) | -10(4) |
| C19A | 119(17) | 52(7) | 47(8) | -17(6) | -12(10) | 15(8) |
| C20 | 66(2) | 57(2) | 31.3(17) | -5.3(15) | -5.9(16) | 0.2(19) |
| C21 | 47.0(19) | 51.3(19) | 29.2(14) | -3.6(14) | -0.1(14) | 3.5(16) |
| C22 | 38.6(16) | 60(2) | 30.4(16) | 3.7(14) | -0.7(13) | 2.1(16) |
| C23 | 36.1(15) | 55.7(19) | 30.8(16) | 1.6(14) | -2.6(13) | -1.0(14) |
| C24 | 41.7(17) | 42.9(16) | 30.5(15) | -3.3(13) | -2.1(13) | 1.8(13) |
| C25 | 38.1(16) | 44.2(17) | 29.1(15) | -2.4(13) | 0.9(12) | -1.3(14) |
| C26 | 40(3) | 96(4) | 28(2) | 10(3) | 0 | 0 |
| C27 | 57(2) | 142(5) | 38(2) | 25(3) | -9.1(18) | -42(3) |
| C29 | 92(6) | 42(4) | 183(10) | 23(5) | -95(7) | -17(4) |
| F1 | 440(20) | 78(4) | 238(11) | -51(6) | -256(13) | 43(8) |
| F2 | 169(8) | 63(4) | 187(9) | -23(5) | -94(7) | -3(5) |
| F3 | 142(9) | 114(8) | 480(20) | 9(12) | -69(13) | -89(7) |
| O3 | 48.0(15) | 59.4(17) | 76(2) | 2.4(15) | -18.2(15) | -7.6(12) |
| O4 | 53.7(18) | 66(2) | 130(4) | 16(2) | -24(2) | -4.4(14) |
| C28 | 61(5) | 59(6) | 102(7) | 2(5) | -41(5) | -16(4) |
| C29A | 93(8) | 59(6) | 129(10) | -6(7) | -33(8) | 15(6) |
| F1A | 178(13) | 55(5) | 130(8) | -36(5) | -59(8) | -22(7) |
| F2A | 196(12) | 62(5) | 184(11) | -7(6) | -105(11) | 35(6) |
| F3A | 143(8) | 53(4) | 126(7) | 34(4) | -77(6) | -29(5) |
| C28A | 60(6) | 53(8) | 178(16) | 42(8) | -19(9) | 6(6) |

Bond Lengths in Å for **3**.

| Atom | Atom | Length/Å | Atom | Atom | Length/Å |
|---|---|---|---|---|---|
| O1 | C9 | 1.405(4) | C17A | C18A | 1.53(2) |
| O1 | C10 | 1.462(5) | C18 | C19 | 1.54(2) |
| O2 | C7 | 1.330(4) | C18A | C19A | 1.49(5) |
| O2 | C8 | 1.445(5) | C19 | C20 | 1.547(9) |
| N1 | C21 | 1.369(5) | C19A | C20 | 1.548(14) |
| N1 | C24 | 1.378(4) | C20 | C21 | 1.499(5) |
| N2 | C5 | 1.330(5) | C21 | C22 | 1.373(5) |
| N2 | C9 | 1.475(4) | C22 | C23 | 1.435(4) |
| N3 | C1 | 1.349(5) | C22 | C26 | 1.518(5) |
| N3 | C4 | 1.380(5) | C23 | C24 | 1.359(5) |
| C1 | C2 | 1.372(7) | C24 | C25 | 1.495(4) |
| C2 | C3 | 1.383(6) | C26 | C27[1] | 1.542(6) |
| C3 | C4 | 1.405(6) | C26 | C27 | 1.542(6) |

| Atom | Atom | Length/Å |
|---|---|---|
| C4 | C5 | 1.413(5) |
| C5 | C6 | 1.447(5) |
| C6 | C7 | 1.348(5) |
| C7 | C9 | 1.504(5) |
| C9 | C25 | 1.542(5) |
| C10 | C11 | 1.517(5) |
| C10 | C12 | 1.508(5) |
| C12 | C13 | 1.537(5) |
| C13 | C14 | 1.536(5) |
| C13 | C25 | 1.546(5) |
| C14 | C15 | 1.522(6) |
| C15 | C16 | 1.526(7) |
| C16 | C17 | 1.547(15) |
| C16 | C17A | 1.53(2) |
| C17 | C18 | 1.548(15) |

| Atom | Atom | Length/Å |
|---|---|---|
| C29 | F1 | 1.3500 |
| C29 | F2 | 1.3506 |
| C29 | F3 | 1.3607 |
| C29 | C28 | 1.5779 |
| O3 | C28 | 1.2374 |
| O3 | C28A | 1.257(10) |
| O4 | C28 | 1.2376 |
| O4 | C28A | 1.193(10) |
| C29A | F1A | 1.3508 |
| C29A | F2A | 1.3604 |
| C29A | F3A | 1.3508 |
| C29A | C28A | 1.5785 |
| F2A | F3A[2] | 1.61(2) |

[1]+x,1-y,1-z; [2]1-x,+y,1/2-z

Bond Angles in ° for **3**.

| Atom | Atom | Atom | Angle/° |
|---|---|---|---|
| C9 | O1 | C10 | 116.7(3) |
| C7 | O2 | C8 | 115.0(3) |
| C21 | N1 | C24 | 109.7(3) |
| C5 | N2 | C9 | 111.5(3) |
| C1 | N3 | C4 | 109.1(3) |
| N3 | C1 | C2 | 109.5(3) |
| C1 | C2 | C3 | 107.0(4) |
| C2 | C3 | C4 | 108.2(4) |
| N3 | C4 | C3 | 106.2(3) |
| N3 | C4 | C5 | 126.1(3) |
| C3 | C4 | C5 | 127.7(3) |
| N2 | C5 | C4 | 127.4(3) |
| N2 | C5 | C6 | 110.3(3) |
| C4 | C5 | C6 | 122.3(3) |
| C7 | C6 | C5 | 105.9(3) |
| O2 | C7 | C6 | 131.0(3) |
| O2 | C7 | C9 | 116.5(3) |
| C6 | C7 | C9 | 112.5(3) |
| O1 | C9 | N2 | 110.9(3) |
| O1 | C9 | C7 | 109.6(3) |
| O1 | C9 | C25 | 113.0(3) |
| N2 | C9 | C7 | 99.8(3) |
| N2 | C9 | C25 | 111.9(3) |
| C7 | C9 | C25 | 111.0(3) |
| O1 | C10 | C11 | 105.0(3) |
| O1 | C10 | C12 | 110.5(3) |
| C12 | C10 | C11 | 114.2(3) |
| C10 | C12 | C13 | 111.5(3) |
| C12 | C13 | C25 | 111.0(3) |
| C14 | C13 | C12 | 114.0(3) |
| C14 | C13 | C25 | 109.3(3) |
| C15 | C14 | C13 | 114.4(3) |
| C14 | C15 | C16 | 115.0(4) |
| C15 | C16 | C17 | 114.1(7) |
| C15 | C16 | C17A | 120.0(12) |
| C16 | C17 | C18 | 120.8(13) |
| C16 | C17A | C18A | 100.4(19) |
| C19 | C18 | C17 | 117.3(14) |
| C19A | C18A | C17A | 109(3) |
| C18 | C19 | C20 | 115.2(12) |
| C18A | C19A | C20 | 114(3) |
| C21 | C20 | C19 | 113.0(4) |

| Atom | Atom | Atom | Angle/° |
|---|---|---|---|
| C21 | C20 | C19A | 114.4(6) |
| N1 | C21 | C20 | 118.6(3) |
| N1 | C21 | C22 | 108.1(3) |
| C22 | C21 | C20 | 133.1(3) |
| C21 | C22 | C23 | 106.6(3) |
| C21 | C22 | C26 | 127.0(3) |
| C23 | C22 | C26 | 126.4(3) |
| C24 | C23 | C22 | 107.9(3) |
| N1 | C24 | C25 | 121.1(3) |
| C23 | C24 | N1 | 107.7(3) |
| C23 | C24 | C25 | 131.1(3) |
| C9 | C25 | C13 | 112.2(3) |
| C24 | C25 | C9 | 112.9(3) |
| C24 | C25 | C13 | 112.1(3) |
| C22 | C26 | C22[1] | 109.4(4) |
| C22 | C26 | C27 | 110.3(2) |
| C22[1] | C26 | C27 | 109.1(3) |
| C22[1] | C26 | C27[1] | 110.3(2) |
| C22 | C26 | C27[1] | 109.1(3) |
| C27[1] | C26 | C27 | 108.6(6) |
| F1 | C29 | F2 | 105.6 |
| F1 | C29 | F3 | 105.7 |
| F1 | C29 | C28 | 114.1 |
| F2 | C29 | F3 | 105.7 |
| F2 | C29 | C28 | 114.0 |
| F3 | C29 | C28 | 111.0 |
| O3 | C28 | C29 | 113.5 |
| O3 | C28 | O4 | 132.9 |
| O4 | C28 | C29 | 113.6 |
| F1A | C29A | F2A | 105.6 |
| F1A | C29A | C28A | 114.1 |
| F2A | C29A | C28A | 111.0 |
| F3A | C29A | F1A | 105.6 |
| F3A | C29A | F2A | 105.7 |
| F3A | C29A | C28A | 114.1 |
| C29A | F2A | F3A[2] | 124.2(10) |
| O3 | C28A | C29A | 108.3(5) |
| O4 | C28A | O3 | 135.6(5) |
| O4 | C28A | C29A | 115.8(5) |

[1]+x,1-y,1-z; [2]1-x,+y,1/2-z

Torsion angles in ° for **3**.

| Atom | Atom | Atom | Atom | Angle/° |
|---|---|---|---|---|
| O1 | C9 | C25 | C13 | -54.0(4) |
| O1 | C9 | C25 | C24 | 178.2(3) |
| O1 | C10 | C12 | C13 | -61.3(4) |
| O2 | C7 | C9 | O1 | -65.5(4) |
| O2 | C7 | C9 | N2 | 178.1(3) |
| O2 | C7 | C9 | C25 | 59.9(4) |
| N1 | C21 | C22 | C23 | -2.3(4) |
| N1 | C21 | C22 | C26 | 176.2(3) |
| N1 | C24 | C25 | C9 | 82.8(4) |
| N1 | C24 | C25 | C13 | -45.1(4) |
| N2 | C5 | C6 | C7 | -2.4(4) |
| N2 | C9 | C25 | C13 | 72.0(3) |
| N2 | C9 | C25 | C24 | -55.8(4) |
| N3 | C1 | C2 | C3 | -1.1(5) |
| N3 | C4 | C5 | N2 | -0.1(6) |
| N3 | C4 | C5 | C6 | -179.3(3) |
| C1 | N3 | C4 | C3 | -0.1(4) |
| C1 | N3 | C4 | C5 | -178.6(3) |
| C1 | C2 | C3 | C4 | 1.0(5) |
| C2 | C3 | C4 | N3 | -0.6(4) |
| C2 | C3 | C4 | C5 | 177.9(4) |
| C3 | C4 | C5 | N2 | -178.2(3) |
| C3 | C4 | C5 | C6 | 2.5(6) |
| C4 | N3 | C1 | C2 | 0.7(4) |
| C4 | C5 | C6 | C7 | 176.9(3) |
| C5 | N2 | C9 | O1 | -116.6(3) |
| C5 | N2 | C9 | C7 | -1.2(3) |
| C5 | N2 | C9 | C25 | 116.3(3) |
| C5 | C6 | C7 | O2 | -176.5(3) |
| C5 | C6 | C7 | C9 | 1.6(4) |
| C6 | C7 | C9 | O1 | 116.1(3) |
| C6 | C7 | C9 | N2 | -0.4(3) |
| C6 | C7 | C9 | C25 | -118.5(3) |
| C7 | C9 | C25 | C13 | -177.5(3) |
| C7 | C9 | C25 | C24 | 54.7(4) |
| C8 | O2 | C7 | C6 | 2.4(6) |
| C8 | O2 | C7 | C9 | -175.7(4) |
| C9 | O1 | C10 | C11 | 170.0(3) |
| C9 | O1 | C10 | C12 | 46.5(4) |
| C9 | N2 | C5 | C4 | -177.0(3) |
| C9 | N2 | C5 | C6 | 2.3(4) |
| C10 | O1 | C9 | N2 | -115.9(3) |
| C10 | O1 | C9 | C7 | 134.9(3) |
| C10 | O1 | C9 | C25 | 10.6(4) |
| C10 | C12 | C13 | C14 | -106.1(4) |
| C10 | C12 | C13 | C25 | 17.8(4) |
| C11 | C10 | C12 | C13 | -179.3(3) |
| C12 | C13 | C14 | C15 | -89.1(4) |
| C12 | C13 | C25 | C9 | 36.8(4) |
| C12 | C13 | C25 | C24 | 165.1(3) |
| C13 | C14 | C15 | C16 | -63.6(5) |
| C14 | C13 | C25 | C9 | 163.4(3) |
| C14 | C13 | C25 | C24 | -68.3(4) |
| C14 | C15 | C16 | C17 | -61.4(9) |
| C14 | C15 | C16 | C17A | -54.5(18) |
| C15 | C16 | C17 | C18 | 177.2(11) |
| C15 | C16 | C17A | C18A | 168.0(14) |
| C16 | C17 | C18 | C19 | -61.8(19) |
| C16 | C17A | C18A | C19A | -164(3) |
| C17 | C18 | C19 | C20 | -85.1(18) |
| C17A | C18A | C19A | C20 | 86(4) |
| C18 | C19 | C20 | C21 | 61.2(13) |
| C18A | C19A | C20 | C21 | -63(3) |
| C19 | C20 | C21 | N1 | 47.4(8) |
| C19 | C20 | C21 | C22 | -125.4(8) |
| C19A | C20 | C21 | N1 | 84.3(14) |
| C19A | C20 | C21 | C22 | -88.5(15) |
| C20 | C21 | C22 | C23 | 171.1(4) |
| C20 | C21 | C22 | C26 | -10.4(7) |
| C21 | N1 | C24 | C23 | -1.6(4) |
| C21 | N1 | C24 | C25 | 174.7(3) |
| C21 | C22 | C23 | C24 | 1.3(4) |
| C21 | C22 | C26 | C22[1] | -42.5(3) |
| C21 | C22 | C26 | C27 | -162.5(4) |
| C21 | C22 | C26 | C27[1] | 78.3(5) |
| C22 | C23 | C24 | N1 | 0.1(4) |
| C22 | C23 | C24 | C25 | -175.7(4) |
| C23 | C22 | C26 | C22[1] | 135.7(4) |
| C23 | C22 | C26 | C27[1] | -103.5(4) |
| C23 | C22 | C26 | C27 | 15.7(6) |
| C23 | C24 | C25 | C9 | -101.9(4) |
| C23 | C24 | C25 | C13 | 130.2(4) |
| C24 | N1 | C21 | C20 | -172.1(3) |
| C24 | N1 | C21 | C22 | 2.4(4) |
| C25 | C13 | C14 | C15 | 146.1(3) |
| C26 | C22 | C23 | C24 | -177.2(3) |
| F1 | C29 | C28 | O3 | 109.1 |
| F1 | C29 | C28 | O4 | -72.7 |
| F2 | C29 | C28 | O3 | -12.3 |
| F2 | C29 | C28 | O4 | 165.9 |
| F3 | C29 | C28 | O3 | -131.6 |
| F3 | C29 | C28 | O4 | 46.6 |
| F1A | C29A | F2A | F3A[2] | 163.9(8) |
| F1A | C29A | F3A | F2A[2] | -147.5(3) |
| F1A | C29A | C28A | O3 | 67.2(8) |
| F1A | C29A | C28A | O4 | -106.9(10) |
| F2A | C29A | F3A | F2A[2] | -35.9(3) |
| F2A | C29A | C28A | O3 | -52.0(8) |
| F2A | C29A | C28A | O4 | 133.9(10) |
| F3A | C29A | F2A | F3A[2] | 52.3(8) |
| F3A | C29A | C28A | O3 | -171.3(8) |
| F3A | C29A | C28A | O4 | 14.5(10) |
| C28A | C29A | F2A | F3A[2] | -71.9(8) |
| C28A | C29A | F3A | F2A[2] | 86.4(3) |

[1]+x,1-y,1-z; [2]1-x,+y,1/2-z

Hydrogen Fractional Atomic Coordinates (×$10^4$) and Equivalent Isotropic Displacement Parameters (Å$^2$×$10^3$) for **3**. $U_{eq}$ is defined as 1/3 of the trace of the orthogonalised $U_{ij}$.

| Atom | x | y | z | $U_{eq}$ |
|---|---|---|---|---|
| H1 | 2640.15 | 6009.73 | 3711.32 | 50 |
| H2 | 3650.53 | 5593.34 | 2512.48 | 46 |
| H3 | 4892.76 | 5343.22 | 3383.65 | 55 |
| H1A | 5931.94 | 4784.27 | 4181.38 | 61 |
| H2A | 5454.23 | 3654.88 | 4268.57 | 64 |
| H3A | 4052.56 | 3555.43 | 3526.63 | 57 |
| H6 | 2794.26 | 3797.17 | 2702.8 | 48 |
| H8A | 641.61 | 3645.84 | 1766.61 | 96 |
| H8B | 1172.98 | 3675.29 | 2563.01 | 96 |
| H8C | 1649.29 | 3438.81 | 1790.69 | 96 |
| H10 | 1606.13 | 5885.76 | 865.34 | 52 |
| H11A | 3288.12 | 6051.09 | 162.52 | 79 |
| H11B | 2407.57 | 6377.79 | -143.73 | 79 |
| H11C | 2518.44 | 5620.44 | -184.75 | 79 |
| H12A | 2191.05 | 6894.73 | 1202.8 | 57 |
| H12B | 3072.06 | 6559.44 | 1496.33 | 57 |
| H13 | 2468.22 | 6532.45 | 2635.43 | 49 |
| H14A | 887.49 | 6870.72 | 1891.38 | 56 |
| H14B | 808.49 | 6581.47 | 2742.83 | 56 |
| H15A | 833.21 | 7710.64 | 2753.48 | 72 |
| H15B | 1785.34 | 7700.97 | 2372.88 | 72 |
| H16C | 1861.89 | 7912.08 | 3699.42 | 86 |
| H16D | 2400.31 | 7289.87 | 3477.34 | 86 |
| H16A | 1924.2 | 7907.82 | 3689.1 | 86 |
| H16B | 2367.09 | 7235.05 | 3504.63 | 86 |
| H17A | 1075.92 | 6714.07 | 3970.45 | 76 |
| H17B | 671.21 | 7389.18 | 4183.98 | 76 |
| H17C | 1200.04 | 6660.33 | 4035.41 | 127 |
| H17D | 826.53 | 7323.39 | 4377.46 | 127 |
| H18A | 1681.8 | 7503.65 | 5204.66 | 89 |
| H18B | 1083.51 | 6887.95 | 5312.51 | 89 |
| H18C | 2656.05 | 7009.35 | 4546.91 | 119 |
| H18D | 2128.57 | 7533.19 | 5036.19 | 119 |
| H19A | 2686.43 | 6813.66 | 5596.79 | 68 |
| H19B | 2788.89 | 6733.04 | 4683.43 | 68 |
| H19C | 1252.52 | 6603.45 | 5483.91 | 87 |
| H19D | 2166.98 | 6754.32 | 5892.92 | 87 |
| H20C | 2860.14 | 5953.36 | 5126.12 | 62 |
| H20D | 2100.96 | 5657.12 | 5649.79 | 62 |
| H20A | 2781.47 | 5721.31 | 5304.19 | 62 |
| H20B | 1832.06 | 5863.98 | 5653.34 | 62 |
| H23 | 472.13 | 5042.14 | 3306.88 | 49 |
| H25 | 1235.8 | 5615.36 | 2099.17 | 45 |
| H27A | -526.62 | 4680.76 | 4292.13 | 118 |
| H27B | -414.49 | 4255.7 | 5053.68 | 118 |
| H27C | 259.66 | 4179.49 | 4351.74 | 118 |

Atomic Occupancies for all atoms that are not fully occupied in **3**.

| Atom | Occupancy | Atom | Occupancy | Atom | Occupancy | Atom | Occupancy |
|---|---|---|---|---|---|---|---|
| H16C | 0.40(3) | C18 | 0.60(3) | H19C | 0.40(3) | C28 | 0.587(9) |
| H16D | 0.40(3) | H18A | 0.60(3) | H19D | 0.40(3) | C29A | 0.413(9) |
| H16A | 0.60(3) | H18B | 0.60(3) | H20C | 0.40(3) | F1A | 0.413(9) |
| H16B | 0.60(3) | C18A | 0.40(3) | H20D | 0.40(3) | F2A | 0.413(9) |
| C17 | 0.60(3) | H18C | 0.40(3) | H20A | 0.60(3) | F3A | 0.413(9) |
| H17A | 0.60(3) | H18D | 0.40(3) | H20B | 0.60(3) | C28A | 0.413(9) |
| H17B | 0.60(3) | C19 | 0.60(3) | C29 | 0.587(9) | | |
| C17A | 0.40(3) | H19A | 0.60(3) | F1 | 0.587(9) | | |
| H17C | 0.40(3) | H19B | 0.60(3) | F2 | 0.587(9) | | |
| H17D | 0.40(3) | C19A | 0.40(3) | F3 | 0.587(9) | | |

#### References

1. Adusumilli, R. & Mallick, P. Data conversion with ProteoWizard msConvert. in *Proteomics: Methods and Protocols* (eds. Comai, L., Katz, J. E. & Mallick, P.) 339–368 (Springer, New York, NY, 2017). doi:10.1007/978-1-4939-6747-6_23

2. Schmid, R. *et al.* Integrative analysis of multimodal mass spectrometry data in MZmine 3. *Nat. Biotechnol.* **41,** 447–449 (2023).

3. Wang, M. *et al.* Sharing and community curation of mass spectrometry data with Global Natural Products Social Molecular Networking. *Nat. Biotechnol.* **34,** 828–837 (2016).

4. Shannon, P. *et al.* Cytoscape: a software environment for integrated models of biomolecular interaction networks. *Genome Res.* **13,** 2498–2504 (2003).

5. Boonlarppradab, C., Kauffman, C. A., Jensen, P. R. & Fenical, W. Marineosins A and B, cytotoxic spiroaminals from a marine-derived actinomycete. *Org. Lett.* **10,** 5505–5508 (2008).

6. Bierman, M. *et al.* Plasmid cloning vectors for the conjugal transfer of DNA from Escherichia coli to Streptomyces spp. *Gene* **116,** 43–49 (1992).

7. Lian, W. *et al.* Genome-wide transcriptome analysis reveals that a pleiotropic antibiotic regulator, AfsS, modulates nutritional stress response in Streptomyces coelicolor A3(2). *BMC Genomics* **9,** 56 (2008).

8. Pereira, F. *et al.* Optimized production of concanamycins using a rational metabolic engineering strategy. *Metab. Eng.* **88,** 63–76 (2025).

9. Gilchrist, C. L. M. & Chooi, Y.-H. clinker & clustermap.js: automatic generation of gene cluster comparison figures. *Bioinformatics* **37,** 2473–2475 (2021).

10. Usón, I. & Sheldrick, G. M. An introduction to experimental phasing of macromolecules illustrated by SHELX; new autotracing features. *Acta Crystallogr. Sect. Struct. Biol.* **74,** 106–116 (2018).

11. Dolomanov, O. V., Bourhis, L. J., Gildea, R. J., Howard, J. A. K. & Puschmann, H. OLEX2: a complete structure solution, refinement and analysis program. *J. Appl. Crystallogr.* **42,** 339–341 (2009).

12. Sheldrick, G. M. Crystal structure refinement with SHELXL. *Acta Crystallogr. Sect. C Struct. Chem.* **71,** 3–8 (2015).

13. PX018 - CrysAlis$^{Pro}$: an all-in-one software package for single crystal X-ray diffraction. https://rigaku.com/products/crystallography/x-ray-diffraction/application-notes/px018-crysalispro-software-package-single-crystal.
